# Global Protein-Ligand Binding Affinity Profiling via Photocatalytic Labeling

**DOI:** 10.1101/2025.07.04.662444

**Authors:** Charles D. Warren, Noah Yardeny, Siyang Peng, Colin S. Burdette, Jacob B. Geri

## Abstract

Protein-ligand binding, selectivity, and affinity dictate the effects of drugs and endogenous molecules in cells. Currently, potential protein-ligand interactions are identified by qualitative interpretation of proteomic, transcriptomic, or genomic data, then binding affinities of hits are measured using purified proteins or engineered reporter systems to validate and quantify the strength of individual interactions. Few methods enable simultaneous target identification and biophysical affinity measurement, and these either apply to specific enzyme classes or proteins with ligand-dependent shifts in stability. Here we describe a general platform, termed Affinity Map, which leverages competitive binding analysis, high fidelity photocatalytic labeling, and high throughput proteomics for global quantitative binding affinity profiling. We show that this method is applicable to major classes of ligands, including small molecules, linear peptides, cyclic peptides, and proteins, and can measure affinities between unmodified ligands and proteins in cell lysates, organ extracts, and live cell surfaces.

## Introduction

Methods for measuring the strength and specificity of protein-ligand interactions are of central importance in biology. Ligands can interact with many different proteins, with aberrant interactions causing disease and undesired therapeutic side effects. Extensive profiles of binding affinity across the proteome are therefore required to understand ligand functions and binding selectivity, especially in drug development. Affinity measurements are typically performed separately for individual protein-ligand pairs, either using purified proteins, cell-based reporter protein systems, or competitive binding with a selective radioactive ligand.^1–5^ These methods can efficiently measure the binding affinities of many different molecules for a single protein target, making them indispensable in drug development. However, target-based assays are only available for a fraction of the proteome, preventing an accurate assessment of binding selectivity in complex biological systems (Fig. 1, top).

**Fig. 1.**
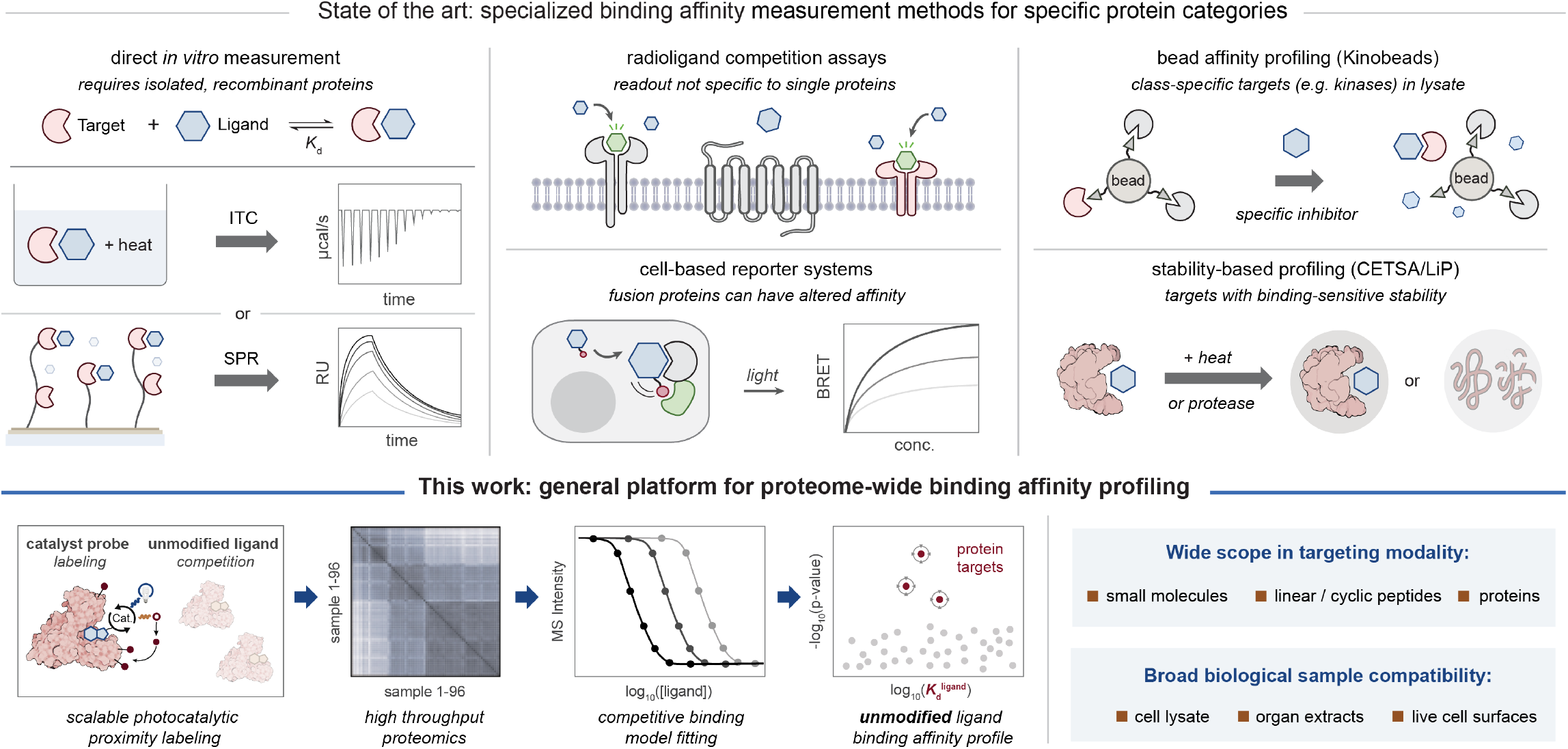
A high fidelity platform for simultaneous protein-ligand interaction identification and affinity measurement. Top: Current methods for measuring protein-ligand binding affinity use recombinant proteins, engineered cells, or provide incomplete coverage of interactions in the proteome. Bottom: Photocatalytic affinity labeling combined with a quantitative competitive ligand binding model enables robust, modality-agnostic *K*d profiling in lysate, organ extracts, and live cells.

Current approaches for proteome-wide affinity measurement rely on competitive binding to bead surfaces or solid support or shifts in protein stability as mass spectrometry (MS)-quantifiable readouts of protein binding site occupancy. Bead-based methods (e.g. Kinobeads)^6–11^ can offer affinity profiles for select categories of proteins such as kinases, but are not easily used with membrane proteins or live cells. While more general in scope, methods based on ligand-altered protein stability (cellular thermal shift assay, CETSA; limited proteolysis, LiP; peptide-centric local stability assay, PELSA) capture the subset of interactions which materially alter protein melting point (*ΔT*_m_) or susceptibility to proteolytic degradation.^12–20^ Specialized methods are also available for measuring DNA/RNA-protein binding affinities^21,22^ and profiling the kinetic reactivity of covalent molecules with nucleophilic amino acids,^23^ but these are not applicable to druglike noncovalent ligands. There remains an unmet need for a general affinity profiling platform equally applicable to all categories of noncovalent ligands, protein targets, and biological samples.

Photoaffinity labeling offers a potentially advantageous proteomic readout for affinity profiling. Like a radioligand binding assay,^24^ photocrosslinking works with membrane and non-membrane proteins, is compatible with live cells, and is not impacted by protein physical properties such as molecular weight or melting point.^25,26^ Recently, photocatalytic labeling platforms (e.g. µMap, Photag, Lux-MS, MultiMap, DarT)^27–32^ have been introduced that amplify crosslinking efficiency, leading to their widespread adoption. Despite their versatility, photolabeling methods have only been used to qualitatively nominate potential binding targets rather than quantitatively measure binding affinity.^33,34^ We hypothesized that labeling by photocatalytic probes could be leveraged to accurately measure the protein binding affinities (*K*_d_) of unmodified, non-photocatalytic ligands, enabling robust affinity profiling. If protein labeling is proportional to binding site occupancy by a photocatalyst-conjugated probe, treatment with a competing ligand would afford a dose-dependent reduction in labeling. The relationship between labeling signal and probe/ligand concentrations can therefore be modeled using standard competitive binding kinetics. Such a method would be akin to a global radioligand competition assay with a protein-level readout, enabling general *K*_d_ determination via MS measurement of the photocatalytically labeled proteome.

Here we report a new method, Affinity Map, which combines efficient labeling chemistry, high-throughput MS proteomics, and a tailored data processing pipeline to measure binding affinities between ligands and the proteome. To demonstrate the generality of this method, we show that it is compatible with major types of therapeutic protein-binding ligands (small molecules, linear peptides, cyclic peptides, and antibodies), is applicable to both membrane and non-membrane protein targets, and can measure binding affinities in cell lysate, organ extracts, and live cells (Fig. 1, bottom).

### Design concept and analysis pipeline

Accurate affinity profiling requires that the MS signal introduced by photocatalytic labeling is highly specific to direct ligand binding targets. We selected the µMap platform, in which highly reactive carbenes selectively biotinylate the direct binding targets of a photocatalyst-probe conjugate, to provide the MS readout used for modeling. We hypothesized that the MS signal from protein labeling would be proportional to binding site occupancy by the photocatalytic probe, such that competitive binding between the probe and an unmodified ligand would result in logistic dose-response curves where half of the probe would be displaced at a measured unmodified ligand concentration (EC_50_). The relationship between EC_50_, probe concentration, and *K*_d_ values for both the probe and ligand are given by the Cheng-Prusoff equation^35^ :

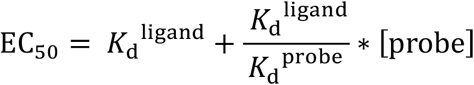

Using this relationship, linear regression can be used to obtain *K*_d_^ligand^ values by measuring EC_50_ at a series of probe concentrations. Notably, modification of a ligand to incorporate a bulky reporter moiety such as a photo-catalyst typically weakens its binding affinity. When the relative affinity of the probe is much weaker than the ligand, its effect on the EC_50_ value approaches zero, such that EC_50_ = *K*_d_^ligand^. Therefore, we expected that accurate *K*_d_^ligand^ measurement for the unmodified ligand would be possible even if the binding affinity of the probe is low, so long as sufficient MS signal from photocatalytic labeling is observed for a given protein.

Based on these principles, we designed a custom data processing pipeline to determine proteome-wide unmodified ligand affinities, which we named Affinity Surveyor (Fig. 2A, S8, detailed description in supplementary materials). After normalization and imputation, MS data for a specific protein is subjected to logistic curve fitting at each probe concentration. EC_50_ values from each curve are then used to fit the ligand binding affinity using the Cheng-Prusoff equation, and p-values for all logistic models are combined to give the aggregate significance of data supporting a *K*_d_^ligand^ measurement. For proteins with significant (p < 0.1) values for *K*_d_^ligand^*/K*_d_^probe^, corrected values of *K*_d_^ligand^ are reported. In cases where *K*_d_^ligand^*/K*_d_^probe^ is indistinguishable from zero (confidence interval < ±0.25) we report average EC_50_ values as *K*_d_^ligand^. For proteins with insufficient data to model *K*_d_^ligand^*/K*_d_^probe^, the EC_50_ measured at the lowest probe concentration is reported as a best estimate of *K*_d_^ligand^ because incorporation of a bulky photocatalyst often makes *K*_d_^probe^ much larger than *K*_d_^ligand^. This is repeated for all detected proteins in the MS dataset, providing a proteome-wide binding affinity profile.

**Fig. 2.**
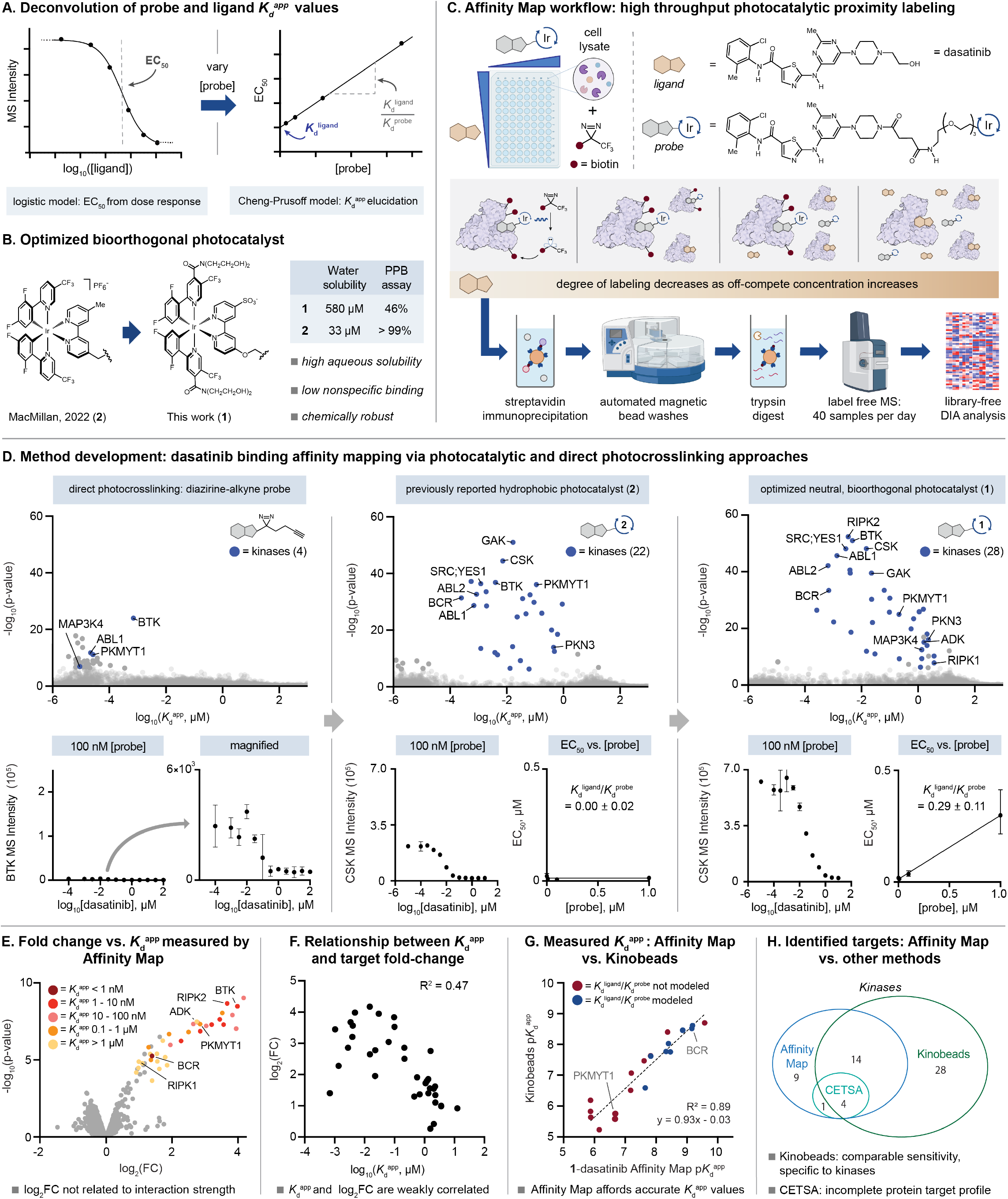
Experimental design and small molecule affinity profiling in lysate. **A)** EC_50_ values measured at each probe concentration are used to obtain the free ligand *K*_d_^app^ value. **B)** Neutral, hydrophilic photocatalyst **1** with low plasma protein binding (PPB). **C)** Affinity Map workflow in lysate. **D)** Comparison of dasatinib affinity profiles using various probes (0.01% false-discovery rate (FDR). **E)** Volcano plot showing enriched proteins in a two-group comparison (log_2_(fold change) (FC) = log_2_(1 µM **1**-dasatinib)-log_2_(1 µM **1**-dasatinib + 10 µM dasatinib) color-coded by measured binding affinity. **F)** Correlation of dasatinib-target log_2_(FC) vs. Affinity Map-measured binding affinity. **G)** Correlation of kinase binding affinities measured by Affinity Map using both Cheng-Prusoff modeled (*K*_d_^ligand^/*K*_d_^probe^ value significant, p < 0.1, or confidence interval < ±0.25) and non-modeled values vs. Kinobeads. **H)** Dasatinib kinase targets identified by Affinity Map, Kinobeads, and CETSA.

### Catalyst development

We developed an optimized bioorthogonal photocatalyst, **1**, for the Affinity Map workflow (Fig. 2B). We first attempted to use previously reported photocatalyst **2**^33,34^ to assess competitive binding of small molecules. However, Ir(dF(CF_3_)ppy)_2_(dmbpy) based catalysts are hydrophobic, leading to nonspecific protein binding, off-target labeling, and low water solubility. Reported catalysts^27,36^ containing carboxylic acid moieties are more hydrophilic, but are challenging to synthesize and use on large scales. Photocatalyst **1** has high aqueous solubility, decreased nonspecific interactions with the proteome by mass spectrometry and plasma protein binding (PPB), no net charge, and robust bench stability (Fig. S1-S5).

### Small molecule-protein affinity profiling

Dasatinib was selected as a model ligand for initial experiments due to its large number of reported kinase binding affinities. To provide sufficient statistical power for useful affinity profiling, we designed an experimental matrix in which twelve ligand concentrations (vehicle, 100 pM-10 µM) were used at four photocatalytic probe (**1**-dasatinib) concentrations (1 nM-1 µM in duplicate, 96 samples, Fig. S19), with a 60-minute incubation time. To facilitate large scale experiments and increase sample-to-sample consistency, we developed a high throughput, 96-well plate format, two-day protocol for parallel photocatalytic labeling driven by 440 nm light, streptavidin enrichment, tryptic digestion, and preparation for MS analysis (Fig. 2C, S18). MS data acquisition (timsTOF Pro 2) and protein quantification (DIA-NN, *in silico* library) generally required three days per affinity profile. After protein quantification, the data was subjected to the Affinity Surveyor pipeline to determine dasatinib targets. Notably, as binding may not have reached equilibrium for some targets and target concentrations are unknown, all measured *K*_d_ values are reported as apparent affinities (*K*_d_^app^) (Fig. S22).

Using a photocatalytic **1**-dasatinib conjugate and the Affinity Map workflow in K562 lysate, we unambiguously measured *K*_d_^app^ for dasatinib against 28 kinases and 8 other proteins (many of which contain nucleotide binding sites) (Fig. 2D). Qualitative inspection of data collected for target proteins such as CSK and BTK revealed clear sigmoidal dose response curves at all **1**-dasatinib probe concentrations (Fig. S10). We also compared the performance of **1**-dasatinib, **2**-dasatinib, and a dasatinib-diazirine-alkyne conjugate as reporter probes for affinity profiling. Using the more hydrophobic photocatalyst **2**, we observed fewer kinases with measurable affinity values. Inspection of dose-response curves reveals lower on-target labeling efficiency at low catalyst concentrations and higher background signal at higher catalyst concentrations, reflecting labeling caused by nonspecific catalyst binding. In contrast, activation of a non-photocatalytic dasatinib-diazirine probe with 375 nm light led to far less efficient protein labeling than observed with **1**-dasatinib, enabling measurement of only four kinase affinities.

Previously reported photoaffinity and photocatalytic labeling methods compare protein crosslinking by a probe against a control, with the most differentially enriched proteins flagged as potential binding targets. We evaluated whether differential enrichment from protein labeling is correlated with affinity. Treating K562 lysate with **1**-dasatinib and either vehicle control or 10-fold excess dasatinib, we found that the difference in enrichment was not well-correlated with affinity (Fig. 2E-F, S12). Indeed, many proteins with high affinity to dasatinib, including the well-characterized target BCR-ABL, have relatively low fold change due to variable and protein-specific MS intensity background signal. Proteins can have significant fold change but lack competitive binding curves, potentially representing false-positive hits caused by nonspecific binding, which are indistinguishable from real targets in standard two-group comparison but easily identified by logistic modeling in Affinity Map. *K*_d_^app^ values obtained using Affinity Map were close to literature Kinobead and *in vitro* affinities (Fig. 2G, S15),^6,7,37^ including both values where the *K*_d_^ligand^*/K*_d_^probe^ is fully modeled and well constrained or estimated due to a lack of sufficient data for Cheng-Prusoff modeling. Notably, we measure *K*_d_^app^ values for more kinase targets than were identified in a recently reported CETSA experiment performed in K562 lysate, which identified only five kinases with measurable *Δ*T_*m*_ values despite MS detection of many kinases that were successfully profiled by Affinity Map (Fig. 2H, S15). One protein identified by CETSA, ADK,^13^ lacked a known affinity for dasatinib. We confirmed this interaction using Affinity Map, measuring an affinity of 1.6 µM. We observed three kinase targets via Affinity Map (PKN3, CSNK1D, and RIPK1) that to our knowledge were previously unknown. We therefore used an *in vitro* activity assay (PhosphoSens) to validate these interactions. Two kinases gave values very similar to those measured via Affinity Map (RIPK1, 3.0 µM vs. 3.6 µM; CSNK1D, 4.1 µM vs. 3.6 µM), while PKN3 was assayed within approximately one log of the Affinity Map value (Fig. 3A, S16).

**Fig. 3.**
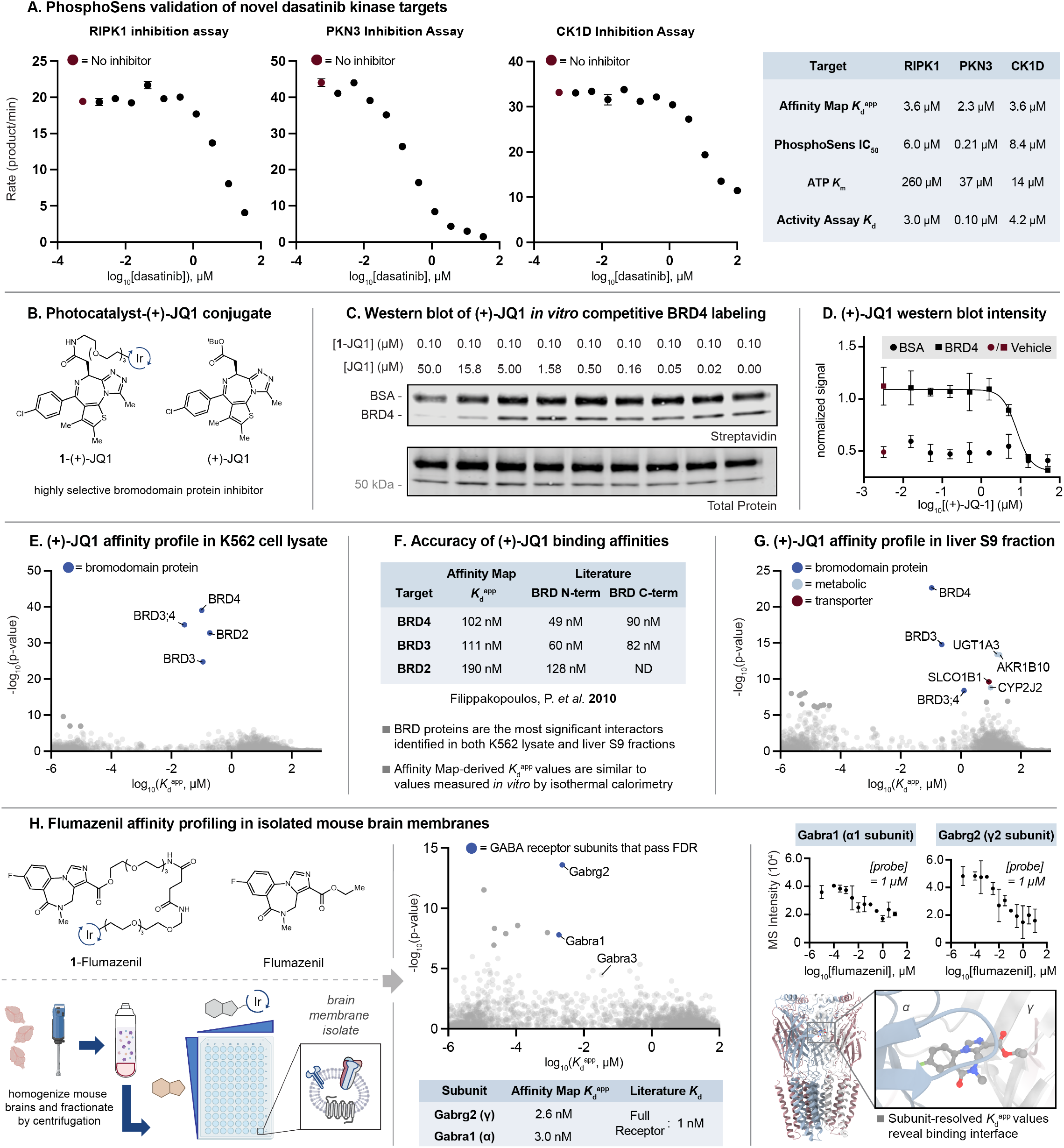
Affinity profiling of (+)-JQ-1, a selective bromodomain-containing protein inhibitor. **A)** Validation of newly identified dasatinib binding targets RIPK1, PKN3, and CK1D and binding affinity via *in vitro* activity assays (PhosphoSens) at [ATP] = K_m_. **B)** Chemical structures of both **1**-(+)-JQ1 and unmodified (+)-JQ1. **C)** Western blot showing a [(+)-JQ1]-dependent reduction in BRD4 biotinylation by **1**-(+)-JQ1. **D)** Normalized streptavidin intensity (relative to total protein) vs. [(+)-JQ1]. **E)** (+)-JQ1 affinity profile in K562 lysate reflects selective binding to BRD proteins. **F)** Comparison of protein binding affinities measured using Affinity Map vs. isothermal calorimetry. **G)** (+)-JQ1 Affinity profile in pooled human liver S9 fraction. **H)** Chemical structures of both **1**-(+)-Flumazenil and unmodified Flumazenil, the workflow for mouse brain membrane isolation, flumazenil affinity profile in mouse brain membrane isolates, and sigmoidal curve fits for the α/γ GABA receptor subunits showing labeling of the flumazenil binding interface (PDB: 6X3U). Affinity profiles shown at 0.01% FDR.

To test the false positive discovery rate of Affinity Map, we profiled a highly selective small molecule, (+)-JQ1 (Fig. 3B), which is an inhibitor of bromodomain containing proteins BRD2, BRD3, and BRD4. Using (+)-JQ1 and **1**-(+)-JQ1, we measured competitive labeling at the intact protein level in the presence of BSA as a nonspecific binding control. We observed a clear sigmoidal dose-response curve by streptavidin western blot (Fig. 3C-D, S7), further validating our logistic model of photocatalytic protein biotinylation. In K562 cell lysate, Affinity Map accurately captures the selectivity of this compound (Fig. 3E), exclusively reporting measurable binding affinities for these three proteins. Measured *K*_d_^app^ values are close to previously reported values obtained by isothermal calorimetry with purified BRD domains (Fig. 3F).^38^

We next evaluated the applicability of the Affinity Map workflow to protein extracts from human organs (Fig. 3G). Liver microsomes generated through mechanical organ disruption contain metabolic enzymes and are commonly used to evaluate the stability of small molecules. We performed (+)-JQ1 targeted Affinity Map using pooled human liver S9 fraction, a minimally processed homogenate containing microsomes and globular proteins. We measured *K*_d_^app^ values for BRD3, BRD4, AKR1B10 (a cytosolic aldo-keto reductase), in addition to microsome constituents CYP2J2 (a membrane-bound P450 enzyme) and UGT1A3 (a transmembrane glucuronosyltransferase). We further measured (+)-JQ1 affinity to SLCO1B1, a liver-specific plasma membrane organic ion transporter. These data highlight the potential utility of Affinity Map for profiling diverse sectors of the proteome directly in patient samples.

To further demonstrate the utility of Affinity Map in organ extracts, we profiled the interactions of flumazenil, a benzodiazepine antagonist, in mouse brain membrane isolates, prepared according to radioligand binding assay protocols where endogenous GABA is washed out (Fig. 3H).^39^ Flumazenil binds heteropentameric GABA receptors at the α and γ subunit interface, known as the benzodiazepine site. In our experiment, significant sigmoidal curve fits and accurate *K*_d_^app^ values^40^ were measured for two GABA receptor subunits, Gabrg2 (γ2 subunit) and Gabra1 (α1 subunit). A single sigmoid was measured for the α3 subunit but did not pass FDR likely due to its low MS intensity. Detection of other subunits (β1−3 and α2/4/6) that lack sigmoidal dose-response curves demonstrates subunit resolved affinity measurements directly at the α/γ binding interface. Notably, the pentameric form of the GABA receptor binding flumazenil in whole mouse brain is composed primarily of α1/3 subunits, consistent with previous literature.^41^ We anticipate that Affinity Map in brain membrane isolates will be instrumental for profiling other neuroactive ligand-receptor interactions and their affinities.

### Peptide-protein affinity profiling

Beyond drug-like small molecules, we next applied Affinity Map to profile the binding affinities of endogenous linear peptides and minimal binding motifs in proteins. We prepared photocatalyst-peptide probes by reacting an iodoacetamide derivative of **1** and peptides bearing an appended N or C-terminal cysteine residue (Fig. 4A), and profiled affinities for each peptide in K562 cell lysate. We began by applying Affinity Map to a fragment of Smac, a mitochondrial protein that promotes cell death when present in the cytosol by binding inhibitor of apoptosis (IAP) proteins, leading to the release of active caspase enzymes. Using a 9 amino acid peptide (Fig. 4B) as a minimal IAP-binding motif from Smac, Affinity Map identified IAP proteins XIAP and BIRC2 as the most significant interactions with affinities of 3.3 µM and 2.0 µM respectively. The measured binding affinities were comparable to reported data obtained *in vitro* with purified BIR domains and the same peptide (XIAP BIR3/BIR2 domains: 0.43 µM/ 6.0 µM).^42^ As our workflow uses protein level enrichment, the obtained affinities represent an ensemble measurement of all peptide-BIR domain interactions within XIAP and BIRC2, each of which contain 3 unique BIR domains with different affinities for Smac.^42^ We did not observe other IAPs, such as BIRC6, which require multi-valent interactions with full-length Smac for binding.^43^ Additionally, we performed Affinity Map for a 14 amino acid minimal binding motif from the protein SOS1 that binds the SH3 domain of GRB2.^44,45^ We identify GRB2 (Fig. 4C) as a low affinity (5.6 µM) interactor of this peptide, consistent with a previous affinity measurement of 4 µM.^44^

**Fig. 4.**
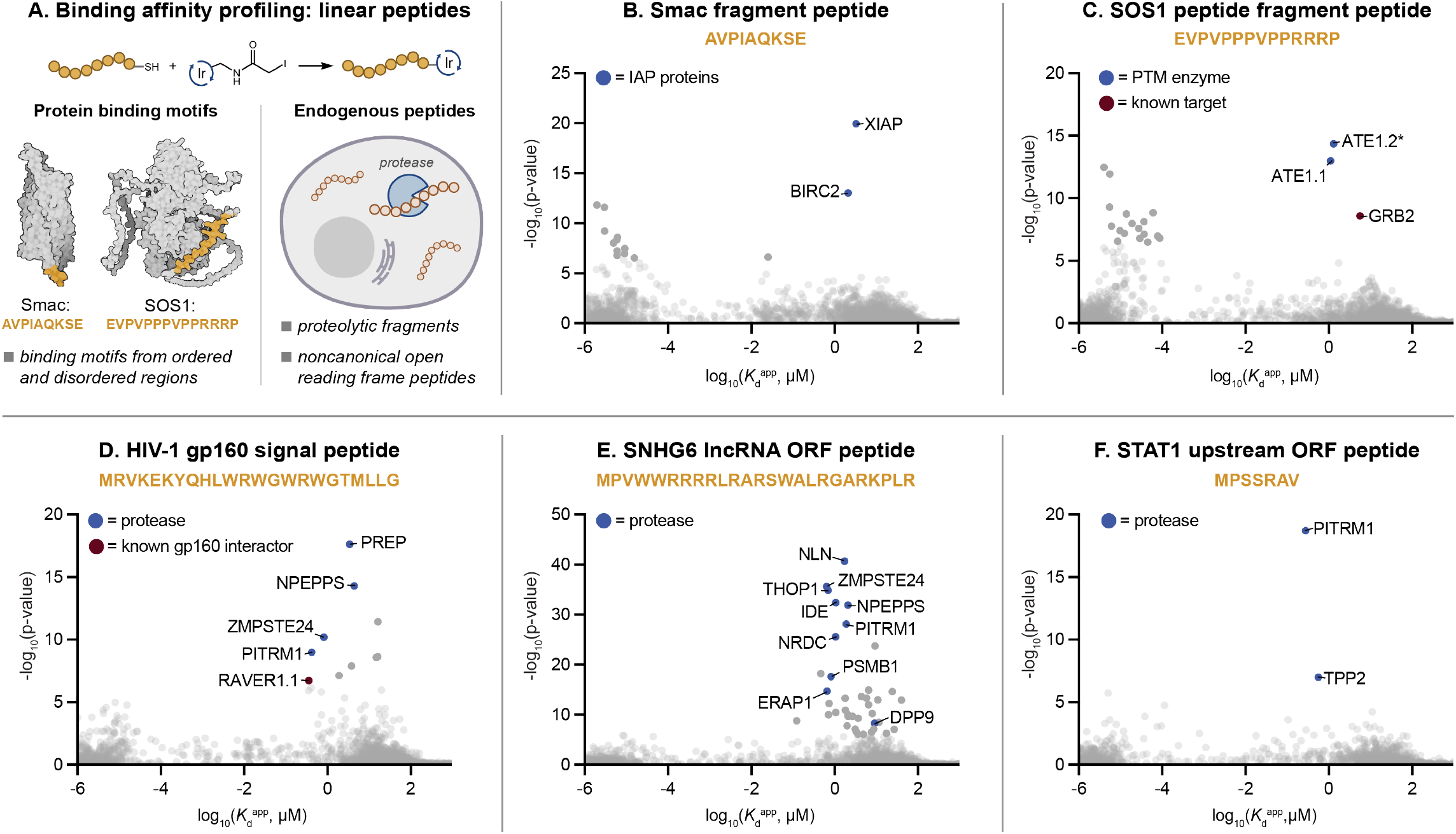
Linear peptide affinity mapping in K562 lysate. **A)** Probe molecules were synthesized by reacting an iodoacetamide-**1** derivative with peptides containing an appended N- or C-terminal cysteine. Peptides selected for analysis were either endogenous (ncORF, proteolytic fragment) or derived from ordered and disordered regions of proteins. Smac pdb: 1FEW; SOS1 structure from AlphaFold. **B)** An IAP-binding peptide fragment from Smac was profiled using Affinity Map. **C)** The affinity of the primary interactor of the SOS1-derived peptide was also measured by Affinity Map. **D-F)** Affinity profiles for endogenous peptides (derived from HIV-1 gp160 signal peptide, SNHG6 lncRNA ORF, and STAT upstream ORF). *ATE1.2: protein group comprising peptides matched to both ATE1 and the ATE1-2 sequence variant. Affinity profiles shown at 0.01% FDR.

We also profiled the binding affinities of several endogenous peptides (Fig. 4D-F): 1) a signal peptide fragment cleaved from the nascent HIV-1 gp160 envelope protein during infection,^46^ 2) a peptide translated from a non-canonical open reading frame (ncORF) from the SNHG6 long noncoding RNA (lncRNA),^47^ and 3) a translated peptide from an upstream ORF (uORF) of the STAT1 gene.^45^ We measure an affinity for a known interactor of the HIV-1 envelope protein^48^ and also measure the affinities of these three peptides to proteases that likely mediate their degradation. Unique proteases were identified for each peptide, consistent with their known cleavage sites (Fig. S17). As an example, the arginine rich peptide from the SNHG6 lncRNA ORF binds NRDC protease, which cleaves sites containing dibasic motifs.^49^ This demonstrates the ability of Affinity Map to identify proteases relevant to the stability of endogenous peptides.

### Protein-protein and cyclic peptide-protein affinity measurement on live cell surfaces

We hypothesized that Affinity Map would be compatible with live cells, enabling the measurement of ligand binding affinities for cell surface proteins in their native environment. Initially, we sought to demonstrate affinity profiling on live cell surfaces using cyclic peptides, an emerging category of therapeutics and tool compounds. The macrocyclic cRGDfK peptide is a high affinity integrin alpha-v beta-3 (αV*Δ*3) antagonist that has been explored as a drug scaffold for glioma treatment.^50^ *In vitro* assays for integrin-RGD peptide binding affinity and selectivity require purified integrins, and reported measurements using surface adhesion to live cells cannot discriminate between different integrin affinities.^51,52^ We prepared a photocatalytic conjugate of cRGDfK by functionalizing the lysine residue with **1** via an amide linkage and treated live U87MG cells adhered to 6-well culture plates (16 plates, 96 wells) with 4 different probe concentrations and 12 concentrations of unmodified cRGDfK in complete media (Fig. 5A). Following irradiation, the cells were harvested and subjected to the Affinity Map workflow. Integrin β3 (ITGB3) had the most significant measurable binding affinity, with clear sigmoidal competition curves at multiple probe concentrations. These data represent the first measurement of cRGDfK selectivity and binding affinity (268 nM) in live cells.

**Fig. 5.**
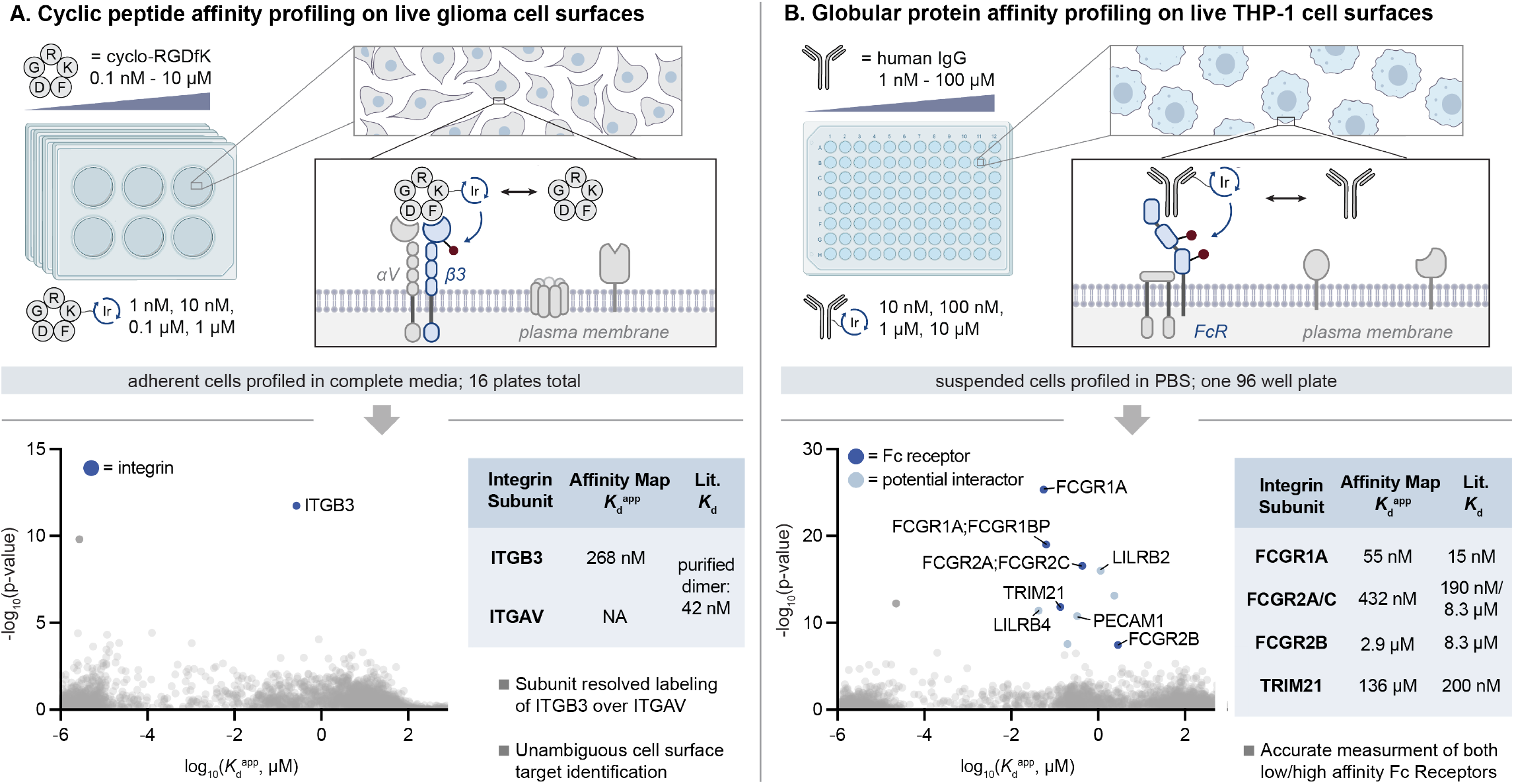
Affinity Map for a cyclic RGD peptide and IgG on live cell surfaces. **A)** Left: cyclo-RGDfK affinity profiling in live U87MG glioma cells was performed in 24 6-well plates, allowing for simultaneous affinity measurement and selectivity assessment on the surfaces of living, adherent cells in complete media. Right: Affinity Map measures binding to integrin β3 (ITGB3), consistent with known cyclo-RGDfK selectivity for the αVβ3 heterodimer. **B)** Left: IgG affinity profiling on live THP-1 cell surfaces was performed with suspended cells using a single 96 cluster-tube plate. Right: the affinity profile shows IgG affinities for several Fc-receptors. All plots shown are at 0.01% FDR.

Beyond small molecules, carbene-based photocatalytic labeling methods have been previously used to identify the binding targets of proteins such as antibodies and the SARS-COV-2 spike protein on live cell surfaces.^27,36^ In these experiments, labeling by photocatalyst-conjugated proteins is contrasted with a control lacking a photocatalyst targeting modality. We reasoned that the Affinity Map platform could be generalized to profile the binding affinity of globular proteins to proteins on live cell surfaces. Currently, protein-protein affinities are measured using cell-based reporter protein systems or recombinant protein assays, and no published methods are available for proteome-wide protein-protein binding affinity measurement.

As a proof of concept, we aimed to profile the binding affinities between human IgG and proteins on live THP-1 cell surfaces (Fig. 5B). We synthesized **1**-IgG as a photocatalytic protein probe using purified IgG from human serum and an activated NHS ester prepared through a strain-promoted click ligation between **1**-DBCO and NHS-PEG_3_-N_3_ and used unmodified IgG as a competitive non-photocatalytic ligand. We then incubated live cells with a 96-sample matrix of **1**-IgG / IgG concentrations (Fig. S21), irradiated the mixtures for 15 minutes, lysed the cells, and subjected the lysate to the standard Affinity Map sample preparation and data processing pipeline. We measured binding affinities closely matching literature values for the major IgG receptors expressed by THP-1 cells, including FCGR1A (55 nM; lit. 15 nM), FCGRIIB (2.9 µM; lit. 8.3 µM) and a set of peptides matching to both FCGRIIA and FCGRIIC (432 nM; lit. 190 nM/8.3 µM).^53^ We also measured the IgG affinity of TRIM21, an intracellular Fc receptor (136 nM; lit. ∼200 nM),^54^ and identified additional proteins that are not known to directly bind IgG, such as the monocyte checkpoint inhibitor proteins LILRB2 and LILRB4 at lower significance. Notably these two proteins have closely aligned affinity values with Fc receptors, suggesting that they may represent distal Fc receptor interactions. Together, these data validate the ability of Affinity Map to measure the affinities between globular proteins and cell surface receptors, which are important metrics in the development of therapeutic anti-bodies.

## Discussion

Affinity Map will potentially enable improved measurement of polypharmacology at critical decision points in drug development, identifying off-target interactions which may contribute to toxicity and undesired side effects. Many such interactions cannot be detected with existing affinity profiling methods due to incompatibility between specific protein physical properties and assay requirements. Hits from qualitative target-ID experiments can either lack activity-based assays or can be challenging to isolate, preventing direct affinity measurement using *in vitro* approaches. Affinity Map bypasses these limitations by offering a completely target-agnostic binding affinity measurement platform. Notably, Affinity Map requires the synthesis of a photocatalytic probe for competition experiments with the corresponding unmodified ligand of interest. Incorporation of the photocatalyst can disrupt target binding, requiring assessment of structure-activity relationships in probe design, and some molecules cannot be easily modified; these represent key limitations of all label-based methods in chemical biology. We view Affinity Map as a complementary approach to label-free methods such as CETSA, which do not require probe molecules as competitive ligands but can fail to identify target proteins if they lack significant ligand-induced stability shifts.

In developing Affinity Map, we also introduce the first scalable photocatalytic proximity labeling workflow (>1,200 samples were used to generate data in Figs. 2-5), enabling high throughput applications in target identification for drug-like molecules. We expect that two-dimensional competitive binding assays with MS-proteomic readout will translate to other labeling methods beyond µMap and become a standard, first-line strategy for simultaneous target identification and affinity measurement as the throughput and accessibility of proteomic measurement methods continue to improve.

## Acknowledgements

The authors thank the Sanders Tri-Institutional Therapeutics Discovery Institute for supplying chemical reagents and its members for fruitful discussion.

## Author Contributions

J.B.G. supervised the work. J.B.G., N.Y., and C.D.W. developed the theoretical model. J.B.G. and S.P. wrote the Affinity Surveyor code. C.D.W. designed the Affinity Map experimental pipeline. N.Y. designed and synthesized molecules. C.S.B. characterized photocatalyst activity. C.D.W., N.Y., and S.P. performed Affinity Map experiments. J.B.G. and C.D.W. wrote the manuscript with input from all authors.

## Methods

These methods outline the mass spectrometry-based proteomic workflows and data analysis pipeline (Affinity Surveyor) used to generate Affinity Map-related data. An exhaustive description of methods, including detailed methods for chemical syntheses of all compounds and details concerning materials and reagents (*e*.*g*., suppliers, part no.), additional methods (e.g. cell culture techniques), and additional MS-based proteomics information are available in Supplementary Information (SI).

### Pooled cell lysate preparation

For a single 96-sample experiment, 2.2 x 10^8^ K562 cells were collected by centrifugation (800 x g, 5 min, 20 °C), and washed twice with 40 mL of cold PBS. Cells were resuspended in 10.6 mL cold (4 °C) PBS in a 50 mL conical tube and lysed by sonication (QSonica Q500 with cup horn, 4 °C, 3 cycles of 60% amplitude, 4 seconds on/off for 64 seconds). Lysate was examined under a brightfield microscope (1:1 lysate:trypan blue 0.4%) to ensure complete lysis. The lysate was cleared by centrifugation (2,500 x g, 15 min, 4 °C). Concentrated diazrine-PEG3-biotin (Dz-Bt) dissolved in cold PBS (600 *μ*M, 7.04 mL) was added to the cleared lysate and diluted with cold PBS to 21.1 mL to give a final concentration of 200*μ*M Dz-Bt and a 2 mg/mL protein concentration, determined by bicinchoninic acid (BCA) assay. Lysate was kept fresh on ice and was not frozen.

### *In vitro* lysate treatment with photocatalyst-conjugated small molecules/peptides and free small molecules/peptides

200 *μ*L (400 *μ*g protein) of pooled K562 cell lysate was aliquoted into 12 8-strip cluster tubes (96 tubes) in a rack and kept on ice. Preprepared 96-well plates (compound plates, **Fig. S19**) containing 3 *μ*L of a DMSO solution containing appropriate concentrations of compounds (or vehicle DMSO) were used for 1) free small molecule addition and 2) probe addition. 1) All wells in the first compound plate (free small molecules) were diluted with 27 *μ*L room temperature PBS and then 20 *μ*L of each well was added to the correspond cluster tube containing 200 *μ*L of lysate (120-fold dilution) and incubated on ice for 20 minutes. 2) All wells in the second compound plate (probe) were diluted with 34.5 *μ*L room temperature PBS and 20 *μ*L of each well was added to the correspond cluster tube containing 200 *μ*L of lysate (150-fold dilution) and incubated on ice for 20 minutes (final volume 240 *μ*L, final concentrations small molecule/probe tabulated below). The plate setup presented here, and the concentration range used for most molecules in this manuscript (0.1 nM-10 µM ligand, 1 nM – 1 µM probe) is designed to capture affinities in the 1 nM-10 µM range. Based on predicted ligand binding regime a shifted or extended concentration range could be used. For example, for ligands with weaker affinities a higher concentration regime can be used (100 nM-1 mM range for ligand, 1 µM-100 µM for probe). In these experiments, ligand concentration may be increased as high as solubility allows. For the IgG and dasatinib-diazirine-alkyne experiments (**Fig. S21**) concentrations were increased by an order of magnitude in this way. Samples were then argon degassed, irradiated with 440 nm light, and protein was precipitated as detailed in the methods below.

### Preparation of liver S9 fraction for (+)-JQ1 Affinity Map

Frozen, mixed gender pooled liver S9 fractions were purchased from BioIVT (X008011). The purchased samples contained 40 mg of protein total, which was diluted to ∼2 mg/mL with PBS containing 200 *μ*M diazrine-PEG3-biotin (Dz-Bt) and was aliquoted across 96 cluster tubes (200 *μ*L homogenate, 400 *μ*g protein /cluster tube). For compound treatment, the *in vitro* lysate treatment method above was followed (plate layout in **Fig. S19**).

### Live U87MG cell treatment with 1-cRGDfK and free cRGDfK and labeling with 440 nm light irradiation

15 cm plates with U87MG cells were washed once with DPBS (10 mL), detached by incubation with TrypLE™ Express Enzyme (Gibco, 5 mL, 5 min, 37 °C), and collected by centrifugation (800 x g, 5 min, 20 °C). Cells were resuspended in DMEM 1640 complete medium (Gibco) supplemented with 10% fetal bovine serum (Gibco) and penicillin-streptomycin (100 U/mL final concentration, Gibco) and passed through cell strainers to help dissociate clumping cells. 4 x 6-well treated cell culture plates were seeded with 5 x 105 U87MG cells and allowed to grow until confluent (∼3 days). The media was aspirated and 400 *μ*L of phenol red-free media containing 200 *μ*M diazrine-PEG3-biotin (Dz-Bt) was pipetted over the adherent cells in each well. The appropriate concentration of unconjugated cRGDfK was added (40 *μ*L) or a vehicle control to each well and cells incubated on ice for 20 minutes. Cells were then treated with the appropriate concentration of **1**-cRGDfK (40 *μ*L) and incubated on ice for 20 minutes (final volume 240 *μ*L, 160 *μ*M Dz-Bt, final concentrations free cRGDfK/probe tabulated in **SI Method 16**). Each 6-well plate was then irradiated for 15 minutes at 4 °C. Following irradiation, cells were released from the plate with 1 mL of cold PBS, transferred to a 1.5 mL tube, and pelleted by centrifugation (800 x g, 5 min, 4 °C). The supernatant was aspirated, and cell pellets were frozen at −80 °C. This experiment was repeated for each concentration of **1**-cRGDfK (1 *μ*M, 100 nM, 10 nM, or 1 nM), thus four separate groups of four plates were seeded then treated on separate days (4 concentrations x 4 plates x 6 wells = 96 wells total).

### U87MG cell lysis, reduction, alkylation, and streptavidin enrichment

Cell pellets were thawed for 1 hour at 4 °C before resuspension in RIPA lysis buffer containing 1% SDS and protease inhibitor cocktail (200 *μ*L). Cells were incubated in RIPA buffer for 30 minutes at room temperature followed by sonication (QSonica Q500 with cup horn, 4 °C, 4 cycles of 60% amplitude, 4 seconds on/off for 64 seconds). Immediately following lysis, insoluble material was pelleted (800 x g, 5 min, 4 °C) and the supernatant was transferred to cluster tubes. Dithiothreitol (DTT) in water (50 μL, 50 mM stock, 10 mM final concentration) was added and the mixture incubated at 95 °C for 10 minutes. After the mixture was allowed to cool to room temperature, iodoacetamide in water (50 μL, 120 mM stock, 20 mM final concentration) was added and incubated at in the dark at room temperature for 30 minutes. Samples were centrifuged once more (2,500 x g, 10 min), to ensure any insoluble material was removed, transferred to a 0.5 mL/well 96-well plate, and 50 μL of Sera-MagTM Medium Capacity Streptavidin beads (Cytiva, 30152104010350) were added to each sample. The plate was sealed with a silicone 96-well plate sealing mat (VWR, 76311-636) and incubated overnight with continuous inversion. Sample were then subject to automated streptavidin bead washes detailed below.

### Treatment of live THP-1 cells with 1-IgG and free IgG, labeling with 440 nm light irradiation

2.1 x 108 THP-1 cells were collected by centrifugation (800 x g, 5 min, 20 °C), and washed twice with 40 mL of cold PBS. Cells were resuspended in concentrated diazrine-PEG3-biotin (Dz-Bt) dissolved in cold PBS (570 *μ*M, 2.2 mL) and aliquoted into 12 8-strip cluster tubes (96 tubes) in a rack and kept on ice (21 *μ*L / aliquot, approximately 400 *μ*g of protein after lysis). The appropriate concentration of free IgG was added (29 *μ*L) or a vehicle PBS control to each cluster tube and incubated on ice for 20 minutes. Cells were then treated with the appropriate concentration of Ir-IgG (10 *μ*L / aliquot) and incubated on ice for 20 minutes (final volume 60 *μ*L, 200 *μ*M Dz-Bt, final concentrations free IgG/probe tabulated in **Fig. S21**). Samples were then irradiated for 15 minutes at 4 °C. Following irradiation, 200 *μ*L of cold PBS was added to each cluster tube and cells were pelleted by centrifugation (800 x g, 5 min, 4 °C). The supernatant was aspirated, and cells were pelleted and washed with another 200 *μ*L of cold PBS. Cell pellets were frozen at −80 °C prior to lysis.

### THP-1 cell lysis

Cell pellets were thawed for 1 hour at 4 °C before resuspension in RIPA lysis buffer containing 1% SDS and protease inhibitor cocktail (100 *μ*L). Cells were incubated in RIPA buffer for 30 minutes at room temperature followed by sonication (QSonica Q500 with cup horn, 4 °C, 4 cycles of 60% amplitude, 4 seconds on/off for 64 seconds). Immediately following lysis, 140 *μ*L of additional H_2_O was added to each cluster tube (final volume 240 *μ*L) and MeOH/ CHCl_3_ precipitation was carried out according to the protein precipitation method below.

### Pooled membrane preparation from whole mouse brain (Flumazenil Affinity Map)

Based on a modified literature procedure,^39^ 3 male CD-1 (ICR) mouse brains (BioIVT, MSE00BRAIN-0102904) were thawed on ice for 25 minutes then resuspended in 10 volumes ice cold lysis buffer (50 mM Tris-HCl pH 7.4, containing protease inhibitor cocktail - Roche, 11836170001, approximately 5 mL per brain). The organs were homogenized (Bio-Gen PRO200 Homogenizer) on ice using at medium intensity (6 pulses x 10 seconds). As an initial clearing step, the homogenate was centrifuged (1,000 x g for 10 min at 4 °C) to obtain supernatant. The supernatant was transferred to ultracentrifuge tubes and centrifuged (40,000 x g for 20 minutes at 4 °C; Beckman SW 32 Ti swinging bucket rotor). The supernatant was decanted and discarded and replaced with fresh ice-cold lysis buffer. The pellet was resuspended by homogenization (5 pulses x 10 seconds) and the centrifugation-homogenization was repeated 2 additional times for a total of 3 ultracentrifugations to ensure complete homogenization and wash out of endogenous ligands (specifically GABA). The final pellet was resuspended in a minimal amount of ice-cold protease-inhibitor-free lysis buffer and protein concentration was determined by bicinchoninic acid (BCA) assay. The freshly prepared membrane suspension was never frozen and used as prepared. Concentrated diazrine-PEG_3_-biotin (Dz-Bt) dissolved in cold Tris-HCl was added for a final concentration of 200 µM and a 1 mg/mL protein concentration. The membrane suspension was then aliquoted across 96 cluster tubes (200 µL, 200 µg per sample). For compound treatment, the *in vitro* lysate treatment method above was followed (plate layout in **Fig. S19**).

### Dasatinib-diazirine-alkyne labeling

K562 cell lysate was prepared according to Method 1. Lysate (0.2 mL, 2 mg/mL) was transferred to cluster tubes and treated with compounds (two experiments detailed in **SI Method 13 and Fig. S9)** and irradiated using the same apparatus as Fig. S20 with 100-watt 375 nm LEDs for 5 minutes. 34 µL of 7x protease inhibitor cocktail (Roche, 11836170001) was then added, and biotin-PEG4-azide (2.74 µL, 10 mM in DMSO), CuSO4 (1.37 µL, 50 mM in H_2_O), THPTA (6.85 µL of 10 mM in H_2_O), and sodium ascorbate (13.7 µL, 50 mM in H_2_O, freshly made) added, in that order. Note that order of addition is important, adding the first three reagents together as a cocktail and sodium ascorbate last. Samples were inverted end over end for 1 hour at room temperature. Then 280 *μ*L of −20 °C MeOH and 70 *μ*L of CHCl_3_ were added to each cluster tube and mixed, precipitating denatured protein. Samples were centrifuged (2,500 x g, 10 min, 4 °C), affording a protein disk suspended between the MeOH and CHCl_3_ layers. Solvent was carefully aspirated without disrupting the protein disk, then 500 *μ*L of −20 °C MeOH was added to each cluster tube and the protein disk disrupted via sonication (Qsonica Q500 with cup horn, 4 °C, 3 cycles of 60% Amplitude, 4 sec on/off). Samples were centrifuged (2,500 x g, 10 min, 4 °C), resulting in a protein pellet at the bottom of each cluster tube. The MeOH was aspirated, and protein pellets were stored at −80 °C overnight then processed by the protein pellet redissolving, reduction, alkylation, and streptavidin bead enrichment method below.

### Argon degassing, 440 nm light irradiation, and protein precipitation

Samples were placed in a vessel that was sealed and purged with water-saturated argon (**Fig. S20**) for 15 minutes. Samples were then irradiated using the apparatus shown below. This setup was shown to achieve uniform irradiation of 96 cluster tubes using ferrioxalate-based actinometry (**Fig. S20**). Immediately following irradiation, 240 *μ*L of −20 °C MeOH and 60 *μ*L of CHCl_3_ were added to each cluster tube and mixed, precipitating denatured protein. Samples were centrifuged (2,500 x g, 10 min, 4 °C), affording a protein disk suspended between the MeOH and CHCl_3_ layers. Solvent was carefully aspirated without disrupting the protein disk, then 500 *μ*L of −20 °C MeOH was added to each cluster tube and the protein disk disrupted via sonication (Qsonica Q500 with cup horn, 4 °C, 3 cycles of 60% Amplitude, 4 sec on/off). Samples were centrifuged (2,500 x g, 10 min, 4 °C), resulting in a protein pellet at the bottom of each cluster tube. The MeOH was aspirated, and protein pellets were either carried forward or stored at −80 °C.

### Protein pellet redissolving, reduction, alkylation, and streptavidin bead enrichment

Cluster tubes containing protein pellets were resuspended in 80 uL 600 mM Tris-HCl Buffer with 4% SDS (pH = 8). Samples were boiled at 95 °C for 15 minutes, sonicated in a bath sonicator at room temperature for 10 minutes, boiled again at 95°C for 5 minutes to dissolve the protein pellet, and then 140 uL of MQ H_2_O was added to each sample. Dithiothreitol (DTT) in water (50 μL, 54 mM stock, 10 mM final concentration) was added and the mixture incubated at 95 °C for 10 minutes. After the mixture was allowed to cool to room temperature, iodoacetamide in water (50 μL, 128 mM stock, 20 mM final concentration) was added and incubated in the dark at room temperature for 30 minutes. Samples were centrifuged (2,500 x g, 10 min) to ensure any insoluble material was removed, transferred to a 0.5 mL/well 96-well plate, and 50 μL of Sera-MagTM Medium Capacity Streptavidin beads (Cytiva, 30152104010350) were added to each sample. The plate was sealed with a silicone 96-well plate sealing mat (VWR, 76311-636) and incubated overnight with continuous inversion.

### Automated streptavidin bead washing

Samples were transferred to a KingFisher 96 deep-well plate (Thermo Fisher Scientific 95040450B) (plate 1). Beads were washed using a Qiagen BioSprint 96 workstation (Qiagen 9000852) with KingFisher 96 deep-well plates loaded with four wash buffers: 1) 3 x PBS with 1% SDS washes (plate 2, 3, and 4, 250 μL/well), 2) 3 x PBS with 1 M NaCl (plate 5, 6, and 7, 250 μL/well), 3) 3 x PBS with 10% EtOH (plate 8, 9, 10, 250 μL/well), and 4) 3 x 50 mM ammonium bicarbonate in MQ H_2_O (plate, 11, 12, 13, 250 μL/well). The following steps were implemented for each washing buffer (4 rounds with plate 1-4; 4-7; 7-10; and 10-13): (a) first plate (1, 4, 7, or 10) collect beads (premix), (b) plate 2, 5, 8, or 11, wash (release beads, 30 sec, medium; wash beads, 1 min, medium), wash (release beads, 30 sec, fast dual mix; wash beads, 1 min, fast dual mix), wash (release beads, 30 sec, medium; wash beads, 1 min, medium), (c) plate 3, 6, 9, or 12 wash (release beads, 30 sec, medium; wash beads, 1 min, medium), wash (release beads, 30 sec, fast dual mix; wash beads, 1 min, fast dual mix), wash (release beads, 30 sec, medium; wash beads, 1 min, medium), (d) plate 4, 7, 10, or 13, wash (release beads, 30 sec, medium; wash beads, 1 min, medium), wash (release beads, 30 sec, fast dual mix; wash beads, 1 min, fast dual mix), wash (release beads, 30 sec, medium; wash beads, 1 min, medium) (Fig. S18). After the final wash, beads from plate 13 were released in a KingFisher 96 deep-well plate loaded with trypsin (0.2 μg/well, Pierce, Thermo part #90058) in 50 mM ammonium bicarbonate (plate 14, 60 μL/well, 3.3 ng/μL final trypsin concentration in 50 mM ammonium bicarbonate). The following steps were implemented: (a) plate 4, collect beads (premix), (b) plate 5, wash (release beads, 30 sec, medium; wash beads, 30 sec, medium), wash (release beads, 30 sec, fast dual mix; wash beads, 30 sec, fast dual mix), wash (release beads, 30 sec, medium; wash beads, 30 sec, medium).

### Trypsin digestion

Beads suspended in trypsin-containing 50 mM ammonium bicarbonate (0.2 µg of trypsin per sample) were transferred to a full-skirted 96-well PCR plate, capped, and incubated at 37 °C overnight with continuous inversion. The plate was centrifuged (200 x g, 20 °C, 1 min) to collect liquid/beads at the bottom of each well, then beads pelleted using a magnetic rack (DynaMag™ 96 Side Skirted Magnet) for 5 minutes. Formic acid (5 μl/well, 5% *v/v* in water) was added to a fresh full-skirted 96-well PCR plate and the supernatant from the trypsin digest was transferred to this plate.

### In-plate desalting

An Oasis HLB 96-well plate (Waters) was conditioned by washing once with MeCN+0.1% formic acid (300 μL/well, elution at 200 x g for one minute) and twice with H_2_O+0.1% formic acid (300 μL/well, elution at 200 x g for 1 min). The tryptic peptide solutions from on-bead digestion were loaded and eluted twice at 200 x g for 1 min. Adsorbed tryptic peptides were washed twice with H_2_O+0.1% v/v formic acid (300 μL/well, elution at 200 x g for 1 min), then eluted once with 80% MeCN+0.1% v/v formic acid (100 μL/well, elution at 200 x g for 1 min). Volatiles were removed by vacuum concentration, and the samples reconstituted in 40 µL 2% MeCN+0.1% formic acid.

### Proteomic data acquisition (standard sensitivity)

Samples (all dasatinib, JQ1, and linear peptide Affinity Map samples) were analyzed using a Bruker nanoElute 2 / timsTOF Pro 2 nano-UHPLC-IM/MS/MS system using a two-column nano-UHPLC separation method and data independent analysis (DIA) MS method. Acidified and digested samples (5 µL) were loaded on a trap column (Thermo-Fisher, Cat. No. 174500, PepMap Neo C18, 5 µm particle size, 300 µm ID, 5 mm length) using 12 volume equivalents of H_2_O/0.1% FA. Flow through the trap was then reversed, and peptides eluted through a separation column (Bruker, Cat. No. 1893472, Bruker 10 C18, 1.9 µm particle size, 75 µm ID, 10 cm length) using a 2-35% gradient (H_2_O/0.1% FA – MeCN/0.1% FA) at a 500 nL/min flow rate. A 20 µm ID CaptiveSpray emitter was used at 1400 V capillary voltage. MS analysis was performed using the instrument default short gradient DIA method in HyStar v. 6.2 without alteration.

### Proteomic data acquisition (medium sensitivity)

Samples (flumazenil Affinity Map experiment only) were analyzed using a Bruker nanoElute 2 / timsTOF Pro 2 nano-UHPLC-IM/MS/MS system using a two-column nano-UHPLC separation method and data independent analysis (DIA) MS method. Oasis HLB plate desalted samples (5 µL) were loaded on a trap column (Waters nanoEase M/Z Symmetry C18 Trap, 2 cm x 180 μm, 5 μm particle size, Part No.: 186008821) using 12 volume equivalents of H_2_O/0.1% FA. Flow through the trap was then reversed, and peptides eluted through a separation column with an integrated emitter tip (IonOptiks Aurora Ultimate CSI C18 (15 cm x 75 μm, 1.7 μm particle size, Part No.: AUR3-15075C18-CSI) using a 40-minute 2-35% gradient (H_2_O/0.1% FA – MeCN/0.1% FA) at 150 nL/min flow rate with 1400 V capillary voltage. MS analysis was performed using the default short gradient DIA method in HyStar v. 6.2 without alteration.

### Proteomic data acquisition (high sensitivity)

Samples (IgG Affinity Map experiment only) were analyzed using a Bruker nanoElute 2 / timsTOF Pro 2 nano-UHPLC-IM/MS/MS system using a two-column nano-UHPLC separation method and data independent analysis (DIA) MS method. Oasis HLB plate desalted samples (5 µL) were loaded on a trap column (Waters nanoEase M/Z Symmetry C18 Trap, 2 cm x 180 μm, 5 μm particle size, Part No.: 186008821) using 12 volume equivalents of H_2_O/0.1% FA. Flow through the trap was then reversed, and peptides eluted through a separation column with an integrated emitter tip (IonOptiks Aurora Ultimate CSI C18 (25 cm x 75 μm, 1.7 μm particle size, Part No.: AUR3-25075C18-CSI) using a 60-minute 2-35% gradient (H_2_O/0.1% FA – MeCN/0.1% FA) at 150 nL/min flow rate with 1400 V capillary voltage. MS analysis was performed using the default short gradient DIA method in HyStar v. 6.2 without alteration.

### Proteomic data acquisition (high sensitivity, 90 minute gradient)

Samples (cycloRGDfK Affinity Map experiment only) were analyzed using a Bruker nanoElute 2 / timsTOF Pro 2 LC/IM/MS2 system using a two-column nano-UHPLC separation method and data independent analysis (DIA) MS method. Oasis HLB plate desalted samples (5 µL) were loaded on a trap column (Waters nanoEase M/Z Symmetry C18 Trap, 2 cm x 180 μm, 5 μm particle size, Part No.: 186008821) using 12 volume equivalents of H_2_O/0.1% FA. Flow through the trap was then reversed, and peptides eluted through a separation column with an integrated emitter tip (IonOptiks AuroraTM Ultimate CSI C18 (25 cm x 75 μm, 1.7 μm particle size, Part No.: AUR3-25075C18-CSI) using a 90-minute 2-35% gradient (H_2_O/0.1% FA – MeCN/0.1% FA) at 150 nL/min flow rate with 1400 V capillary voltage. MS analysis was performed using the default short gradient DIA method in HyStar v. 6.2 without alteration.

### Proteomics data independent analysis using DIA-NN

Data was analyzed using DIA-NN 1.8, 1.9, or 2.0.^55^ A Human FASTA sequence database from UniProt (accession date: 12-05-2022) was used. A trypsin-digested *in silico* spectral library generated in DIA-NN was used for analysis. DIA-NN settings: MS1 accuracy and mass accuracy tolerance: ±12.5 ppm; missed cleavages: 1; maximum number of variable modifications: 1; peptide length: 7-30; precursor charge range: 1-4; precursor m/z range: 300-1800; fragment ion m/z range: 200-1800; modifications: C carbamidomethylation, N-term M excision, Ox(M), Ac(N-term); Precursor FDR (%) = 1.0; match between runs enabled, neural network classifier: double-pass mode; protein inference: genes; quantification strategy: robust LC (high precision); cross-run normalization: off; library-generation: smart profiling.

### Volcano plot analysis

Processing of DIANN output files (report.pg_matrix.tsv, report.pr_matrix.tsv) was done in R (4.4.1) using the MSnbase package separately for each set of samples treated with given concentration of photocatalyst conjugate and off-compete molecule or vehicle. Proteomics data was subjected to variance stabilization normalization (‘vsn’) in independent experimental groups, ^56,57^ imputation using MinProb in MSnbase, and the number of peptides used for each protein inference determined and added as metadata. Once complete, individually processed MSnSet datasets were combined into a single table. Differential expression analysis was performed using linear models (‘limma’)^58^ with Benjamini-Hochburg FDR adjustment to identify significantly enriched proteins. A volcano plot comparing the log_2_ fold change and −log_10_(p-value) was generated, with significantly enriched proteins colored by affinity measured using Affinity Map.

### Affinity Surveyor data preprocessing

Preprocessing of DIANN output files (report.pg_matrix.tsv, report.pr_matrix.tsv) was done in R (4.4.1) using the MSnbase package separately for each set of samples treated with a given concentration of photocatalyst conjugate. Proteomics data was subjected to variance stabilization normalization (‘vsn’) and left-censored missing value imputation (‘MinProb’) using default settings,^56,57^ and the number of peptides used to quantify each protein determined from the peptide-level report and then added as metadata. Once complete, individually processed MSnSet datasets were combined into a single table.

### Affinity Surveyor

Our approach for apparent (non-equilibrium) binding affinity (*K*_d_^app^) elucidation is analogous to a proteome-wide radioligand competitive binding assay, in which streptavidin IP/MS intensity is used as an indirect readout of competitive protein binding site occupancy by a photocatalyst conjugate probe and a competing ligand molecule with distinct *K*_d_^app^ values (*K*_d_^ligand^ and *K*_d_^probe^). As photocatalytic proximity labeling measured by mass spectrometry is more complex than a simple radioactivity readout corresponding to receptor occupancy, two assumptions are made to constrain the kinetic model: 1) the MS intensity difference of proteins obtained from streptavidin/biotin IP induced by photocatalyst off-compete by a non-photocatalytic ligand is directly proportional to the occupancy of a single binding site by the photocatalyst conjugate, and 2) protein labeling does not materially alter affinity for the probe or ligand molecules. With these constraints in place, a rearranged Cheng-Prusoff equation can be used to independently solve for the *K*_d_^app^ values for each molecule^59^ using protein intensities derived from photocatalytic labeling experiments where the concentrations of both the probe and ligand molecules are systematically varied. All reported affinity profiling experiments use four concentrations of the photocatalyst conjugate and 12 concentrations of competitive non-photocatalytic molecule in duplicate, for a total of 96 samples.

Affinities were measured in Python (3.12.2). The EC_50_ of competitive binding of a ligand molecule to each protein in the proteome at each photocatalytic probe concentration for which < 25% of values are imputed is modeled using Equation 1, assuming a Hill coefficient of 1:

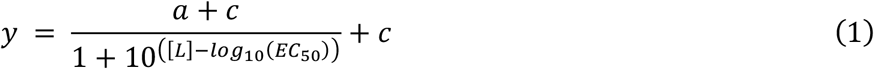

where [L] = competitive ligand molecule concentration, y = integrated MS intensity for a protein, and *a* (maximum intensity minus minimum intensity), *c* (minimum intensity), and the EC_50_ (concentration of ligand molecule displacing 50% of the quantity of probe molecule present in the absence of competition) are all freely optimized parameters. Nonlinear fitting for Equation 1 was performed using lmfit.^60^

During optimization, the initial value for *c* is set to the global minimum intensity detected from all proteins and samples and the initial *a* value is set to the maximum of the signal intensity value in samples used for curve fitting. The significance (p-value) of the fit is determined by performing a one-way F-test with the SciPy ‘stats’ module, comparing Equation 1 to the reduced model y = a, representing a null hypothesis in which there is no relationship between ligand molecule concentration and signal intensity. Each competitive model is fit 30 times with different initial values set for the EC_50_ parameter drawn from a random uniform distribution between −5 and 3 (two order of magnitude beyond the range of concentrations used), retaining the result with the most significant fit. We then test the impact of removing the data point corresponding to treatment with vehicle in place of the unmodified ligand by repeating the nonlinear fitting process and evaluating whether inclusion of vehicle is required for the fit to remain significant. Fits whose significance depends on the inclusion of the vehicle are removed.

The relationship between *K*_d_ and EC_50_ for pseudo-first order binding kinetics, i.e. at low concentrations of protein ([Protein] << [Ligand]), is described by the Cheng-Prusoff equation^35^ according to Equation 2:

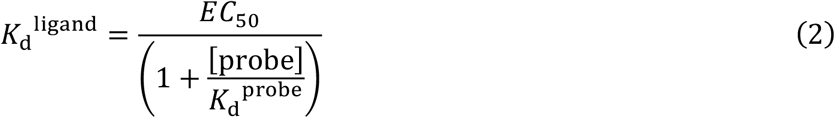

where [probe] = probe concentration, *K*_d_^ligand^ *=* dissociation constant of ligand molecule, *K*_d_^probe^ = dissociation constant of photocatalytic probe, and EC_50_ = EC_50_ of competitive binding by the ligand molecule at each probe molecule concentration. Equation 2 can be expressed as Equation 3, a Schild plot:

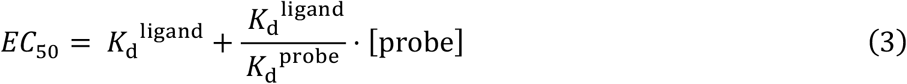

As EC_50_ errors are measured in logarithmic space, the Schild plot is modeled in logarithmic space as well (Equation 4) using curve_fit in the scipy.optimize package^61^ :

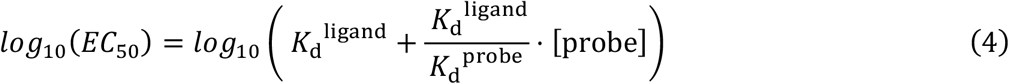

The significance of the modeled *K*_d_^ligand^ */ K*_d_^probe^ value is measured with a one-way F-test using the SciPy ‘stats’ module, comparing Equation 4 to a reduced model log_10_(EC_50_) = a, representing the null hypothesis that there is no relationship between the value of EC_50_ and the concentration of the probe photocatalyst conjugate. The *K*_d_^app^ of the free molecule (often a drug, metabolite, peptide, or protein of interest) will be different from an observed EC_50_ by an amount proportional to the ratio of the probe and ligand affinities multiplied by the concentration of its photocatalyst conjugate. As the affinity or concentration of a probe photocatalyst conjugate becomes weaker, the *K*_d_^ligand^ value of the unconjugated molecule approaches the measured EC_50_ value.

For each protein observed, we perform EC_50_ determination at each photocatalytic probe concentration and use significant (p < 0.001) values of the EC_50_ to measure *K*_d_^probe^ and *K*_d_^ligand^ via the Cheng-Prusoff equation. Individual proteins have widely varying affinity for the photocatalyst conjugated probe and unmodified ligand molecule; the vast majority do not display any significant EC_50_ values at any value of [probe], reflecting that they do not bind either the probe or ligand molecules. Other proteins which do bind both the probe and ligand molecules within the range of concentrations used show significant EC_50_ values. These are then subjected to Cheng-Prusoff analysis to calculate *K*_d_^probe^ and *K*_d_^ligand^. A tiered approach is used to estimate binding affinities for proteins based on the number of significant EC_50_ values observed for each protein:

1. Three or more significant EC_50_ values at different probe concentrations, measurable *K*_d_^ligand^ / *K*_d_^probe^: *K*_d_^probe^ and *K*_d_^ligand^ are directly measured by fitting the data to Equation 4. If the significance of the modeled slope (*K*_d_^ligand^ / *K*_d_^probe^) is significant (p < 0.1), both values are reported along with the error from fitting. If the slope is not significant (p > 0.1) and data is tightly distributed (confidence interval < ±0.25), the relative affinity for the probe ligand vs. the ligand molecule is too low to accurately measure. At this limit (*K*_d_^ligand^ / *K*_d_^probe^ ≈ 0), each EC_50_ ≈ *K*_d_^ligand^; *K*_d_^ligand^ is therefore reported as the weighted average of the EC_50_ values in these cases. The weighted average of EC_50_ values is determined by Equation 5, where 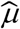 is the weighted average, N is the total number of EC_50_ used for fitting, *i* indicates the EC_50_ values and *σ*(standard error) of EC_50_ from fitting in Equation 1 at each value of [probe]. All such proteins are flagged as having modeled *K*_d_^ligand^ / *K*_d_^probe^ values.

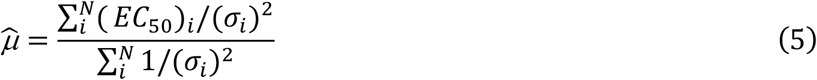

2. Three or more significant EC_50_ values at different probe concentrations, insufficient data to measure *K*_d_^ligand^ / *K*_d_^probe^: Where the slope is not significant (p > 0.1) and data is not tightly distributed (confidence interval > ±0.25), *K*_d_^ligand^ / *K*_d_^probe^ cannot be empirically determined. The weighted average EC_50_ value (calculated using Equation 5) is therefore reported, reflecting the null hypothesis *K*_d_^ligand^ / *K*_d_^probe^ ≈ 0).

3. Two significant EC_50_ values at different probe concentrations: Curve fitting with two points cannot afford an accurate estimation of error in the resulting fit. The value measured at the lowest photocatalytic probe concentration is therefore reported as a best estimate of *K*_d_^ligand^ as it is the closest empirical value to the y-intercept in the Schild plot.

4. One significant EC_50_ value at one probe concentration: *K*_d_^ligand^ is reported as the EC_50_.

5. Zero significant EC_50_ values: The most significant EC_50_ value is reported as *K*_d_^ligand^.

For each protein, the total significance of the data used for *K*_d_^ligand^ estimation is reported as the Fisher combined p-values of all fitted EC_50_ values observed, and this aggregated p-value is then subjected to Benjamini-Hochburg FDR estimation to account for multiple hypothesis testing and represents the significance of all evidence used to support a reported *K*_d_^ligand^ value. For most proteins, *K*_d_^ligand^ / *K*_d_^probe^ is small because attaching the photocatalyst to a ligand significantly reduces its affinity (*K*_d_^ligand^ << *K*_d_^probe^), consistent with the impact of attaching other types of photoaffinity labels; in such cases, weighted average EC_50_ values give very good estimates of *K*_d_^app^ because the influence of photocatalytic probe concentration on any observed EC_50_ value is small. Since proteins that have measurable EC_50_ values only at high probe concentration (scenarios 2, 3, and 4 above) likely have small *K*_d_^ligand^ / *K*_d_^probe^ ratios, and most protein-ligand systems have small *K*_d_^ligand^ / *K*_d_^probe^ ratios (see supplementary dataframe reporting all fitted parameters, Supplementary_Data_1.xlsx), single EC_50_ values (scenario 4) can be used as acceptable best estimates of *K*_d_^ligand^. In scenarios with exactly two significant EC_50_ values, using the EC_50_ value obtained at the lower value of photocatalyst concentration provides an improved estimate as it is measured closer to the value of *K*_d_^ligand^ in the Schild plot.

We find that the assumption of small values for *K*_d_^ligand^ / *K*_d_^probe^ is valid in data generated using photocatalytic affinity profiling for dasatinib, which shows good correlation with reported affinities for kinases with *K*_d_^ligand^ values measured using three or more, two, or one significant EC_50_ value (Fig. S14, Figure 2G). Indeed, correlation between *K*_d_^ligand^ values measured by photocatalytic affinity profiling and literature values is good and has a slope near one, reflecting the validity of both the broader assumptions made to constrain the kinetic model used for analysis and our tiered data processing pipeline approach.

