## Supplementary Information for "Global Protein-Ligand Binding Affinity Profiling via Photocatalytic Labeling"

#### Supplementary Figures

##### Fig. S1. Development of an improved photocatalyst for Affinity Map

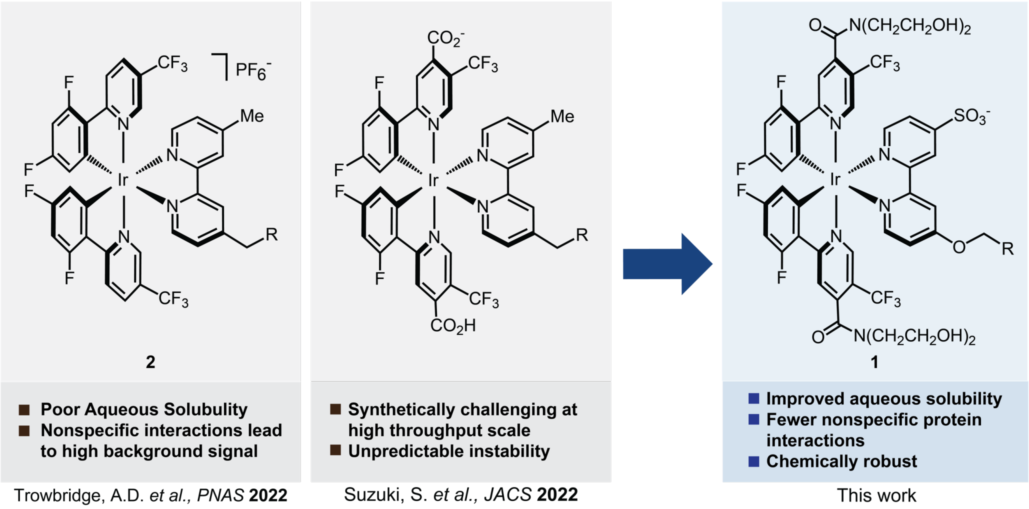

In photocatalytic proximity labeling, a photocatalyst near a protein of interest is used to generate reactive intermediates. These reactive species then covalently tag proteins in close proximity to the photocatalyst, enabling their identification by MS. High spatial resolution methods use carbenes generated by sensitized photolysis of diazirines for proximity labeling due to their tight labeling radius (< 4 nm) and promiscuous reactivity. By using a small molecule targeting modality for the photocatalyst, small molecule binding targets are selectively labeled, while using a larger (ca. 30 nm) primary/secondary antibody targeting modality enables protein-protein interaction discovery. However, current-generation photocatalysts for diazirine activation are either highly hydrophobic and water-insoluble (**2**) or are anionic at physiological pH, are challenging to synthesize, and can present stability liabilities. These shortcomings have hindered widespread use of diazirine-based proximity labeling, preventing high throughput applications. Catalysts based on Ir(dF(CF3)ppy)2(dmbpy) such as **2** are highly stable and synthetically tractable, leading to their common use in small molecule target-ID applications. However, hydrophobic photocatalysts exhibit high background labeling caused by nonspecific protein binding, limiting the maximum achievable difference in MS signal observed in competitive binding assays using small molecule photocatalyst conjugates and reducing the signal-to-noise ratio. An ideal photocatalyst for high throughput applications, such as binding affinity profiling, requires high solubility, bench stability, and good photocatalytic activity.

We report the synthesis and characterization of **1**, a novel iridium photocatalyst for proximity labeling. The complex is charge-neutral, light and heat stable, hydrophilic, and able to activate diazirine probes for catalytic labeling. We demonstrate that this new catalyst has a high aqueous solubility (**Figs. S2, S3**), activates diazirine probes and maintains diazirine-sensitization activity after 30 minutes of blue light irradiation (**Figs. S2, S3**), has reduced nonspecific protein binding and associated MS background signal (**Fig. S3, S5**), can effectively label the binding targets of small molecules using a small molecule targeting modality (**Fig. S5**), and identifies protein-protein interactions using a primary/secondary antibody targeting modality (**Fig. S6**). Further, we show that these improved properties enable enhanced binding affinity profiling (**Fig. S7, S10,** and main text) in comparison with **2**. Overall, **1** represents a next-generation photocatalyst which exhibits improved physical properties and robust diazirine-sensitization activity for carbene-based proximity labeling applications in lysates/extracts and on cell surfaces, and is especially well-suited to high throughput methods such as binding affinity profiling.

**Fig. S2. Diazirine photosensitization yields and logP for synthesized photocatalysts**

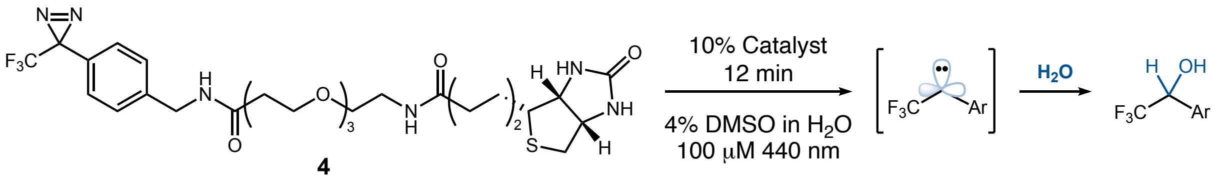

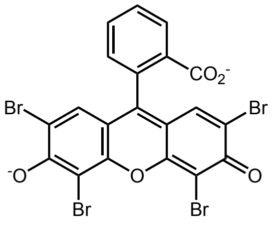

Eosin Y
Dz-Sensitization = 0%

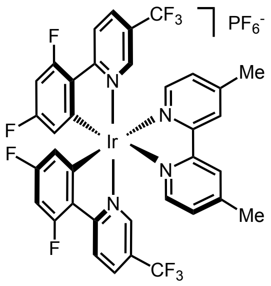

**16**
Dz-Sensitization = 73%
logP = 1.12

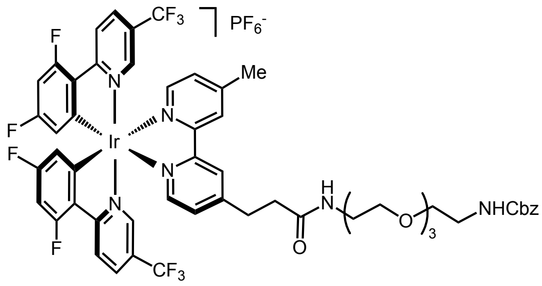

**2**
Dz-Sensitization = 77%
logP = 1.27

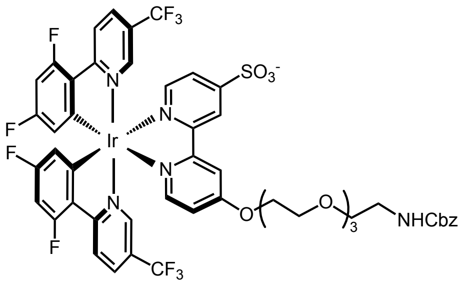

**20**
Dz-Sensitization = 9%
logP = 0.92

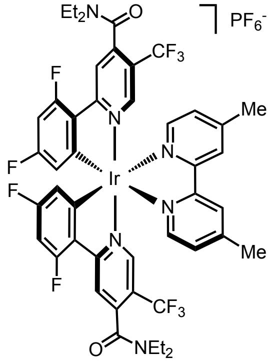

**18**
Dz-Sensitization = 22%

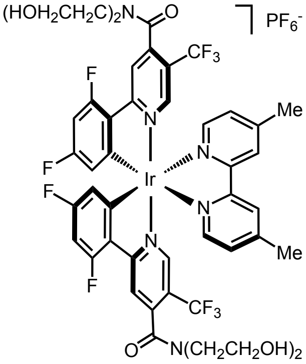

**19**
Dz-Sensitization = 29%
logP = -0.44

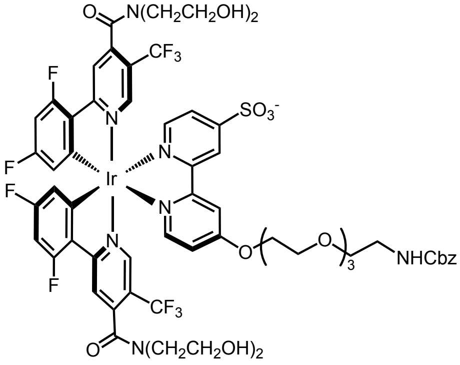

**1**
Dz-Sensitization = 26%
logP = 0.75

**Photocatalytic diazirine sensitization assay:**
Diazirine-biotin (**4**, 100 μM), photocatalyst (10 μM), and internal standard (IS) (2-chloro-5-trifluoromethylpyridine, 100 μM) were dissolved in 350 μL H2O in a 1.5 mL capped centrifuge tube. The tube was degassed with argon and placed 17.8 cm above two 440 nm, 100 W COB LED light (Chanzon, 1DGL-JC-100W-440) placed side-by-side (**Fig. S20**). The samples were then irradiated for the appropriate amount of time. After irradiation, $D_2O$ (100 μL) was added, and the concentration of diazirine measured by $^{19}F$ NMR spectroscopy relative to the IS, comparing against a negative control in which the photocatalyst is absent. Experiments were performed in triplicate and average values are reported.

**LogP measurement:**
$H_2O$ (150 μL) and 1-octanol (150 μL) were combined in a 1.5 mL capped centrifuge tube to which appropriate catalyst (3 μL of a 100 μM stock, final concentration 1 μM, 1% DMSO) was added. The sample was vortexed for 7 seconds, let equilibrate for 2 min., then centrifuged at 21,000 x g for 10 min. Organic and aqueous layers were separated, solvent was evaporated, and solids redissolved in DMSO (100 μL). The photocatalyst concentration was measured by absorbance at 375 nm. Each measurement was performed in triplicate and averaged for the reported value.

**Thermodynamic solubility measurement:**
1.125 mg of catalyst (as a 5 mM stock in DMSO) was added to a 1.5 mL capped centrifuge tube and solvent was evaporated. The powder was resuspended in $H_2O$ or PBS (100 μL), sonicated for 5 min., then inverted end over end for 24 hours at 37 °C. The sample was centrifuged (21,000 x g for 10 min.) and supernatant was transferred to a fresh tube. Solvent was evaporated and solid redissolved in DMSO (100 μL). The photocatalyst concentration was determined by absorbance at 375 nm. Each measurement was performed in triplicate and averaged for the reported value.

**Fig. S3. Catalyst 1 has higher aqueous solubility and less nonspecific photocatalytic labeling than 2**

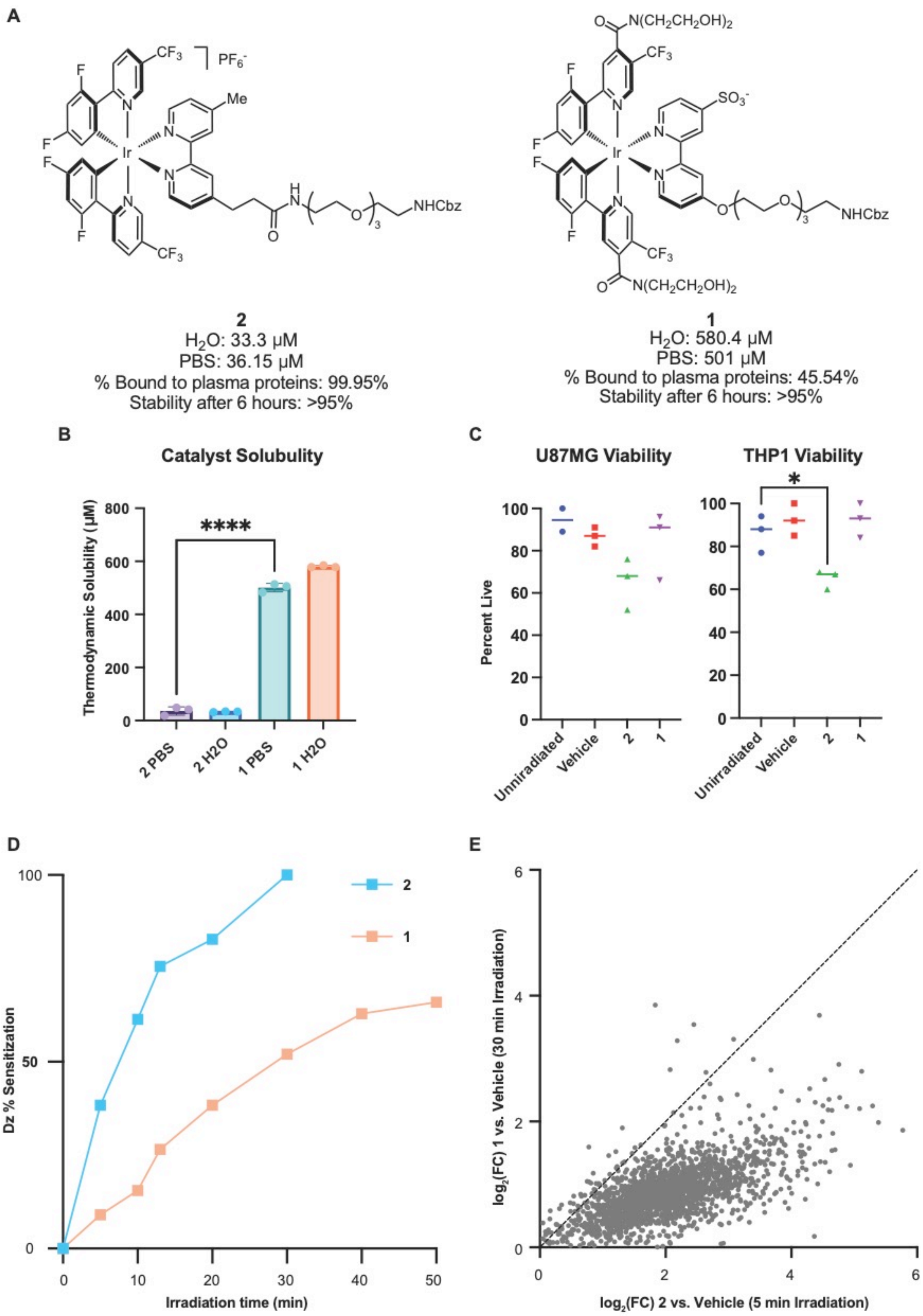

**A.** Structure of catalysts used for photophysical property comparison. Thermodynamic solubilities listed in PBS and DI $H_2O$. Binding data of each catalyst to human plasma proteins demonstrating increased non-specific binding of **2** compared to **1**. Both catalysts are stable after incubation at 37 °C for 6 hours. **B.** Thermodynamic solubility comparison in PBS and $H_2O$. **C**. U87MG and THP1 cell viability following incubation with **1** or **2** and 15-minute irradiation. **1** is not toxic to cells. **D.** Diazirine sensitization yields over time. **E.** Comparison of protein biotinylation measured by streptavidin enrichment and bottom-up MS proteomics by **2** and **1** vs. no-catalyst control at timepoints where **1** has greater diazirine consumption than **2** shows increased nonspecific protein labeling by **2**, which results from its greater nonspecific protein binding.

**Experimental protocol:**
**C.** THP-1
3 x $10^5$ THP-1 cells were collected by centrifugation (800 x g, 5 min, 20 °C), and washed twice with 40 mL of cold PBS. Cells were resuspended in cold PBS (0.75 mL) and aliquoted into 8-strip cluster tubes (12 tubes) in a rack and kept on ice (50 µL / aliquot, approximately 20,000 cells) for 20 minutes. Cells were then treated with free catalyst (**1** and **2**) or a vehicle control (10 µL / aliquot, 6 uM) and incubated on ice for 20 minutes (final volume 60 µL, final concentration of catalyst 1 µM). Samples were then irradiated with 440 nm light for 15 minutes at 4 °C. Following irradiation, cells were stained with trypan blue (1:1 with 0.4%, VWR) and cell viability was calculated with a Countess 3 (Invitrogen).

U87MG
2 x 6-well treated cell culture plates were seeded with 5 x $10^5$ U87MG cells and allowed to grow until confluent (~3 days). The media was aspirated and 400 µL of phenol red-free media was pipetted over the adherent cells in each well. DMSO (40 µL) was added to each well and cells were incubated on ice for 20 minutes. Cells were then treated with the free catalysts (1 and 2) or a vehicle control (40 µL, 120 µM) and incubated on ice for 20 minutes (final volume 480 µL, 10 µM catalyst). Each 6-well plate was then irradiated for 15 minutes at 4 °C. Following irradiation, cells were released from the plate, stained with trypan blue (1:1 with 0.4%, VWR), and cell viability was calculated with a Countess 3 (Invitrogen).

**E.** K562 cell lysate was prepared according to **method 1**. 200 µL of 1.5 mg/mL lysate containing 200 µM Dz-Bt was added to 1.5 mL capped centrifuge tube. Added 2 µL of catalyst (100 µM, DMSO) or vehicle and incubated on ice for 30 min. Samples were degassed with argon according to **method 3** and irradiated for appropriate amount of time using the apparatus in **method 3**. Samples were then processed according to **methods 4-7**. Each experiment was performed in triplicate. **Human Plasma Protein Binding:** Assay was conducted by Pharmaron.

**Fig. S4. UV-Vis Absorption and Emission Spectra for 1 and Diazirine Sensitization at Multiple Concentrations**

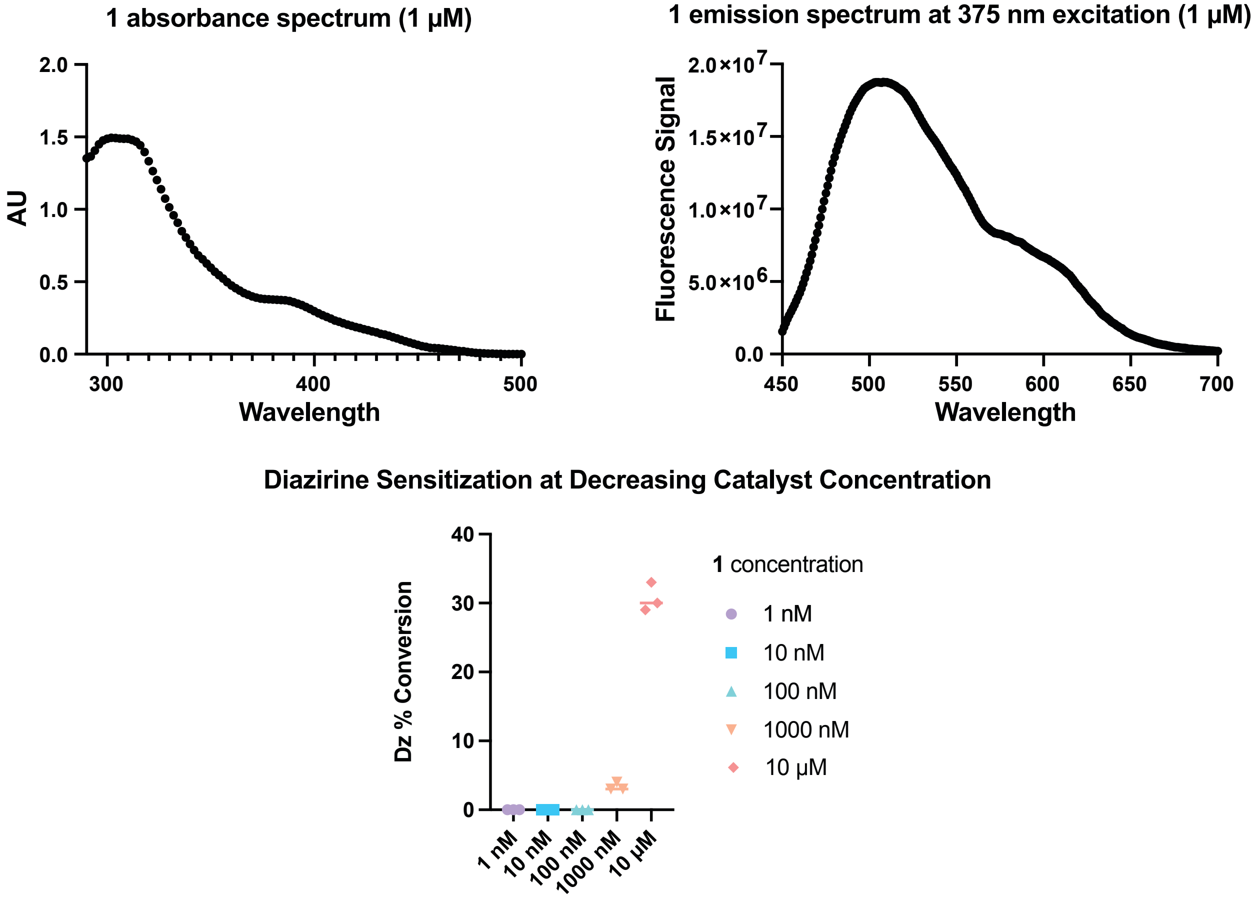

**Photocatalytic diazirine sensitization assay with different concentrations:**
A stock solution of diazirine-biotin (**4**, final concentration 100 µM), 2-chloro-5-trifluoromethylpyridine IS (final concentration 100 µM), and water was aliquoted into 15 1.5 mL centrifuge tubes. The catalyst (**1**) was then added to the tubes at the appropriate concentration (final volume 350 µL), vortexed for 7 seconds to mix, and spun down at 200 x g for 10 seconds. The tubes were degassed with argon, placed 17.8 cm above two 440 nm 100 W COB LED lights (**Fig. S20**), and irradiated for 15 minutes. 100 µL of deuterium oxide was added to the tubes, and samples were transferred to NMR tubes and analyzed by $^{19}F$ NMR spectroscopy. Quantitation was done relative to IS and comparing against a negative control in which the photocatalyst is absent.

**Fig. S5. Small molecule target-ID: comparison between 1 and 2**

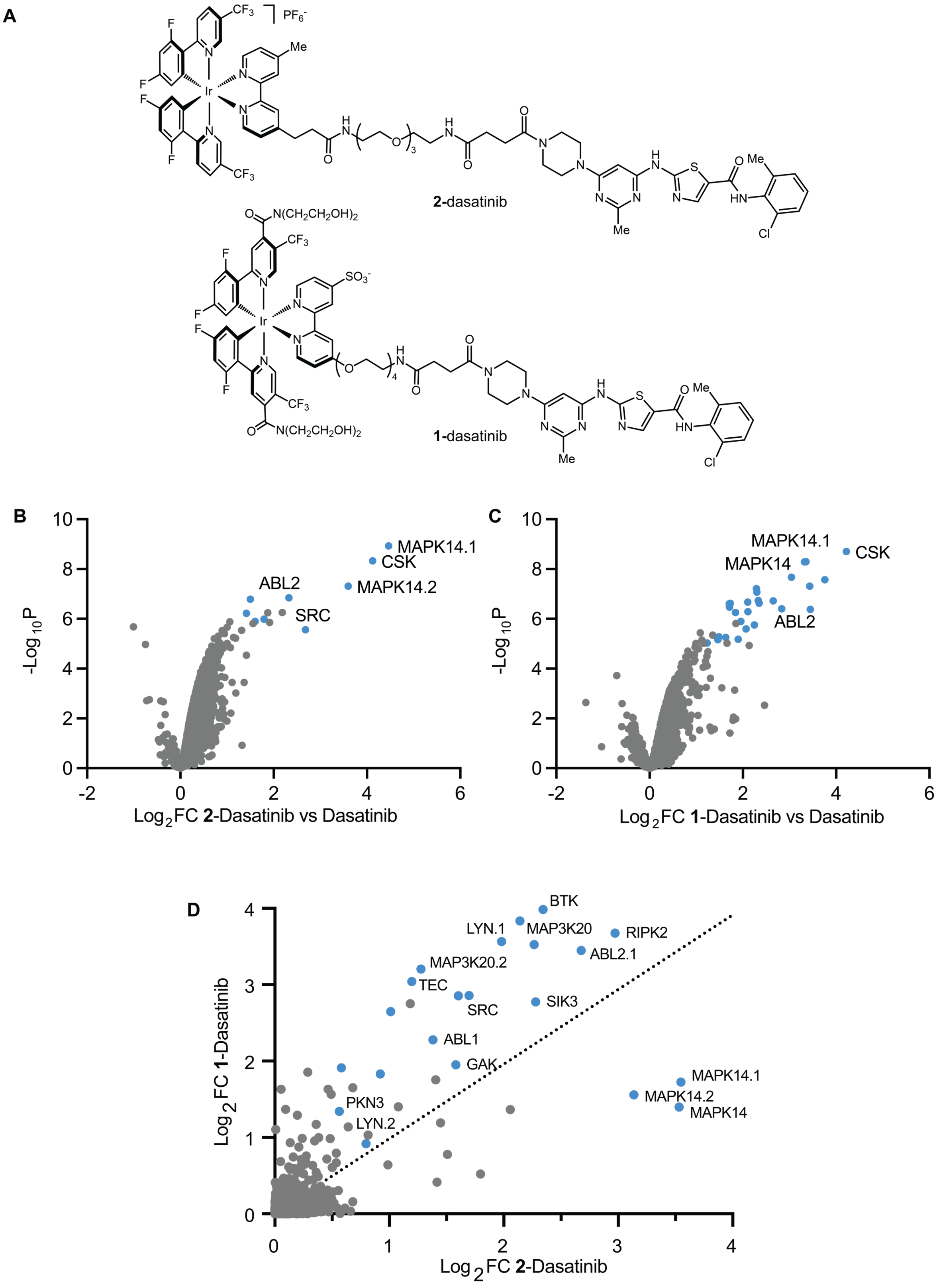

**A.** Structures of **1** and **2** conjugated to the SRC/ABL kinase inhibitor dasatnib **B.** Volcano plot comparing protein labeling by **2**-dasatinib (1 μM) vs **2-**dasatinib (1 μM) + dasatinib (10 μM, off-compete control). Kinases highlighted blue. **C.** Volcano plot of **1**-dasatinib (1 μM) vs **1**-dasatinib (1 μM) + dasatinib (10 μM, off-compete control). Kinases highlighted blue **D.** Comparison of $log_2FC$ from **B** and **C.** Kinases and nucleotide-binding proteins are more enriched by **1** than **2** despite lower diazirine sensitization yield for **1** (**Figs. S2, S3**) due to lower background signal caused by nonspecific protein labeling. **B** and **C** performed according to **methods 1-7** with a single off-compete concentration (10 μM).

**Fig. S6. Antibody-targeted photocatalytic proximity labeling at CD19 using 1 and 2**

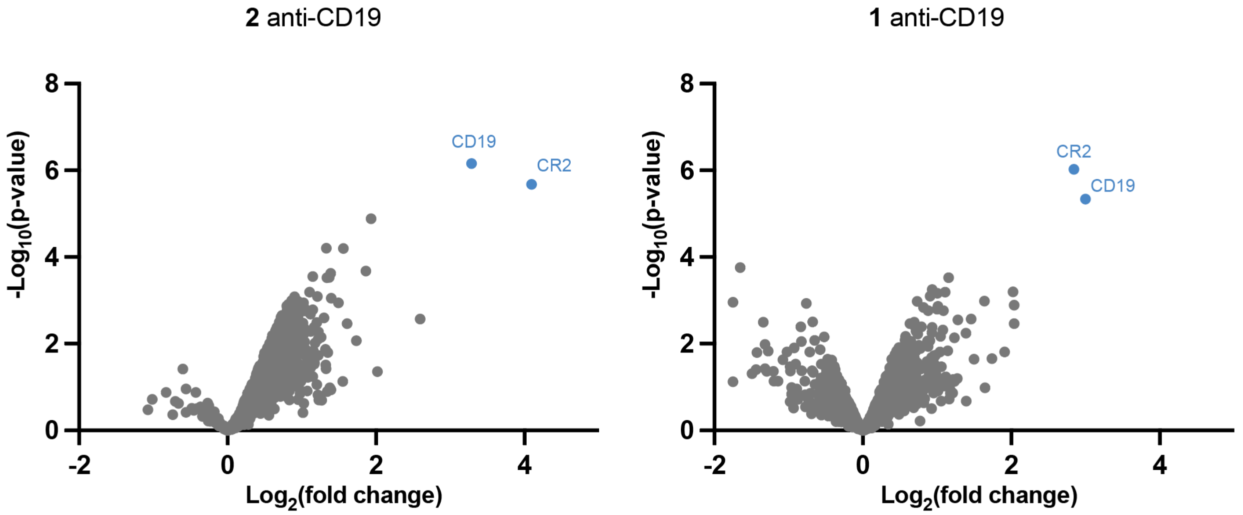

Both catalysts **1** and **2** are able to enrich the target (CD19) and a known interactor (CR2) [1,2] when conjugated to a goat anti-mouse secondary antibody targeting live cells stained with mouse anti-CD19 vs. isotype control.

**Preparation of photocatalyst-antibody conjugate:**

Catalyst **1**: **1**-DBCO (20 µL, 5 mM DMSO) and azido-PEG3-NHS ester (BroadPharm, BP-21605) (1 µL, 100 mM DMSO) were mixed in a vial and allowed to react for 10 minutes. Separately, goat anti-mouse polyclonal antibody (Jackson ImmunoResearch 115-005-205) (180 µL, 0.5 mg/mL, PBS) was combined with $NaHCO_3$ (18 µL, 1 M). The solution of **1**-NHS ester (19 µL) generated through strain-promoted cycloaddition was added and allowed to react with the antibody for 1 hour at room temperature. The reaction was then passed through a ZEBA 40 kDa 0.5 mL desalting column (Thermo Fisher Scientific, A57760) that had been equilibrated with PBS. The antibody-catalyst solution was analyzed by BCA (50:1 solution A:B; A: 10 mg/mL Bicinchonic acid, 20 mg/mL $Na_2CO_3$, 1.6 mg/mL sodium tartarate, 4 mg/mL NaOH, 9.5 mg/mL $NaHCO_3$. B: 40 mg/mL $CuSO_4$) vs. a serial dilution of BSA. Catalyst concentration was measured by fluorescence of the sample at 485 nm following excitation at 380 nm compared to a serial dilution of **1**-NHS. The measured concentration was 0.52 mg/mL antibody, with 12 catalysts/antibody.

Catalyst **2**: **2**-DBCO (20 µL, 5 mM DMSO) and azido-PEG24-NHS ester (BroadPharm, BP-23542) (1 µL, 100 mM DMSO) were mixed in a vial and allowed to react for 10 minutes. Separately, goat anti-mouse polyclonal antibody (Jackson ImmunoResearch 115-005-205) (450 µL, 0.5 mg/mL, 1:1 PBS/DMSO) was combined with $NaHCO_3$ (45 µL, 1 M). **2**-NHS ester solution (5.625 µL) was added and allowed to react for 1 hour at room temperature. The reaction was then passed through a ZEBA 40 kDa 0.5 mL desalting column (Thermo Fisher Scientific, A57760) that had been equilibrated with 1:1 PBS/DMSO. The antibody-catalyst solution was analyzed by BCA for protein concentration (50:1 solution A:B; A: 10 mg/mL Bicinchonic acid, 20 mg/mL $Na_2CO_3$, 1.6 mg/mL sodium tartarate, 4 mg/mL NaOH, 9.5 mg/mL $NaHCO_3$. B: 40 mg/mL $CuSO_4$) compared to a serial dilution of BSA. Catalyst concentration was measured by fluorescence of the sample at 485 nm following excitation at 380 nm compared to a serial dilution of **2**-NHS. The measured concentration was 0.14 mg/mL antibody, with 3.85 catalysts/antibody.

**Live cell labeling:**

Comparison of catalysts each to isotype control in triplicate, 6 samples per catalyst, 12 samples total. $10^8$ Ramos (RA 1, CRL-1596) cells were harvested, cooled to 4 °C, pelleted, and washed 2x (DPBS 1% FBS), and pelleted (800 x g, 5 min). The following steps were all performed at 4 °C. The cell pellet was resuspended in DPBS 1% FBS (10 mL) and evenly split into 0.5 mL aliquots in 1.5 mL capped centrifuge tubes. Cells were pelleted (800 x g, 5 min) and resuspended in DPBS 1% FBS (0.2 mL) containing 5 µg anti-CD19 primary (SJ25C1, IgG1) or isotype control (mouse IgG, Santa Cruz sc-2025) and continuously inverted end over end at 4 °C for 1 hour. Samples were then pelleted (800 x g, 5 min), washed with DPBS 1% FBS (0.6 mL, 1x), and resuspended in DPBS 1% FBS (0.2 mL) containing 10 µg goat-anti-mouse secondary antibody conjugated to appropriate catalyst and continuously inverted end over end at 4

°C for 2 hours. Samples were then pelleted (800 x g, 5 min), washed with DPBS (0.6 mL, 1x), and resuspended in DPBS (0.2 mL) containing 250 µM Dz-Bt. Samples were irradiated with 440 nm light using the apparatus in **Method 3** for 15 minutes, then washed with DPBS (0.6 mL, 3x). Cells were resuspended in RIPA (Millipore, 20-188) containing 1% SDS (0.4 mL) and sonicated in a cup horn sonicator (**Method 1**) for 15 minutes (20 seconds on, 10 seconds off at 60% intensity). Samples were heated to 95 °C for 5 minutes, then centrifuged (15,000 x g, 10 minutes) to remove insoluble debris. 375 µL of supernatant was transferred to new 1.5 mL tubes. DTT (20 µL, 100 mM) was added to each sample, and the samples heated to 95 °C for 5 minutes. After allowing the sample to cool to room temperature, iodoacetamide was added (15 µL, 200 mM) and incubated in the dark. After 30 minutes, 150 µL streptavidin beads (Cytiva, 30152104010350) were added to each sample and the samples inverted end over end for 2 days at room temperature. Samples were then processed according to **Methods 5-9** and **Method 11** and protein abundances separately compared between the anti-CD19 and isotype treated samples for each catalyst.

**Fig. S7. Dose-responsive competetive photocatalytic labeling of recombinant BRD4 (AAs 49-460)**

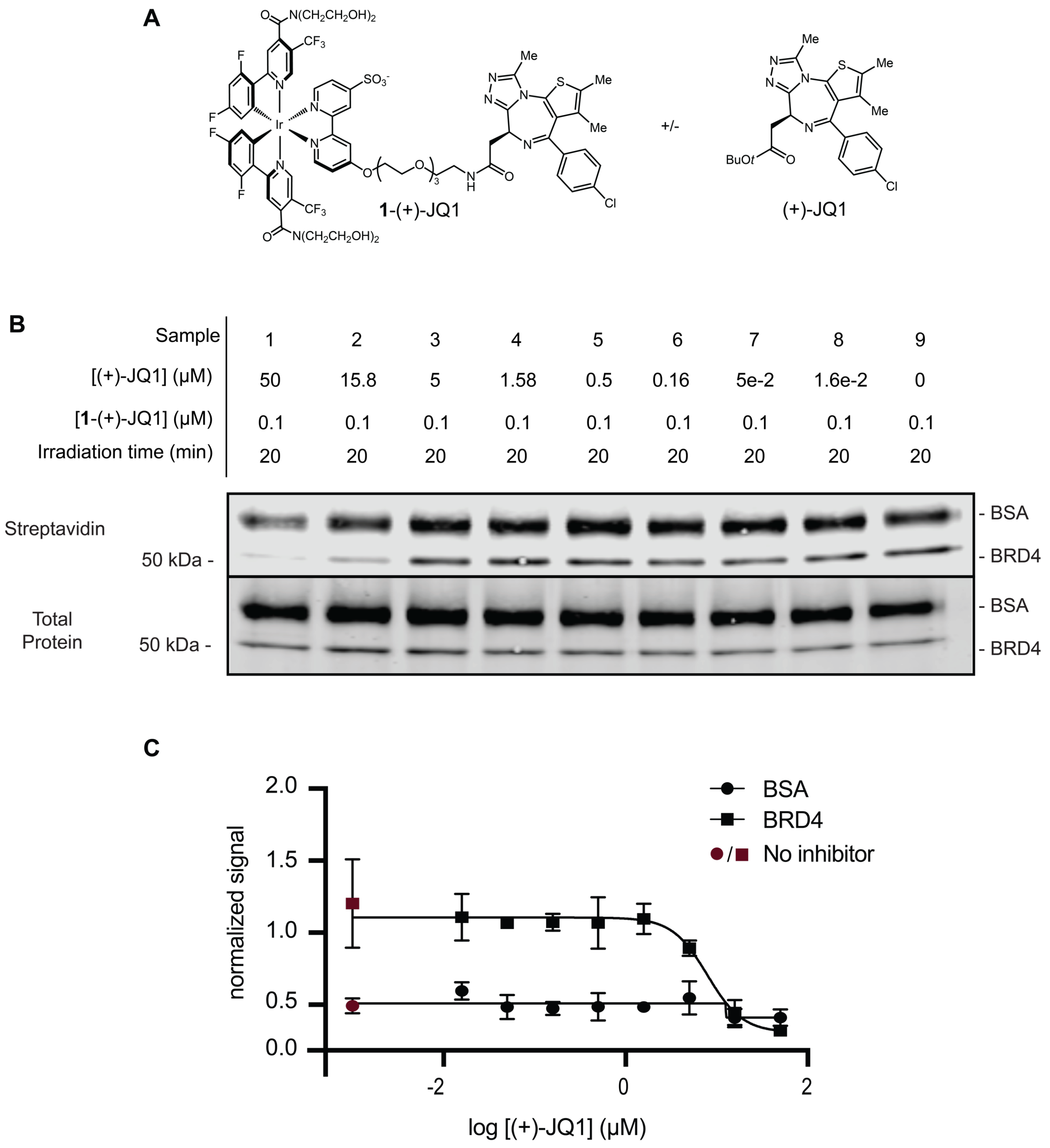

**A.** Structures of **1**-(+)-JQ1 and (+)-JQ1 **B.** Representative western blot showing dose-dependent labeling of recombinant BRD4 BD1+BD2 (AAs 49-460) domains and BSA **C.** Quantified densitometry values for BSA and BRD4 bands from the blot in **B** normalized against total protein intensity. Experiment performed in triplicate.

**Photocatalytic labeling of recombinant BRD4 BD1+BD2:**
To a 1.5 mL capped centrifuge tube was added $(His)_6$-BRD4 (BD1+BD2 domains, AAs 49-460, BPSBioscience catalog no. 31045) (2 µL, 31.76 µM), bovine serum albumin (5.08 µL, 13.79 µM), Dz-Bt (186.7 µL, 400 µM) and 1x PBS (46.12 µL, pH 7.4). To a separate 1.5 mL capped centrifuge tube was added (+)-JQ1 (12 µL of 4x desired concentration, in PBS + 2% DMSO) followed by a solution of the proteins described above (24 µL). This was incubated on ice for 15 minutes. **1**-(+)-JQ1 (12 µL, 0.4 µM, PBS + 1% DMSO) was added, and the mixture incubated on ice for 15 minutes (final volume 48 µL, 0.75% DMSO). Samples were then degassed with argon and irradiated with 440 nm light for 20 minutes according to **Method 3**. The sample was diluted with 4x Laemlli buffer with 10% BME (12 µL) and heated to 95 °C for 10 minutes. The samples were cooled to room temperature and centrifuged. The samples were loaded onto a Novex 10-20% tris-glycine gel with protein ladder (Biorad Cat. 1610373) in freshly prepared Tris running buffer and subjected to electrophoresis (200 V, 45 min). The gel was washed (3x DI $H_2O$) and transferred to nitrocellulose membrane using a Trans-Blot Turbo Transfer System (BioRad 1704150). The membrane was immersed in REVERT total protein stain (Li-Cor 926-11011) for 10 minutes. Stain was removed, the membrane washed (33.5:150:316.5, AcOH:MeOH:$H_2O$, 3x 5 minutes), rinsed with water, and imaged using a Li-Cor Odyssey CLx scanner at 700 nm. The membrane was immersed in 30% MeOH in 100 mM aqueous NaOH for 10 minutes, washed with this solution (2x), rinsed with water (3x), and incubated with Intercept Blocking Buffer (Li-Cor, 927-60001) for 1 hour. The blocking buffer was decanted and the membrane was incubated with 10 mL 1X TBST containing 1:10,000 dilution of IRDye 800 CW streptavidin (Li-Cor 926-32230) for 1 hour. The buffer was decanted and the membrane washed with 1X TBST (3x 5 minutes) and water before imaging using a Li-Cor Odyssey CLx scanner at 800 nm. Pixel densitometry was performed in Image Studio V. 5.5.4. Streptavidin 800 nm channel signal was divided by total protein signal in the 700 nm channel to determine the protein abundance-normalized biotinylation signal for each protein band. Experiment performed in triplicate, whole gel images shown below.

Gel 1, samples 1-9 replicate 1 stained for streptavidin (left) and total protein loading control (right).

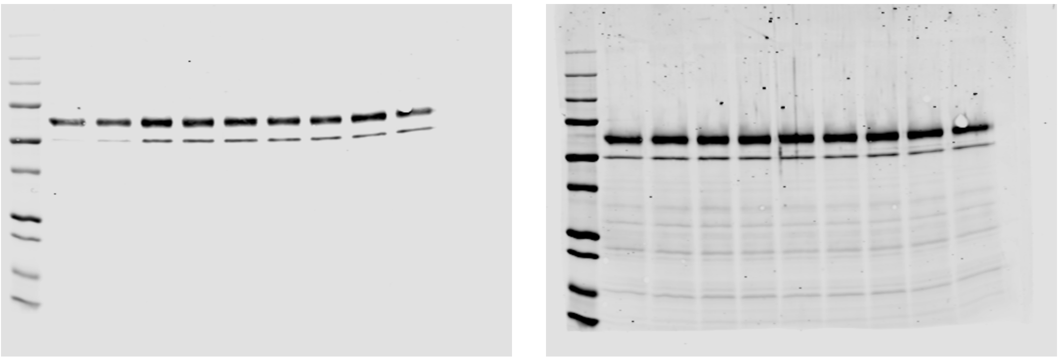

Gel 2, samples 1-9 replicate 2 stained for streptavidin (left) and total protein loading control (right).

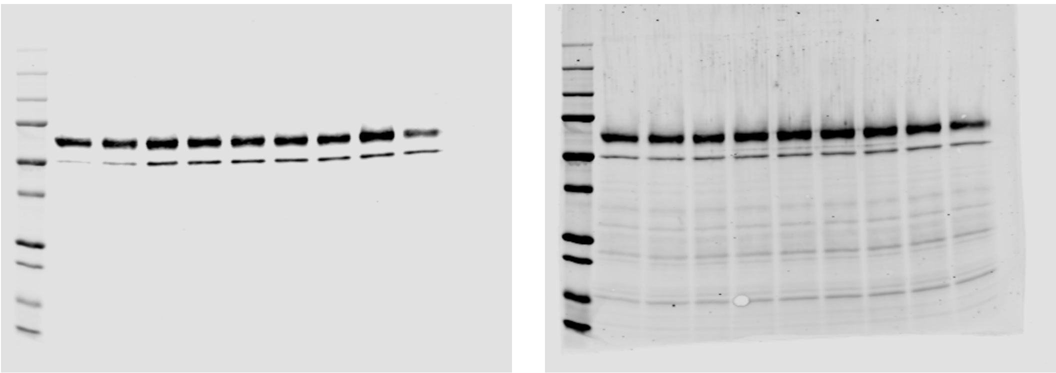

Gel 3, samples 1-9 replicate 3 stained for streptavidin (left) and total protein loading control (right).

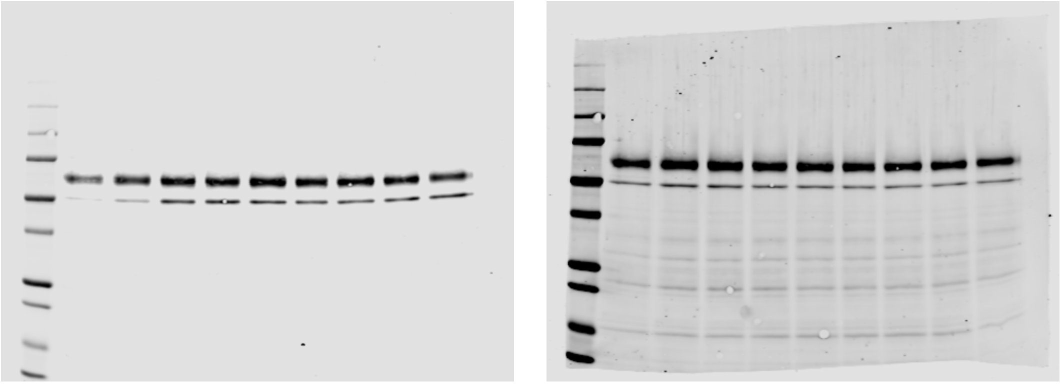

**Fig. S8. Binding affinity profiling data processing workflow**

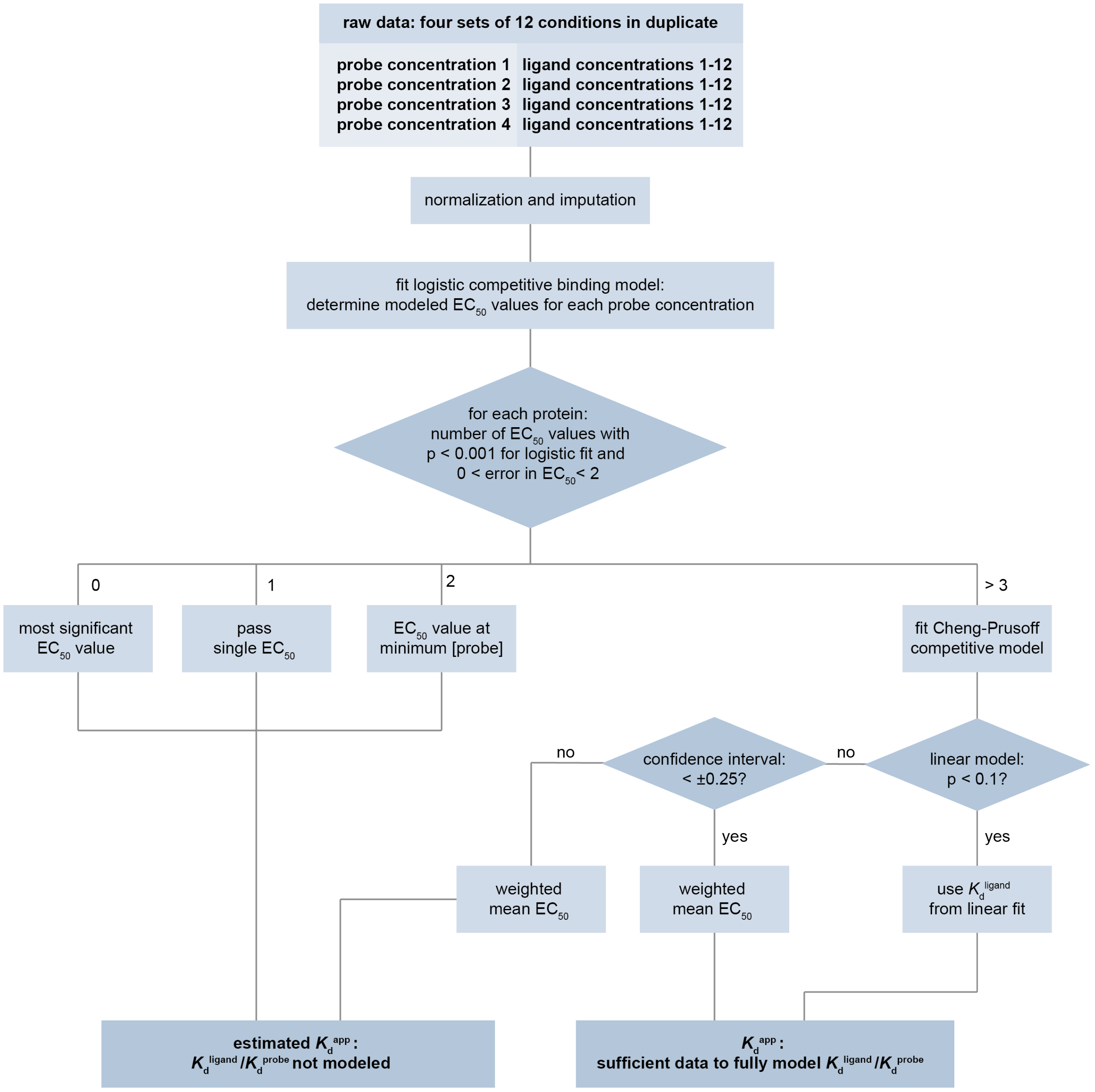

**Fig. S9. Dasatinib-Diazirine-Alkyne Photoaffinity Probe Labeling Optimization**

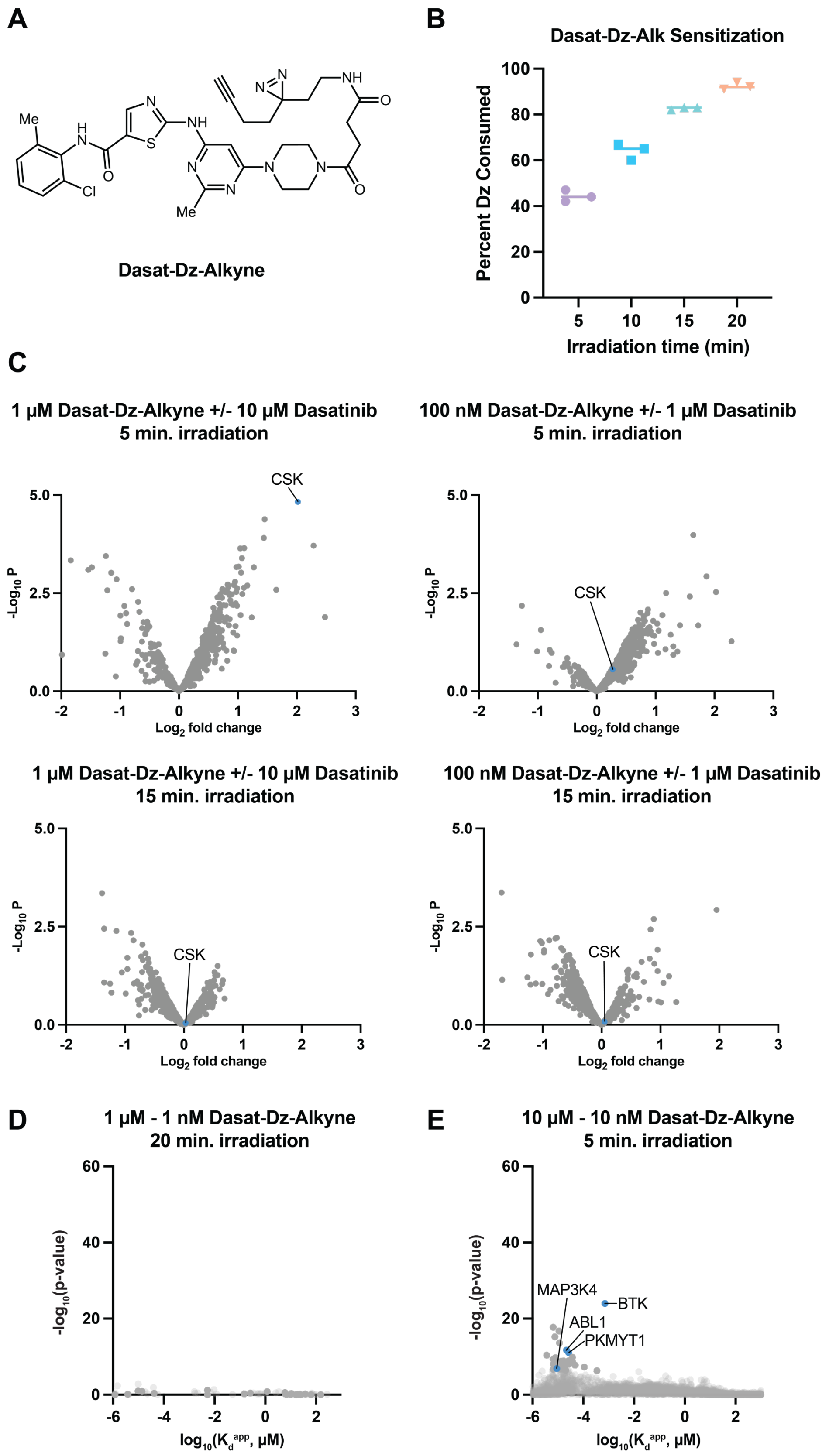

**A.** Structures of Dasatinib-Diazirine-Alkyne photoaffinity probe**. B.** Diazirine probe sensitization time course at 375 nm. Experimental procedures described in method below **C.** Volcano plots for labeling optimization of dasatinib photoaffinity probe irradiation time and concentration with CSK highlighted as a target kinase. CSK was selected to benchmark labeling as it is the only kinase detected in all samples. The Affinity Map in the main text **Fig. 2** and **Fig. S9E** used for comparison to photocatalytic probes was done with 5-minute irradiation at a 10-fold higher starting concentration as this result gave the best enrichment of CSK. Data was collected according to **Method 13** with a two-arm comparison between probe and vehicle versus probe and dasatinib at the indicated concentrations. Each arm of each volcano plot was conducted in triplicate. **D.** Affinity Map performed using Dasatinib-Diazirine-Alkyne probe at concentration identical to photocatalytic probes with 20-minute irradiation when >95% of probe is consumed. Data was obtained following **Method 13** with the concentrations shown below. There is significant measurement of any kinases. **E.** Affinity Map performed with the same probe and conditions optimized for photoaffinity probe. The same map is depicted in **Fig. 2D** and used to compare catalytic and noncatalytic probes. Kinase targets highlighted in blue.

**Sensitization of Das-Dz-alk with UV in Fig S9B:**

200 µL Dasatinib-dz-alkyne and 4,4'-dimethyl-2,2'-bipyridine IS (as a 50 µM stock in PBS) were aliquoted into 8-strip cluster tubes (15 tubes). The samples were irradiated 17.8 cm above two 375 nm 100 W COB LED lights for the appropriate amount of time (**Fig. S20**). The samples were then analyzed by HPLC, and peaks were quantified by absorbance at 325 nm and standardized based off IS. Each measurement was performed in triplicate, comparing against a negative control with no irradiation.

**Probe and off-compete concentrations for the dasatinib-dz-alkyne Affinity Map experiment shown in Fig. S9D**

| Probe [ ] (µM) | Off-compete [ ] (µM) (in duplicate) | Number of samples |
|---|---|---|
| Row A/B : 1 | 0, 10, 3.16, 1, 0.316, 0.1, 0.0316, 0.01, 0.00316, 0.001, 0.000316, 0.0001 | 24 |
| Row C/D : 0.1 | 0, 10, 3.16, 1, 0.316, 0.1, 0.0316, 0.01, 0.00316, 0.001, 0.000316, 0.0001 | 24 |
| Row E/F : 0.01 | 0, 10, 3.16, 1, 0.316, 0.1, 0.0316, 0.01, 0.00316, 0.001, 0.000316, 0.0001 | 24 |
| Row G/H: 0.001 | 0, 10, 3.16, 1, 0.316, 0.1, 0.0316, 0.01, 0.00316, 0.001, 0.000316, 0.0001 | 24 |

**Fig. S10. Competitive photolabeling of BTK by 1, 2, and diazirine-alkyne conjugates of dasatinib in lysate measured via mass spectrometry**

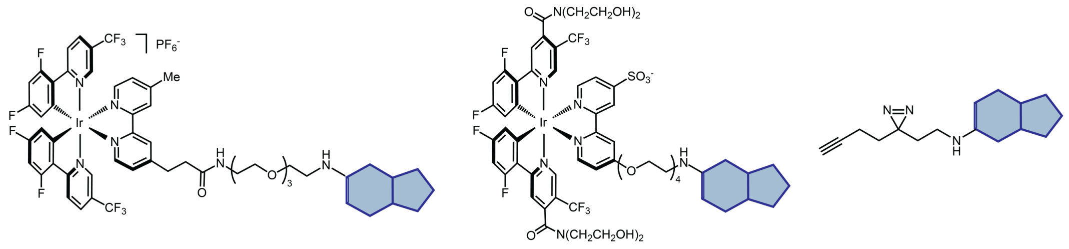

2-Dasatinib    1-Dasatinib    Dasatinib-Dz-Alkyne

**Dose response curves and Schild plot for BTK: 1-Dasatinib**

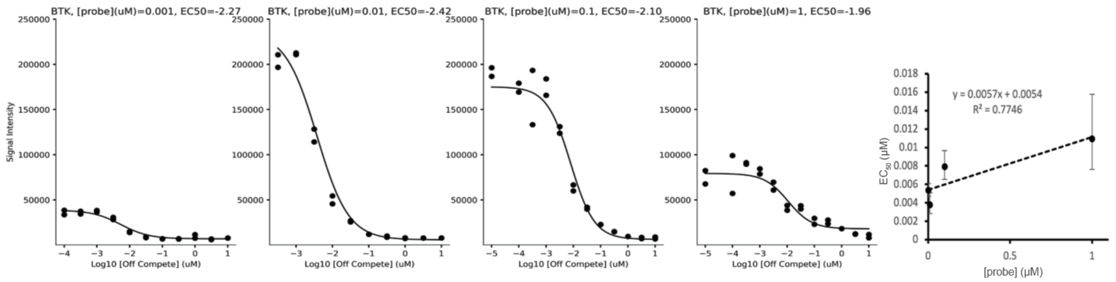

**Dose response curves and Schild plot for BTK: 2-Dasatinib**

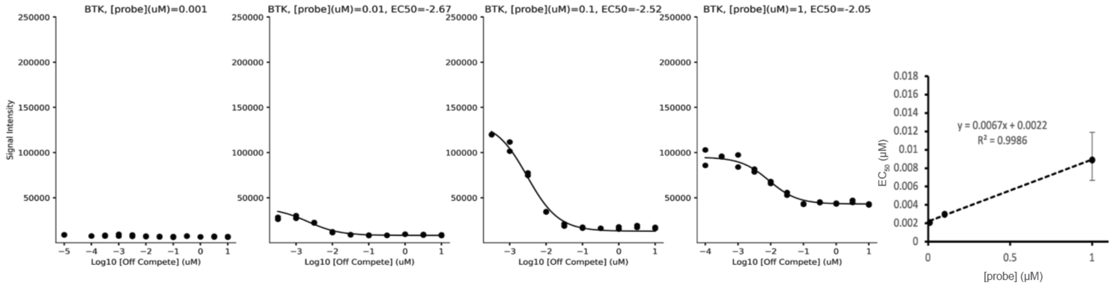

**Dose response curves and Schild plot for BTK: Dasatinib-diazirine-alkyne**

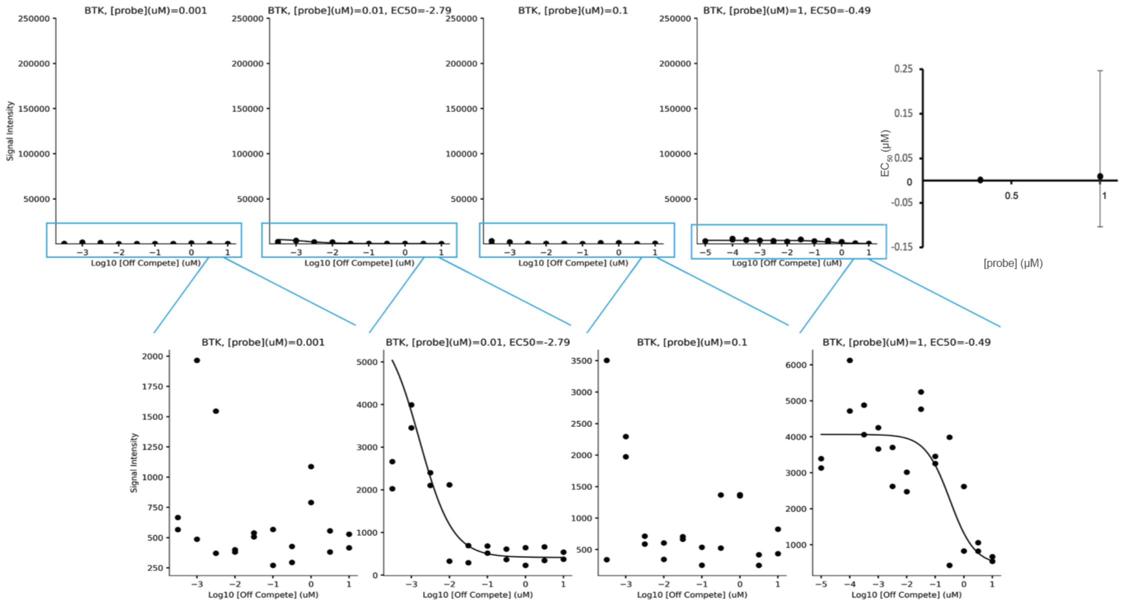

Top: Structures of **1**, **2**, and diazirine-dasatinib conjugates. Bottom: K562 cell lysate was treated with diazirine-biotin at 200 μM, **1**, **2**, or diazirine alkyne conjugates of dasatinib at 1 μM, 100 nM, 10 nM, or 1 nM, and unmodified dasatinib at concentrations ranging from 100 pM to 10 μM in half-log increments in duplicate (96 total samples, **Method 1-2**, **Fig. S18**). After irradiation, protein precipitation, resuspension, reduction/alkylation, IP, and tryptic digestion, peptides were analyzed via nano-UHPLC/IM/MS/MS in DIA mode using a nanoElute 2 / timsTOF Pro 2 MS system and the resulting data processed using DIANN 1.8 with an *in-silico* generated spectral library (**Methods 3-7 and 11**, **Fig. S18**). After imputation (MinProb, MSnbase), and normalization (vsn, MSnbase) within each photocatalyst-dasatinib concentration sample group, integrated intensity data was fit to a logistic model to measure a $EC_{50}$ for the dose-responsive reduction in intensity induced by competitive binding of unmodified dasatinib at each photocatalyst concentration for each protein identified in the proteome assuming a Hill coefficient of 1 for all proteins. These $EC_{50}$ values were then used to generate a Schild plot to model competitive binding between dasatinib and dasatinib-photocatalyst conjugates, with the slope affording the ratio of dasatinib-protein and conjugate-protein $K_d^{app}$ and the intercept affording the $K_d^{app}$ of unmodified dasatinib. **1** performs better than **2**, which gives reduced signal at low concentrations and elevated background signal at high concentrations due to high nonspecific binding, while diazirine-alkyne conjugate of dasatinib yields only weak signal intensity that enables marginal detection of the BTK-dasatinib interaction.

**Fig. S11. Performance of 1, 2, and diazirine-alkyne for binding affinity profiling of dasatinib in cell lysate**

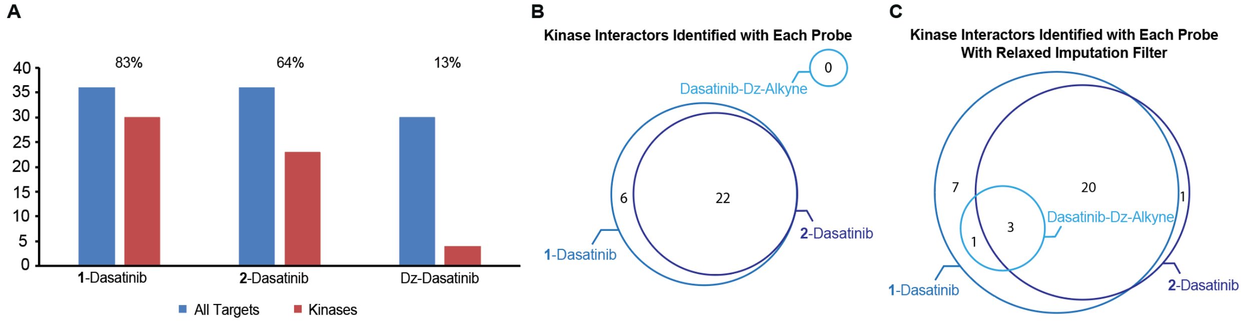

**A.** Proteins and kinases with affinities passing FDR obtained using **1**, **2**, and diazirine-alkyne conjugates of dasatinib in K562 lysate, with the percentage of kinases among identified interactions. For dasatinib-diazirine-alkyne, a relaxed imputation filter was used to enable detection of affinities. **B.** Venn diagram of all kinases identified by each dasatinib conjugate. **C.** Venn diagram of kinases with $K_d^{app}$ identified using a relaxed imputation filter. Relaxed imputation filter refers to constructing competition models on imputed intensity data for a given protein at a given probe concentration if at least one measured value is available, rather than >25% as is typically required for Affinity Map profiles.

**1** afforded the largest number of proteins with significant measured $K_d^{app}$ values, the highest percentage of kinases among measured proteins, and identified all other kinases identified by **2** and diazirine-alkyne. The reduced number of measured $K_d^{app}$ values by **2** reflects its higher nonspecific binding, as demonstrated by lower competitive fold-changes for target proteins (**Fig. S5)**, while the small number of identified values by non-photocatalytic diazirine-alkyne reflects inefficient crosslinking.

**Fig. S12. Comparison of 1 vs. 2 for protein $K_d^{app}$ measurement, benchmarking against literature $K_d^{app}$ values measured using Kinobeads, and comparing log(FC) vs. measured affinity**

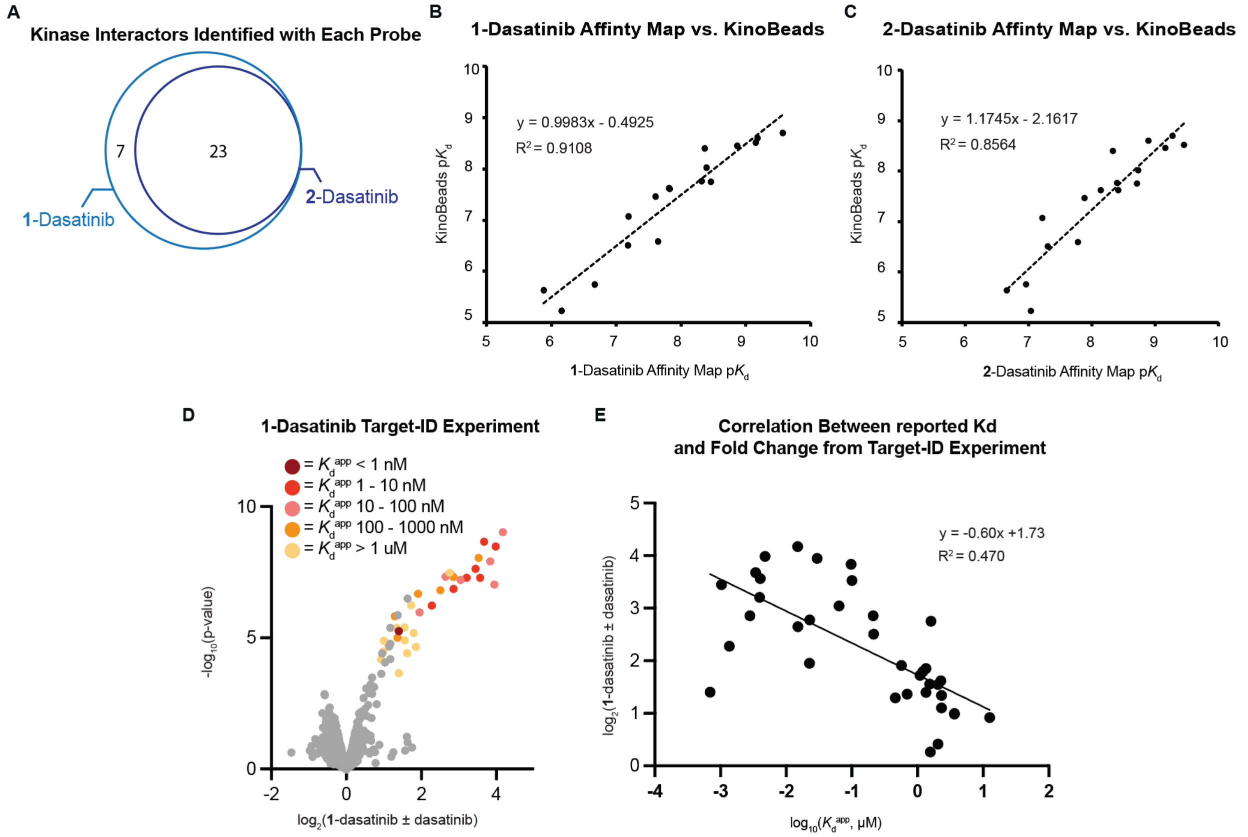

**A.** Venn diagram of protein targets with significant measured $K_d^{app}$ values identified in K562 cell lysate with either **1**-dasatinib or **2**-dasatinib. **B, C.** Comparison of $pK_d^{app}$ values determined by **1**-dasatinib and **2**-dasatinib and published $pK_d^{app}$ values measured using the Kinobeads method. **1** enables the measurement of more kinases than is possible using **2**, and reports slightly more accurate $pK_d^{ap}$ values. Notably, the slope is near one, indicating that photocatalytic binding affinity profiling yields accurate $K_d^{app}$ values [3]. **D.** Volcano plot showing enriched proteins in a two-group comparison ($\log_2$(FC) = $\log_2$(1 µM **1**-dasatinib)-$\log_2$(1 µM **1**-dasatinib + 10 µM dasatinib)) color-coded by Affinity Map-measured binding affinity. **E.** Comparison of $\log_{10}(K_d^{app})$ values determined by probe **1** and fold change measured for enriched proteins in **D**.

**Fig. S13. Benchmarking $K_d^{app}$ estimation workflows for proteins with two significant $EC_{50}$ measurements**

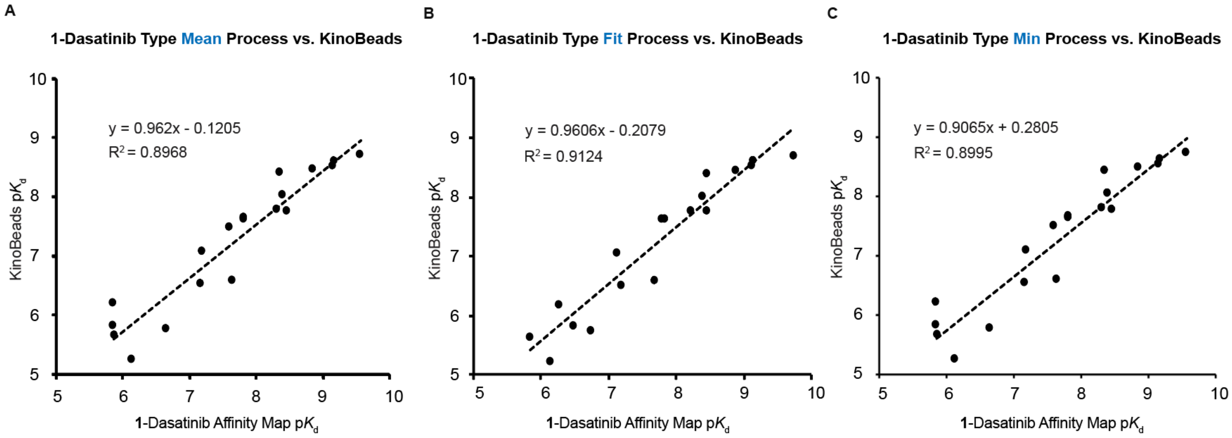

Comparison of how different methods of estimating $K_d^{app}$ perform for proteins that have only two significant $EC_{50}$ determinations, benchmarked against reported $pK_d^{app}$ values measured using the Kinobeads method [3]. As the error in a linear fit cannot be accurately determined using two data points, we benchmarked three methods for extracting estimated $K_d^{app}$ values in such cases using dasatinib binding affinity profiling data generated from K562 cell lysate using **1-**dasatinib. **A.** Error-weighted mean of the two $EC_{50}$ values from Equation 5 reported as $K_d^{app}$. **B.** Fitted $K_d^{app}$ from Equation 4 reported as $K_d^{app}$. TNK2 is excluded from comparison as $pK_d^{app}$ reported is 19.08 due to erroneous results that can arise from two-point linear fitting. **C.** $EC_{50}$ value at the minimum photocatalytic probe concentration reported as $K_d^{app}$. Based on these data, which showed similar performance for all methods, we chose to use the minimum value reporting method as the measurement made at the lowest concentration of the photocatalytic probe is the closest value to the y-intercept of the Schild plot, affording a more accurate estimate than would be produced by a weighted average for the $K_d^{app}$ of proteins that may have comparable affinities for both photocatalyst conjugates and free molecules [3] while avoiding nonphysical results and statistical uncertainties inherent in two-point line fitting.

**Fig. S14. $K_d^{app}$ values from photocatalytic vs. Kinobeads profiling by number of $EC_{50}$ values used for Affinity Map measurement**

| Number of $EC_{50}$ values used for Affinity Map | Average absolute $pK_d^{app}$ difference | Number of proteins |
|---|---|---|
| 4 | 0.501 | 8 |
| 3 | 0.682 | 5 |
| 2 | 0.349 | 3 |
| 1 | 0.192 | 1 |

Relationship between the number of $EC_{50}$ values used for $pK_d^{app}$ determination by **1**-dasatinib in K562 cell lysate and the average $pK_d^{app}$ deviance between measured and literature $pK_d^{app}$ data (Kinobeads) [3].

**Fig. S15. Benchmarking Affinity Map vs. literature Kinobeads, recombinant protein assay, and CETSA datasets**

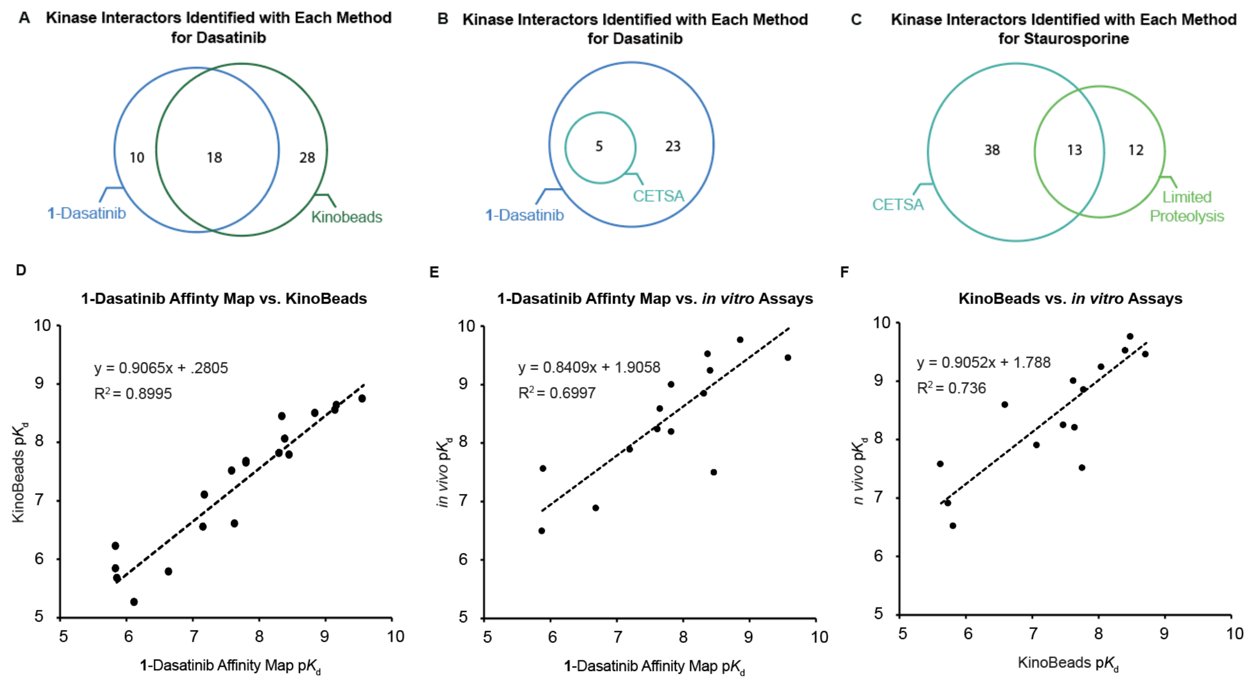

**A.** Venn diagram of protein kinases with significant $K_d^{app}$ measurements made via **1**-dasatinib and protein kinases with significant $K_d^{app}$ measurements obtained in a published dataset generated using the Kinobeads method [4]. **B.** Comparison of protein kinases in K562 lysate with significant measured $K_d^{app}$ values made via **1**-dasatinib and protein kinases with measurable $\Delta T_m$ values from a literature dataset generated using the CETSA method in lysate and live cells [5]. **C.** Comparison of protein kinases in K562 lysate identified by Gao et. al. as interactors of a more promiscuous kinase ligand, staurosporine, using a limited proteolysis (LiP)-based method and reported protein kinases with measurable $\Delta T_m$ values identified via CETSA [5,6]. **D.** Comparison of $pK_d^{app}$ values determined using **1**-dasatinib and reported $pK_d^{app}$ values measured using the Kinobeads method. **E.** Comparison of $pK_d^{app}$ values determined using Affinity Map and **1**-dasatinib and reported $pK_d^{app}$ values measured using recombinant protein assays [7]. **F.** Comparison of reported $pK_d^{app}$ values measured using Kinobeads and reported $pK_d$ values measured by recombinant protein assays [7,8]. In comparison with previously reported data, binding affinity profiling via **1**-dasatinib affords a distinct set of kinases and proteins associated with measurable $K_d^{app}$ values and performs well when compared against assays using activity-based measurement and recombinant proteins. No protein targets of dasatinib identified exclusively by CETSA are kinases. The accuracy of Affinity Map via **1**-dasatinib benchmarked against *in-vitro* assays using activity-based measurements and recombinant proteins is similar to that obtained using Kinobeads.

**Fig. S16. Validation of novel dasatinib-kinase interactions and affinities**

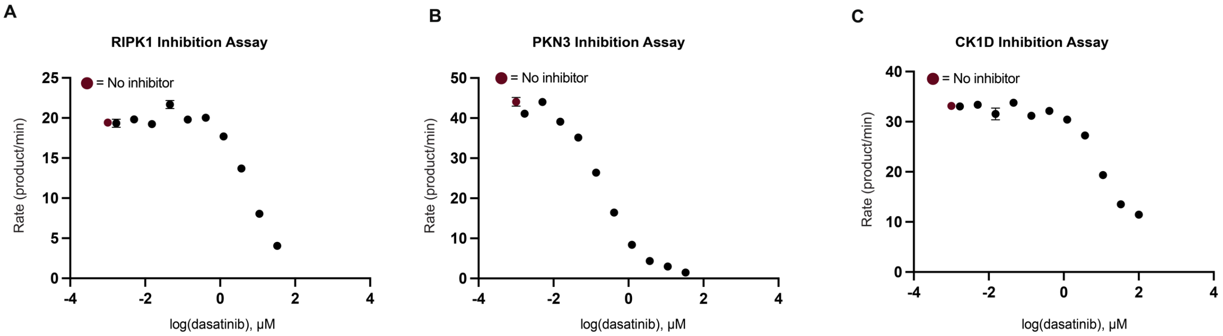

D

| Kinase | ATP $K_m$ | Activity Assay $IC_{50}$ | Activity Assay $K_i$ | 1-das. Affinity Map $K_d$ |
|---|---|---|---|---|
| RIPK1 | 256 µM | 6013 nM | 3007 nM | 3643 nM |
| PKN3 | 36.8 µM | 215 nM | 108 nM | 2311 nM |
| CSNK1D | 14.2 µM | 8362 nM | 4181 nM | 3634 nM |

Activity assay conducted at ATP [ ] = $K_m$ for each kinase
*$K_i$ values estimated derived from Cheng-Prusoff kinetics, assuming ATP-competitive inhibition

**A.** RIPK1 PhosphoSense activity assay dose response curve at [ATP] = $K_m$. **B.** PKN3 PhosphoSense activity assay dose response curve at [ATP] = $K_m$. **C.** CK1D/CSNK1D PhosphoSense activity assay dose response curve at [ATP] = $K_m$. **D.** Summary of PhosphoSense assay $IC_{50}$, ATP $K_m$, calculated $K_i$ value (Cheng-Prusoff), and $K_d^{app}$ values measured using **1**-dasatinib via photocatalytic binding affinity profiling for each kinase.

**AssayQuant Kinase Activity Assay:**
Three kinases, PKN3, CK1D, and RIPK1 were assayed for inhibition by dasatinib using the PhosphoSens CSox-based kinetic assay. Each kinase was treated with either vehicle or dasatinib at 11 concentrations across a three-fold dilution series (final concentration of dasatinib: 100 µM, 33.3 µM, … , 1.69 nM) in technical duplicate. Individual reactions were set up as follows: 11.7 µL reaction mixture (HEPES, Brij-35, EGTA, $MgCl_2$, DTT, ATP, & CSx Substrate), 0.3 µL 50X inhibitor in 100% DMSO, 3.0 uL enzyme dilution buffer (EDB - 20 mM HEPES, pH 7.5, 0.01% Brij-35, 5% Glycerol, 1mM EGTA, 1 mM DTT, 1 mg/ml BSA) or a 5X kinase in EDB. Final reaction volume was 15 µL and final reaction concentrations were the following: 50 mM HEPES, pH 7.5, ATP Km, 1.0 mM DTT, 0.01% Brij-35, 0.5 mM EGTA, 1% glycerol (from EDB), 10 mM $MgCl_2$, 0.20 mg/mL BSA (*from EDB*), 15 µM AQT sensor substrate (AQT0150; AQT0509; AQT0709), 1X kinase (PKN3: 5 nM; CK1D: 1 nM; and RIPK1: 15 nM), and 2% DMSO. Reactions were run at 30 °C for 120 minutes, measured in Perkin Elmer 384-well low volume white ProxiPlates (Cat. #6059480) after sealing using optically clear adhesive film (TopSealA-Plus plate seal, PerkinElmer, Cat. #6050185) in a Biotek Synergy Neo 2 microplate reader with excitation at 360 nm and emission at 485 nm. Rates were fit to the following four parameter logistic equation, and nonlinear regression was performed using the Solver algorithm in Excel [9].

$$Y = Bottom + \frac{Top - Bottom}{1 + 10^{(logIC_{50} - \log[dasatinib]xHill\ Slope}}$$

$K_i$ values were calculated using the Cheng-Prusoff equation, assuming a reversible, substrate-competitive mode of inhibition:

$$K_i = \frac{IC_{50}}{1 + \frac{[S]}{K_m}}$$

**AssayQuant Recombinant Kinase Constructs**
RIPK1, recombinant human protein amino acids (1-327), N-terminal GST tag, SignalChem (Cat/Lot#: R07-11G/ E4165-6). PNK3, full length amino acids (1-889), N-terminal GST tag, Carna Biosciences (Cat/Lot #: 01-146 / 18CBS-0337 B). CK1D (CSNK1D), catalytic domain amino acids (1-294), N-terminal GST tag, Carna Biosciences (Cat/Lot# 03-103 / 09CBS-1197 L).

**Fig. S17. Potential protease cleavage sites/PTM sites in each peptide screened by Affinity Map**

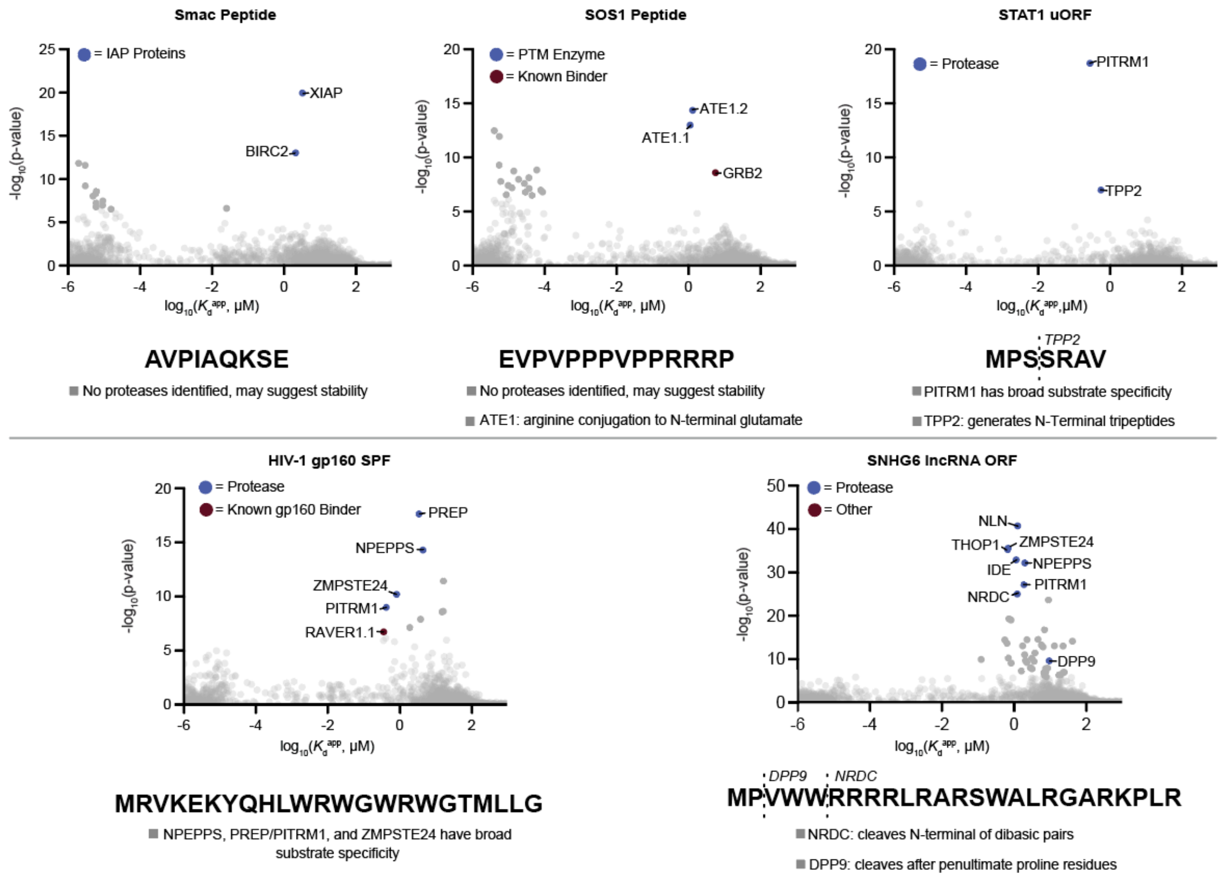

Each peptide has affinity for a unique set of proteases, consistent with known protease substrate specificity. Potential cleavage sites are noted for unique proteases that have affinity for specific peptides [10–16].

### Experimental Methods

##### Fig S18. Sample preparation workflow

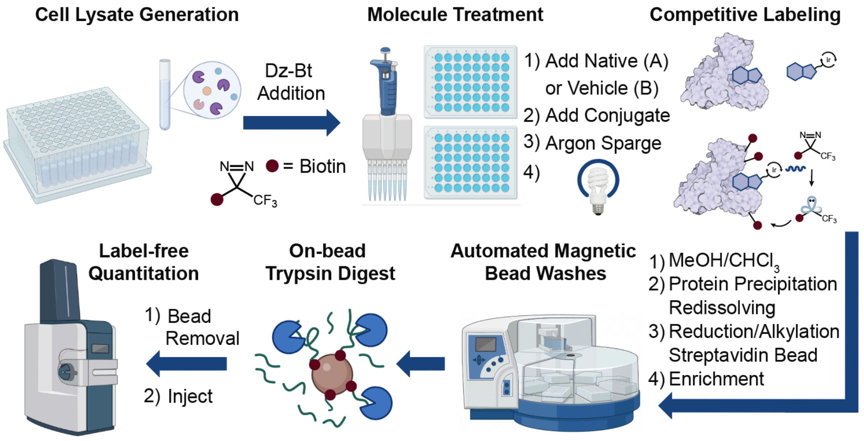

##### Method 1. Pooled cell lysate preparation

For a single 96-sample experiment, 2.2 x $10^8$ K562 cells were collected by centrifugation (800 x g, 5 min, 20 °C), and washed twice with 40 mL of cold PBS. Cells were resuspended in 10.6 mL cold (4 °C) PBS in a 50 mL conical tube and lysed by sonication (QSonica Q500 with cup horn, 4 °C, 3 cycles of 60% amplitude, 4 seconds on/off for 64 seconds). Lysate was examined under a brightfield microscope (1:1 lysate:trypan blue 0.4%) to ensure complete lysis. The lysate was cleared by centrifugation (2,500 x g, 15 min, 4 °C). Concentrated diazrine-PEG$_3$-biotin (Dz-Bt) dissolved in cold PBS (600 µM, 7.04 mL) was added to the cleared lysate and diluted with cold PBS to 21.1 mL to give a final concentration of 200µM Dz-Bt and a 2 mg/mL protein concentration, determined by bicinchoninic acid (BCA) assay. Lysate was kept fresh on ice and was not frozen.

##### Method 2. *In vitro* lysate treatment with probes and free small molecules

200 µL (400 µg protein) of pooled K562 cell lysate was aliquoted into 12 8-strip cluster tubes (96 tubes) in a rack and kept on ice. Preprepared 96-well plates (compound plates) containing 3 µL of a DMSO solution containing appropriate concentrations of compounds (or vehicle DMSO) were used for 1) free small molecule addition and 2) probe addition.

1) All wells in the first compound plate (free small molecules) were diluted with 27 µL room temperature PBS and then 20 µL of each well was added to the correspond cluster tube containing 200 µL of lysate (120-fold dilution) and incubated on ice for 20 minutes.
2) All wells in the second compound plate (probe) were diluted with 34.5 µL room temperature PBS and 20 µL of each well was added to the correspond cluster tube containing 200 µL of lysate (150-fold dilution) and incubated on ice for 20 minutes (final volume 240 µL, final concentrations small molecule/probe tabulated below).

The plate setup presented here and the concentration range used for most molecules in this manuscript (0.1 nM-10 µM ligand, 1 nM – 1 µM probe) is designed to capture affinities in the 1 nM-10 µM range. Based on predicted ligand binding regime a shifted or extended concentration range could be used. For example, for ligands with weaker affinities a higher concentration regime can be used (100 nM-1 mM range for ligand, 1 µM-100 µM for probe). In these experiments, ligand concentration may be increased as high as solubility allows. For the IgG and Dasatinib-diazirine-alkyne experiments (**Methods 13 and 18**) concentrations were increased by an order of magnitude in this way.

**Probe and off-compete small molecule/peptide concentrations in a typical 96-well plate**

| Probe [ ] (µM) | Off-compete [ ] (µM) (in duplicate) | Number of samples |
|---|---|---|
| Row A/B : 1 | 0, 10, 3.16, 1, 0.316, 0.1, 0.0316, 0.01, 0.00316, 0.001, 0.000316, 0.0001 | 24 |
| Row C/D : 0.1 | 0, 10, 3.16, 1, 0.316, 0.1, 0.0316, 0.01, 0.00316, 0.001, 0.000316, 0.0001 | 24 |
| Row E/F : 0.01 | 0, 10, 3.16, 1, 0.316, 0.1, 0.0316, 0.01, 0.00316, 0.001, 0.000316, 0.0001 | 24 |
| Row G/H: 0.001 | 0, 10, 3.16, 1, 0.316, 0.1, 0.0316, 0.01, 0.00316, 0.001, 0.000316, 0.0001 | 24 |

**Fig S19. Small molecule and peptide photocatalytic binding affinity profiling plate setup**

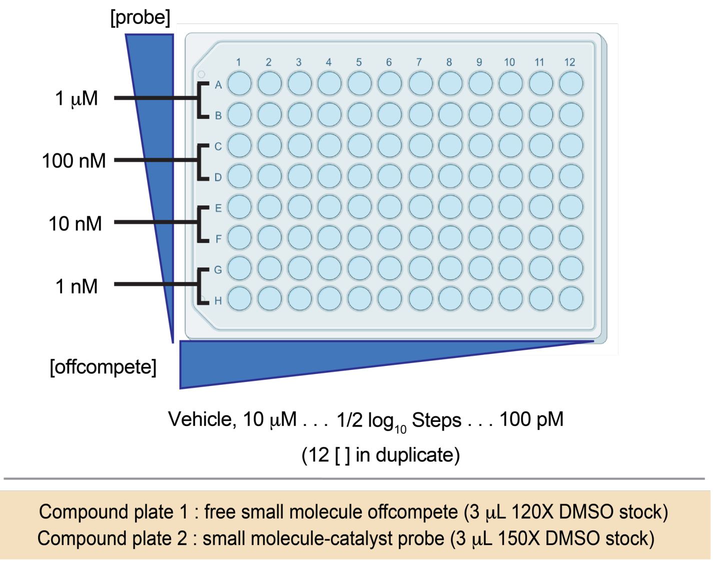

**Method 3. Argon degassing, 440 nm light irradiation, and protein precipitation**

Samples were placed in a vessel that was sealed and purged with water-saturated argon (**Fig. S20**) for 15 minutes. Samples were then irradiated using the apparatus shown below. This setup was shown to achieve uniform irradiation of 96 cluster tubes using ferrioxalate-based actinometry (**Fig. S20**). Immediately following irradiation, 240 µL of -20 °C MeOH and 60 µL of $CHCl_3$ were added to each cluster tube and mixed, precipitating denatured protein. Samples were centrifuged (2,500 x g, 10 min, 4 °C), affording a protein disk suspended between the MeOH and $CHCl_3$ layers. Solvent was carefully aspirated without disrupting the protein disk, then 500 µL of -20 °C MeOH was added to each cluster tube and the protein disk disrupted via sonication (Qsonica Q500 with cup horn, 4 °C, 3 cycles of 60% Amplitude, 4 sec on/off). Samples were centrifuged (2,500 x g, 10 min, 4 °C), resulting in a protein pellet at the bottom of each cluster tube. The MeOH was aspirated, and protein pellets were either carried forward or stored at -80 °C.

**Fig S20. 96-Well plate irradiation apparatus**

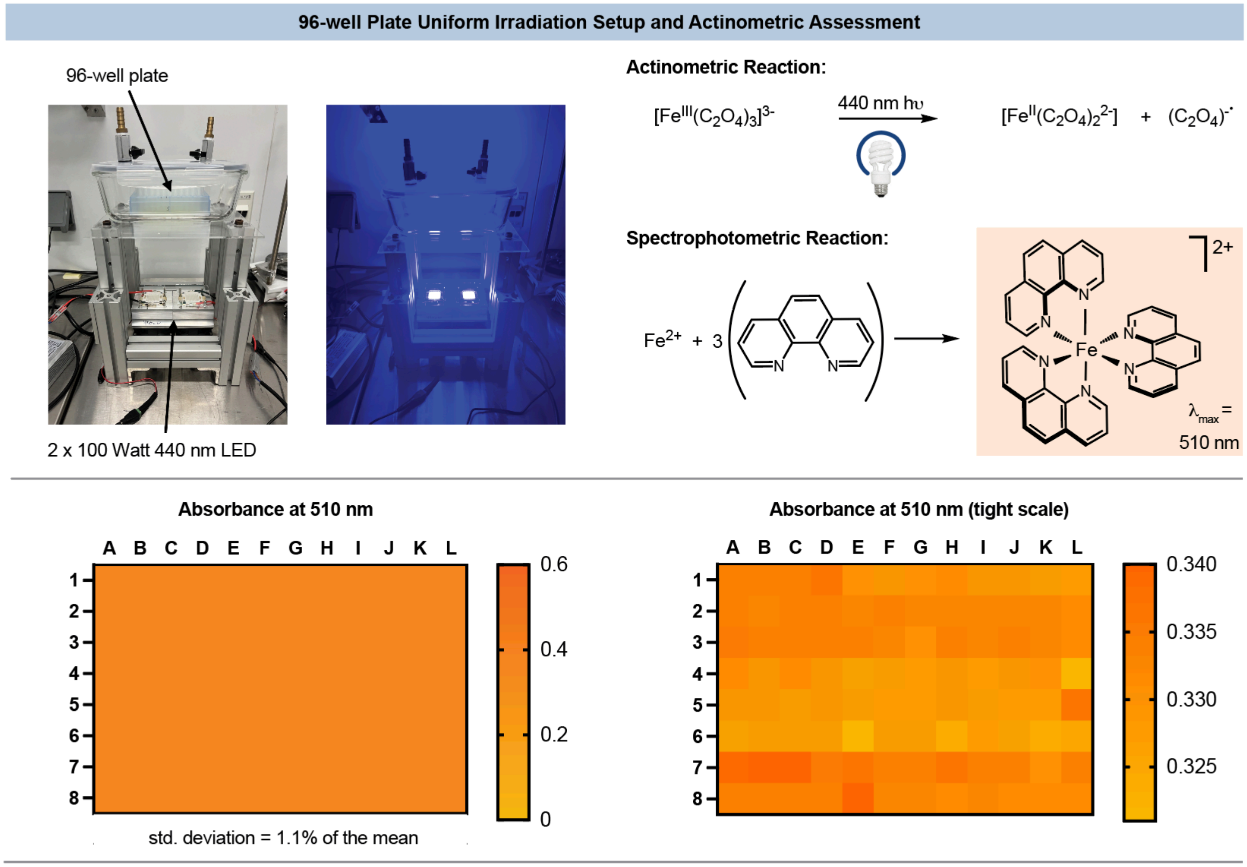

Top: Uniform 96-well plate irradiation setup and ferrioxalate-based actinometry principle. Bottom: a 96-well plate was irradiated and the resulting iron(II) generated by irradiation (actinometric reaction) was measured by iron(II)-phenanthroline complex absorbance at 510 nm. Absorbance values were uniform across the plate, showing a standard deviation that was 1.1% of the mean absorbance, demonstrating even plate irradiation.

**Method 4. Protein pellet redissolving, reduction, alkylation, and streptavidin bead enrichment**
Cluster tubes containing protein pellets were resuspended in 80 uL 600 mM Tris-HCl Buffer with 4% SDS (pH = 8). Samples were boiled at 95 °C for 15 minutes, sonicated in a bath sonicator at room temperature for 10 minutes, boiled again at 95°C for 5 minutes to dissolve the protein pellet, and then 140 uL of MQ $H_2O$ was added to each sample. Dithiothreitol (DTT) in water (50 µL, 54 mM stock, 10 mM final concentration) was added and the mixture incubated at 95 °C for 10 minutes. After the mixture was allowed to cool to room temperature, iodoacetamide in water (50 µL, 128 mM stock, 20 mM final concentration) was added and incubated in the dark at room temperature for 30 minutes. Samples were centrifuged (2,500 x g, 10 min) to ensure any insoluble material was removed, transferred to a 0.5 mL/well 96-well plate, and 50 µL of Sera-Mag$^{TM}$ Medium Capacity Streptavidin beads (Cytiva, 30152104010350) were added to each sample. The plate was sealed with a silicone 96-well plate sealing mat (VWR, 76311-636) and incubated overnight with continuous inversion.

**Method 5. Automated streptavidin bead washing**
Samples were transferred to a KingFisher 96 deep-well plate (Thermo Fisher Scientific 95040450B) (plate 1). Beads were washed using a Qiagen BioSprint 96 workstation (Qiagen 9000852) with KingFisher 96 deep-well plates loaded with four wash buffers: 1) 3 x PBS with 1% SDS washes (plate 2, 3, and 4, 250 µL/well), 2) 3 x PBS with 1 M NaCl (plate 5, 6, and 7, 250 µL/well), 3) 3 x PBS with 10% EtOH (plate 8, 9, 10, 250 µL/well), and 4) 3 x 50 mM ammonium bicarbonate in MQ $H_2O$ (plate, 11, 12, 13, 250 µL/well). The following steps were implemented for each washing buffer (4 rounds with plate 1-4; 4-7; 7-10; and 10-13): (a) first plate (1, 4, 7, or 10) collect beads (premix), (b) plate 2, 5, 8, or 11, wash (release beads, 30 sec, medium; wash beads, 1 min, medium), wash (release beads, 30 sec, fast dual mix; wash beads, 1 min, fast dual mix), wash (release beads, 30 sec, medium; wash beads, 1 min, medium), (c) plate 3, 6, 9, or 12 wash (release beads, 30 sec, medium; wash beads, 1 min, medium), wash (release beads, 30 sec, fast dual mix; wash beads, 1 min, fast dual mix), wash (release beads, 30 sec, medium; wash beads, 1 min, medium), (d) plate 4, 7, 10, or 13, wash (release beads, 30 sec, medium; wash beads, 1 min, medium), wash (release beads, 30 sec, fast dual mix; wash beads, 1 min, fast dual mix), wash (release beads, 30 sec, medium; wash beads, 1 min, medium) (**Fig. S18**). After the final wash, beads from plate 13 were released in a KingFisher 96 deep-well plate loaded with trypsin (0.2 µg/well, Pierce, Thermo part #90058) in 50 mM ammonium bicarbonate (plate 14, 60 µL/well, 3.3 ng/µL final trypsin concentration in 50 mM ammonium bicarbonate). The following steps were implemented: (a) plate 4, collect beads (premix), (b) plate 5, wash (release beads, 30 sec, medium; wash beads, 30 sec, medium), wash (release beads, 30 sec, fast dual mix; wash beads, 30 sec, fast dual mix), wash (release beads, 30 sec, medium; wash beads, 30 sec, medium).

**Method 6. Trypsin digestion**
Beads suspended in trypsin-containing 50 mM ammonium bicarbonate (0.2 µg of trypsin per sample) were transferred to a full-skirted 96-well PCR plate, capped, and incubated at 37 °C overnight with continuous inversion. The plate was centrifuged (200 x g, 20 °C, 1 min) to collect liquid/beads at the bottom of each well, then beads pelleted using a magnetic rack (DynaMag™ 96 Side Skirted Magnet) for 5 minutes. Formic acid (5 µl/well, 5% *v/v* in water) was added to a fresh full-skirted 96-well PCR plate and the supernatant from the trypsin digest was transferred to this plate.

**Method 7. Proteomic data acquisition (standard sensitivity)**
All proteomic samples were analyzed using a Bruker nanoElute 2 / timsTOF Pro 2 nano-UHPLC-IM/MS/MS system using a two-column nano-UHPLC separation method and data independent analysis (DIA) MS method. Acidified and digested samples (5 µL) were loaded on a trap column (Thermo-Fisher, Cat. No. 174500, PepMap Neo C18, 5 µm particle size, 300 µm ID, 5 mm length) using 12 volume equivalents of $H_2O$/0.1% FA. Flow through the trap was then reversed, and peptides eluted through a separation column (Bruker, Cat. No. 1893472, Bruker 10 C18, 1.9 µm particle size, 75 µm ID, 10 cm length) using a 2-35% gradient ($H_2O$/0.1% FA – MeCN/0.1% FA) at a 500 nL/min flow rate. A 20 µm ID CaptiveSpray emitter was used at 1400 V capillary voltage. MS analysis was performed using the instrument default short gradient DIA method in HyStar v. 6.2 without alteration.

**Method 8. In-plate desalting**
An Oasis HLB 96-well plate (Waters) was conditioned by washing once with MeCN+0.1% formic acid (300 µL/well , elution at 200 x g for one minute) and twice with $H_2O$+0.1% formic acid (300 µL/well, elution at 200 x g for 1 min). The tryptic peptide solutions from on-bead digestion were loaded and eluted twice at 200 x g for 1 min. Adsorbed tryptic peptides were washed twice with $H_2O$+0.1% v/v formic acid (300 µL/well, elution at 200 x g for 1 min), then eluted once with 80% MeCN+0.1% v/v formic acid (100 µL/well, elution at 200 x g for 1 min). Volatiles were removed by vacuum concentration, and the samples reconstituted in 40 µL 2% MeCN+0.1% formic acid.

**Method 9. Proteomic data acquisition (high sensitivity)**

Samples were analyzed using a Bruker nanoElute 2 / timsTOF Pro 2 nano-UHPLC-IM/MS/MS system using a two-column nano-UHPLC separation method and data independent analysis (DIA) MS method. Oasis HLB plate desalted samples (5 µL) were loaded on a trap column (Waters nanoEase M/Z Symmetry C18 Trap, 2 cm x 180 µm, 5 µm particle size, Part No.: 186008821) using 12 volume equivalents of H2O/0.1% FA. Flow through the trap was then reversed, and peptides eluted through a separation column with an integrated emitter tip (IonOptiks Aurora Ultimate CSI C18 (25 cm x 75 µm, 1.7 µm particle size, Part No.: AUR3-25075C18-CSI) using a 60-minute 2-35% gradient (H2O/0.1% FA – MeCN/0.1% FA) at 150 nL/min flow rate with 1400 V capillary voltage. MS analysis was performed using the default short gradient DIA method in HyStar v. 6.2 without alteration.

**Method 10. Proteomic data acquisition (high sensitivity, 90 minute gradient)**

Samples were analyzed using a Bruker nanoElute 2 / timsTOF Pro 2 LC/IM/$MS^2$ system using a two-column nano-UHPLC separation method and data independent analysis (DIA) MS method. Oasis HLB plate desalted samples (5 µL) were loaded on a trap column (Waters nanoEase M/Z Symmetry C18 Trap, 2 cm x 180 µm, 5 µm particle size, Part No.: 186008821) using 12 volume equivalents of $H_2O$/0.1% FA. Flow through the trap was then reversed, and peptides eluted through a separation column with an integrated emitter tip (IonOptiks Aurora™ Ultimate CSI C18 (25 cm x 75 µm, 1.7 µm particle size, Part No.: AUR3-25075C18-CSI) using a 90-minute 2-35% gradient ($H_2O$/0.1% FA – MeCN/0.1% FA) at 150 nL/min flow rate with 1400 V capillary voltage. MS analysis was performed using the default short gradient DIA method in HyStar v. 6.2 without alteration.

**Method 11. Proteomics data independent analysis using DIA-NN**

Data was analyzed using DIA-NN 1.8. A Human FASTA sequence database from UniProt (accession date: 12-05-2022) was used. A trypsin-digested *in silico* spectral library generated in DIA-NN was used for analysis. DIA-NN settings: MS1 accuracy and mass accuracy tolerance: ±12.5 ppm; missed cleavages: 1; maximum number of variable modifications: 1; peptide length: 7-30; precursor charge range: 1-4; precursor m/z range: 300-1800; fragment ion m/z range: 200-1800; modifications: C carbamidomethylation, N-term M excision, Ox(M), Ac(N-term); Precursor FDR (%) = 1.0; match between runs enabled, neural network classifier: double-pass mode; protein inference: genes; quantification strategy: robust LC (high precision); cross-run normalization: off; library-generation: smart profiling.

**Method 12. Cell culture (K562 and THP-1 cells)**

K-562 (CCL-243) and THP-1 (TIB-202) cells were purchased from ATCC. K-562 and THP-1 cells were cultured at 37 °C and 5% CO2 in RPMI 1640 complete medium (Gibco) supplemented with 10% fetal bovine serum (Gibco) and penicillin-streptomycin (100 U/mL final concentration, Gibco). Subculturing and medium renewal were performed according to ATCC's product sheets for each cell line. Unless otherwise noted, cells were grown in 25 $cm^2$, 75 $cm^2$ or 150 $cm^2$ canted neck, vented cap sterile cell culture flasks.

**Method 13. Dasatinib-diazirine-alkyne labeling**

K562 cell lysate was prepared according to **Method 1**. Lysate (0.2 mL, 2 mg/mL) was transferred to cluster tubes and treated with compounds as described in **Method 2** (using dasatinib-dz-alkyne instead of **1**-dasatinib at the concentrations shown) and irradiated using the same apparatus as **Fig. S20** with 100-watt 375 nm LEDs for 5 minutes. 34 µL of 7x protease inhibitor cocktail (Roche, 11836170001) was then added, and biotin-$PEG_4$-azide (2.74 µL, 10 mM in DMSO), $CuSO_4$ (1.37 µL, 50 mM in $H_2O$), THPTA (6.85 µL of 10 mM in $H_2O$), and sodium ascorbate (13.7 µL, 50 mM in $H_2O$, freshly made) added, in that order. Note that order of addition is important, adding the first three reagents together as a cocktail and sodium ascorbate last. Samples were inverted end over end for 1 hour at room temperature. Then 280 µL of -20 °C MeOH and 70 µL of $CHCl_3$ were added to each cluster tube and mixed, precipitating denatured protein. Samples were centrifuged (2,500 x g, 10 min, 4 °C), affording a protein disk suspended between the MeOH and $CHCl_3$ layers. Solvent was carefully aspirated without disrupting the protein disk, then 500 µL of -20 °C MeOH was added to each cluster tube and the protein disk disrupted via sonication (Qsonica Q500 with cup horn, 4 °C, 3 cycles of 60% Amplitude, 4 sec on/off). Samples were centrifuged (2,500 x g, 10 min, 4 °C), resulting in a protein pellet at the bottom of each cluster tube. The MeOH was aspirated, and protein pellets were stored at -80 °C overnight then processed and analyzed using **Methods 4**-**7** and **11**.

**Probe and off-compete concentrations for the dasatinib-dz-alkyne Affinity Map experiment**

| Probe [ ] (μM) | Off-compete [ ] (μM) (in duplicate) | Number of samples |
|---|---|---|
| Row A/B : 10 | 0, 100, 31.6, 10, 3.16, 1, 0.316, 0.1, 0.0316, 0.01, 0.00316, 0.001 | 24 |
| Row C/D : 1 | 0, 100, 31.6, 10, 3.16, 1, 0.316, 0.1, 0.0316, 0.01, 0.00316, 0.001 | 24 |
| Row E/F : 0.1 | 0, 10, 3.16, 1, 0.316, 0.1, 0.0316, 0.01, 0.00316, 0.001, 0.000316, 0.0001 | 24 |
| Row G/H: 0.01 | 0, 10, 3.16, 1, 0.316, 0.1, 0.0316, 0.01, 0.00316, 0.001, 0.000316, 0.0001 | 24 |

**Method 14. Preparation of liver S9 fraction for (+)-JQ1 Affinity Map**
Frozen, mixed gender pooled liver S9 fractions were purchased from BioIVT (X008011). The purchased samples contained 40 mg of protein total, which was diluted to ~2 mg/mL with PBS containing 200 μM diazrine-PEG$_3$-biotin (Dz-Bt) and was aliquoted across 96 cluster tubes (200 μL homogenate, 400 μg protein /cluster tube). At this point, for compound treatment method **2** was followed. The samples were then processed using methods **3**-**7**, and **11**.

**Method 15. Cell culture (U87MG cells)**
U87MG cells (HTB-14) were purchased from ATCC. Cells were cultured at 37 °C and 5% CO2 in DMEM 1640 complete medium (Gibco) supplemented with 10% fetal bovine serum (Gibco) and penicillin-streptomycin (100 U/mL final concentration, Gibco). Subculturing and medium renewal were performed according to ATCC's product sheets for each cell line. Unless otherwise noted, cells were grown in 10 cm or 15 cm diameter cell culture treated plates.

**Method 16. Live U87MG cell treatment with 1-cRGDfK and free cRGDfK and labeling with 440 nm light irradiation**
15 cm plates with U87MG cells were washed once with DPBS (10 mL), detached by incubation with TrypLE™ Express Enzyme (Gibco, 5 mL, 5 min, 37 °C), and collected by centrifugation (800 x g, 5 min, 20 °C). Cells were resuspended in DMEM 1640 complete medium (Gibco) supplemented with 10% fetal bovine serum (Gibco) and penicillin-streptomycin (100 U/mL final concentration, Gibco) and passed through cell strainers to help dissociate clumping cells. 4 x 6-well treated cell culture plates were seeded with 5 x $10^5$ U87MG cells and allowed to grow until confluent (~3 days). The media was aspirated and 400 μL of phenol red-free media containing 200 μM diazrine-PEG$_3$-biotin (Dz-Bt) was pipetted over the adherent cells in each well. The appropriate concentration of unconjugated cRGDfK was added (40 μL) or a vehicle control to each well and cells incubated on ice for 20 minutes. Cells were then treated with the appropriate concentration of **1**-cRGDfK (40 μL) and incubated on ice for 20 minutes (final volume 240 μL, 160 μM Dz-Bt, final concentrations free cRGDFk/probe tabulated below). Each 6-well plate was then irradiated for 15-minutes at 4 °C. Following irradiation, cells were released from the plate with 1 mL of cold PBS, transferred to a 1.5 mL tube, and pelleted by centrifugation (800 x g, 5 min, 4 °C). The supernatant was aspirated, and cell pellets were frozen at -80 °C. This experiment was repeated for each concentration of **1**-cRGDfK (1 μM, 100 nM, 10 nM, or 1 nM), thus four separate groups of four plates were seeded then treated on separate days (4 concentrations x 4 plates x 6 wells = 96 wells total).

**Probe and off-compete concentrations for cRGDfK affinity profiling**

| Probe [ ] (μM) | Off-compete [ ] (μM) (in duplicate) | Number of samples |
|---|---|---|
| Plate 1 (day 1): 1 | 0, 100, 31.6, 10, 3.16, 1, 0.316, 0.1, 0.0316, 0.01, 0.00316, 0.001 | 24 |
| Plate 2 (day 2): 0.1 | 0, 100, 31.6, 10, 3.16, 1, 0.316, 0.1, 0.0316, 0.01, 0.00316, 0.001 | 24 |
| Plate 3 (day 3): 0.01 | 0, 100, 31.6, 10, 3.16, 1, 0.316, 0.1, 0.0316, 0.01, 0.00316, 0.001 | 24 |
| Plate 4 (day 4): 0.001 | 0, 100, 31.6, 10, 3.16, 1, 0.316, 0.1, 0.0316, 0.01, 0.00316, 0.001 | 24 |

**Method 17. U87MG cell lysis, reduction, alkylation, and streptavidin enrichment**
Cell pellets were thawed for 1 hour at 4 °C before resuspension in RIPA lysis buffer containing 1% SDS and protease inhibitor cocktail (200 μL). Cells were incubated in RIPA buffer for 30-minutes at room temperature followed by sonication (QSonica Q500 with cup horn, 4 °C, 4 cycles of 60% amplitude, 4 seconds on/off for 64 seconds). Immediately following lysis, insoluble material was pelleted (800 x g, 5 min, 4 °C) and the supernatant was transferred to cluster tubes. Dithiothreitol (DTT) in water (50 μL, 50 mM stock, 10 mM final concentration) was added and the mixture incubated at 95 °C for 10 minutes. After the mixture was allowed to cool to room temperature, iodoacetamide in water (50 μL, 120 mM stock, 20 mM final concentration) was added and incubated at in the dark at room temperature for 30 minutes. Samples were centrifuged once more (2,500 x g, 10 min), to ensure any insoluble material was removed, transferred to a 0.5 mL/well 96-well plate, and 50 μL of Sera-Mag$^{TM}$ Medium Capacity Streptavidin beads (Cytiva, 30152104010350) were added to each sample. The plate was sealed with a silicone 96-well plate sealing mat (VWR, 76311-636) and incubated overnight with continuous inversion. Sample were then processed using methods **5, 6, 8, 10,** and **11**.

**Method 18. Conjugation of 1 to human serum-derived IgG**
**1**-DBCO (120 μL, 5 mM in DMSO) and azido-PEG$_3$-NHS ester (BroadPharm, BP-21605, 6 μL, 100 mM in DMSO) were mixed and allowed to react for 10 minutes. Purified IgG (Sigma Aldrich, I4506, 300 μL, 20 mg/mL in PBS) was mixed with $NaHCO_3$ (30 μL, 1M in $H_2O$) and the click reaction mixture (120 uL, 4.76 mM in DMSO) and allowed to react for 2 hours in the dark. The resulting iridium labeled IgG was desalted twice (to ensure no free catalyst remained) using Thermo Scientific$^{TM}$ Zeba$^{TM}$ Spin Desalting Columns, 40K MWCO (catalog no. PIA57759) that had been equilibrated with PBS. The desalted mixture was analyzed by BCA for protein concentration (50:1 solution A:B; A: 10 mg/mL bicinchonic acid, 20 mg/mL $Na_2CO_3$, 1.6 mg/mL sodium tartarate, 4 mg/mL NaOH, 9.5 mg/mL $NaHCO_3$. B: 40 mg/mL $CuSO_4$) compared to a BSA standard. Photocatalyst concentration was measured by fluorescence of the sample at 485 nm following excitation at 380 nm compared to a standard of **1**. An extent of labeling of 4 **1**/IgG was obtained.

**Method 19. Treatment of live THP-1 cells with 1-IgG and free IgG, labeling with 440 nm light irradiation**
2.1 x $10^8$ THP-1 cells were collected by centrifugation (800 x g, 5 min, 20 °C), and washed twice with 40 mL of cold PBS. Cells were resuspended in concentrated diazrine-PEG$_3$-biotin (Dz-Bt) dissolved in cold PBS (570 μM, 2.2 mL) and aliquoted into 12 8-strip cluster tubes (96 tubes) in a rack and kept on ice (21 μL / aliquot, approximately 400 μg of protein after lysis). The appropriate concentration of free IgG was added (29 μL) or a vehicle PBS control to each cluster tube and incubated on ice for 20 minutes. Cells were then treated with the appropriate concentration of IgG-Ir (10 μL / aliquot) and incubated on ice for 20 minutes (final volume 60 μL, 200 μM Dz-Bt, final concentrations free IgG/probe tabulated below). Samples were then irradiated for 15 minutes at 4 °C. Following irradiation, 200 μL of cold PBS was added to each cluster tube and cells were pelleted by centrifugation (800 x g, 5 min, 4 °C). The supernatant was aspirated, and cells were pelleted and washed with another 200 μL of cold PBS. Cell pellets were frozen at -80 °C.

**Probe and off-compete concentrations for the IgG Affinity Map experiment**

| Probe [ ] (µM) | Off-compete [ ] (µM) (in duplicate) | Number of samples |
|---|---|---|
| Row A/B : 10 | 0, 100, 31.6, 10, 3.16, 1, 0.316, 0.1, 0.0316, 0.01, 0.00316, 0.001 | 24 |
| Row C/D : 1 | 0, 100, 31.6, 10, 3.16, 1, 0.316, 0.1, 0.0316, 0.01, 0.00316, 0.001 | 24 |
| Row E/F : 0.1 | 0, 100, 31.6, 10, 3.16, 1, 0.316, 0.1, 0.0316, 0.01, 0.00316, 0.001 | 24 |
| Row G/H: 0.01 | 0, 100, 31.6, 10, 3.16, 1, 0.316, 0.1, 0.0316, 0.01, 0.00316, 0.001 | 24 |

**Fig S21. IgG Affinity Profiling Plate Setup**

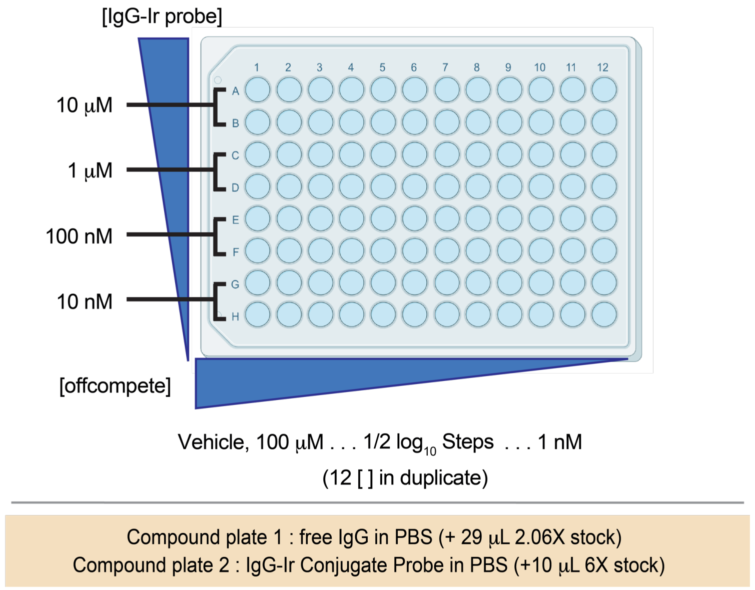

**Method 20. THP-1 cell lysis**

Cell pellets were thawed for 1 hour at 4 °C before resuspension in RIPA lysis buffer containing 1% SDS and protease inhibitor cocktail (100 μL). Cells were incubated in RIPA buffer for 30-minutes at room temperature followed by sonication (QSonica Q500 with cup horn, 4 °C, 4 cycles of 60% amplitude, 4 seconds on/off for 64 seconds). Immediately following lysis, 140 μL of additional $H_2O$ was added to each cluster tube (final volume 240 μL) and $MeOH/CHCl_3$ precipitation was carried out according to method **3**. The samples were then processed using methods **4, 5, 6, 8, 9,** and **11**.

**Method 21. Ferrioxalate actinometry for assessment of plate-based 440 nm irradiation**

Based on a modified literature procedure [17], 240 μL of actinometric solution containing potassium ferrioxalate (200 mM $H_2SO_4$, 20 mM $K_3Fe(C_2O_4)_3$ in $H_2O$) was aliquoted into 12 8-strip cluster tubes (96 tubes) in a rack and irradiated with 440 nm light using the plate irradiation apparatus (**Fig. S20**) for 30 seconds. During the irradiation, 187 uL of spectrophotometric analysis solution (300 mM AcONa, 130 mM $H_2SO_4$, and 277 μM phenanthroline) was added to a 96-well plate. Following irradiation, the actinometric solution was diluted with 760 μL $H_2O$ and 25 μL of this diluted solution transferred to the plate containing the spectrophotometric solution. After the solutions were mixed, the plate was incubated in the dark at room temperature for 1 hour to equilibrate. Absorbance of each individual well was read at 510 nm to detect the light induced formation of iron(II) phenanthroline complexes and assess the evenness of plate irradiation.

**Fig S22. Dasatinib Affinity Map with extended pre-irradiation incubation affords similar $K_d^{app}$ values**

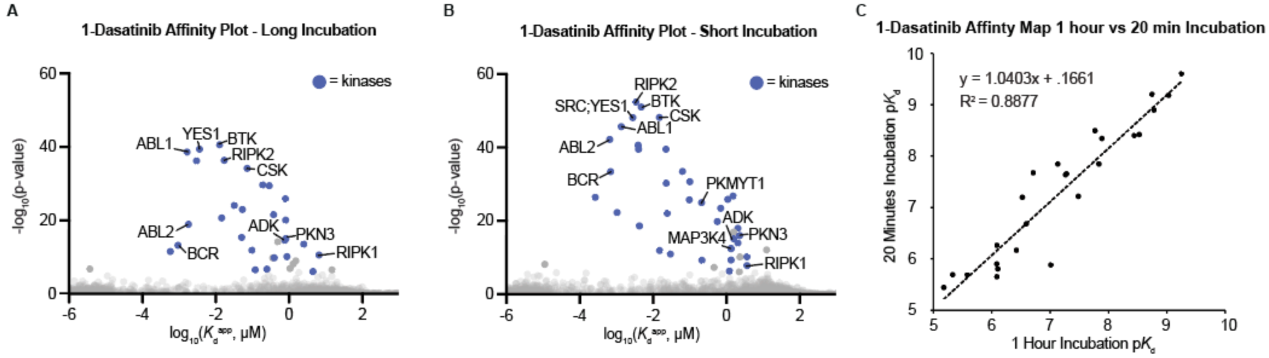

**A.** Dasatinib affinity profile in K562 lysate where cell lysate was incubated for 1 hour with dasatinib then 1 hour with **1**-dasatinib. **B.** Dasatinib affinity profile in K562 lysate where cell lysate was incubated for 20 minutes with dasatinib then 20 minutes with **1**-dasatinib. **C.** Comparison of p$K_d^{app}$ values determined by either long incubation shown in A or short incubation shown in B. The detected p$K_d^{app}$ is similar at 1 hour and 20-minute incubation timepoints, although a reduction in signal intensity was observed with longer incubation time (potentially due to protein degradation). All reported affinity measurements represent apparent binding affinities ($K_d^{app}$), because as the extent of dasatinib-target equilibration is not assessed, very slow on or off rates for ligand association/dissociation are possible.

**Method 22. Affinity profiling dasatinib with longer incubation time**

K562 cell lysate was prepared according to **Method 1**. Lysate (0.2 mL, 4 mg/mL) was transferred to cluster tubes and treated with compounds as described in **Method 2,** but incubation periods were extended to 1 hour instead of 20 minutes for each compound treatment. Sample were irradiated, prepped, and subjected to mass spectrometry according to **Methods 3-7**, and analyzed using **Method 11**.

**Method 23. Pooled membrane preparation from whole mouse brains (Flumazenil Affinity Map)**

Based on a modified literature procedure [18], 3 male CD-1 (ICR) mouse brains (BioIVT, MSE00BRAIN-0102904) were thawed on ice for 25 minutes then resuspended in 10 volumes ice cold lysis buffer (50 mM Tris-HCl pH 7.4, containing protease inhibitor cocktail - Roche, 11836170001, approximately 5 mL per brain). The organs were homogenized (Bio-Gen PRO200 Homogenizer) on ice using at medium intensity (6 pulses x 10 seconds). As an initial clearing step, the homogenate was centrifuged (1,000 x g for 10 min at 4 °C) to obtain supernatant. The supernatant was transferred to ultracentrifuge tubes and centrifuged (40,000 x g for 20 minutes at 4 °C; Beckman SW 32 Ti swinging bucket rotor). The supernatant was decanted and discarded and replaced with fresh ice-cold lysis buffer. The pellet was resuspended by homogenization (5 pulses x 10 seconds) and the centrifugation-homogenization was repeated 2 additional times for a total of 3 ultracentrifugations to ensure complete homogenization and wash out of endogenous ligands (specifically GABA). The final pellet was resuspended in a minimal amount of ice-cold protease-inhibitor-free lysis buffer and protein concentration was determined by bicinchoninic acid (BCA) assay. The freshly prepared membrane suspension was never frozen and used as prepared. Concentrated diazrine-PEG$_3$-biotin (Dz-Bt) dissolved in cold Tris-HCl was added for a final concentration of 200 µM and a 1 mg/mL protein concentration. The membrane suspension was then aliquoted across 96 cluster tubes (200 µL, 200 µg per sample). Compound treatment and sample processing was then performed according to **methods 2-6, 8, 24, and 11.**

**Method 24. Proteomic data acquisition (medium sensitivity)**

Samples were analyzed using a Bruker nanoElute 2 / timsTOF Pro 2 nano-UHPLC-IM/MS/MS system using a two-column nano-UHPLC separation method and data independent analysis (DIA) MS method. Oasis HLB plate desalted samples (5 µL) were loaded on a trap column (Waters nanoEase M/Z Symmetry C18 Trap, 2 cm x 180 µm, 5 µm particle size, Part No.: 186008821) using 12 volume equivalents of H2O/0.1% FA. Flow through the trap was then reversed, and peptides eluted through a separation column with an integrated emitter tip (IonOptiks Aurora Ultimate CSI C18 (15 cm x 75 µm, 1.7 µm particle size, Part No.: AUR3-15075C18-CSI) using a 40-minute 2-35% gradient (H2O/0.1% FA – MeCN/0.1% FA) at 150 nL/min flow rate with 1400 V capillary voltage. MS analysis was performed using the default short gradient DIA method in HyStar v. 6.2 without alteration.

**1-Flumazenil probe and flumazenil off-compete concentrations in a 96-well plate**

| Probe [ ] (μM) | Off-compete [ ] (μM) (in duplicate) | Number of samples |
|---|---|---|
| Row A/B : 1 | 0, 10, 3.16, 1, 0.316, 0.1, 0.0316, 0.01, 0.00316, 0.001, 0.000316, 0.0001 | 24 |
| Row C/D : 0.1 | 0, 10, 3.16, 1, 0.316, 0.1, 0.0316, 0.01, 0.00316, 0.001, 0.000316, 0.0001 | 24 |
| Row E/F : 0.01 | 0, 10, 3.16, 1, 0.316, 0.1, 0.0316, 0.01, 0.00316, 0.001, 0.000316, 0.0001 | 24 |
| Row G/H: 0.001 | 0, 10, 3.16, 1, 0.316, 0.1, 0.0316, 0.01, 0.00316, 0.001, 0.000316, 0.0001 | 24 |

#### Computational Methods – Affinity Surveyor

##### Peptide identification and quantification – DIA

Peptide identification and quantification were performed using DIANN 1.8 or 1.9 [19,20] using a library generated *in-silico* from the human proteome (Uniprot, 12/05/2022). Digestion was set as "Trypsin/P" (K/R cuts), a maximum of one missed cleavage was allowed, peptide length set to 7-30, charge set to 1-4, precursor mass range set to 300-1800, and using a fragment ion m/z range of 200-1800. Modifications were considered as variable for methionine oxidation, acetylation of N-terminus, N-terminal methionine excision, and cysteine carbamidomethylation. The precursor false discovery rate was set to 1%, mass accuracy and MS1 accuracy to 12.5 ppm, and scan window set to 0 with likely inferences removed. Isotopologues and match between runs were enabled. Searches were conducted in double-pass mode using robust, high precision liquid chromatography quantification strategy, protein inference set to genes, library generation set to smart profiling, speed and RAM usage set to optimal results, and with cross-run normalization disabled.

##### Preprocessing

Preprocessing of DIANN output files (report.pg_matrix.tsv, report.pr_matrix.tsv) was done in R (4.4.1) using the MSnbase package separately for each set of samples treated with a given concentration of photocatalyst conjugate. Proteomics data was subjected to variance stabilization normalization ('vsn') and left-censored missing value imputation ('MinProb') using default settings [21,22], and the number of peptides used to quantify each protein determined from the peptide-level report and then added as metadata. Once complete, individually processed MSnSet datasets were combined into a single table.

##### Affinity profiling.

Our approach for apparent (non-equilibrium) binding affinity ($K_d^{app}$) elucidation is analogous to a proteome-wide radioligand competitive binding assay, in which streptavidin IP/MS intensity is used as an indirect readout of competitive protein binding site occupancy by a photocatalyst conjugate probe and a competing ligand molecule with distinct $K_d^{app}$ values ($K_d^{ligand}$ and $K_d^{probe}$). As photocatalytic proximity labeling measured by mass spectrometry is more complex than a simple radioactivity readout corresponding to receptor occupancy, two assumptions are made to constrain the kinetic model: 1) the MS intensity difference of proteins obtained from streptavidin/biotin IP induced by photocatalyst off-compete by a non-photocatalytic ligand is directly proportional to the occupancy of a single binding site by the photocatalyst conjugate, and 2) protein labeling does not materially alter affinity for the probe or ligand molecules. With these constraints in place, a rearranged Cheng-Prusoff equation can be used to independently solve for the $K_d^{app}$ values for each molecule [23] using protein intensities derived from photocatalytic labeling experiments where the concentrations of both the probe and ligand molecules are systematically varied. All reported affinity profiling experiments use four concentrations of the photocatalyst conjugate and 12 concentrations of competitive non-photocatalytic molecule in duplicate, for a total of 96 samples.

Affinities were measured in Python (3.12.2). The $EC_{50}$ of competitive binding of a ligand molecule to each protein in the proteome at each photocatalytic probe concentration for which < 25% of values are imputed is modeled using Equation 1, assuming a Hill coefficient of 1:

$$y = \frac{a + c}{1 + 10^{([L] - log_{10}(EC_{50}))}} + c \tag{1}$$

where [L] = competitive ligand molecule concentration, y = integrated MS intensity for a protein, and *a* (maximum intensity minus minimum intensity), *c* (minimum intensity), and the $EC_{50}$ (concentration of ligand molecule displacing 50% of the quantity of probe molecule present in the absence of competition) are all freely optimized parameters. Nonlinear fitting for Equation 1 was performed using lmfit [24].

During optimization, the initial value for *c* is set to the global minimum intensity detected from all proteins and samples and the initial *a* value is set to the maximum of the signal intensity value in samples used for curve fitting. The significance (p-value) of the fit is determined by performing a one-way F-test with the SciPy 'stats' module, comparing Equation 1 to the reduced model y = a, representing a null hypothesis in which there is no relationship between ligand molecule concentration and signal intensity. Each competitive model is fit 30 times with different initial values set for the $EC_{50}$ parameter drawn from a random uniform distribution between -5 and 3 (two order of magnitude beyond the range of concentrations used), retaining the result with the most significant fit. We then test the impact of removing the data point corresponding to treatment with vehicle in place of the unmodified ligand by repeating the nonlinear fitting process and evaluating whether inclusion of vehicle is required for the fit to remain significant. Fits whose significance depends on the inclusion of the vehicle are removed.

The relationship between $K_d$ and $EC_{50}$ for pseudo-first order binding kinetics, i.e. at low concentrations of protein ([Protein] << [Ligand]), is described by the Cheng-Prusoff equation [25] according to Equation 2:

$$K_{\mathrm{d}}{}^{\mathrm{ligand}} = \frac{EC_{50}}{\left(1 + \frac{[\mathrm{probe}]}{K_{\mathrm{d}}{}^{\mathrm{probe}}}\right)} \tag{2}$$

where [probe] = probe concentration, $K_d^{\mathrm{ligand}}$ = dissociation constant of ligand molecule, $K_d^{\mathrm{probe}}$ = dissociation constant of photocatalytic probe, and $EC_{50}$ = $EC_{50}$ of competitive binding by the ligand molecule at each probe molecule concentration. Equation 2 can be expressed as Equation 3, a Schild plot:

$$EC_{50} = K_{\mathrm{d}}{}^{\mathrm{ligand}} + \frac{K_{\mathrm{d}}{}^{\mathrm{ligand}}}{K_{\mathrm{d}}{}^{\mathrm{probe}}} \cdot [\mathrm{probe}] \tag{3}$$

As $EC_{50}$ errors are measured in logarithmic space, the Schild plot is modeled in logarithmic space as well (Equation 4) using curve_fit in the scipy.optimize package [26]:

$$log_{10}(EC_{50}) = log_{10}\left( K_{\mathrm{d}}{}^{\mathrm{ligand}} + \frac{K_{\mathrm{d}}{}^{\mathrm{ligand}}}{K_{\mathrm{d}}{}^{\mathrm{probe}}} \cdot [\mathrm{probe}]\right) \tag{4}$$

The significance of the modeled $K_d^{\mathrm{ligand}} / K_d^{\mathrm{probe}}$ value is measured with a one-way F-test using the SciPy 'stats' module, comparing Equation 4 to a reduced model $\log_{10}(EC_{50}) = a$, representing the null hypothesis that there is no relationship between the value of $EC_{50}$ and the concentration of the probe photocatalyst conjugate. The $K_d^{\mathrm{app}}$ of the free molecule (often a drug, metabolite, peptide, or protein of interest) will be different from an observed $EC_{50}$ by an amount proportional to the ratio of the probe and ligand affinities multiplied by the concentration of its photocatalyst conjugate. As the affinity or concentration of a probe photocatalyst conjugate becomes weaker, the $K_d^{\mathrm{ligand}}$ value of the unconjugated molecule approaches the measured $EC_{50}$ value.

For each protein observed, we perform $EC_{50}$ determination at each photocatalytic probe concentration and use significant ($p < 0.001$) values of the $EC_{50}$ to measure $K_d^{\mathrm{probe}}$ and $K_d^{\mathrm{ligand}}$ via the Cheng-Prusoff equation. Individual proteins have widely varying affinity for the photocatalyst conjugated probe and unmodified ligand molecule; the vast majority do not display any significant $EC_{50}$ values at any value of [probe], reflecting that they do not bind either the probe or ligand molecules. Other proteins which do bind both the probe and ligand molecules within the range of concentrations used show significant $EC_{50}$ values. These are then subjected to Cheng-Prusoff analysis to calculate $K_d^{\mathrm{probe}}$ and $K_d^{\mathrm{ligand}}$. A tiered approach is used to estimate binding affinities for proteins based on the number of significant $EC_{50}$ values observed for each protein:

1. Three or more significant $EC_{50}$ values at different probe concentrations, measurable $K_d^{\mathrm{ligand}} / K_d^{\mathrm{probe}}$: $K_d^{\mathrm{probe}}$ and $K_d^{\mathrm{ligand}}$ are directly measured by fitting the data to Equation 4. If the significance of the modeled slope ($K_d^{\mathrm{ligand}} / K_d^{\mathrm{probe}}$) is significant ($p < 0.1$), both values are reported along with the error from fitting. If the slope is not significant ($p > 0.1$) and data is tightly distributed (confidence interval < $\pm 0.25$), the relative affinity for the probe ligand vs. the ligand molecule is too low to accurately measure. At this limit ($K_d^{\mathrm{ligand}} / K_d^{\mathrm{probe}} \approx 0$), each $EC_{50} \approx K_d^{\mathrm{ligand}}$ ; $K_d^{\mathrm{ligand}}$ is therefore reported as the weighted average of the $EC_{50}$ values in these cases. The weighted average of $EC_{50}$ values is determined by Equation 5, where $\widehat{\mu}$ is the weighted average, N is the total number of $EC_{50}$ used for fitting, $i$ indicates the $EC_{50}$ values and $\sigma$ (standard error) of $EC_{50}$ from fitting in Equation 1 at each value of [probe]. All such proteins are flagged as having modeled $K_d^{\mathrm{ligand}} / K_d^{\mathrm{probe}}$ values.

$$\widehat{\mu} = \frac{\sum_i^N (EC_{50})_i/(\sigma_i)^2}{\sum_i^N 1/(\sigma_i)^2} \tag{5}$$

2. Three or more significant $EC_{50}$ values at different probe concentrations, insufficient data to measure $K_d^{ligand}$ / $K_d^{probe}$: Where the slope is not significant ($p > 0.1$) and data is not tightly distributed (confidence interval > ±0.25), $K_d^{ligand}$ / $K_d^{probe}$ cannot be empirically determined. The weighted average $EC_{50}$ value (calculated using Equation 5) is therefore reported, reflecting the null hypothesis $K_d^{ligand} / K_d^{probe} \approx 0$).

3. Two significant $EC_{50}$ values at different probe concentrations: Curve fitting with two points cannot afford an accurate estimation of error in the resulting fit. The value measured at the lowest photocatalytic probe concentration is therefore reported as a best estimate of $K_d^{ligand}$ as it is the closest empirical value to the y-intercept in the Schild plot.

4. One significant $EC_{50}$ value at one probe concentration: $K_d^{ligand}$ is reported as the $EC_{50}$.

5. Zero significant $EC_{50}$ values: The most significant $EC_{50}$ value is reported as $K_d^{ligand}$.

For each protein, the total significance of the data used for $K_d^{ligand}$ estimation is reported as the Fisher combined p-values of all fitted $EC_{50}$ values observed, and this aggregated p-value is then subjected to Benmamini-Hochburg FDR estimation to account for multiple hypothesis testing and represents the significance of all evidence used to support a reported $K_d^{ligand}$ value. For most proteins, $K_d^{ligand}$ / $K_d^{probe}$ is small because attaching the photocatalyst to a ligand significantly reduces its affinity ($K_d^{ligand} << K_d^{probe}$), consistent with the impact of attaching other types of photoaffinity labels; in such cases, weighted average $EC_{50}$ values give very good estimates of $K_d^{app}$ because the influence of photocatalytic probe concentration on any observed $EC_{50}$ value is small. Since proteins that have measurable $EC_{50}$ values only at high probe concentration (scenarios 2, 3, and 4 above) likely have small $K_d^{ligand}$ / $K_d^{probe}$ ratios, and most protein-ligand systems have small $K_d^{ligand}$ / $K_d^{probe}$ ratios (see supplementary dataframe reporting all fitted parameters, Supplementary_Data_1.xlsx), single $EC_{50}$ values (scenario 4) can be used as acceptable best estimates of $K_d^{ligand}$. In scenarios with exactly two significant $EC_{50}$ values, using the $EC_{50}$ value obtained at the lower value of photocatalyst concentration provides an improved estimate as it is measured closer to the value of $K_d^{ligand}$ in the Schild plot.

We find that the assumption of small values for $K_d^{ligand}$ / $K_d^{probe}$ is valid in data generated using photocatalytic affinity profiling for dasatinib, which shows good correlation with reported affinities for kinases with $K_d^{ligand}$ values measured using three or more, two, or one significant $EC_{50}$ value (**Fig. S14, Figure 2G**). Indeed, correlation between $K_d^{ligand}$ values measured by photocatalytic affinity profiling and literature values is good and has a slope near one, reflecting the validity of both the broader assumptions made to constrain the kinetic model used for analysis and our tiered data processing pipeline approach.

**Volcano plot construction: µMap target-ID processing for two-sample group comparison**

Processing of DIANN output files (report.pg_matrix.tsv, report.pr_matrix.tsv) was done in R (4.4.1) using the MSnbase package separately for each set of samples treated with given concentration of photocatalyst conjugate and off-compete molecule or vehicle. Proteomics data was subjected to variance stabilization normalization ('vsn') in independent experimental groups [21,22], imputation using MinProb in MSnbase, and the number of peptides used for each protein inference determined and added as metadata. Once complete, individually processed MSnSet datasets were combined into a single table. Differential expression analysis was performed using linear models ('limma') [27] with Benmamini-Hochburg FDR adjustment to identify significantly enriched proteins. A volcano plot comparing the $\log_2$ fold change and $-\log_{10}$(p-value) was generated, with significantly enriched proteins colored by affinity measured using Affinity Map.

#### Synthesis and Characterization

**General information concerning synthesis and characterization**.

Reaction temperatures refer to heating/cooling media (heating block, oil bath, or cryogenic bath). After preparative HPLC purification, fractions containing the product were evaporated using a SPGenevac EZ-2 4.0. Final compounds were characterized by $^{1}$H NMR, $^{13}$C NMR, $^{19}$F NMR (when applicable), and ESI-MS. Retention times refer to analytical LC-MS chromatography, which was performed via HPLC-MS-ELSD using a 5%-95% solvent B gradient over 10 minutes, unless otherwise stated. Isolated molecules are reported with complete analytical characterization when not previously reported in literature. Commercially available reagents were purchased from Ambeed, Oakwood, Tokyo Chemical Industry, Sigma Aldrich, or Thermo Fisher Scientific.

**Solvents:**

Hexanes (VWR, ACS reagent), $CH_2Cl_2$ (VWR, ACS reagent), $CHCl_3$ (Merck, > 98%), ethyl acetate (VWR, ACS reagent), methanol (Sigma-Aldrich, HPLC grade, > 99.8%), DMF (Oakwood, ACS reagent), DMSO (Alfa-Aesar, > 99.8%), MeCN (Sigma-Aldrich, HPLC grade, > 99.9%), acetone (VWR, ACS reagent), ethanol (KOPTEC, USP 200 proof) were used without additional purification. Anhydrous THF was purchased from Sigma-Aldrich (anhydrous, > 99.9%) and distilled over sodium/benzophenone, then stored in a Schlenk tube under argon in the presence of activated 4Å molecular sieves.

**NMR spectroscopy**:

Experiments were performed at the CLC Nuclear Magnetic Resonance Core Facility at Weill Cornell Medicine using a Bruker Avance III HD 500 MHz equipped with a 5 mm CPTCI CryoProbe or an Agilent Varian INOVA 600 MHz with a 5 mm BBO600S3 probe. The following sequences were used: $^{1}$H NMR: zg30; $^{13}$C NMR: zgpg30; $^{19}$F NMR: zg30. The following solvents were used: $CDCl_3$ (Sigma-Aldrich, 99.8% *d*), $H_2O$ (Cambridge Isotope Laboratories, 99.9% *d*), DMSO-$d_6$ (Cambridge Isotope Laboratories, 99.9% *d* + 0.05 *v/v* TMS), $CD_3OD$ (Cambridge Isotope Laboratories, 99.8% *d*). Spectra were analyzed using MestReNova 14.3 by applying standard baseline and phase correction methods. Chemical shifts (δ) for $^{1}$H and $^{13}$C NMR spectra are given in parts per million (ppm) relative to residual solvent peaks. $^{19}$F NMR spectra were calibrated using an absolute referencing system, as suggested by IUPAC. $^{1}$H and $^{13}$C NMR multiplicities that can be analyzed as first order multiplets are reported using the following abbreviations (or combinations thereof): s = singlet, d = doublet, t = triplet, q = quartet, p = quintet, h =sextet; hept = heptet; m = multiplet, br = broad.

**Flash column chromatography**:

Purifications were performed using a Teledyne ISCO CombiFlash RF$^{+}$ Lumen system equipped with reusable cartridges loaded with Cleanert Silica (particle size: 40 – 60 µm, pore size: 60 Å). Column volumes (CV) are noted which scale according to appropriate cartridge size.

**HPLC purification details:**

**General method development:**
For all HPLC analysis and purification a Waters AutoPurification System with 2424-ELS Detector, 2998-Photodiode Array Detector and SQ Detector 2 was used. All gradients begin at 5% solvent B and hold this for 30 seconds during compound loading. The gradient then increases to the specified percentage solvent B over the specified time window. For isocratic methods, the percentage solvent B is reached in 90 seconds using a linearly increasing gradient and held for the specified duration of the run. Sample sandwiching refers to the use of a specified volume of a solvent before and after sample uptake by the syringe directly prior to injection in order to prevent hydrophobic compound precipitation during column loading.

Solvent A: $H_2O$:Formic acid (999:1)
Solvent B: MeCN:Formic acid (999:1)

Preparative column: Xbridge BEH C18 OBD Prep Column, 130Å, 5 µm, 19 mm x 150 mm, 186002979
Preparative flow rate: 20 mL/min

Semi-preparative column: Xbridge BEH C18 OBD Prep Column, 130Å, 5 µm, 10 mm x 150 mm, 186008165
Semi-preparative flow rate: 10 mL/min

Analytical column: Xbridge C18 column, 130Å, 5 µm, 4.6 x 159 mm, 186003116
Analytical flow rate: 1.4 mL/min

**Common reagents and starting materials**

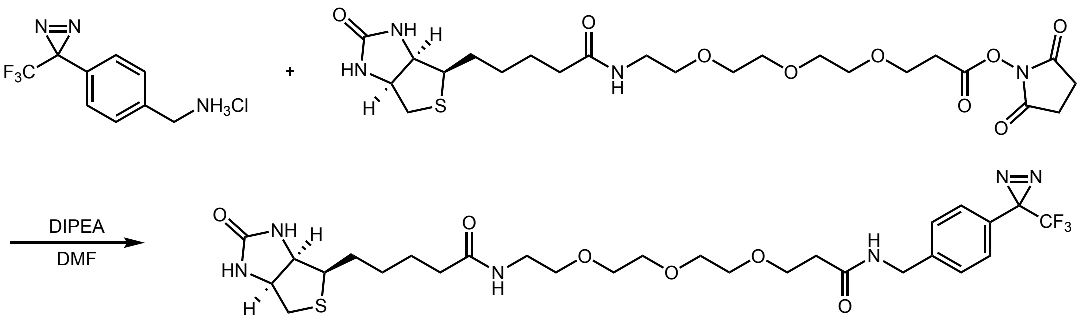

**Diazirine-biotin (4)**

The reaction was performed according to a modified literature procedure [28]. A scintillation vial containing a PTFE-coated stir bar was charged with diazirine amine (139 mg, 0.551 mmol, 1.0 equiv.) and biotin-PEG$_3$-NHS ester (300 mg, 0.551 mmol, 1.0 equiv.). The solids were dissolved in DMF (2.2 mL) and diisopropylethylamine (DIPEA, 192 µL, 1.102 mmol, 2.0 equiv.) was added. The reaction was stirred at room temperature for 16 hours. Solvent was removed and the residue was purified by flash column chromatography on silica (DCM:MeOH = 0% to 80% over 12 CVs) to afford the title compound as an opaque oil (0.330 g, 93%). Spectral data are consistent with existing literature [28].

Note: Trifluoromethyl diazirines can easily isomerize to diazo compounds under typical laboratory conditions (ambient lighting, room temperature, exposure to Lewis acids and transition metals). The diazo isomers are electrophilic, have shifted absorption spectra, and can cause non-specific background protein biotinylation in the presence or absence of catalyst. This, in turn, decreases observed fold change and can obfuscate results. Care should be taken to handle this compound in the dark and avoid extended storage at room temperature, as even trace quantities of diazo isomer can cause irreproducible results. The presence of diazo isomers in stock solutions must be routinely assayed via $^{19}$F NMR (-CF$_3$: -65.5 ppm for diazirine, -57.6 ppm for diazo).

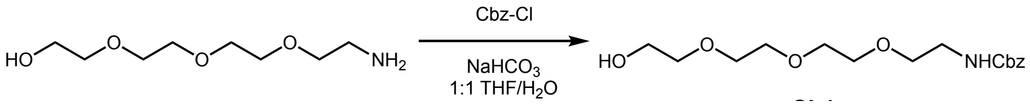

**Benzyl (2-(2-(2-(2-hydroxyethoxy)ethoxy)ethoxy)ethyl)carbamate (6)**. The reaction was performed according to a modified literature procedure [29]. A round-bottom flask equipped with a PTFE-coated stir bar was charged with amino-$PEG_4$-alcohol (2.8 mL, 15.49 mmol, 1.0 equiv.), sodium bicarbonate (1.952 g, 23.24 mmol, 1.5 equiv.), and $H_2O$ (31 mL). The mixture was stirred at room temperature until homogeneous, then cooled to 0 °C for 10 minutes. Separately, benzyl chloroformate (2.65 mL, 18.59 mmol, 1.5 equiv.) was dissolved in THF (31 mL) and this solution was added dropwise to the amine solution with vigorous stirring at 0 °C. The reaction was stirred at 0 °C for 1 hour, then allowed to warm to room temperature and stirred overnight. Solvent was removed and the crude mixture was redissolved in EtOAc:$H_2O$ (1:1, 150 mL). The aqueous layer was extracted 3x with EtOAc (75 mL), and the EtOAc extracts washed with brine, dried with $Na_2SO_4$, and concentrated. The oil was diluted in DCM and purified by flash column chromatography on silica (hexanes:EtOAc = 0% to 100% over 10 CVs) to afford the title compound as a clear oil (2.709 g, 53%). Spectral data are consistent with existing literature [29].

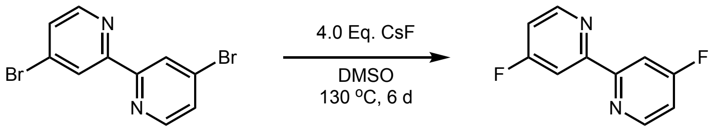

**4,4'-Difluoro-2,2'-bipyridine (7).**
A scintillation vial equipped with a PTFE-coated stir bar was charged with 4,4'-dibromo-2,2'-bipyridine (10 g, 32 mmol, 1.0 equiv.), cesium fluoride (19.44 g, 128 mmol, 4 equiv.), and DMSO (25.6 mL). The mixture was heated to 150 °C and stirred vigorously for 6 days. The reaction mixture was cooled to room temperature and partitioned between $H_2O$ (200 mL) and DCM (200 mL). The aqueous layer was extracted 3x with DCM (100 mL) and the DCM extracts were washed 5x $H_2O$ (75 mL) to remove all DMSO. The DCM extract was washed with brine, dried with $Na_2SO_4$, and concentrated. The residue was purified by flash column chromatography on silica (hexanes:EtOAc = 0% to 10% over 7 CVs) to afford the title compound as a white solid (2.75 g, 45%). The spectral data were consistent with existing literature [30].

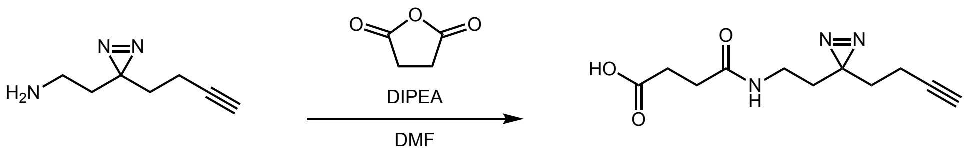

**4-((2-(3-(But-3-yn-1-yl)-3H-diazirin-3-yl)ethyl)amino)-4-oxobutanoic acid (Carboxyl-Dz-Alkyne)**
A scintillation vial equipped with a PTFE-coated stir bar was charged with amino diazirine alkyne (0.021 g, 0.15 mmol, 1.0 equiv.) and succinic anhydride (0.017 g, 0.165 mmol, 1.1 equiv.). The mixture was dissolved in DMF (1 mL). After stirring for 5 minutes DIPEA (0.130 mL, 0.75 mmol, 5 equiv.) was added and the reaction was stirred overnight at room temperature in the dark. The reaction was purified by semipreparative HPLC using a 20%-60% solvent B over 10-minute method. The product eluted at 3.6 minutes and solvent was removed to afford the title compound. (12.2 mg, 34%).

**$^{1}$H NMR** (500 MHz, $CDCl_3$) δ 6.23 (s, 1H), 3.09 (q, $J$ = 6.6 Hz, 2H), 2.58 (dt, $J$ = 89.5, 6.6 Hz, 4H), 2.05 – 1.97 (m, 3H), 1.72 – 1.61 (m, 4H).
**$^{13}$C NMR** (126 MHz, $CDCl_3$) δ 172.54, 82.89, 77.41, 77.16, 76.91, 69.59, 34.62, 32.52, 32.19, 31.03, 26.93, 13.33.
**LC-MS** $R_t$ = 4.68 min; m/z calcd. For $C_{11}H_{15}N_3O_3$ ($[M+H]^+$) = 238.1, found 238.1

Carboxyl-Dz-alkyne $^{1}$H-NMR

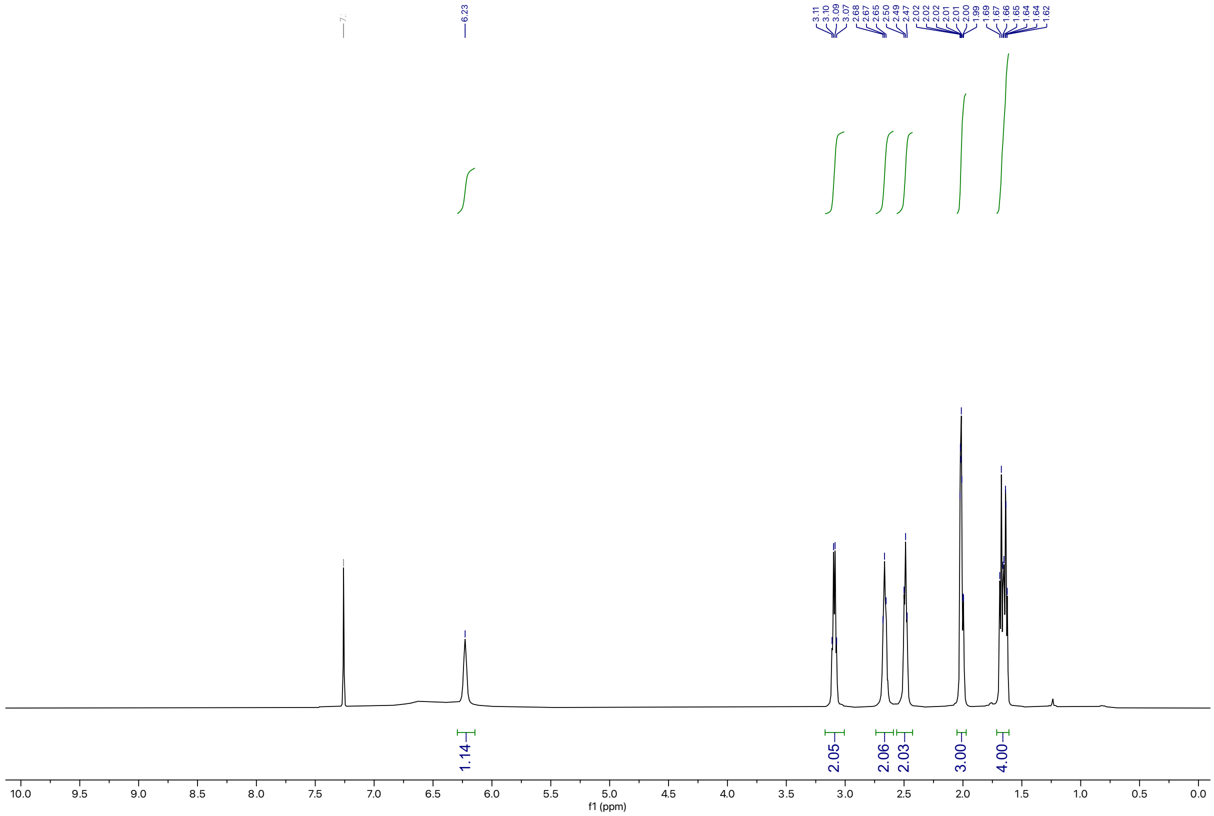

Carboxyl-Dz-alkyne $^{13}$C-NMR

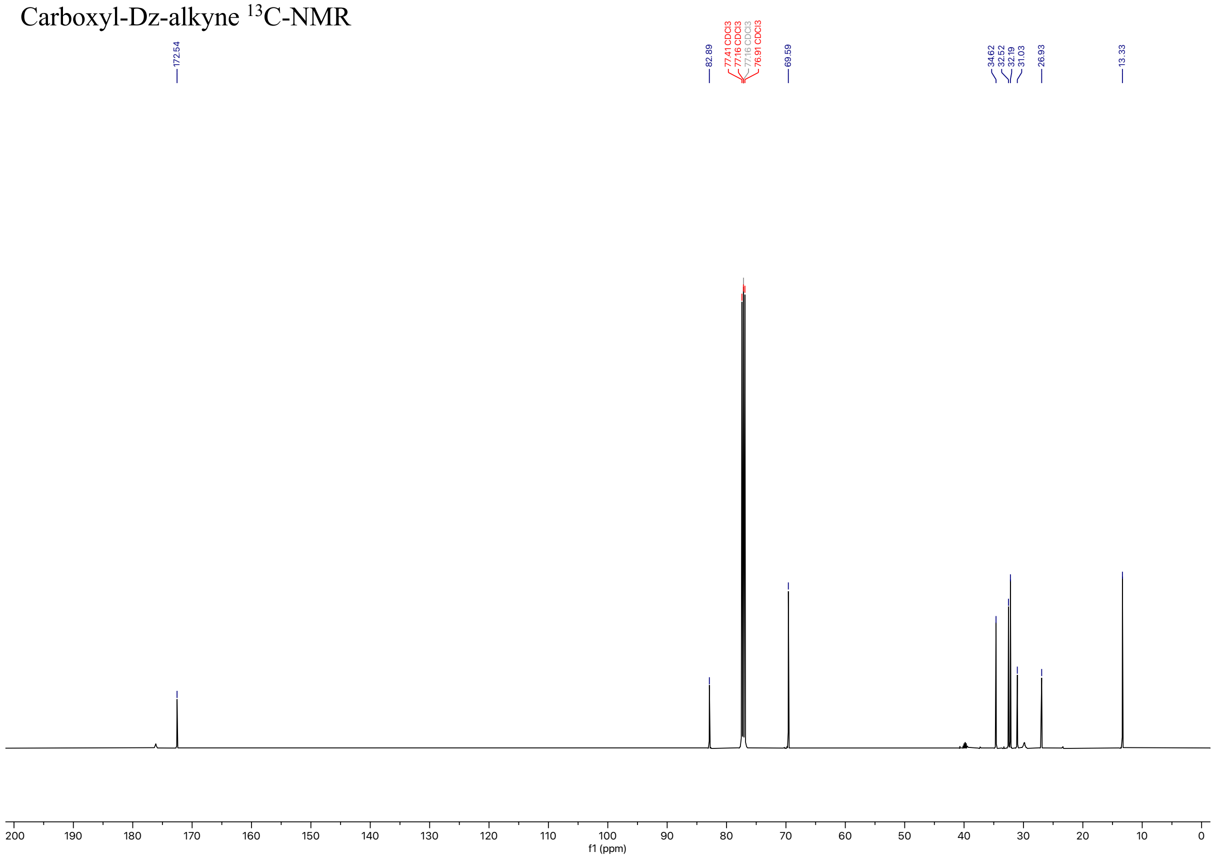

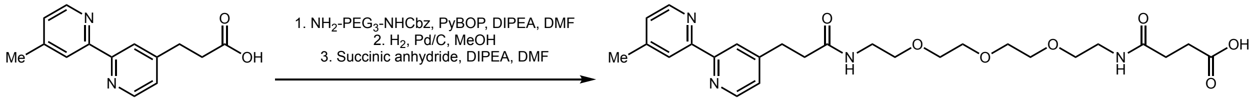

**1-(4’-Methyl-[2,2’-bipyridin]-4-yl)-3,17-dioxo-7,10,13-trioxa-4,16-diazaicosan-20-oic acid (BiPy-Me-PEG3-$CO_2$H)**

Amide coupling: A scintillation vial equipped with a PTFE-coated stir bar was charged with 3-(4’-methyl-[2,2’-bipyridin]-4-yl)propanoic acid (0.280 g, 1.157 mmol, 1.0 equiv.), 3-(2-(2-(3-benzyloxycarbonylaminopropoxy)ethoxy)ethoxy)propylamine (0.415 g, 1.378 mmol, 1.1 equiv.), 50chlenk50azole-1-yloxytripyrrolidinophosphonium hexafluorophosphate (PyBOP) (1.806 g, 3.471 mmol, 3.0 equiv.), and dissolved in DMF (5.79 mL). The solution was stirred for 5 minutes, then DIPEA was added (0.806 mL, 4.628 mmol, 4.0 equiv.). After 16 hours solvent was removed and the residue partitioned between $H_2O$ (100 mL) and DCM (100 mL). The aqueous layer was extracted 3x with DCM (50 mL) and the DCM extract was washed 5x $H_2O$ (50 mL) to remove all DMF. The DCM extract was washed with brine, dried with $Na_2SO_4$, and concentrated to remove solvent. The residue was purified by flash column chromatography on silica (MeOH:DCM = 5% over 8 CVs). The oil was redissolved in DMF (2.0 mL) and purified by preparative HPLC using a 30%-50% solvent B over 10 minutes. The product eluted at 3.5 minutes and solvent was removed to afford the protected intermediate (276 mg, 43%).

**$^1$H NMR** (598 MHz, $CD_3OD$_SPE) δ 8.53 (dd, $J$ = 16.9, 5.1 Hz, 3H), 8.17 (dd, $J$ = 29.2, 1.8 Hz, 3H), 7.38 – 7.33 (m, 8H), 7.32 (dd, $J$ = 5.2, 1.5 Hz, 2H), 7.29 (d, $J$ = 5.1 Hz, 2H), 5.09 (d, $J$ = 14.1 Hz, 4H), 3.60 (d, $J$ = 5.6 Hz, 0H), 3.58 (s, 6H), 3.56 (dd, $J$ = 5.9, 3.1 Hz, 3H), 3.53 – 3.49 (m, 6H), 3.45 (t, $J$ = 5.4 Hz, 3H), 3.36 – 3.33 (m, 4H), 3.05 (t, $J$ = 7.5 Hz, 3H), 2.62 (t, $J$ = 7.5 Hz, 3H), 2.46 (s, 5H), 1.96 (s, 1H).
**$^{13}$C NMR** (150 MHz, $CD_3OD$_SPE) δ 174.18, 173.15, 158.64, 156.73, 156.48, 153.26, 150.57, 150.03, 149.76, 138.27, 129.40, 128.91, 128.75, 126.17, 125.44, 123.54, 122.78, 71.46, 71.41, 71.11, 71.09, 71.06, 70.83, 70.45, 70.41, 67.31, 67.29, 49.43, 49.28, 49.14, 49.00, 48.86, 48.71, 48.57, 41.68, 40.37, 40.31, 37.02, 32.08, 22.54, 21.25.
**LC-MS** $R_t$ = 4.95 min; m/z calcd. For $C_{30}H_{38}N_4O_6$ ($[M+H]^+$) = 551.3, found 551.5

Cbz deprotection: A scintillation vial equipped with a PTFE-coated stir bar was charged with protected intermediate (0.073 g, 0.132 mmol, 1.0 equiv.) and 10% Pd/C, type 487 (1.4 mg) and suspended in MeOH (1.32 mL). The solution was placed under $H_2$ using a balloon and stirred at room temperature for 16 hours. The mixture was filtered over celite and the filter cake was washed with MeOH (80 mL). Solvent was removed to afford the deprotected intermediate as a clear oil which was carried forward without additional purification (52.8 mg, 96%)

Succinylation: A scintillation vial equipped with a PTFE-coated stir bar was charged with the deprotected intermediate (0.052 g, 0.125 mmol, 1.0 equiv.) and succinic anhydride (0.015 g, 0.15 mmol, 1.2 equiv.), and dissolved in DMF (0.83 mL). The reaction was stirred at room temperature for 5 minutes, then DIPEA (0.109 mL, 0.625 mmol, 5.0 equiv.) was added and the reaction was stirred for 16 hours. The product was purified by preparative HPLC using a 5%-55% solvent B gradient over 10 minutes. The product eluted at 4.1 minutes and solvent was removed to afford the title compound as a clear oil (36.2 mg, 56%).

**$^1$H NMR** (598 MHz, $CDCl_3$) δ 8.51 (dd, $J$ = 21.6, 5.0 Hz, 2H), 8.16 (dd, $J$ = 4.4, 1.7 Hz, 2H), 7.17 (ddd, $J$ = 9.2, 5.1, 1.7 Hz, 2H), 6.93 (t, $J$ = 5.7 Hz, 1H), 6.87 (t, $J$ = 5.7 Hz, 1H), 3.61 – 3.50 (m, 10H), 3.47 (t, $J$ = 5.1 Hz, 2H), 3.38 (dq, $J$ = 7.5, 5.3 Hz, 4H), 3.01 (t, $J$ = 7.6 Hz, 2H), 2.66 (t, $J$ = 6.7 Hz, 2H), 2.58 (t, $J$ = 7.7 Hz, 2H), 2.48 (t, $J$ = 6.8 Hz, 2H), 2.43 (s, 3H).
**$^{13}$C NMR** (150 MHz, $CDCl_3$) δ 175.29, 172.39, 172.15, 155.62, 155.44, 151.49, 149.43, 149.27, 148.32, 125.08, 124.61, 122.97, 121.64, 77.37, 77.16, 76.95, 70.57, 70.32, 70.14, 70.10, 69.99, 69.79, 39.49, 39.35, 39.28, 36.60, 31.30, 31.07, 29.99, 21.36.
**LC-MS** $R_t$ = 3.99 min; m/z calcd. For $C_{26}H_{36}N_4O_7$ ($[M+H]^+$) = 517.3, found 517.5

Amide Coupling Product $^{1}$H-NMR

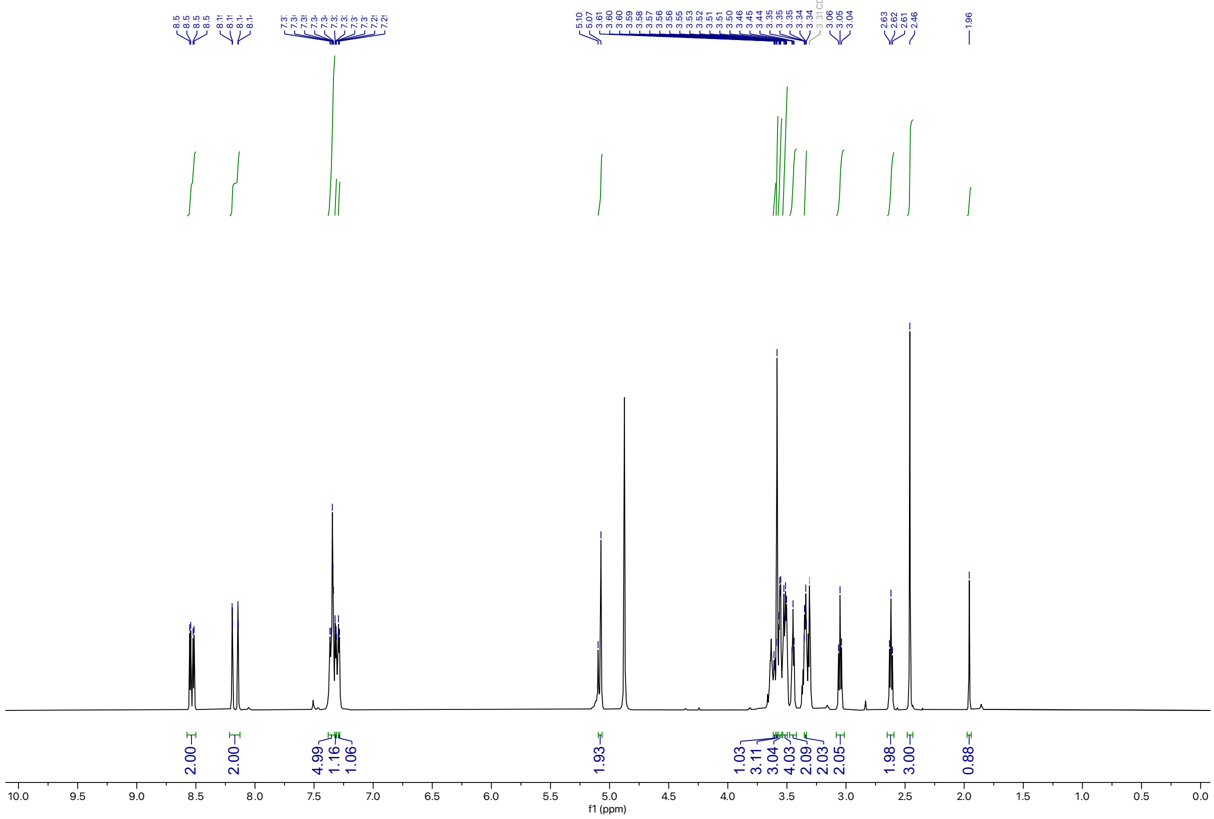

Amide Coupling Product $^{13}$C-NMR

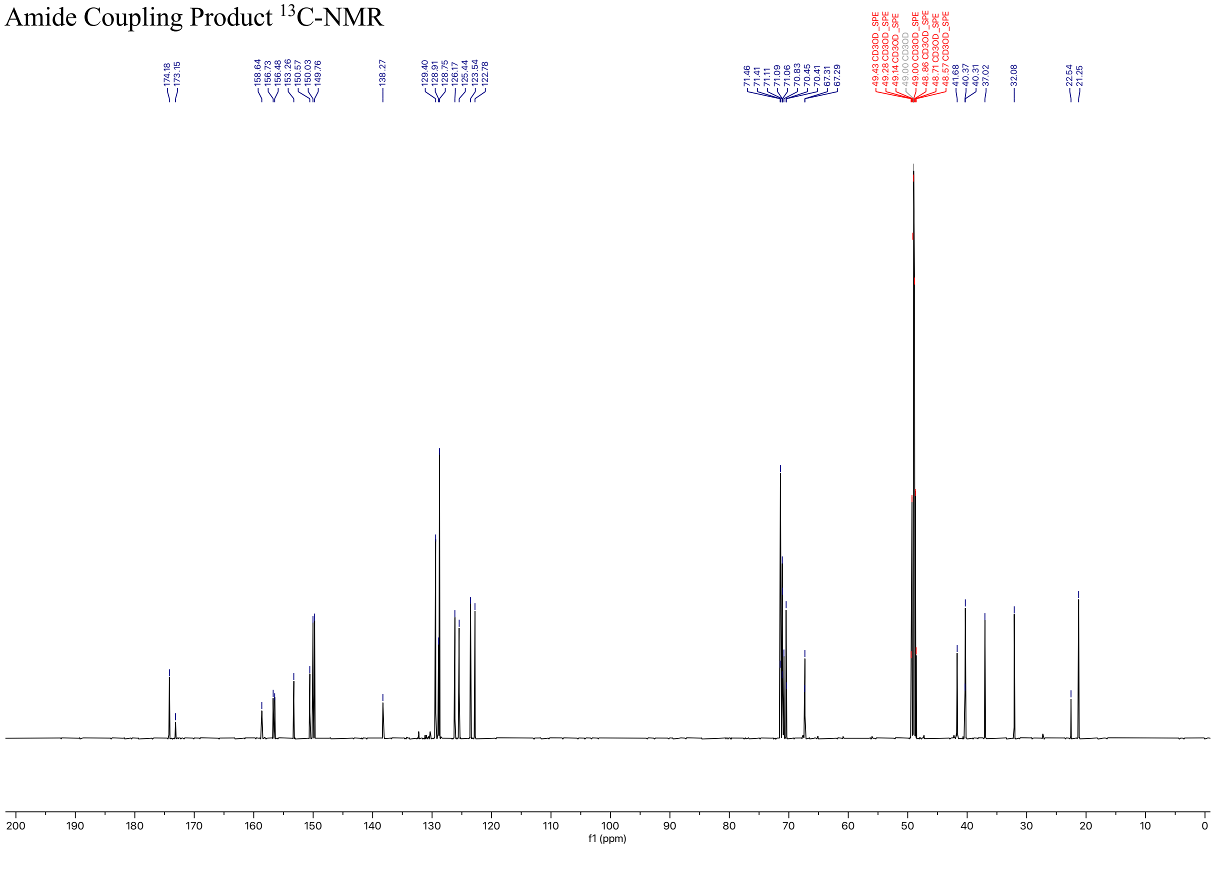

Succinylated Product $^{1}$H-NMR

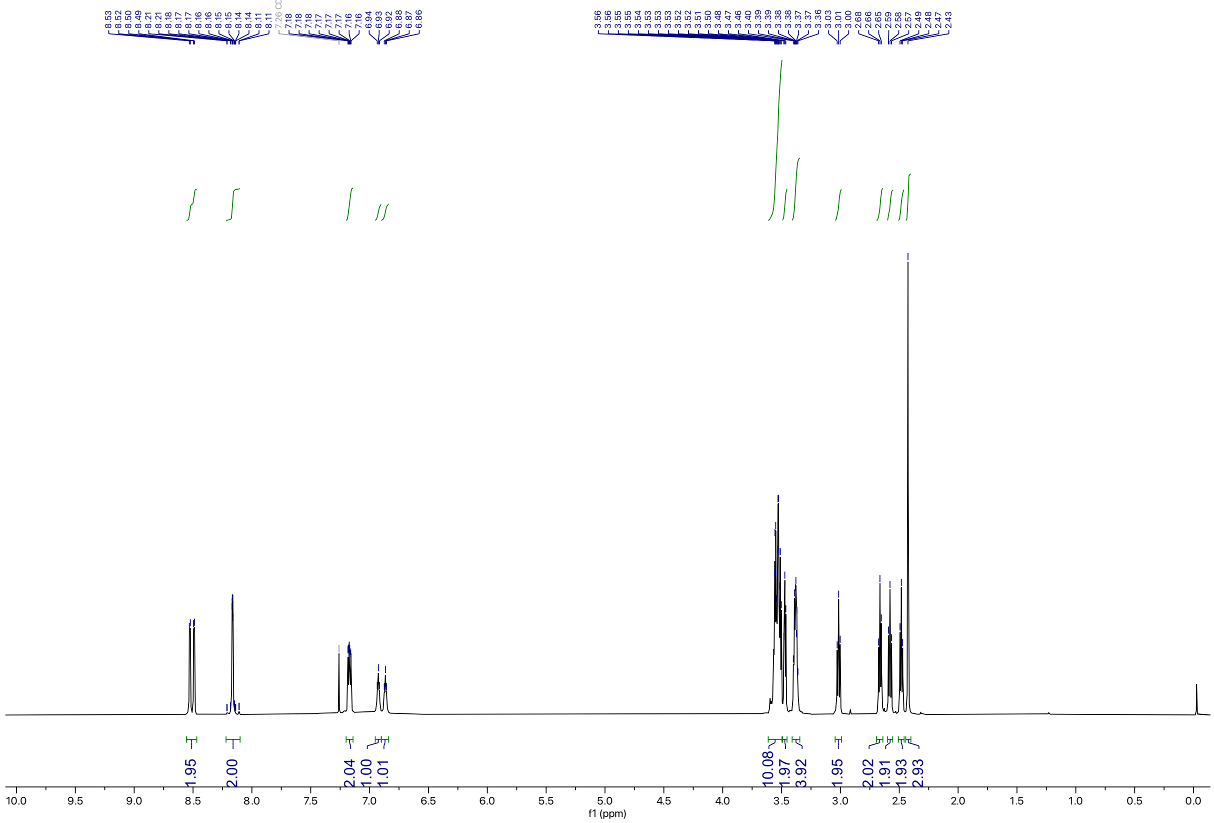

Succinylated Product $^{13}$C-NMR

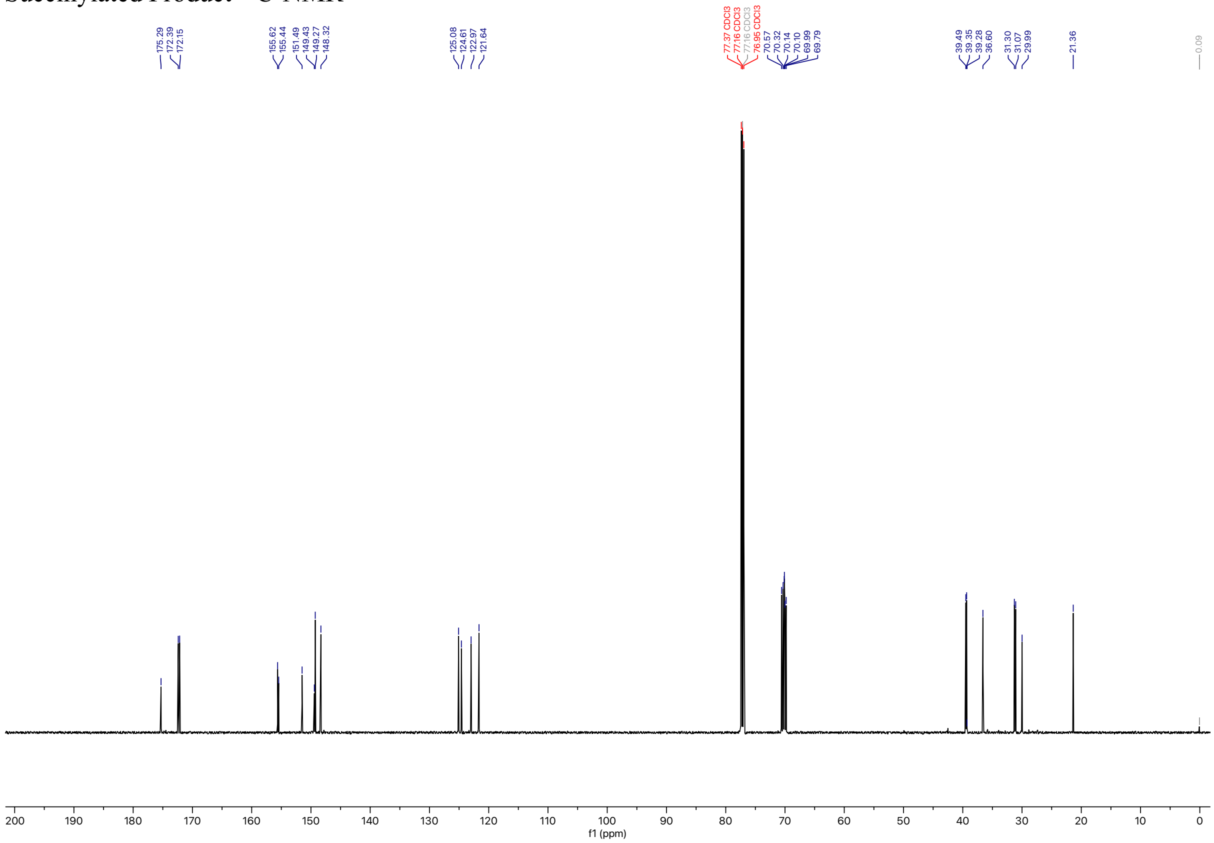

#### Synthesis of 1: Overview

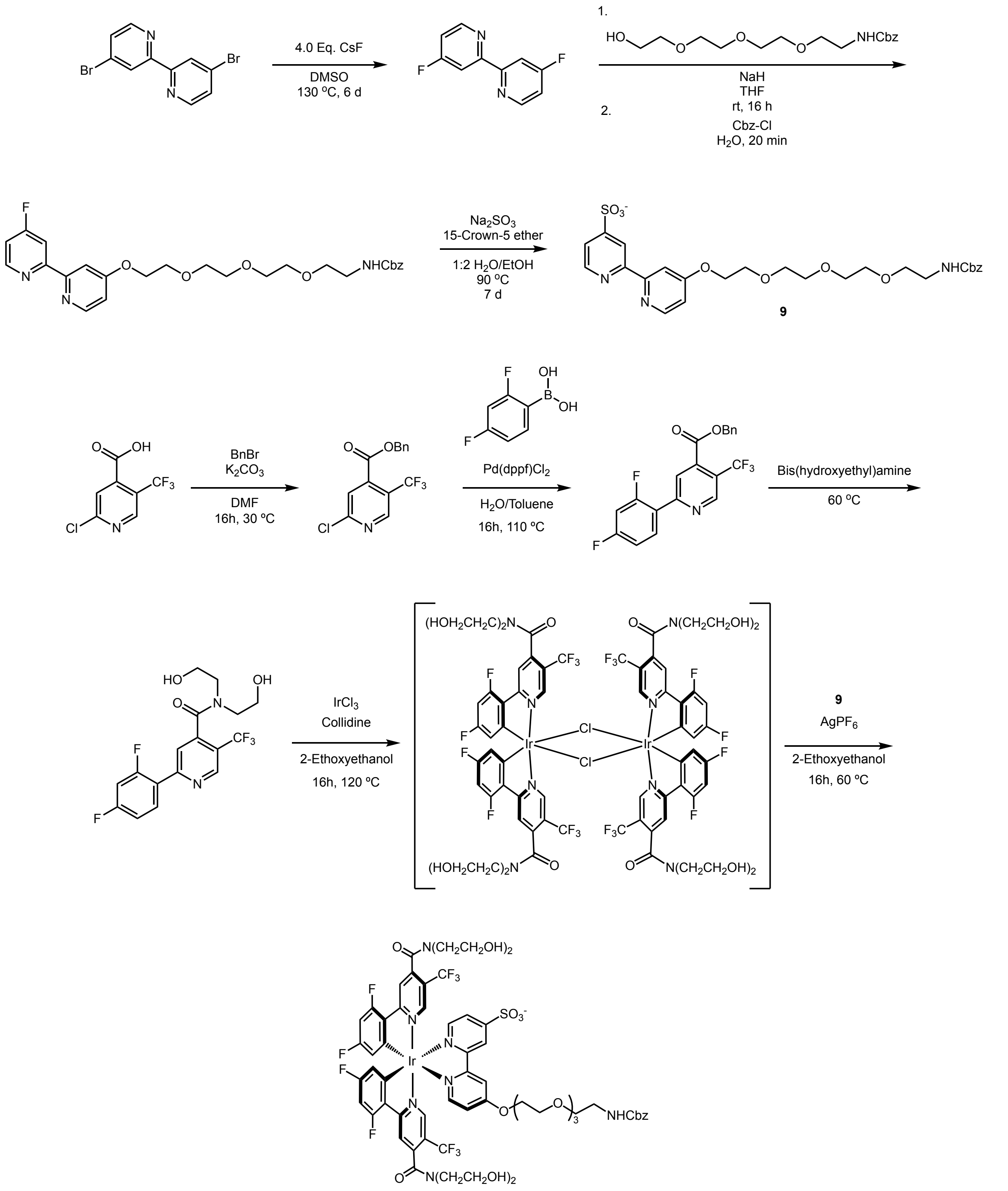

**1**

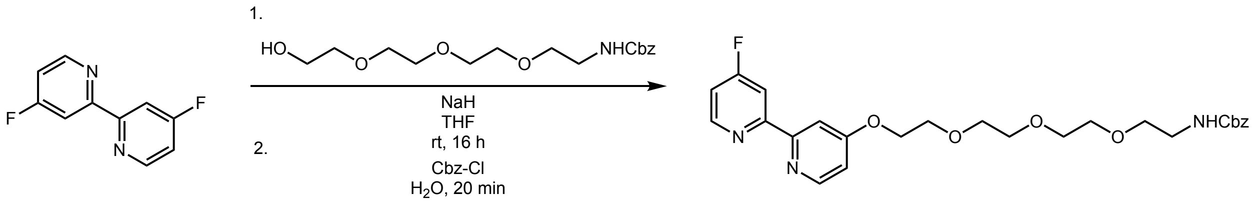

**Benzyl (2-(2-(2-(2-((4’-fluoro-[2,2’-bipyridin]-4-yl)oxy)ethoxy)ethoxy)ethoxy)ethyl)carbamate (8).**
A scintillation vial equipped with a PTFE-coated stir bar was charged with **6** (1.98 g, 6.0 mmol, 1.0 equiv.), **7** (2.93 g, 15.2 mmol, 2.5 equiv.), sodium hydride (60% w/w in mineral oil, 1.16 g, 30.4 mmol, 5 equiv.) and cooled to 0 °C. The vial was sparged with argon. Anhydrous THF (12 mL) was added and the mixture stirred at 0 °C for 15 minutes. The mixture was warmed to room temperature and stirred for 16 hours. The reaction was then cooled to 0 °C and quenched with water (5 mL). Benzyl chloroformate (0.507 mL, 3.58 mmol, 0.59 equiv.) was added dropwise, then the reaction was stirred at room temperature for 30 minutes and solvent was removed. The crude mixture was then partitioned between $H_2O$ (100 mL) and DCM (100 mL). The aqueous layer was extracted 3x with DCM (100 mL) and the DCM extracts were washed with brine, dried with $Na_2SO_4$, and concentrated to remove solvent. The resulting residue was purified by flash column chromatography on silica (hexanes:EtOAc = 0% to 100% over 7 CVs) to afford the title compound as a clear oil (1.55 g, 51%).

**$^1$H NMR** (598 MHz, $CDCl_3$) δ 8.65 – 8.53 (m, 1H), 8.45 (d, $J$ = 5.7 Hz, 1H), 8.14 (d, $J$ = 10.4 Hz, 1H), 7.96 (s, 1H), 7.34 – 7.27 (m, 5H), 7.02 (s, 1H), 6.85 (s, 1H), 5.40 (br s, 1H), 5.08 (s, 2H), 4.23 (s, 2H), 3.84 (s, 2H), 3.70 (s, 2H), 3.68 – 3.63 (m, 2H), 3.60 (d, $J$ = 6.9 Hz, 4H), 3.54 (s, 2H), 3.37 (d, $J$ = 5.5 Hz, 2H).
**$^{13}$C NMR** (150 MHz, $CDCl_3$) δ 170.48, 168.74, 165.91, 159.40 (d, $J$ = 7.2 Hz), 156.68 (d, $J$ = 3.8 Hz), 156.48, 151.33 (d, $J$ = 6.8 Hz), 150.36, 136.63, 128.50, 128.10 (d, $J$ = 7.5 Hz), 111.69, 111.59 (d, $J$ = 16.6 Hz), 109.00 (d, $J$ = 18.6 Hz), 106.96, 70.91, 70.60 (d, $J$ = 9.3 Hz), 70.26, 70.01, 69.32, 67.50, 66.63, 40.88.
**$^{19}$F NMR** (563 MHz, $CDCl_3$) δ -102.00 (d, $J$ = 9.1 Hz).
**LC-MS** $R_t$ = 5.16 min; m/z calcd. For $C_{26}H_{30}FN_3O_6$ ($[M+H]^+$) = 500.2, found 500.5

**8** $^{1}$H-NMR

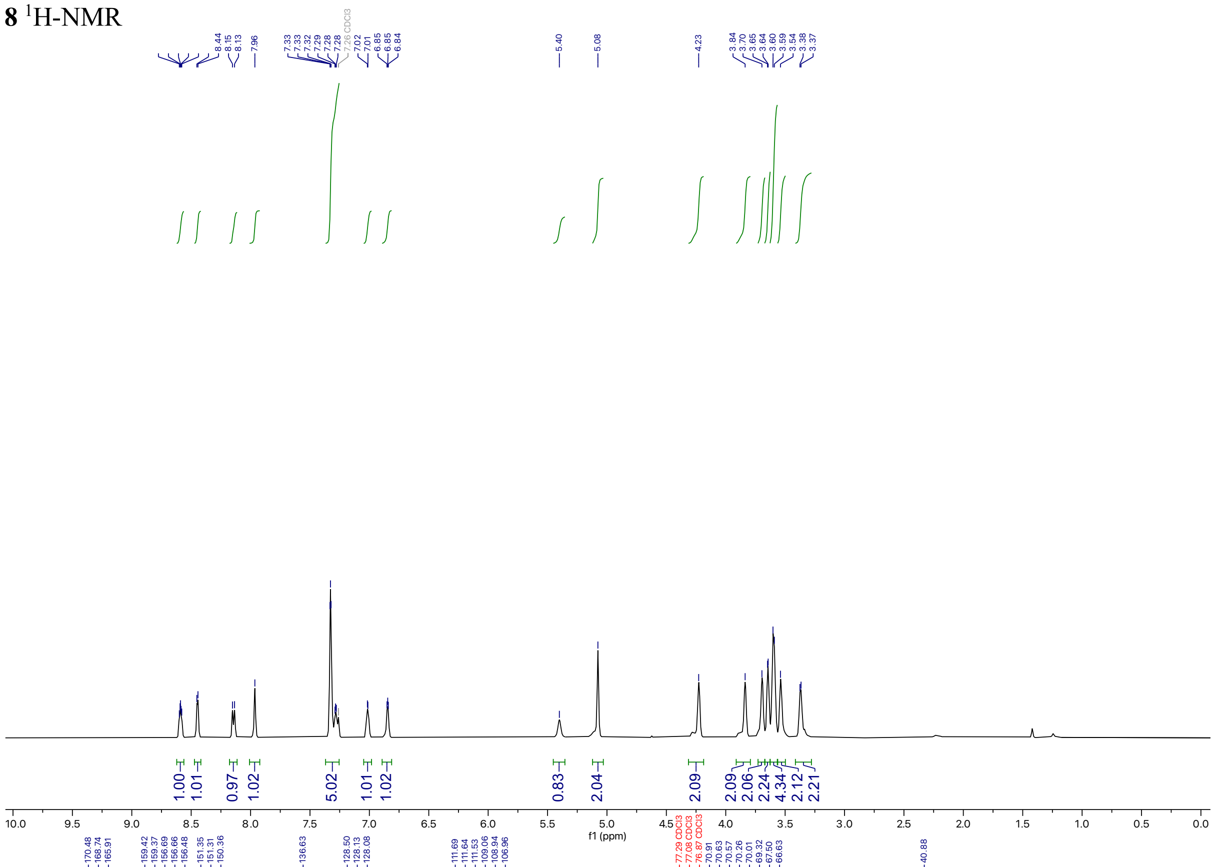

**8** $^{13}$C-NMR

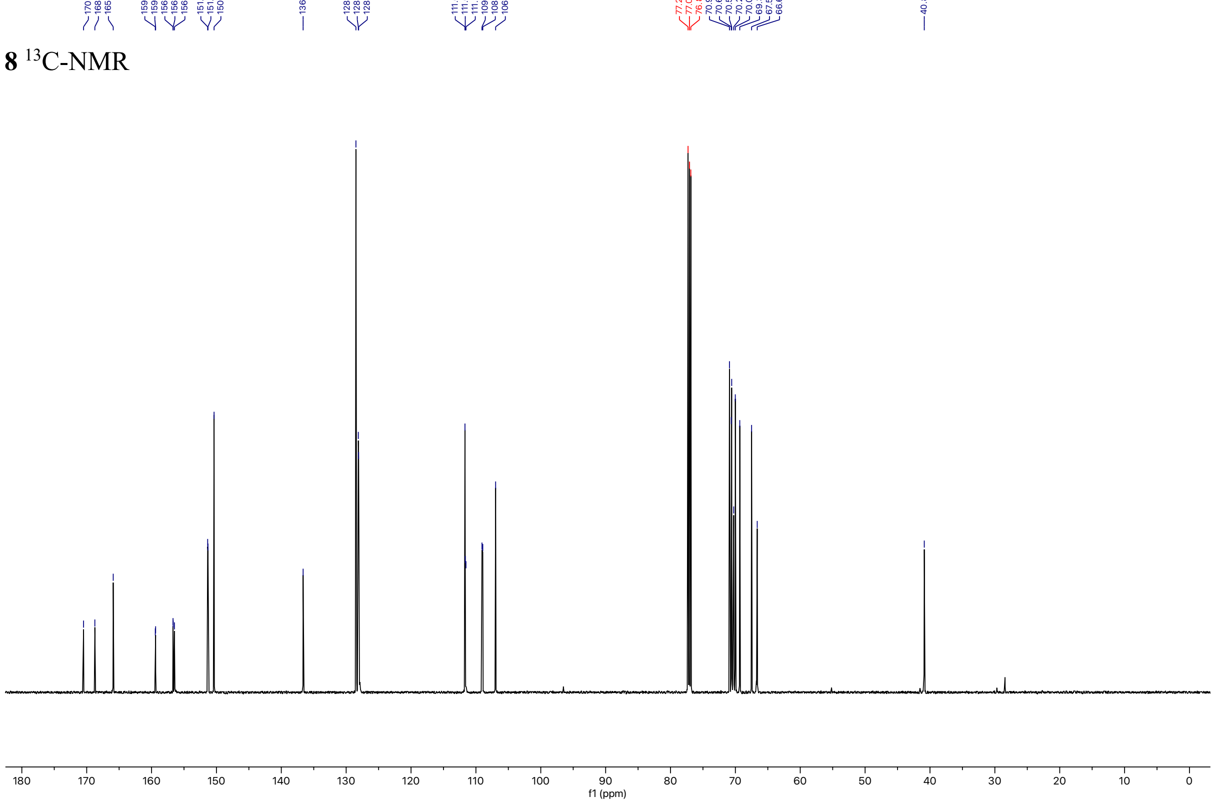

**8** $^{19}$F-NMR

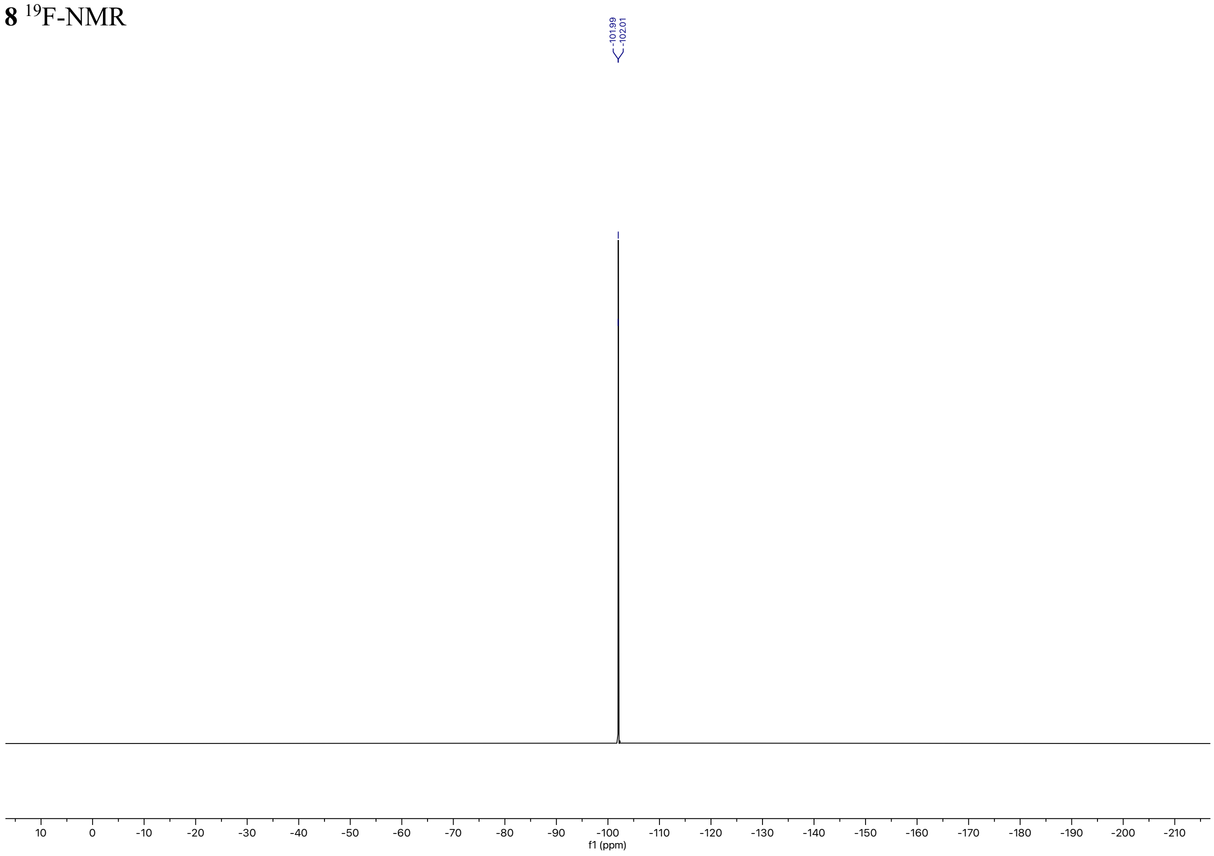

-101.99
-102.01
10
0
-10
-20
-30
-40
-50
-60
-70
-80
-90
-100
-110
-120
-130
-140
-150
-160
-170
-180
-190
-200
-210
f1 (ppm)

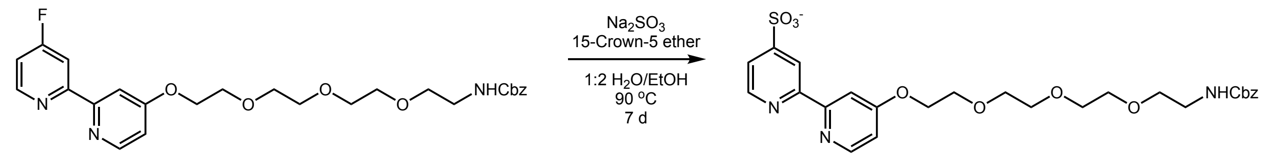

**4’-((3-Oxo-1-phenyl-2,7,10,13-tetraoxa-4-azapentadecan-15-yl)oxy)-[2,2’-bipyridine]-4-sulfonate (9).**
A scintillation vial equipped with a PTFE-coated stir bar was charged with **8** (0.81 g, 1.6 mmol, 1.0 equiv.), sodium sulfite (0.61 g, 4.86 mmol, 3 equiv.), 15-crown-5 (1.98 mL, 9.7 mmol, 6 equiv.) and EtOH:$H_2O$ (2:1, 13 mL). The vial was sparged with argon for 15 min. The reaction was then heated to 90 °C and stirred for 7 days. The mixture was cooled to room temperature, solvent was removed, and the residue purified by flash column chromatography on silica (EtOAc:MeOH (0.1% formic acid) = 0% to 25% over 8 CVs) to afford the title compound as an opaque oil (0.79 g, 87%).

**$^1$H NMR** (598 MHz, DMSO) δ 8.64 (d, $J$ = 4.9 Hz, 1H), 8.59 (s, 1H), 8.50 (d, $J$ = 5.6 Hz, 1H), 7.93 (s, 1H), 7.57 (d, $J$ = 3.1 Hz, 1H), 7.37 – 7.27 (m, 5H), 7.04 – 7.01 (m, 1H), 5.02 (s, 2H), 4.29 (t, 2H), 3.80 (t, $J$ = 4.6 Hz, 2H), 3.64 – 3.60 (m, 2H), 3.54 – 3.49 (m, 6H), 3.43 (t, $J$ = 6.0 Hz, 2H), 3.16 (t, $J$ = 6.0 Hz, 2H).
**$^{13}$C NMR** (150 MHz, DMSO) δ 166.07, 157.51, 157.31, 155.79, 151.06, 149.62, 137.74, 128.75, 128.13, 128.05, 120.88, 117.78, 111.45, 107.07, 72.84, 70.52, 70.34, 70.29, 70.25, 70.10, 69.66, 69.17, 67.99, 65.73, 60.81.
**LC-MS** $R_t$ = 4.69 min; m/z calcd. For $C_{26}H_{30}N_3O_9S$ ($[M+H]^+$) = 561.2, found 562.3.

**9** $^{1}$H-NMR

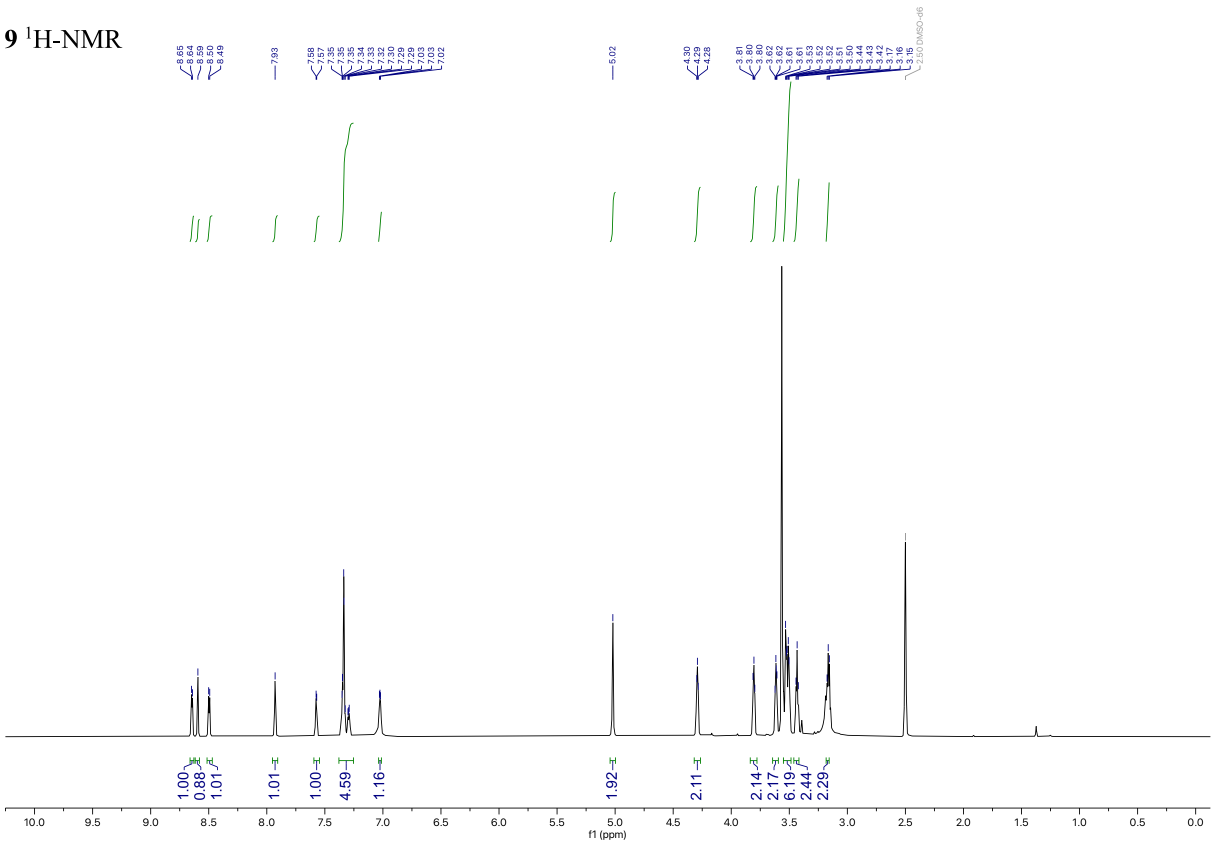

**9** $^{13}$C-NMR

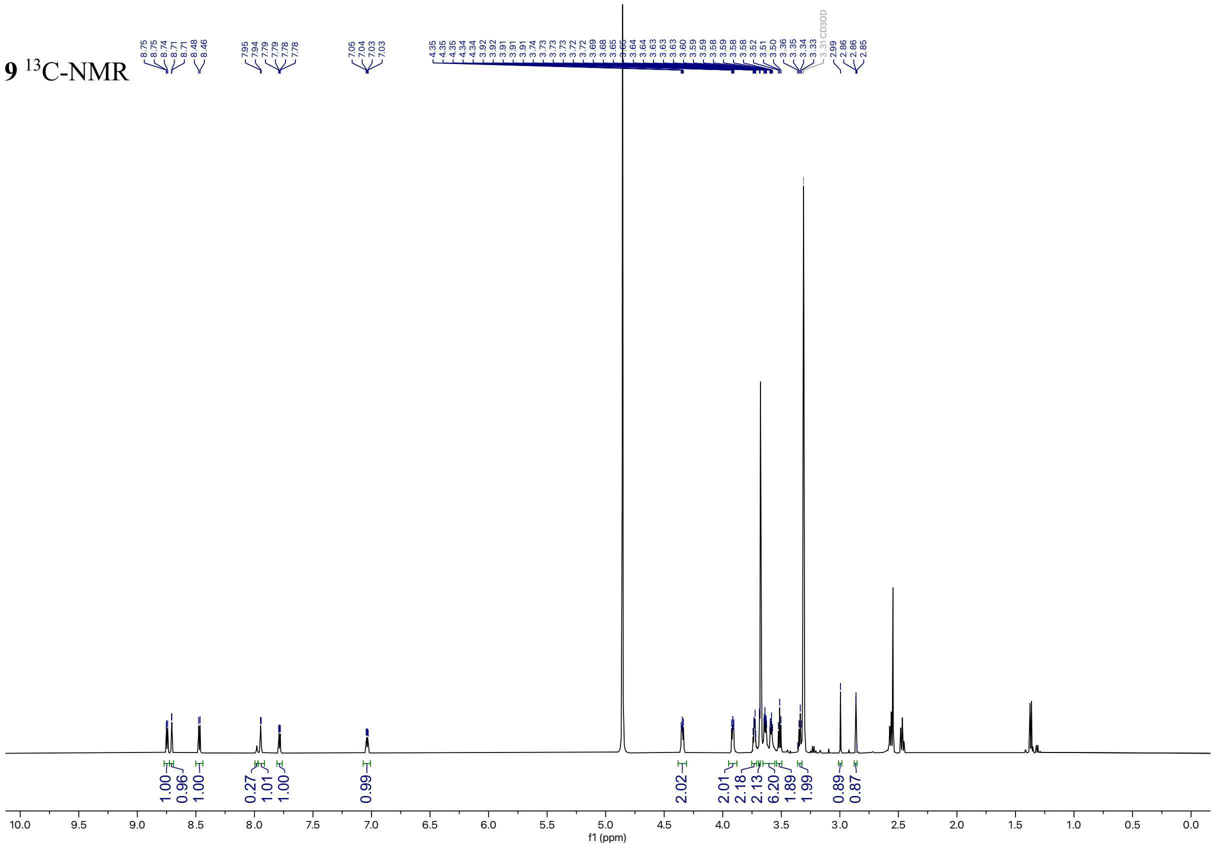

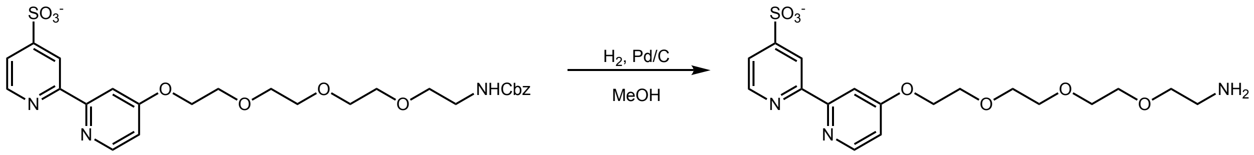

**4’-(2-(2-(2-(2-Aminoethoxy)ethoxy)ethoxy)ethoxy)-[2,2’-bipyridine]-4-sulfonate (10)**
A scintillation vial equipped with a PTFE-coated stir bar was charged with **9** (0.63 g, 0.41 mmol, 1.0 equiv.) and 10% Pd/C, type 487 (4.5 mg) and MeOH (4 mL) added. The solution was placed under $H_2$ atmosphere using a balloon and stirred at room temperature for 16 hours. The mixture was filtered over celite and the filter cake was washed with MeOH (150 mL). Solvent was removed and the title compound was carried forward without additional purification.

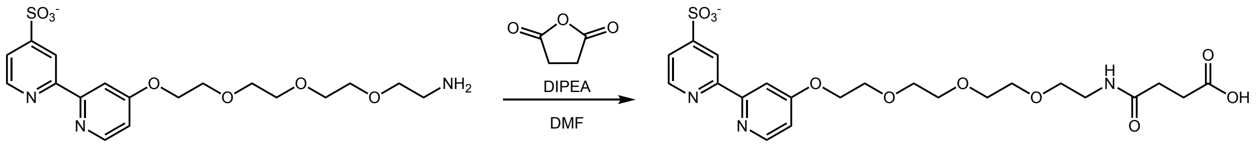

**4’-((15-Carboxy-13-oxo-3,6,9-trioxa-12-azapentadecyl)oxy)-[2,2’-bipyridine]-4-sulfonate (11)**
A scintillation vial equipped with a PTFE-coated stir bar was charged with **10** (0.19 g, 0.44 mmol, 1.0 equiv.), succinic anhydride (0.048 g, 0.49 mmol, 1.1 equiv.), and DMF (3 mL). The reaction was stirred at room temperature for 5 minutes. DIPEA (0.38 mL, 2.23 mmol, 5.0 equiv.) was added and the reaction was stirred for 1 hour. Solvent was then removed to afford the title compound without additional purification.

**$^{1}$H NMR** (500 MHz, $CD_3OD$_SPE) δ 8.77 – 8.73 (m, 1H), 8.71 (d, $J$ = 0.8 Hz, 1H), 8.47 (d, $J$ = 5.7 Hz, 1H), 7.94 (d, $J$ = 2.6 Hz, 1H), 7.79 (dd, $J$ = 5.0, 1.6 Hz, 1H), 7.04 (dd, $J$ = 5.8, 2.6 Hz, 1H), 4.38 – 4.31 (m, 2H), 3.95 – 3.88 (m, 2H), 3.73 (dt, $J$ = 4.8, 3.0 Hz, 2H), 3.68 (d, $J$ = 2.0 Hz, 2H), 3.66 – 3.56 (m, 6H), 3.51 (t, $J$ = 5.5 Hz, 2H), 3.34 (t, $J$ = 5.6 Hz, 2H).
**$^{13}$C NMR** (126 MHz, $CD_3OD$_SPE) δ 177.09, 176.49, 174.71, 167.84, 158.31, 157.78, 155.73, 151.66, 150.81, 121.65, 119.12, 112.32, 108.88, 71.81, 71.61, 71.59, 71.53, 71.29, 70.56, 70.48, 69.92, 68.95, 57.65, 57.48, 57.30, 55.85, 52.15, 49.63, 49.45, 49.28, 49.11, 43.79, 40.43, 31.71, 30.74, 30.62, 30.24, 30.04, 17.43, 17.28, 17.13, 13.17.
**LC-MS** $R_t$ = 3.73 min; m/z calcd. For $C_{22}H_{28}N_3O_{10}S$ ($[M+H]^+$) = 528.2, found 528.3

**11** $^{1}$H-NMR

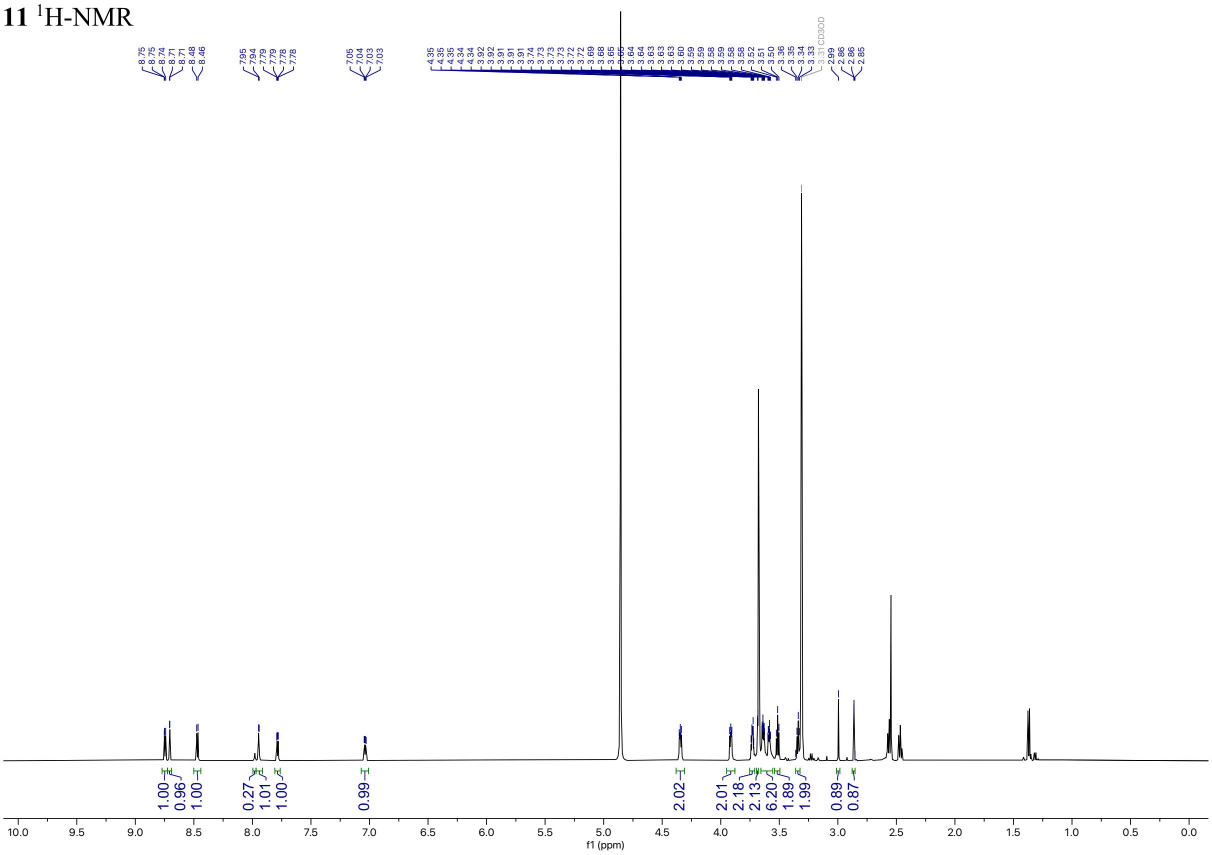

**11** $^{13}$C-NMR

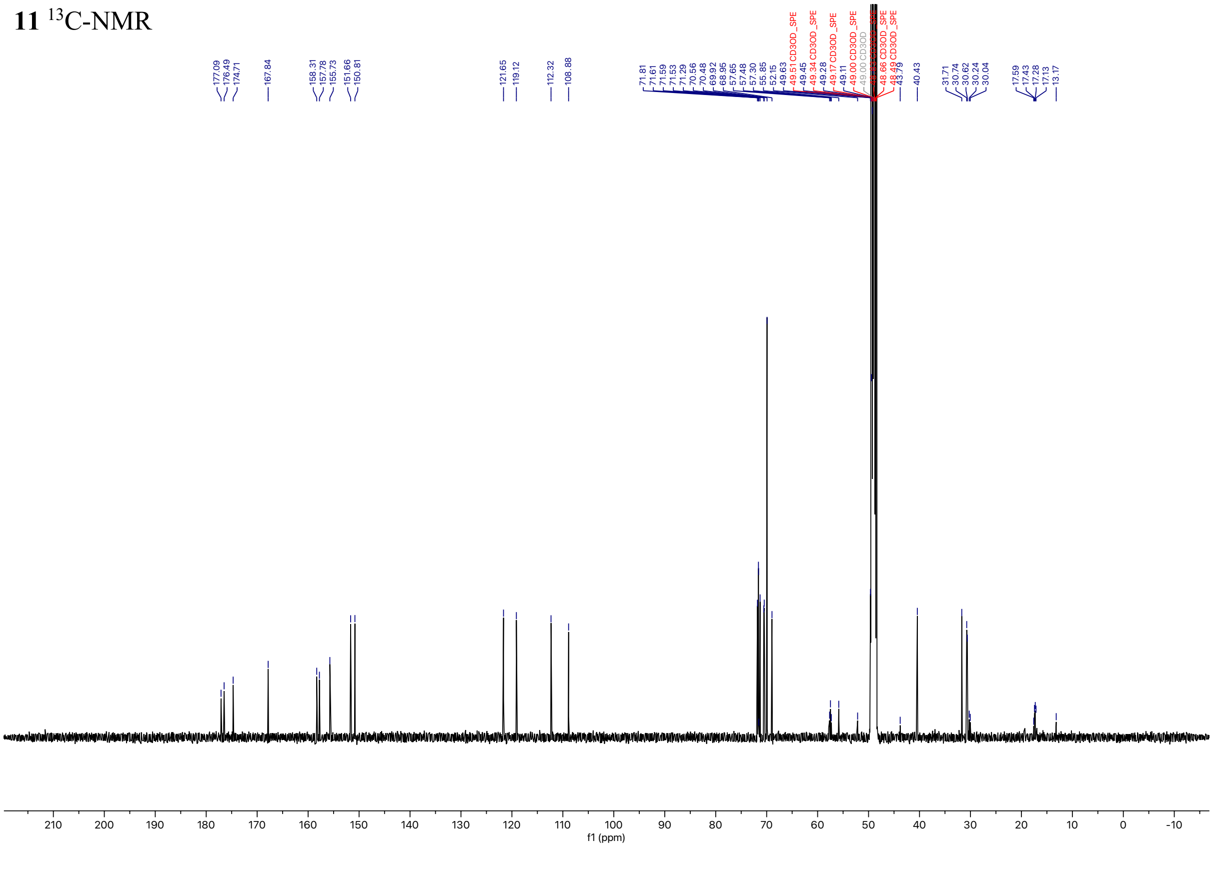

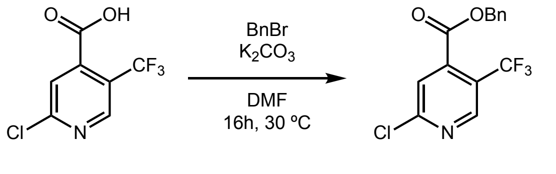

**Benzyl 2-chloro-5-(trifluoromethyl)isonicotinate (12).**
A round-bottom flask equipped with a PTFE-coated stir bar was charged with 2-chloro-5-(trifluoromethyl)isonicotinic acid (7.0 g, 31.0 mmol, 1.0 equiv.), potassium carbonate (5.1 g, 37.2 mmol, 1.2 equiv.), and DMF (51 mL). The solution was cooled to 0 °C for 15 minutes and benzyl bromide (3.7 mL, 31.0 mmol, 1.0 equiv.) was added dropwise. The reaction was stirred at 0 °C for 30 minutes then warmed to room temperature and stirred for 3 hours. The solvent was removed and the crude mixture partitioned between EtOAc and $H_2O$. The aqueous layer was extracted 4x with EtOAc (125 mL). The EtOAc extracts were washed with brine, dried with $Na_2SO_4$, and concentrated to remove solvent. The residue was purified by flash column chromatography on silica (hexanes:EtOAc = 0% to 10% over 7 CVs) to afford the title compound as a white powder (8.5 g, 87%).

**$^1$H NMR** (500 MHz, $CDCl_3$) δ 8.76 (s, 1H), 7.67 (s, 1H), 7.40 (dt, *J* = 22.2, 7.3 Hz, 5H), 5.39 (s, 2H).
**$^{19}$F NMR** (471 MHz, $CDCl_3$) δ -59.27.
**LC-MS** $R_t$ = 7.51 min; m/z calcd. For $C_{14}H_9ClF_3NO_2$ ($[M+H]^+$) = 316.0, found 316.1

**12** $^{1}$H-NMR

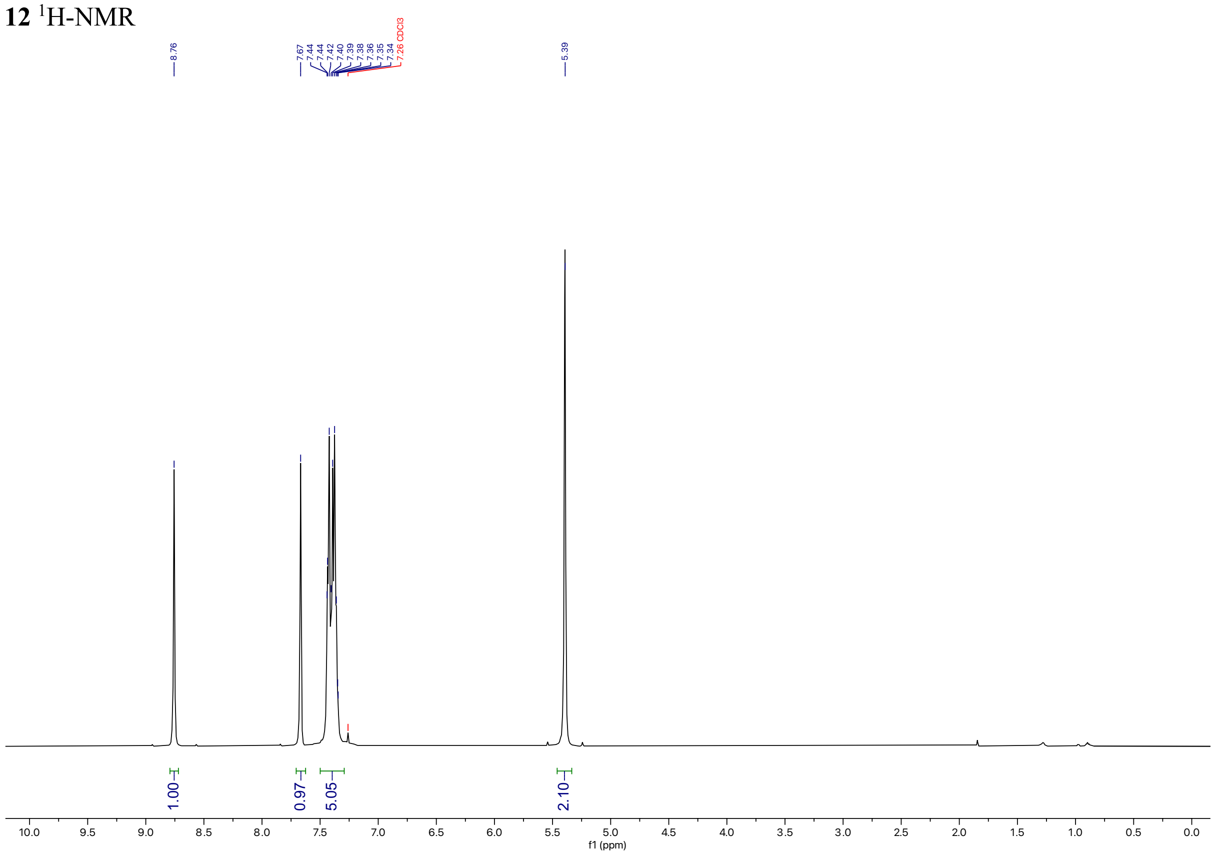

**12** $^{19}$F-NMR

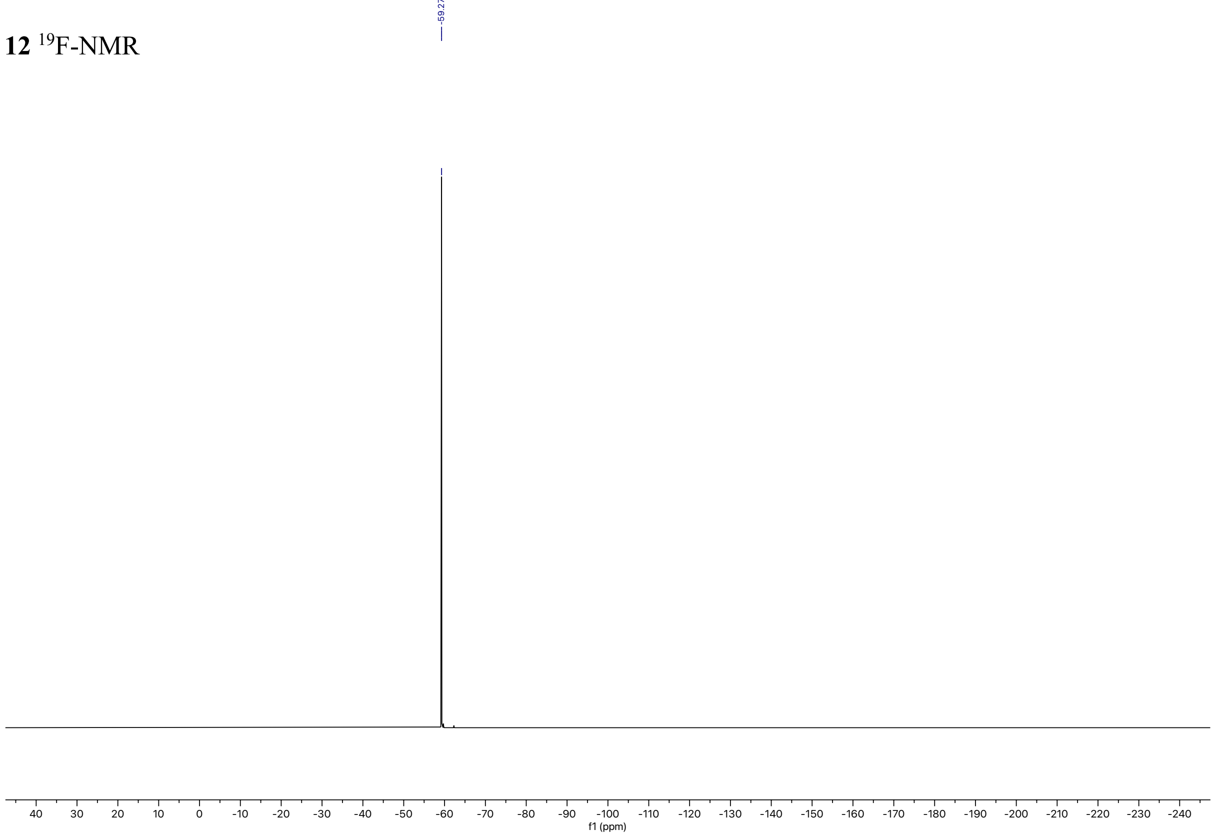

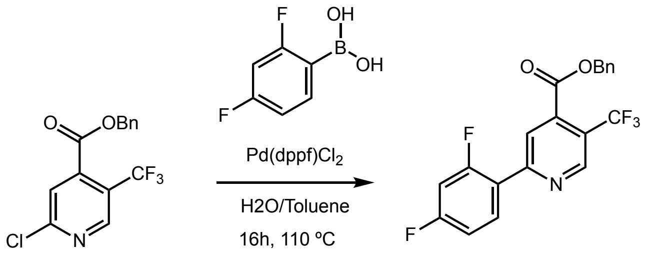

**Benzyl 2-(2,4-difluorophenyl)-5-(trifluoromethyl)isonicotinate (13).**
A flame-dried three-neck round-bottom flask equipped with a PTFE-coated stir bar was charged with **12** (6.04 g, 19.2 mmol, 1.0 equiv.), 2,4-difluorophenyl boronic acid (3.33 g, 21.1 mmol, 1.1 equiv.) potassium carbonate (7.95 g, 57.5 mmol, 3.0 equiv.), and [1,1′-bis(diphenylphosphino)ferrocene]dichloropalladium(II) dichloromethane complex (0.42 g, 0.58 mmol, 0.03 equiv.). The flask was placed under argon atmosphere, toluene:$H_2O$ (2:1, 75 mL) was added, and the solvent further sparged for 10 minutes. The solution was heated to 85 °C and stirred for 16 hours. The reaction was cooled to room temperature and quenched with cold water (200 mL) and the crude mixture partitioned between EtOAc and water. The aqueous layer was extracted 3x with EtOAc (100 mL) and the EtOAc extracts were washed with brine, dried with $Na_2SO_4$, and concentrated to remove solvent. The residue was purified by flash column chromatography on silica (hexanes:DCM = 0% to 25% over 12 CVs) to afford the title compound as a white powder (6.6 g, 88%).

**$^1$H NMR** (500 MHz, $CDCl_3$) δ 9.05 (s, 1H), 8.11 (dd, $J$ = 10.1, 5.1 Hz, 2H), 7.48 – 7.34 (m, 5H), 7.05 (td, $J$ = 8.2, 2.4 Hz, 1H), 6.96 (ddd, $J$ = 11.3, 8.6, 2.4 Hz, 1H), 5.42 (s, 2H).
**$^{19}$F NMR** (471 MHz, $CDCl_3$) δ -59.25 (s, 3F), -105.97 – -106.05 (m, 1F), -111.21 (q, $J$ = 9.8 Hz, 1F).
**LC-MS** $R_t$ = 7.98 min; m/z calcd. For $C_{20}H_{12}F_5NO_2$ ($[M+H]^+$) = 394.1, found 394.2

**13** $^{1}$H-NMR

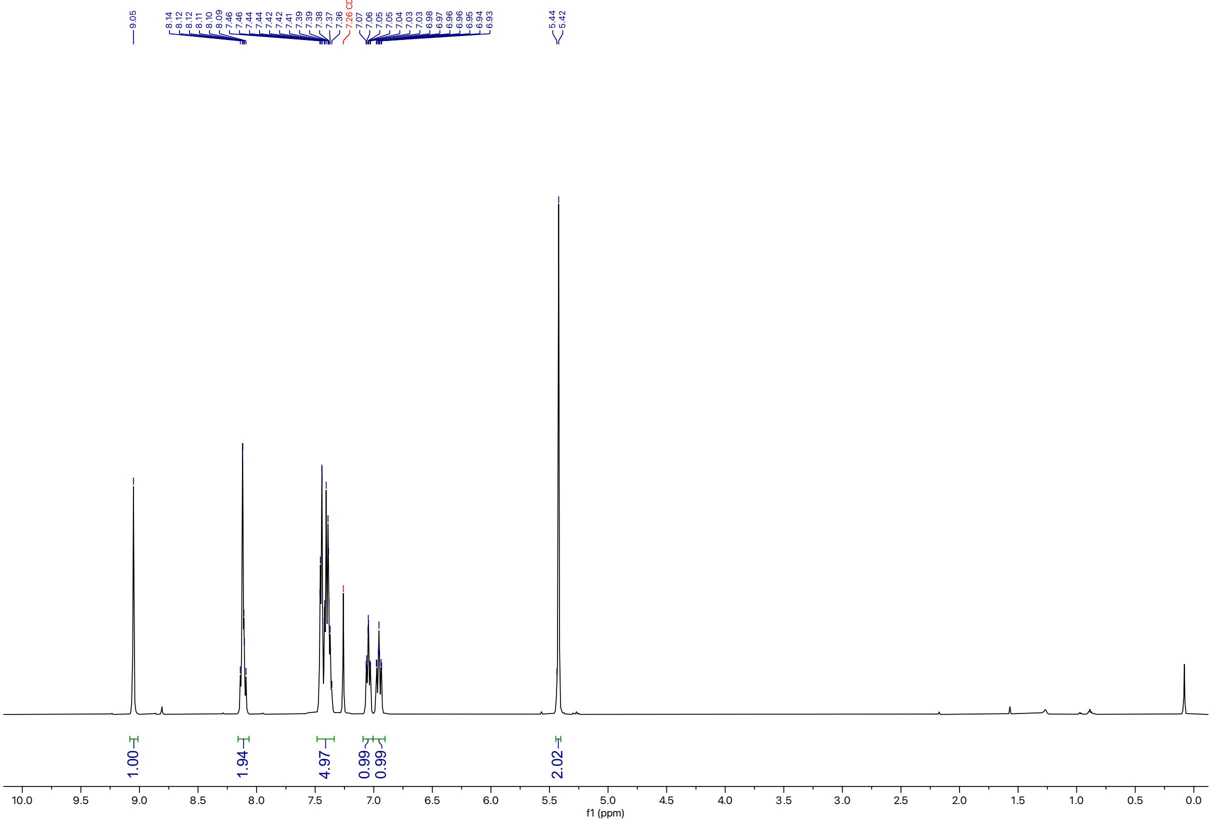

**13** $^{19}$F-NMR

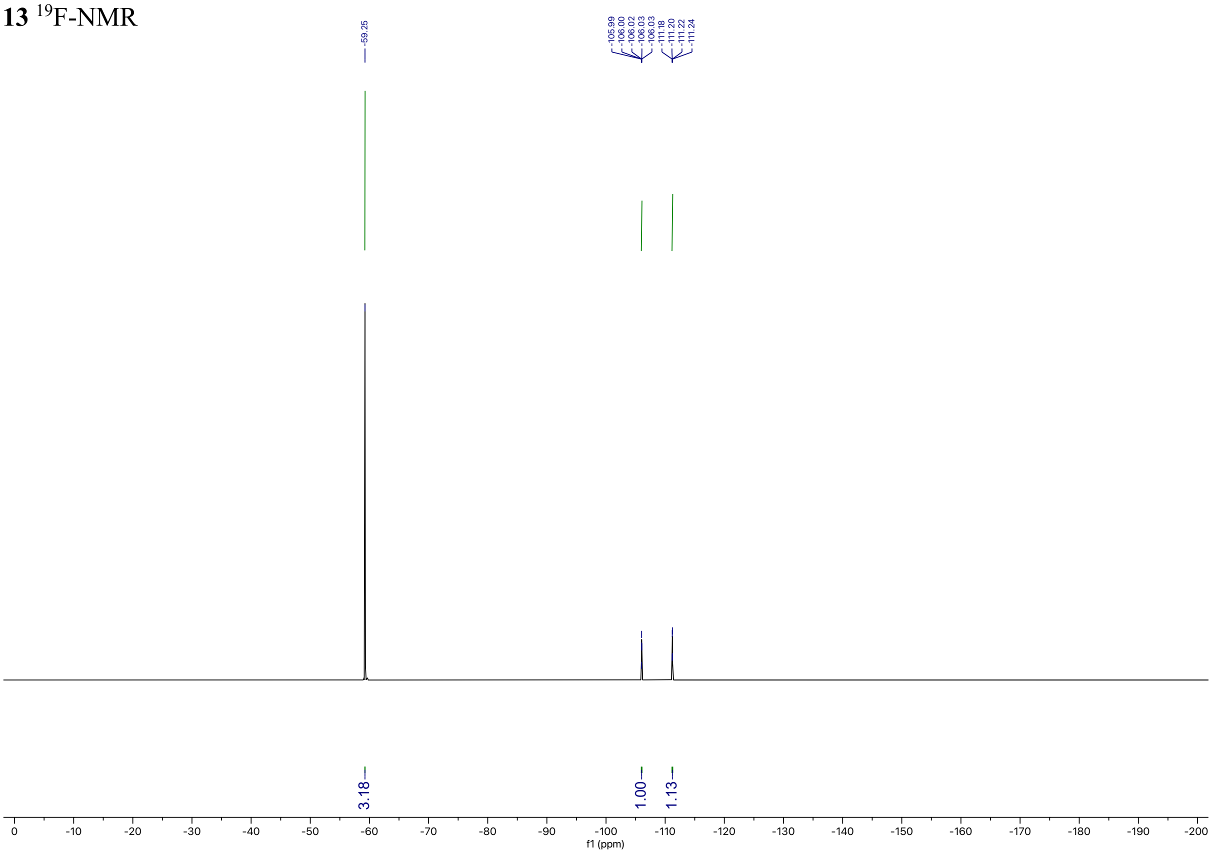

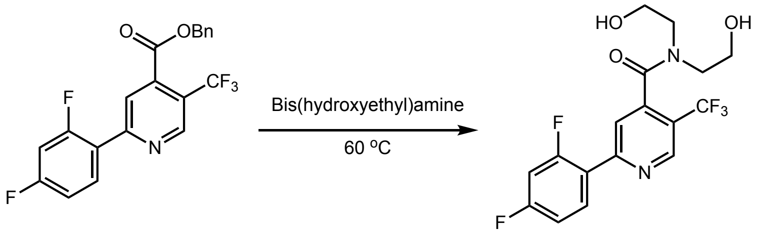

**2-(2,4-Difluorophenyl)-N,N-bis(2-hydroxyethyl)-5-(trifluoromethyl)isonicotinamide (14, dF($CF_3$)(OBN)ppy).** A scintillation vial equipped with a PTFE-coated stir bar was charged with **13** (4.6 g, 11.7 mmol, 1.0 equiv.) and bis(hydroxyethyl)amine (5.6 mL, 58.5 mmol, 5.0 equiv.). The mixture was heated to 70 °C and stirred for 16 hours. The reaction was cooled to room temperature and partitioned between EtOAc and water, and the aqueous layer acidified to pH 7 with HCl (1 M). The aqueous layer was extracted 3x with EtOAc (100 mL) and the EtOAc extracts were washed with brine, dried over $Na_2SO_4$, and concentrated to remove solvent. The residue was purified by flash column chromatography on silica (DCM:MeOH = 0% to 10% over 8 CVs) to afford the title compound as an off-white powder (4.04 g, 88%).

**$^1$H NMR** (500 MHz, $CDCl_3$) δ 9.00 (s, 1H), 8.10 (td, $J$ = 8.8, 6.5 Hz, 1H), 7.84 (s, 1H), 7.04 (td, 1H), 6.93 (ddd, $J$ = 11.3, 8.6, 2.5 Hz, 1H), 4.19 – 3.94 (m, 2H), 3.90 – 3.75 (m, 2H), 3.66 (dt, $J$ = 10.6, 4.5 Hz, 2H), 3.31 – 3.16 (m, 2H), 2.71 (br s, 1H).
**$^{13}$C NMR** (126 MHz, $CDCl_3$) δ 168.07, 165.23, 165.13, 163.21, 163.12, 162.09, 161.99, 160.07, 159.97, 156.21, 147.85, 147.81, 147.77, 147.74, 143.26, 143.24, 132.57, 132.54, 132.49, 132.46, 126.51, 124.33, 122.16, 122.07, 121.89, 121.86, 121.80, 121.77, 121.03, 120.77, 120.51, 120.25, 112.55, 112.52, 112.38, 112.35, 104.95, 104.74, 104.54.
**$^{19}$F NMR** (471 MHz, $CDCl_3$) δ -59.77 (s, 3F), -106.08 – -106.28 (m, 1F), -111.48 (q, $J$ = 9.7 Hz, 1F).
**LC-MS** $R_t$ = 5.11 min; m/z calcd. For $C_{17}H_{15}F_5N_2O_3$ $C_{17}H_{15}F_5N_2O_3$ ([M+H]$^+$) = 391.1, found 391.5

**14** $^{1}$H-NMR

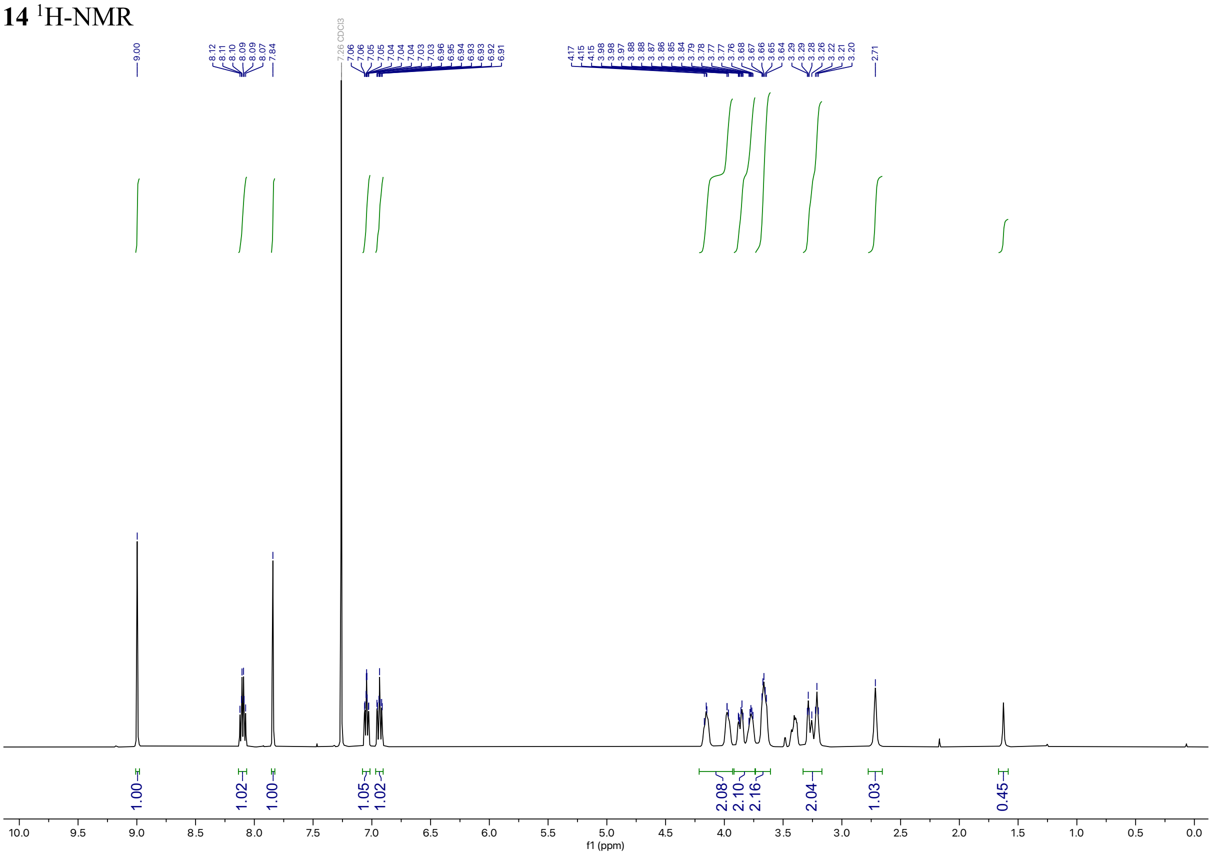

**14** $^{13}$C-NMR

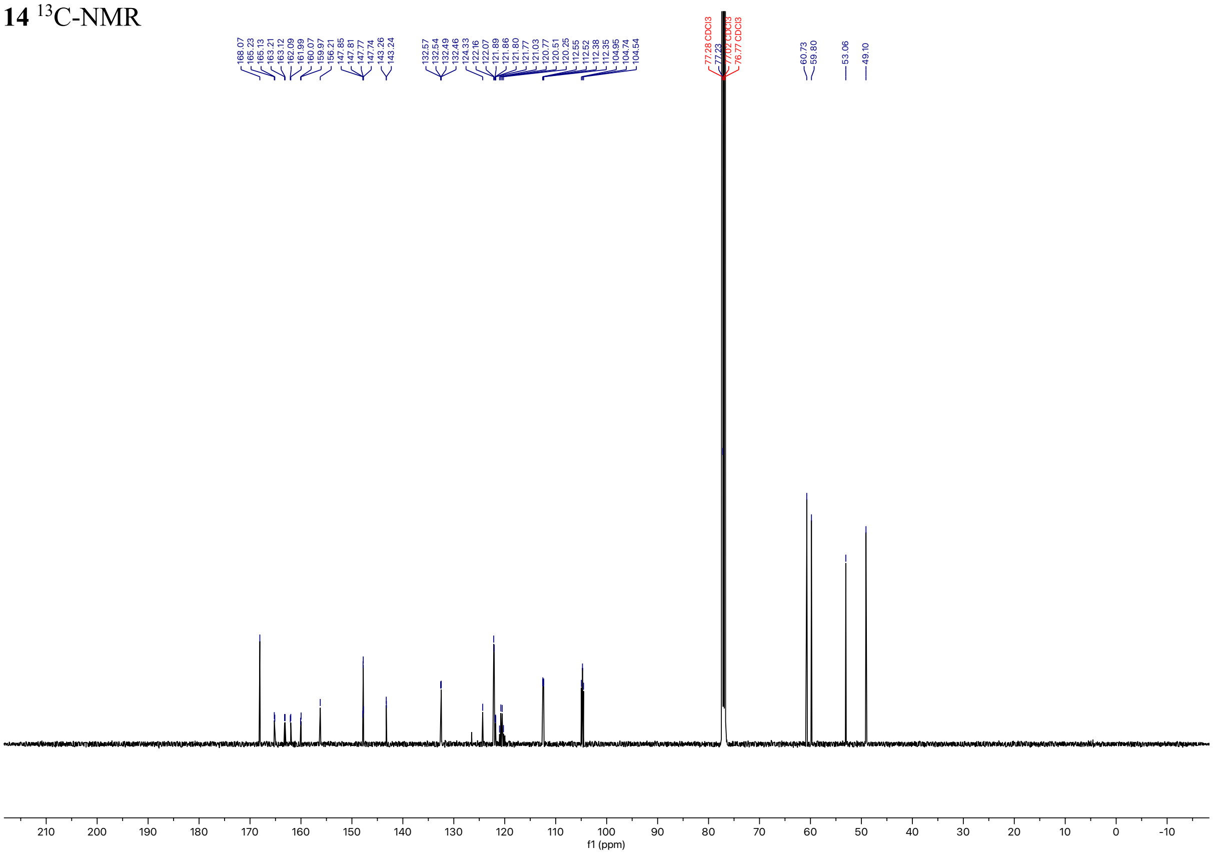

**14** $^{19}$F-NMR

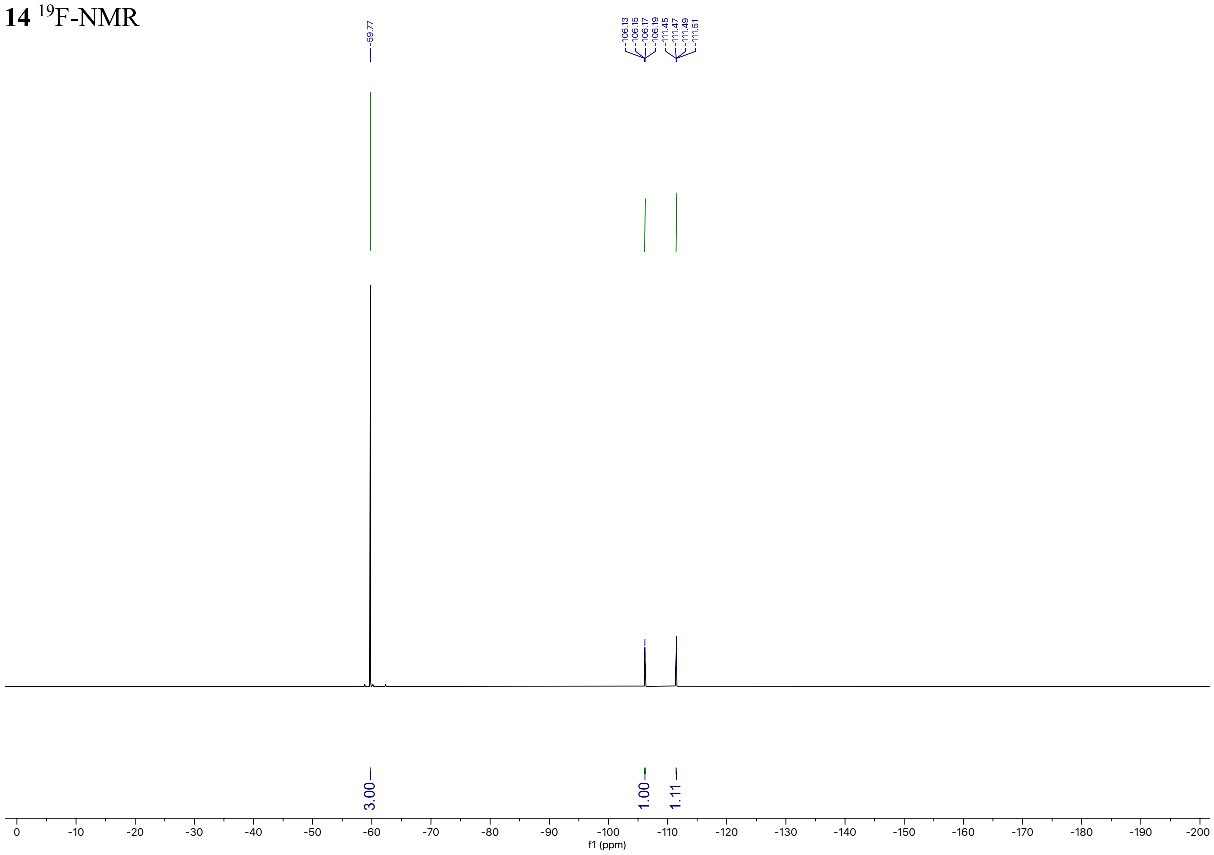
-59.77
-106.13
-106.15
-106.17
-106.19
-111.45
-111.47
-111.49
-111.51
3.00
1.00
1.11
0
-10
-20
-30
-40
-50
-60
-70
-80
-90
-100
-110
-120
-130
-140
-150
-160
-170
-180
-190
-200
f1 (ppm)

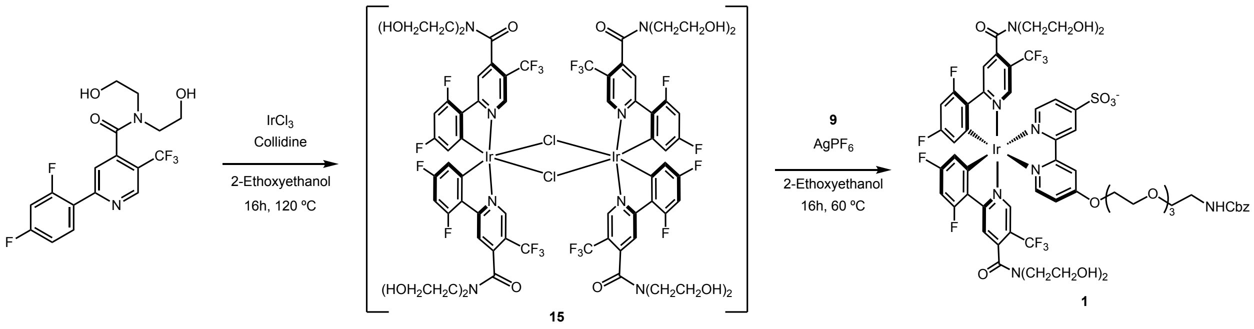

**Ir[dF(CF$_3$)(OBN)ppy] Dimer (15)**

An oven-baked scintillation vial equipped with a PTFE-coated stir bar was charged with **14** (0.1 g, 0.256 mmol, 2.5 equiv.), 2,4,6-Trimethylpyridine (0.050 g, 0.410 mmol, 4 equiv.) and iridium(III) chloride hydrate (0.032 g, 0.103 mmol, 1.0 equiv.). To the mixture 2-ethoxyethanol (4.364 mL, 23.5 mM) was added and the vial was placed under an argon atmosphere by degassing the solvent. The vial was sealed and heated to 120 °C and stirred for 16 hours. The reaction was cooled to room temperature and carried forward without further purification.

**Ir[dF(CF$_3$)(OBN)ppy]bipy-NHCbz complex (1)**

The scintillation vial from the previous reaction containing **15** in 2-ethoxyethanol was charged with **9** (0.690 g, 0.123 mmol, 1.2 equiv.), and silver hexafluorophosphate (0.129 g, 0.513 mmol, 5.0 equiv.). The mixture was heated to 60 °C and stirred for 16 hours. The reaction was cooled to room temperature, filtered over celite, washed with MeOH/acetone (1:1, 70 mL) and the solvent was removed. The crude mixture was purified by flash column chromatography on silica (DCM:MeOH, 0% to 25% over 35 CVs) to afford the title compound as yellow solid (0.152 g, 69% over 2 steps).

**$^1$H NMR** (500 MHz, DMSO) δ 9.06 (s, 1H), 8.58 (s, 1H), 8.50 (br s, 1H), 8.35 (s, 2H), 8.03 (s, 1H), 7.86 (s, 1H), 7.69 (s, 1H), 7.62 (s, 1H), 7.55 (s, 1H), 7.39 – 7.27 (m, 6H), 7.23 (t, $J$ = 5.8 Hz, 1H), 7.12 – 7.00 (m, 1H), 6.69 (s, 1H), 5.98 – 5.54 (m, 2H), 4.98 (s, 3H), 4.83 (t, $J$ = 5.1 Hz, 2H), 4.50 (s, 2H), 3.80 (t, $J$ = 4.4 Hz, 3H), 3.64 – 3.46 (m, 16H), 3.12 (q, $J$ = 5.9 Hz, 4H).
**$^{19}$F NMR** (471 MHz, DMSO) δ -59.69 (s, 6F), -102.68 (t, $J$ = 97.4 Hz, 2F), -105.68 – -106.46 (m, 2F).
**LC-MS** $R_t$ = 5.34 min; m/z calcd. For $C_{60}H_{58}F_{10}IrN_7O_{15}S$ ($[M+H]^+$) = 1532.3, found 1532.6

**1** $^{1}$H-NMR

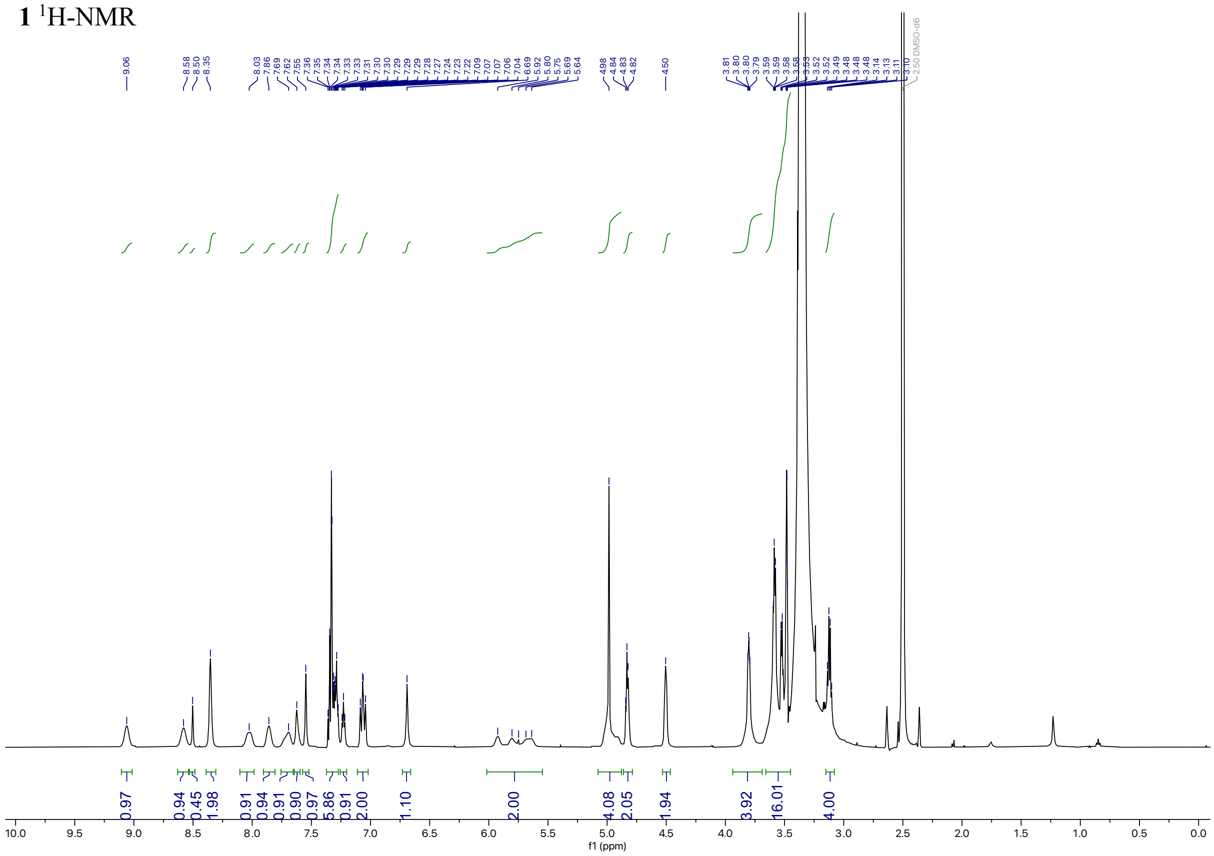

**1** $^{19}$F-NMR

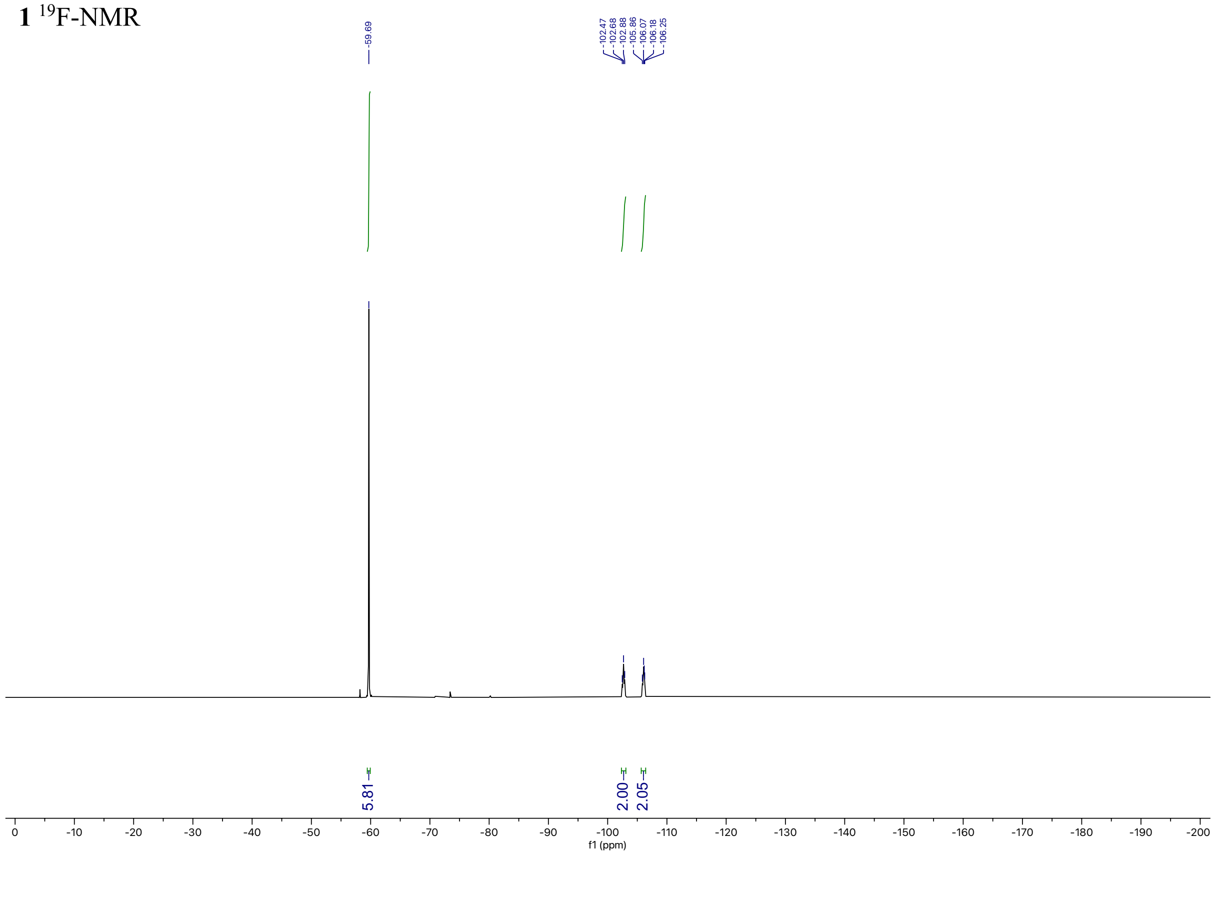

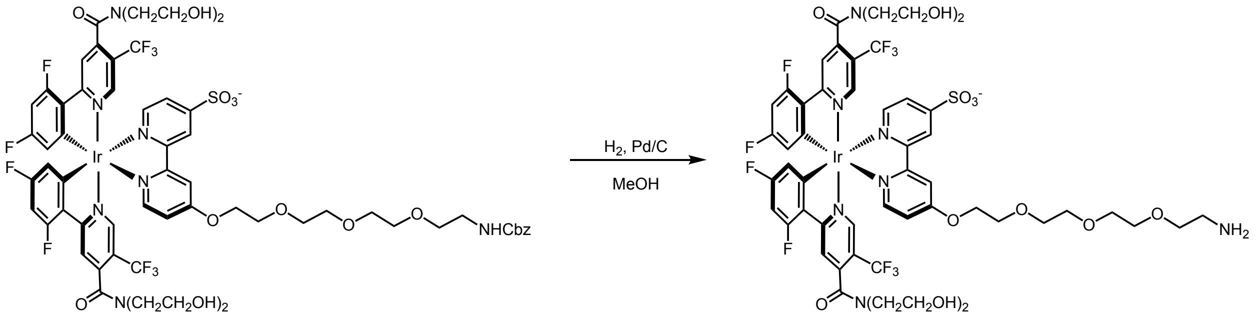

**1-NH₂**

A scintillation vial equipped with a PTFE-coated stir bar was charged with **1** (0.154 g, 0.1 mmol, 1.0 equiv.) and 10% Pd/C, type 487 (1 mg) and suspended in MeOH (1 mL). The mixture was placed under $H_2$ atmosphere using a balloon and stirred at room temperature for 16 hours. The mixture was filtered over celite and the filter cake was washed with MeOH (50 mL). Solvent was removed to afford the title compound which was carried forward without additional purification.

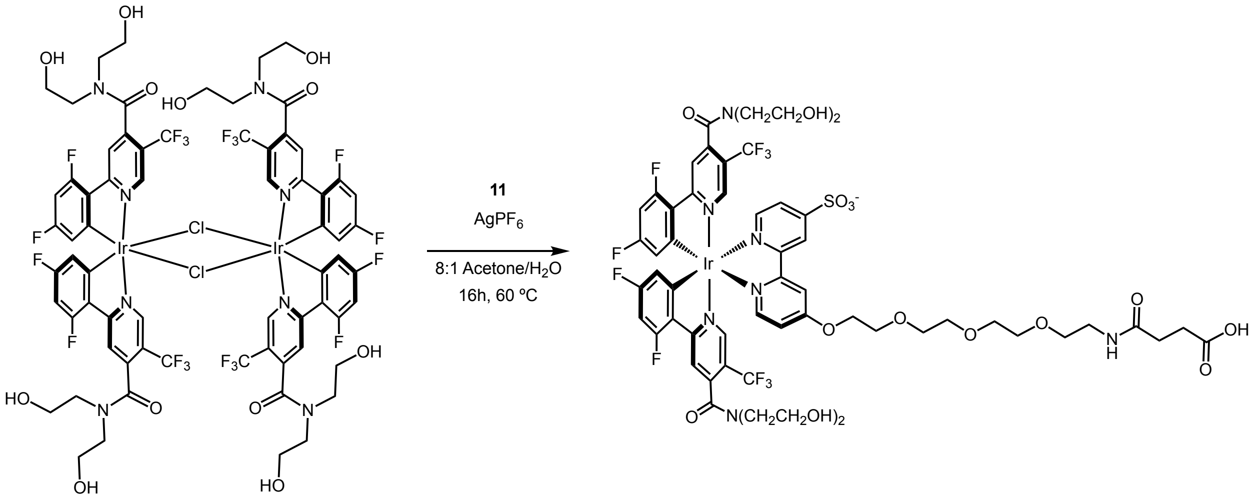

**1-CO₂H**

A scintillation vial equipped with a PTFE-coated stir bar was charged with **15** (0.788 g, 0.39 mmol, 1.0 equiv.), **11** (0.482 g, 0.86 mmol, 2.2 equiv.), and silver hexafluorophosphate (0.297 g, 1.17 mmol, 3.0 equiv.). Acetone:$H_2O$ (8:1, 7.8 mL) was then added, the mixture heated to 60 °C, and stirred at this temperature for 16 hours. The reaction was cooled to room temperature, filtered, and the solvent was removed. The crude mixture was purified by flash column chromatography on silica (EtOAc:MeOH (+0.1% formic acid) = 0% to 35% over 7 CVs) to afford the title compound as yellow solid (0.62 g, >95%).

**$^1$H NMR** (500 MHz, DMSO) δ 9.05 (s, 1H), 8.58 (s, 1H), 8.34 (s, 2H), 8.11 – 7.48 (m, 7H), 7.29 (s, 1H), 7.05 (ddt, $J$ = 12.1, 9.5, 2.5 Hz, 2H), 5.75 (dd, $J$ = 104.6, 41.1 Hz, 2H), 5.16 – 4.62 (br m, 4H), 4.58 – 4.46 (m, 2H), 3.84 – 3.74 (m, 4H), 3.61 – 3.55 (m, 6H), 3.52 (dd, $J$ = 5.9, 3.7 Hz, 2H), 3.49 – 3.42 (m, 8H), 3.14 (q, $J$ = 5.9 Hz, 4H), 2.59 (t, $J$ = 5.7 Hz, 2H), 2.37 – 2.21 (m, 6H).
**$^{19}$F NMR** (471 MHz, DMSO) δ -59.69 (s, 6F), -102.16 – -103.22 (m, 2F), -105.47 – -106.60 (m, 2F).
**LC-MS** $R_t$ = 4.68 min; m/z calcd. For $C_{56}H_{56}F_{10}IrN_7O_{16}S$ ($[M+H]^+$) = 1498.3, found 1498.8

**1-CO₂H** $^{1}$H-NMR

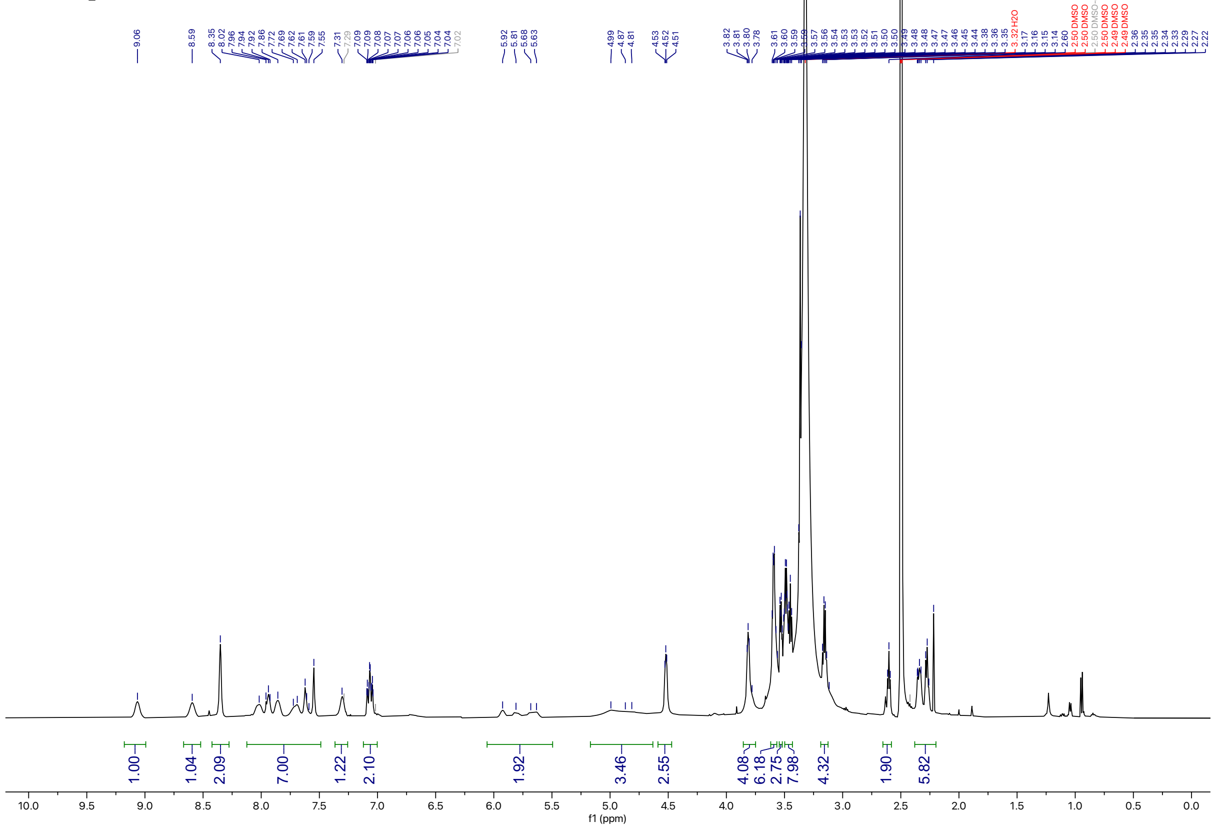

**1-CO₂H** $^{19}$F-NMR

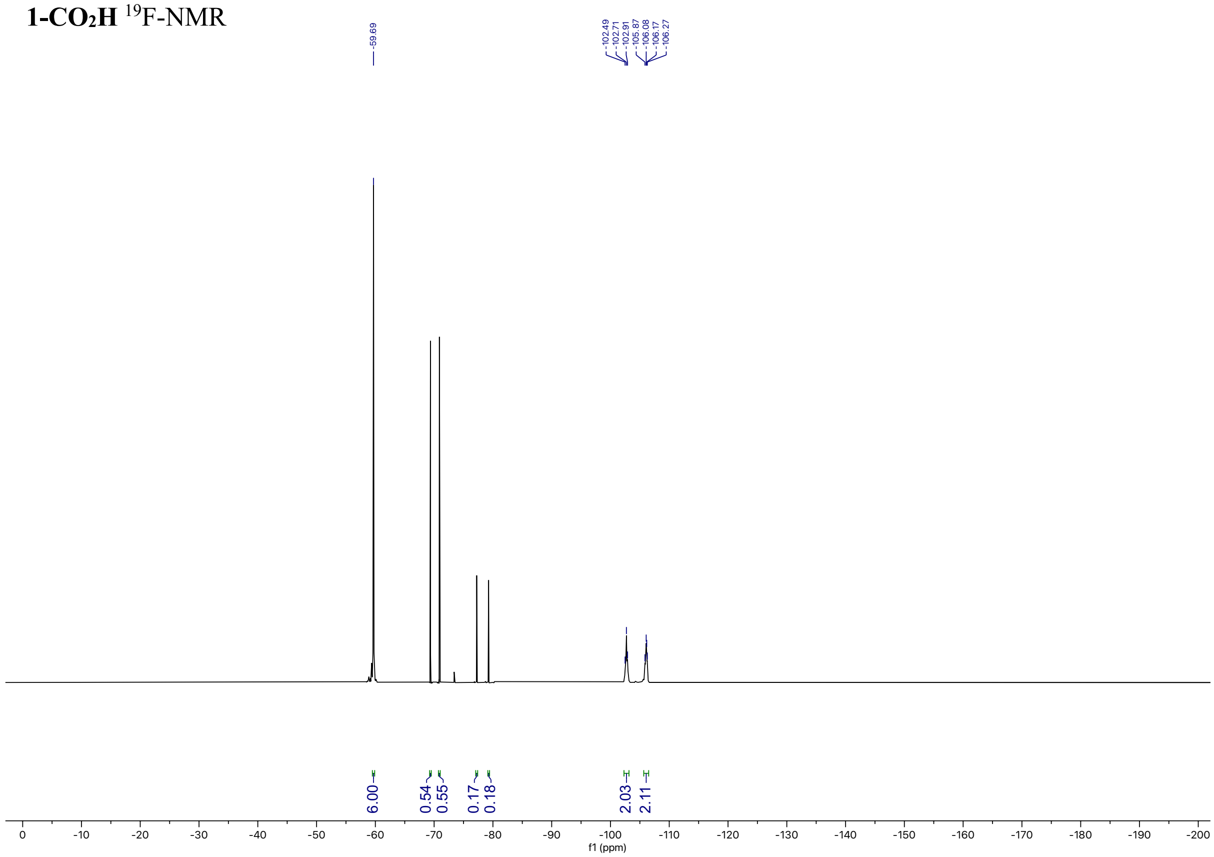

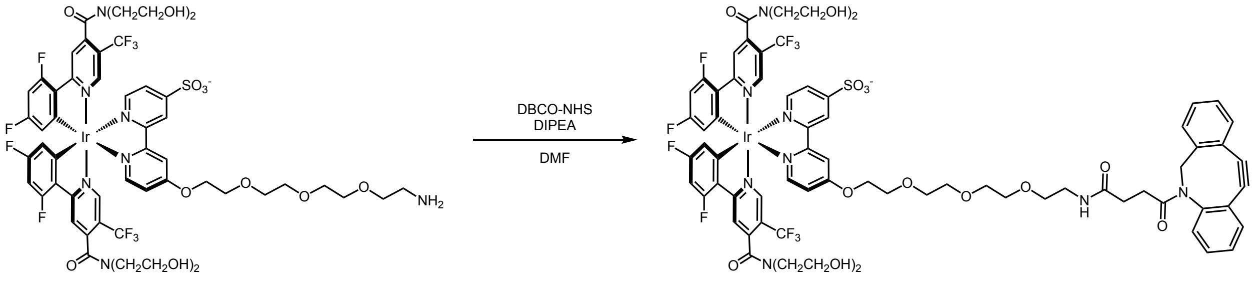

**1-DBCO**

A scintillation vial equipped with a PTFE-coated stir bar was charged with **1-NH₂** (0.025 g, 0.018 mmol, 1.0 equiv.), dibenzocyclooctyne-*N*-hydroxysuccinimidyl ester (DBCO-NHS, 8.0 mg, 0.021 mmol, 1.2 equiv.), and dissolved in DMF (0.4 mL). The reaction was stirred at room temperature for 5 minutes and DIPEA (0.03 mL, 0.09 mmol, 5.0 equiv.) was added. After 16 hours the reaction mixture was diluted with MeOH/$H_2O$ (1:1, 0.4 mL) and purified by preparative HPLC using a 45% solvent B 15-minute isocratic method. The product eluted at 4.25 minutes and was concentrated to afford the title compound as a yellow solid (4.8 mg, 16%).

**$^1$H NMR** (598 MHz, DMSO) δ 9.00 (dd, *J* = 35.7, 20.5 Hz, 1H), 8.65 – 8.25 (m, 4H), 8.00 – 7.91 (m, 1H), 7.85 (d, *J* = 24.9 Hz, 3H), 7.71 (d, *J* = 16.4 Hz, 3H), 7.59 (d, *J* = 33.4 Hz, 4H), 7.46 (s, 1H), 7.31 (s, 3H), 6.90 (d, *J* = 10.9 Hz, 2H), 6.27 (br, s, 1H), 6.13 (s, 1H), 5.60 – 5.39 (m, 1H), 4.71 (br, s, 3H), 4.52 (s, 2H), 3.90 – 3.35 (m, 34H).
**$^{19}$F NMR** (563 MHz, DMSO) δ -61.02 – -62.06 (m, 6F), -95.36 – -97.44 (m, 2F), -102.30 – -103.82 (m, 2F).
**LC-MS** $R_t$ = 5.54 min; m/z calcd. For $C_{71}H_{65}F_{10}IrN_8O_{15}S$ ($[M+H]^+$) = 1685.4, found 1685.6

**1-DBCO** $^{1}$H-NMR

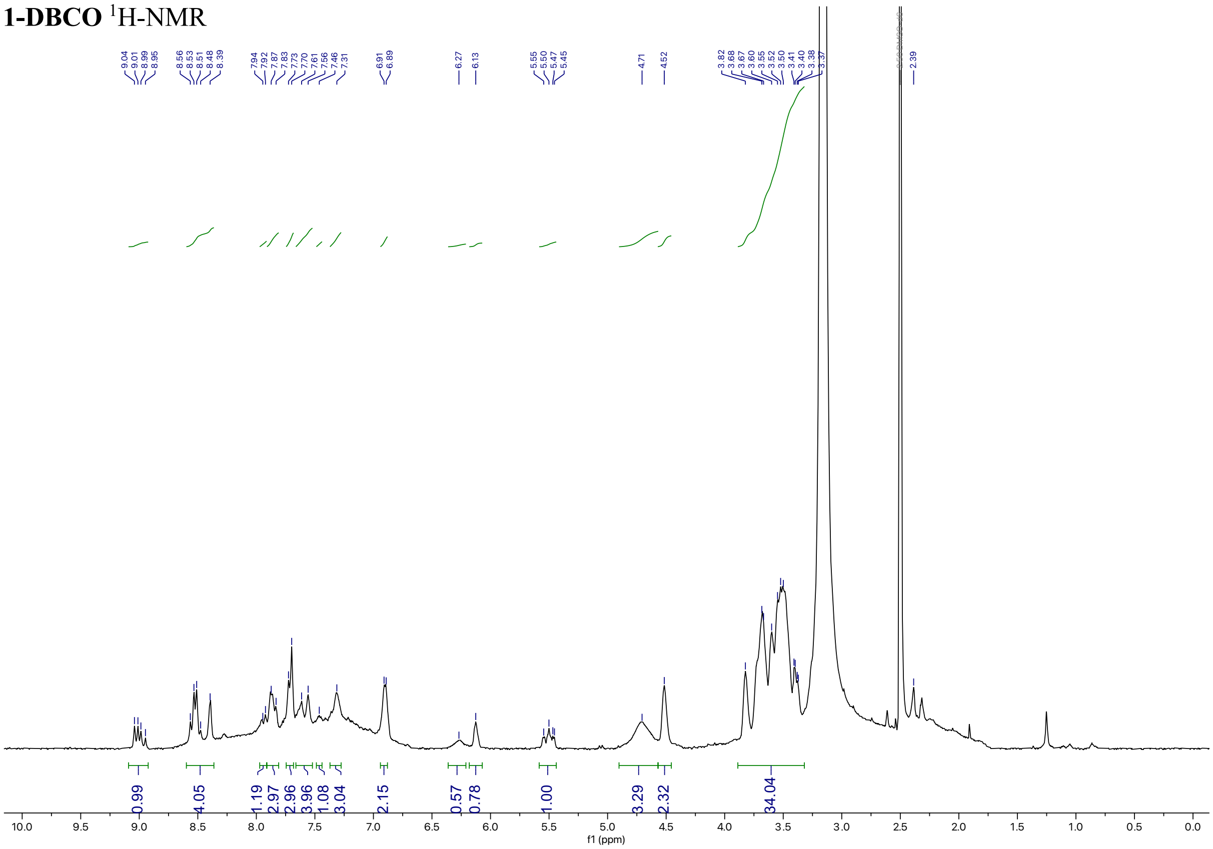

**1-DBCO** $^{19}$F-NMR

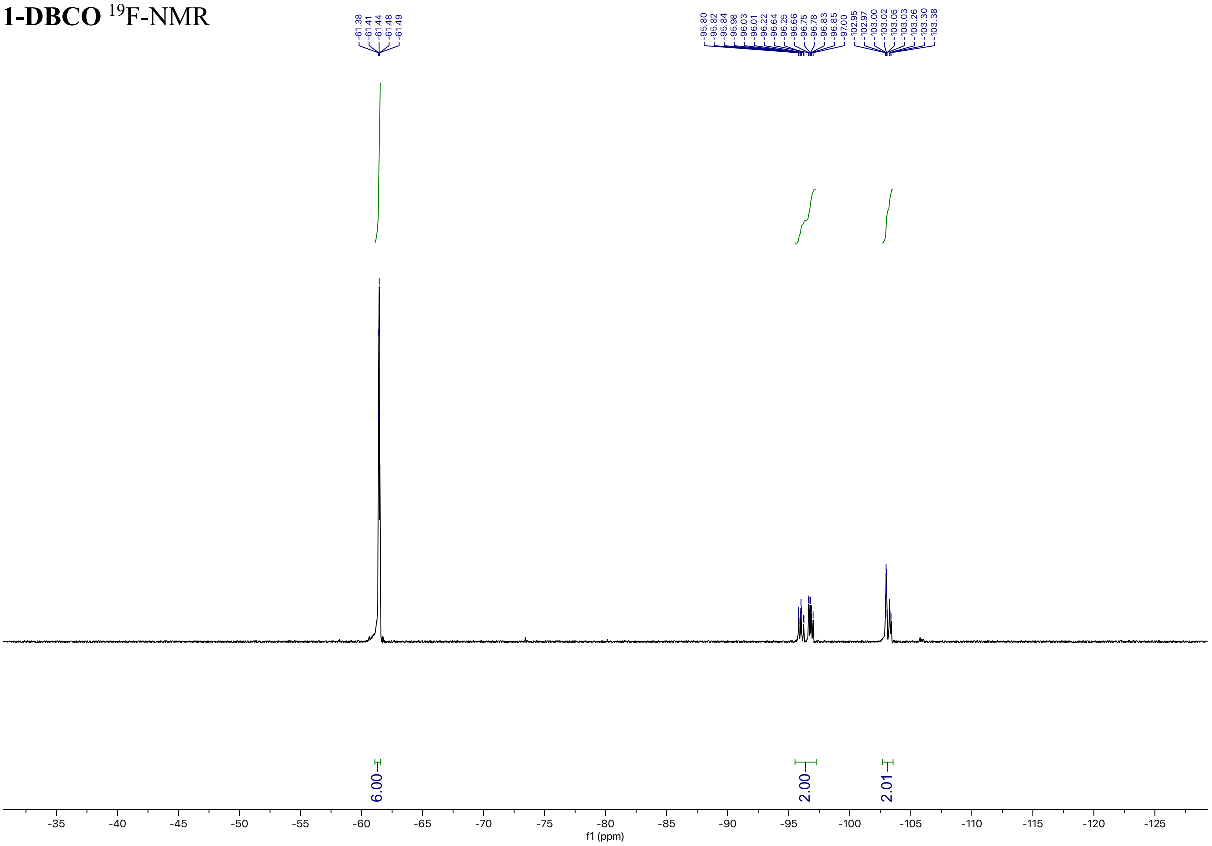

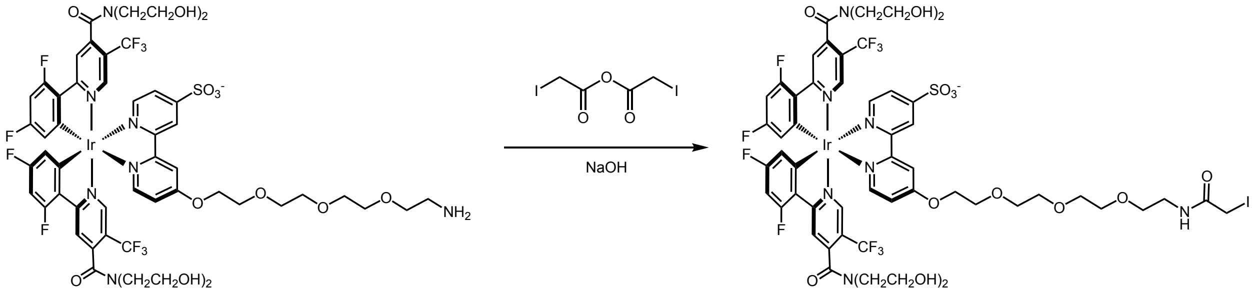

**1-IAA**

A scintillation vial equipped with a PTFE-coated stir bar was charged with **1-NH2** (0.22 g, 0.16 mmol, 1.0 equiv.) and dissolved in 1,4-dioxane:$H_2O$ (3:1, 2.5 mL). The solution was stirred at room temperature for 5 minutes then cooled to 0 °C and NaOH (1 M solution, 0.25 mL, 0.1 M final concentration) was added. Separately, iodoacetic anhydride (0.12 g, 0.35 mmol, 2.2 equiv.) was dissolved in dichloroethane (0.2 mL). The iodoacetic anhydride solution was added dropwise to the **1-NH2** solution at 0 °C with vigorous stirring in 2 additions of 200 µL. The reaction was stirred at 0 °C for 10 minutes in the dark then warmed to room temperature for 50 minutes. Solvent was removed and the residue was redissolved in DMSO:$H_2O$ (9:1, 1 mL). The mixture was purified by preparative HPLC using a 30% solvent B 10-minute isocratic method. The product eluted at 6.0 minutes and was concentrated to afford the title compound as a yellow solid (56.3 mg, 23%).

**Note**: The iodoacetamide containing product is light and temperature sensitive, and should be stored at < -20 °C in the dark.

**$^1$H NMR** (598 MHz, $CD_3OD$_SPE) δ 9.06 – 8.91 (m, 1H), 8.60 (ddd, *J* = 18.2, 8.0, 5.2 Hz, 1H), 8.51 (tt, *J* = 11.0, 2.7 Hz, 1H), 8.45 – 8.34 (m, 1H), 8.26 – 8.06 (m, 2H), 8.05 – 7.64 (m, 5H), 7.30 (ddq, *J* = 10.8, 6.1, 2.7 Hz, 1H), 6.84 – 6.70 (m, 2H), 6.15 (dddd, *J* = 31.9, 24.7, 7.6, 2.7 Hz, 1H), 5.78 – 5.54 (m, 1H), 4.51 (td, *J* = 7.2, 3.3 Hz, 2H), 3.99 – 3.46 (m, 28H), 3.36 (ddd, *J* = 13.6, 6.0, 3.0 Hz, 2H).
**$^{19}$F NMR** (563 MHz, $CD_3OD$_SPE) δ -63.76 – -64.92 (m, 6F), -97.41 – -98.03 (m, 2F), -103.19 – -104.31 (m, 2F).
**LC-MS** $R_t$ = 4.77 min; m/z calcd. For $C_{54}H_{53}IF_{10}IrN_7O_{14}S$ ($[M+H]^+$) = 1566.2, found 1566.3

**1-IA** $^{1}$H-NMR

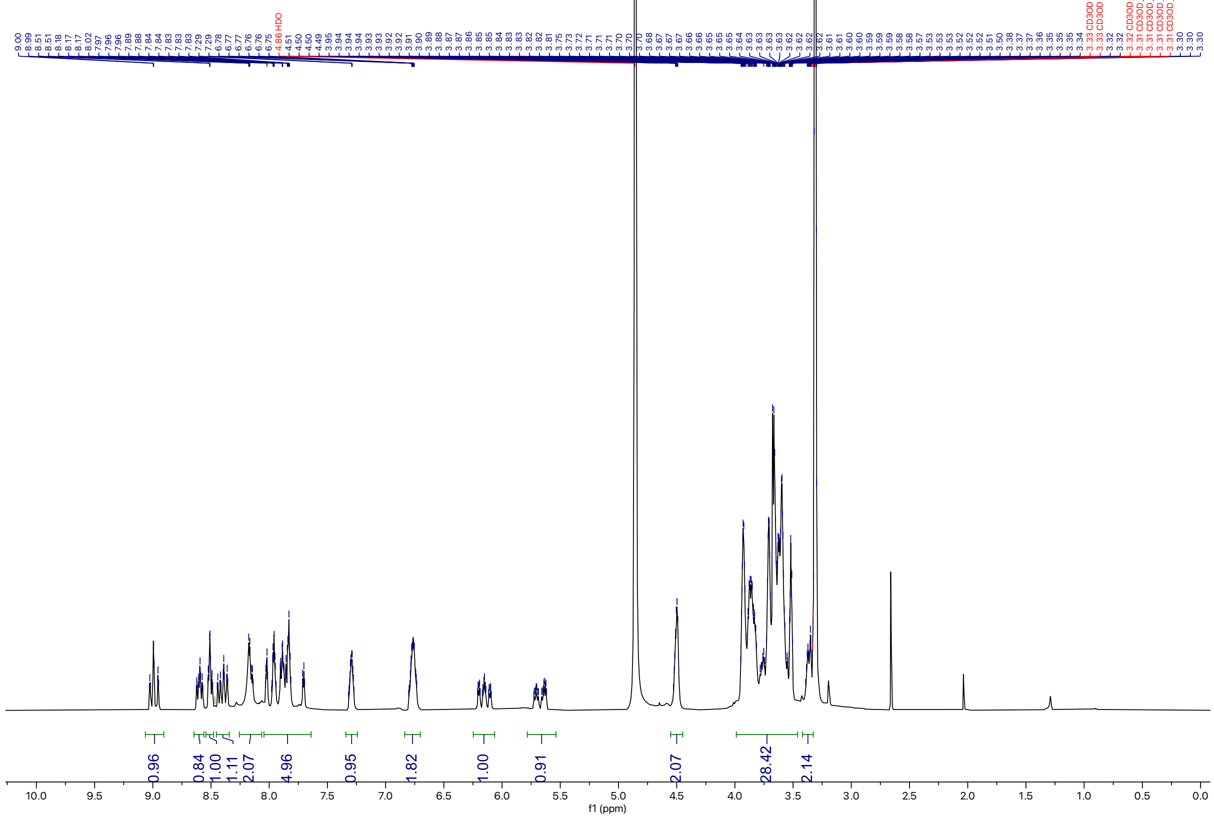

**1-IA** $^{19}$F-NMR

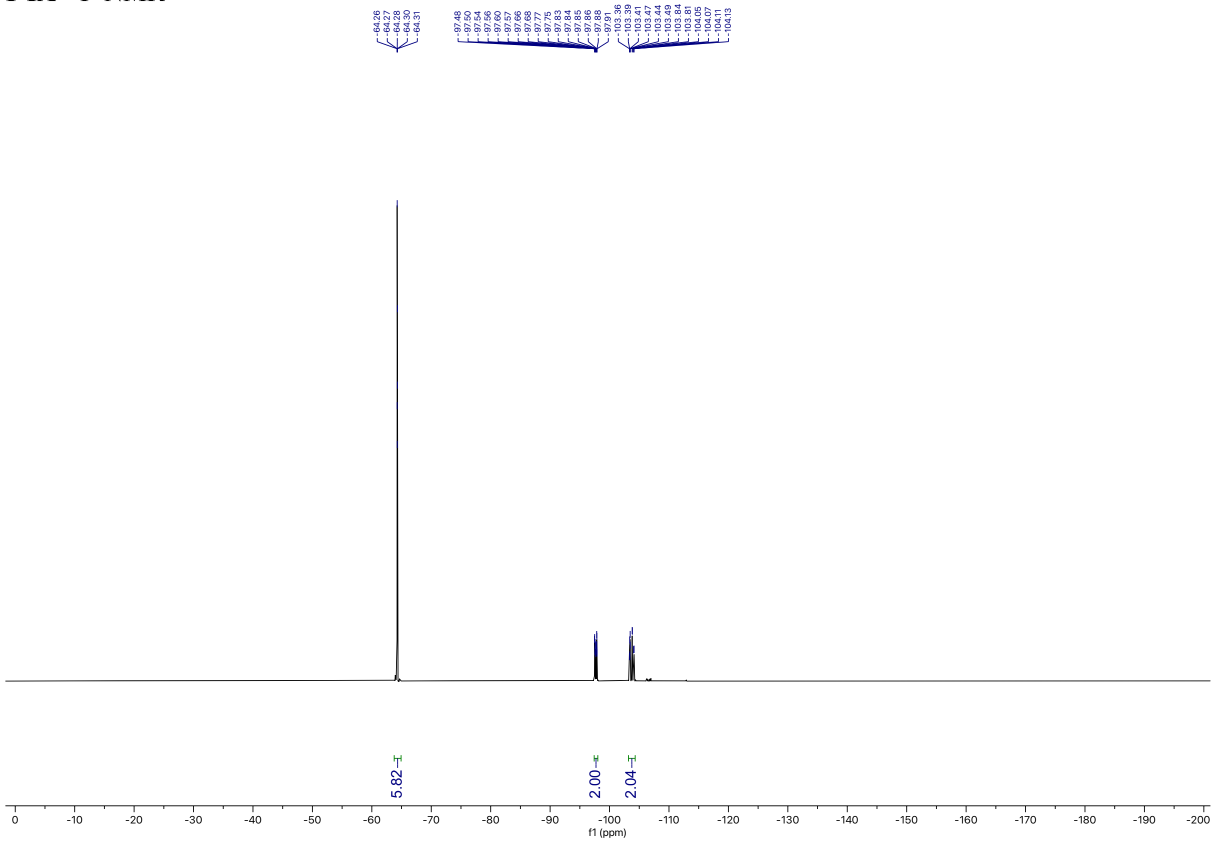

**Small molecule photocatalyst conjugates**

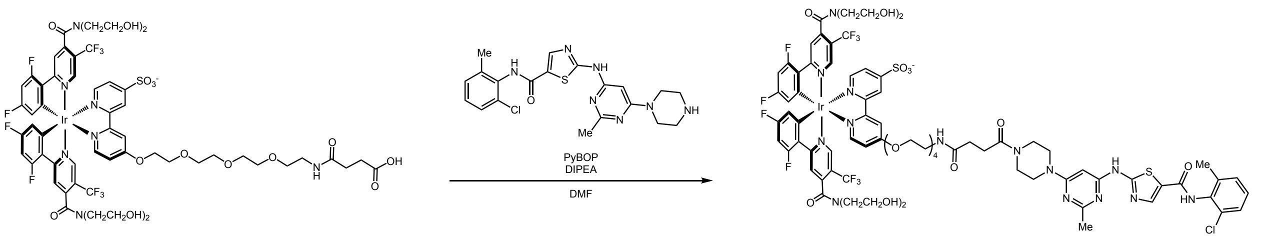

**1-Dasatinib**

A scintillation vial equipped with a PTFE-coated stir bar was charged with **1-$CO_2H$** (4.25 mg, 0.003 mmol, 1.0 equiv.), deshydroxyethyl-dasatinib (382 uL of a 0.01 M solution in DMF, 1.69 mg, 0.0039 mmol, 1.1 equiv.), PyBOP (5.4 mg, 0.01 mmol, 3.0 equiv.), and DMF (0.2 mL). The solution was stirred at room temperature for 5 minutes, then DIPEA (4.83 uL, 0.028 mmol, 8 equiv.) was added and the reaction stirred for 16 hours. The reaction mixture was diluted with MeCN/$H_2O$ (1:1, 0.2 mL) and purified by semi-preparative HPLC using a 5%-60% solvent B 10 minute method. The product eluted at 4.8 minutes and was concentrated to afford the title compound as a yellow solid (1.7 mg, 30%).

**$^1$H NMR** (500 MHz, DMSO) δ 11.51 (s, 1H), 9.88 (s, 1H), 9.07 (s, 1H), 8.60 (s, 1H), 8.39 (d, $J$ = 32.7 Hz, 2H), 8.22 (s, 1H), 8.03 (s, 1H), 7.87 (t, $J$ = 5.7 Hz, 1H), 7.70 (s, 1H), 7.63 (s, 1H), 7.55 (s, 1H), 7.40 (dd, $J$ = 7.7, 1.8 Hz, 1H), 7.33 – 7.20 (m, 3H), 7.07 (ddt, $J$ = 12.1, 9.4, 2.5 Hz, 2H), 6.07 (s, 1H), 5.77 (d, $J$ = 103.5, 42.3 Hz, 3H), 4.90 (d, $J$ = 86.1 Hz, 3H), 4.63 – 4.39 (m, 2H), 3.95 – 3.77 (m, 3H), 3.66 – 3.48 (m, 22H), 3.37 (d, $J$ = 6.0 Hz, 5H), 3.17 (d, $J$ = 5.8 Hz, 3H), 2.55 (t, $J$ = 7.2 Hz, 2H), 2.42 (s, 3H), 2.38 – 2.29 (m, 2H), 2.24 (s, 3H).
**$^{19}$F NMR** (471 MHz, DMSO) δ -59.68 (s, 6F), -102.70 (t, 2F), -106.09 (q, $J$ = 69.6 Hz, 2F).
**LC-MS** $R_t$ = 5.39 min; m/z calcd. For $C_{76}H_{76}ClF_{10}IrN_{14}O_{16}S_2$ ($[M+H]^+$) = 1923.4, found 1923.7

**1-dasatinib** $^{1}$H-NMR

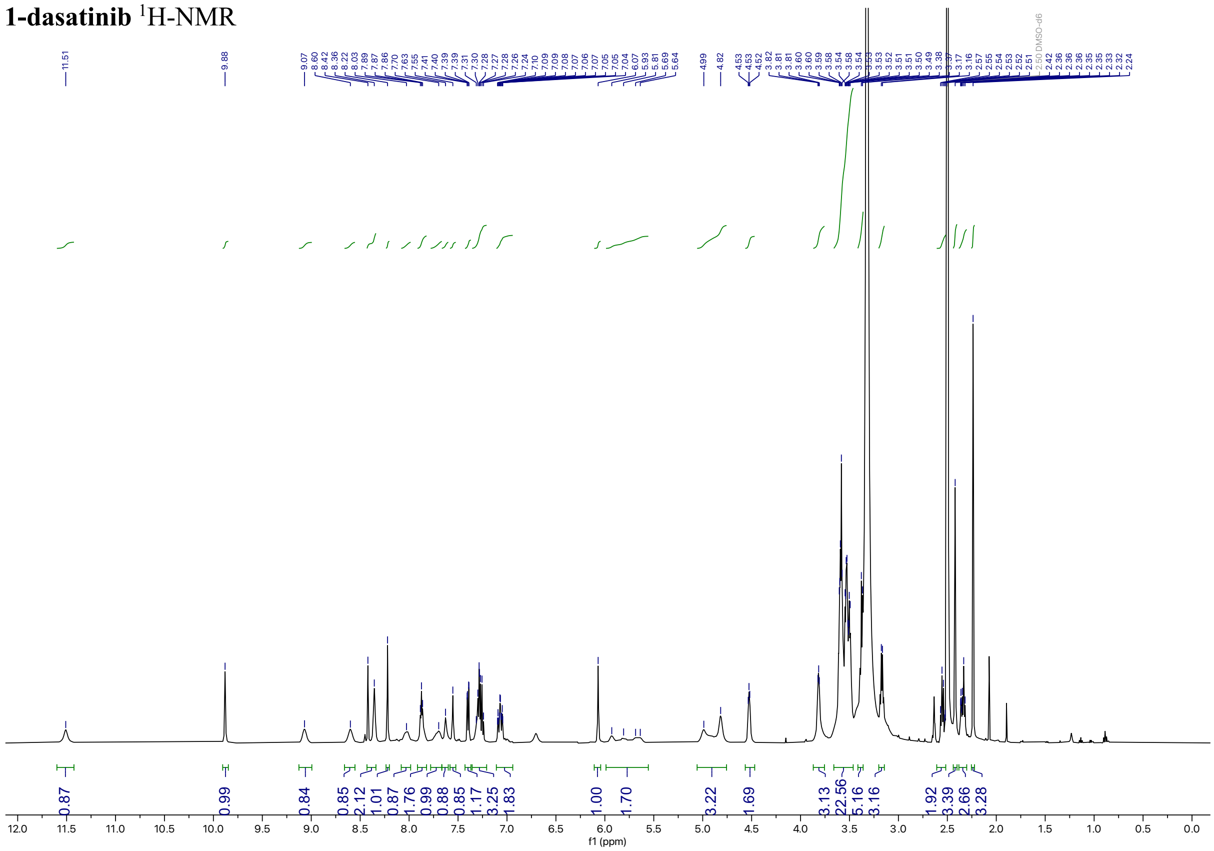

**1-dasatinib** $^{19}$F-NMR

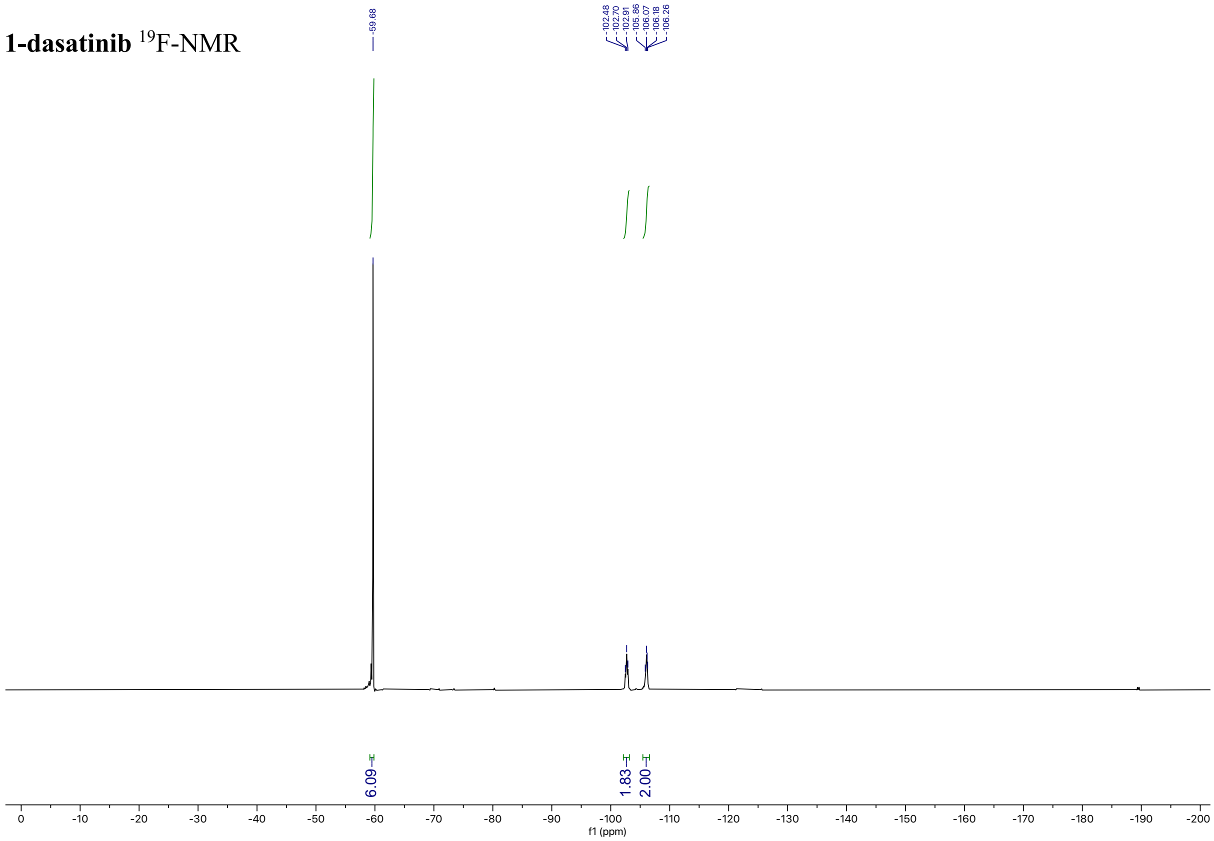

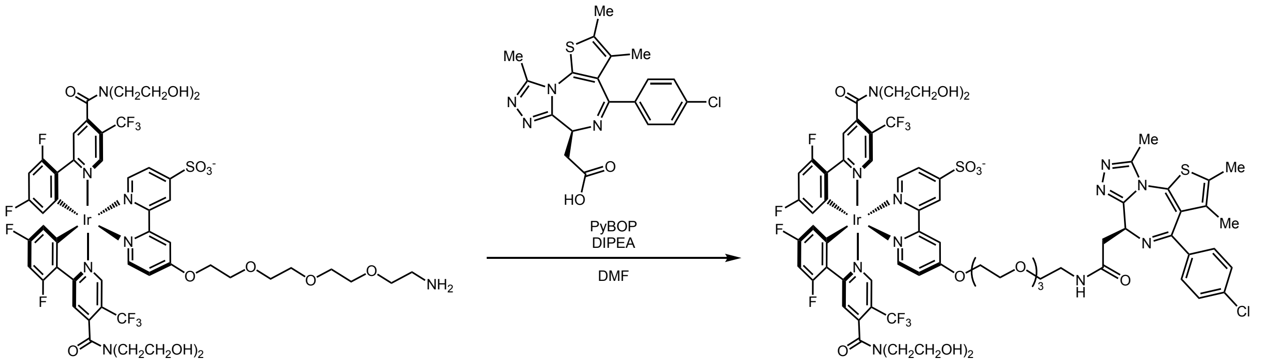

**1-(+)-JQ1**

A scintillation vial equipped with a PTFE-coated stir bar was charged with **1-NH₂** (0.04 g, 0.029 mmol, 1.0 equiv.), (+)-JQ1-COOH (0.012 g, 0.031 mmol, 1.1 equiv.), PyBOP (0.045 g, 0.086 mmol, 3.0 equiv.), and DMF (0.6 mL). The solution was stirred at room temperature for 5 minutes, then DIPEA (0.04 mL, 0.23 mmol, 8 equiv.) was added and the reaction stirred for 16 hours. The reaction mixture was diluted with $H_2O$ (0.2 mL) and purified by semi-preparative HPLC using a 35% solvent B 25-minute isocratic method. The product eluted at 5.3 minutes and solvent was removed to afford the title compound as a yellow solid (3 mg, 6%).

**$^1$H NMR** (500 MHz, $CD_3OD$_SPE) δ 8.97 (d, *J* = 5.8 Hz, 1H), 8.50 (s, 2H), 8.40 (dd, *J* = 5.8, 2.6 Hz, 1H), 8.18 (s, 1H), 7.86 (d, *J* = 97.1 Hz, 4H), 7.42 (ddd, *J* = 33.5, 8.6, 3.2 Hz, 4H), 7.30 (d, *J* = 17.5 Hz, 1H), 6.79 (q, *J* = 10.2 Hz, 2H), 5.92 (d, *J* = 101.5 Hz, 2H), 4.58 (s, 2H), 4.56 – 4.45 (m, 2H), 3.97 – 3.37 (m, 30H), 2.66 (s, 3H), 2.44 (s, 3H), 1.69 (s, 3H).
**$^{19}$F NMR** (563 MHz, $CD_3OD$_SPE) δ -61.75 (d, *J* = 11.6 Hz, 6F), -103.57 (d, *J* = 153.1 Hz, 2F), -107.26 (s, 2F).
**LC-MS** $R_t$ = 5.46 min; m/z calcd. For $C_{71}H_{67}ClF_{10}IrN_{11}O_{14}S_2$ ($[M+H]^+$) = 1780.3, found 1780.7

**1-(+)-JQ1** $^{1}$H-NMR

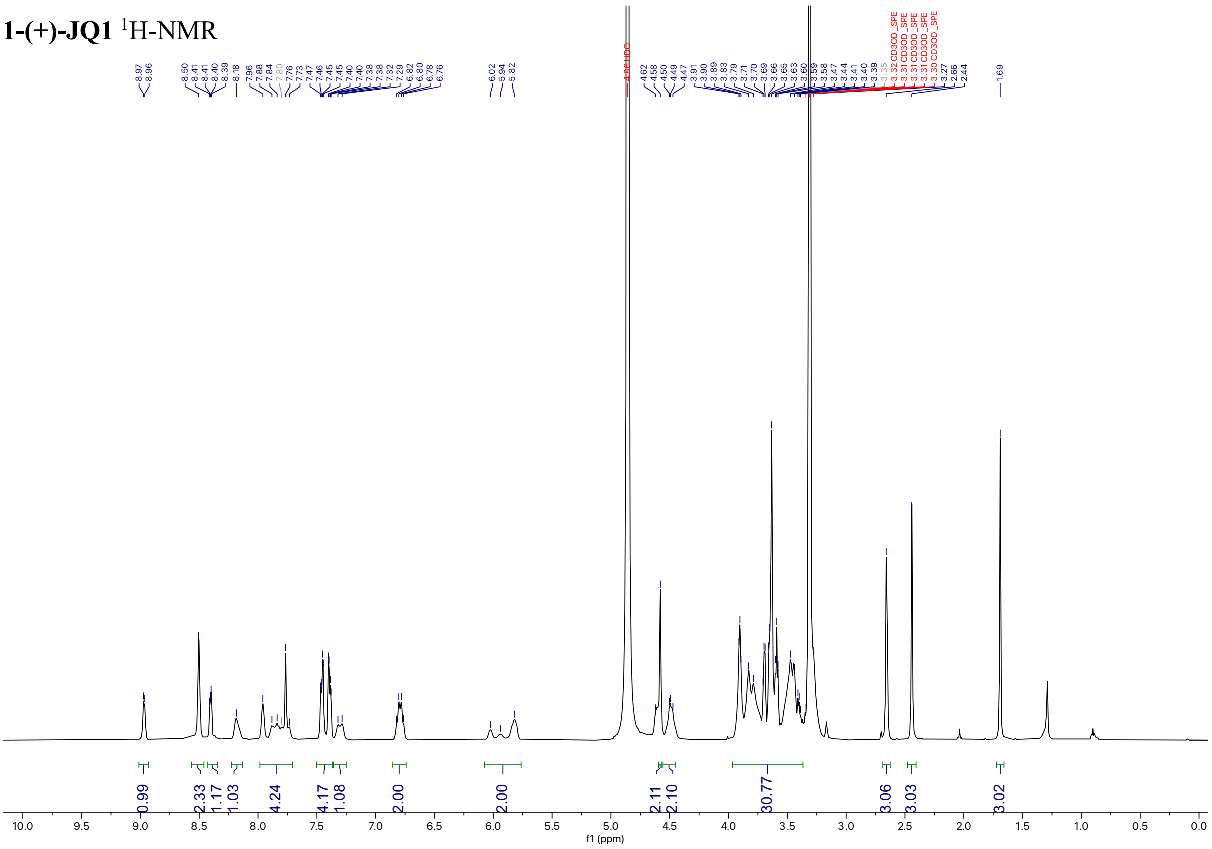

**1-(+)-JQ1** $^{19}$F-NMR

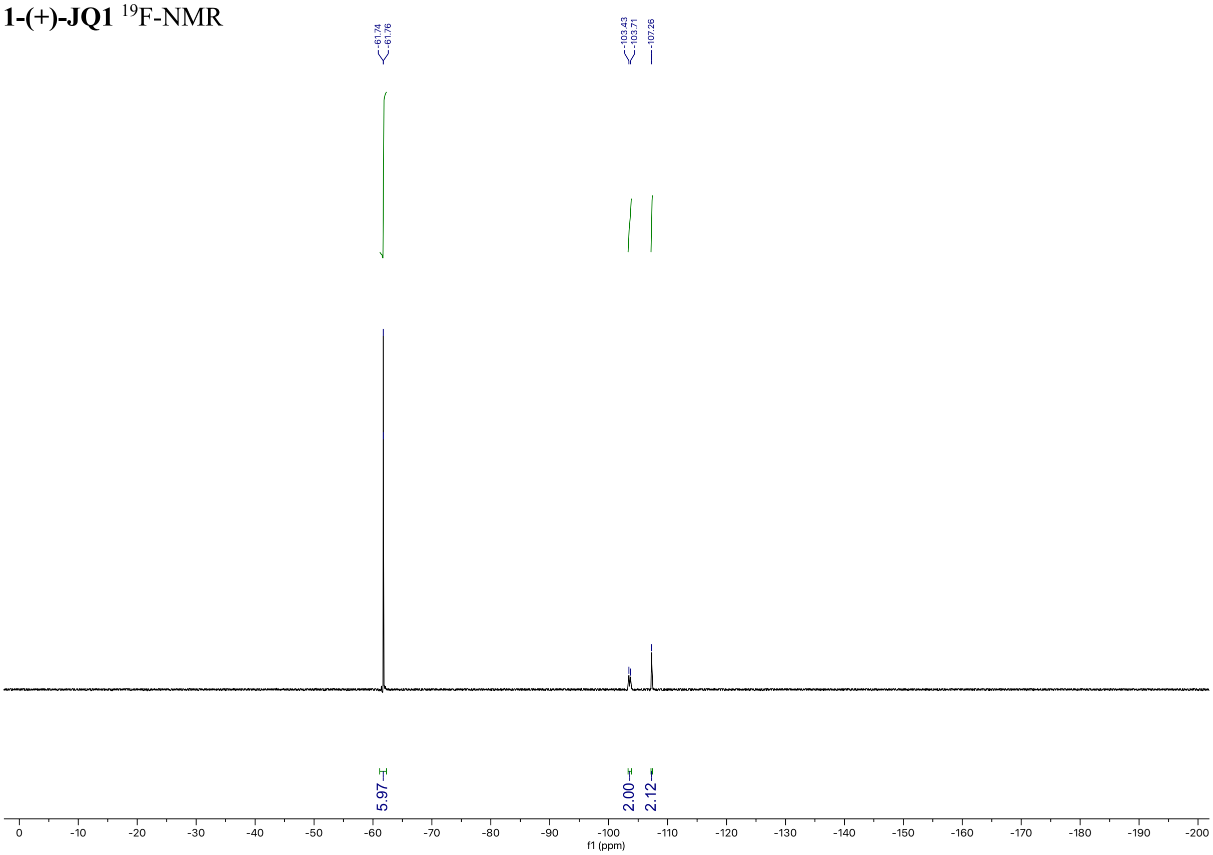

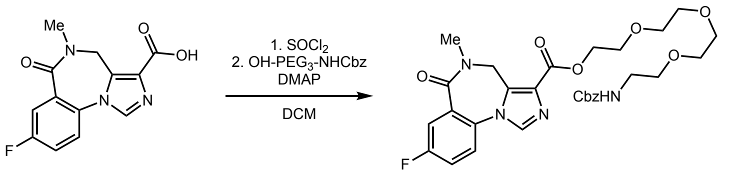

**Flumazenil-Ester-PEG3-NHCbz**
A scintillation vial equipped with a PTFE-coated stir bar was charged with Flumazenil acid (0.013 g, 0.046 mmol, 1.0 equiv.). The vial was placed under an argon atmosphere and thionyl chloride (2.356 mL, 323 mmol, 7,000 equiv.) was added and the mixture refluxed at 65 °C and stirred for 1 hour. The reaction was cooled to room temperature, solvent was removed under a stream of nitrogen, and the residue redissolved in DCM (1 mL). To the reaction added OH-PEG3-NHbz (0.049 g, 0.150 mmol, 3.25 equiv.) and DMAP (0.045 g, 0.369 mmol, 8.0 equiv.) dissolved in DCM (1 mL) and stirred at room temperature for 5 hours. Solvent was removed and the crude mixture was purified by flash column chromatography on silica (EtOAc:MeOH = 0% to 5% over 10 CVs) to afford the title compound as a cloudy oil (0.015 g, 57%).

**$^{1}$H NMR** (500 MHz, $CDCl_3$) δ 7.85 (s, 1H), 7.77 (dd, *J* = 8.7, 2.9 Hz, 1H), 7.41 (dd, *J* = 8.8, 4.5 Hz, 1H), 7.35 – 7.32 (m, 5H), 7.30 – 7.27 (m, 1H), 5.08 (s, 1H), 4.49 (d, *J* = 26.1 Hz, 2H), 3.81 (t, *J* = 4.8 Hz, 2H), 3.69 – 3.57 (m, 10H), 3.54 (t, *J* = 5.1 Hz, 2H), 3.39 – 3.35 (m, 2H), 3.23 (s, 3H).
**$^{19}$F NMR** (471 MHz, $CDCl_3$) δ -111.12.
**LC-MS** $R_t$ = 3.89 min (40%-95% B over 10 minutes); m/z calcd. For $C_{29}H_{33}FN_4O_8$ ($[M+H]^+$) = 585.61, found 585.54

**Flumazenil-Ester-PEG3-NHCbz** $^{1}$H-NMR

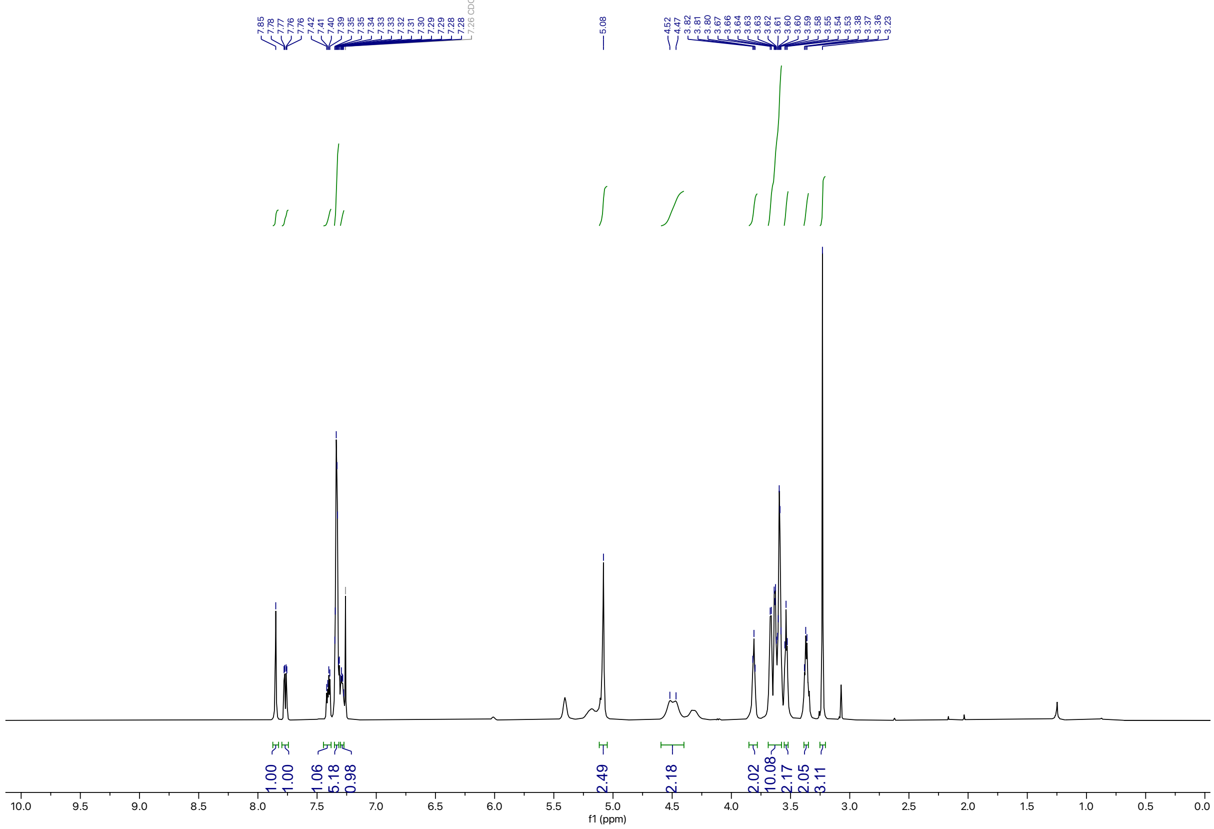

**Flumazenil-Ester-PEG3-NHCbz** $^{19}$F-NMR

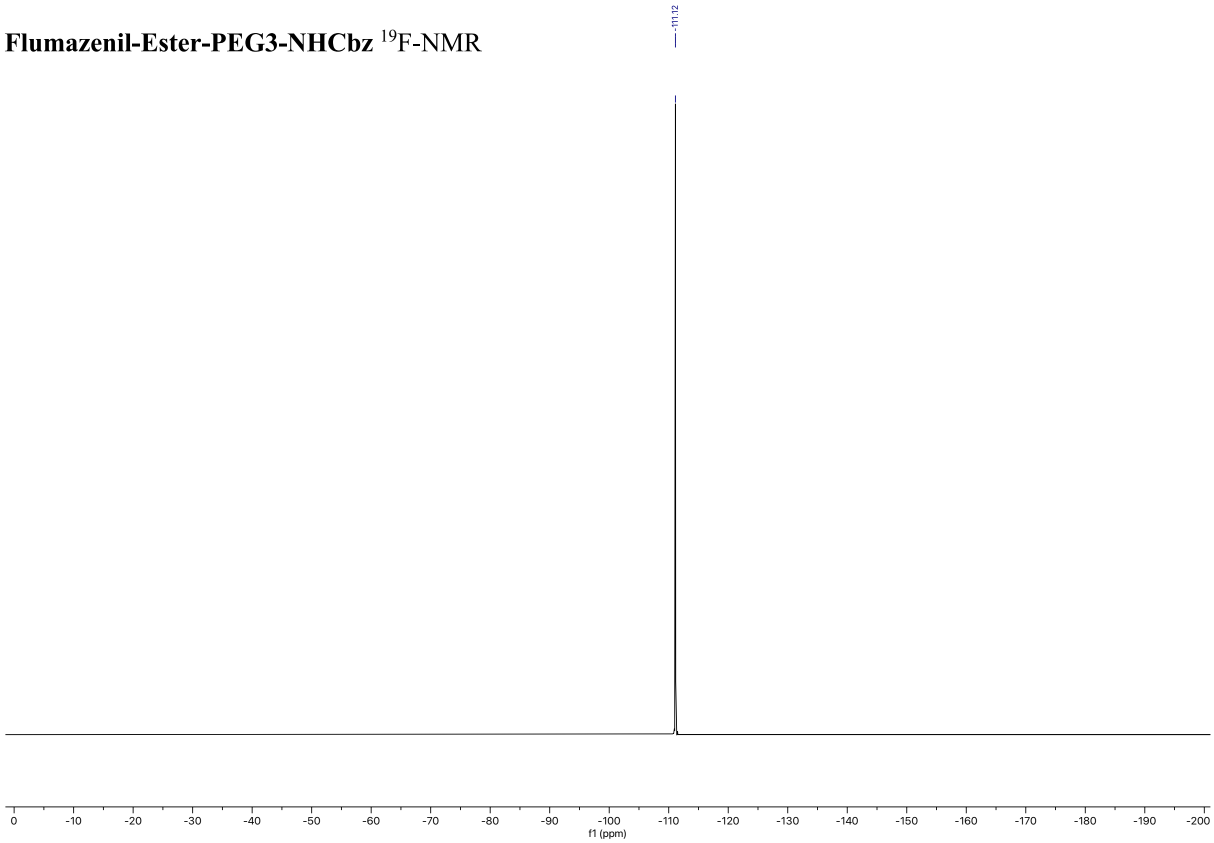

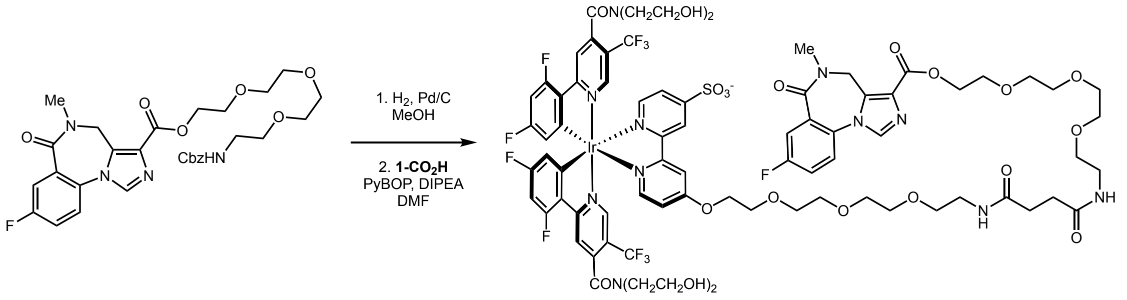

**1-Flumazenil**

A scintillation vial equipped with a PTFE-coated stir bar was charged with **Flumazenil-Ester-PEG3-NHCbz** (0.015 g, 0.026 mmol, 1.0 equiv.) and 10% Pd/C, type 487 (0.279 mg) and suspended in MeOH (1 mL). The mixture was placed under $H_2$ atmosphere and stirred at room temperature for 3 hours. The mixture was filtered over celite and the filter cake was washed with MeOH (4 mL). Solvent was removed to afford the title compound which was carried forward without additional purification.

A scintillation vial equipped with a PTFE-coated stir bar was charged with **1-$CO_2H$** (0.035 g, 0.023 mmol, 1.0 equiv.), crude **Flumazenil-Ester-PEG3-$NH_2$** synthesized above (0.012 g, 0.026 mmol, 1.1 equiv.), PyBOP (0.036 g, 0.070 mmol, 3.0 equiv.), and DMF (0.468 mL). The solution was stirred at room temperature for 5 minutes, then DIPEA (0.032 mL, 0.187 mmol, 8 equiv.) was added and the reaction stirred at room temperature for 16 hours. The reaction mixture was diluted with MeOH (0.12 mL) and purified by semi-preparative HPLC using a 20%-65% solvent B gradient over 10 minutes. The product eluted at 4.63 minutes and solvent was removed to afford the title compound as a yellow solid (8.5 mg, 19%).

**$^{1}$H NMR** (500 MHz, DMSO) δ 9.07 (s, 1H), 8.59 (s, 1H), 8.40 – 8.24 (m, 3H), 8.03 (s, 1H), 7.91 – 7.78 (m, 4H), 7.75 – 7.61 (m, 4H), 7.56 (s, 1H), 7.30 (s, 1H), 7.06 (ddd, $J$ = 12.4, 9.5, 2.7 Hz, 2H), 6.06 – 5.55 (m, 2H), 4.90 (br,m, $J$ = 93.5 Hz, 4H), 4.57 – 4.47 (m, 2H), 4.37 (s, 2H), 3.88 – 3.69 (m, 6H), 3.62 – 3.00 (m, 48H), 2.27 (s, 3H).
**$^{19}$F NMR** (471 MHz, DMSO) δ -59.68 (s, 6F), -102.33 – -103.08 (m, 2F), -106.09 (q, $J$ = 65.5 Hz, 2F), -113.37 (s, 1F).
**LC-MS** $R_t$ = 3.99 min (25%-95% B over 10 minutes); m/z calcd. For $C_{77}H_{81}F_{11}IrN_{11}O_{21}S$ ($[M+H]^+$) = 1930.49, found 1930.22

**1-Flumazenil** $^{1}$H-NMR

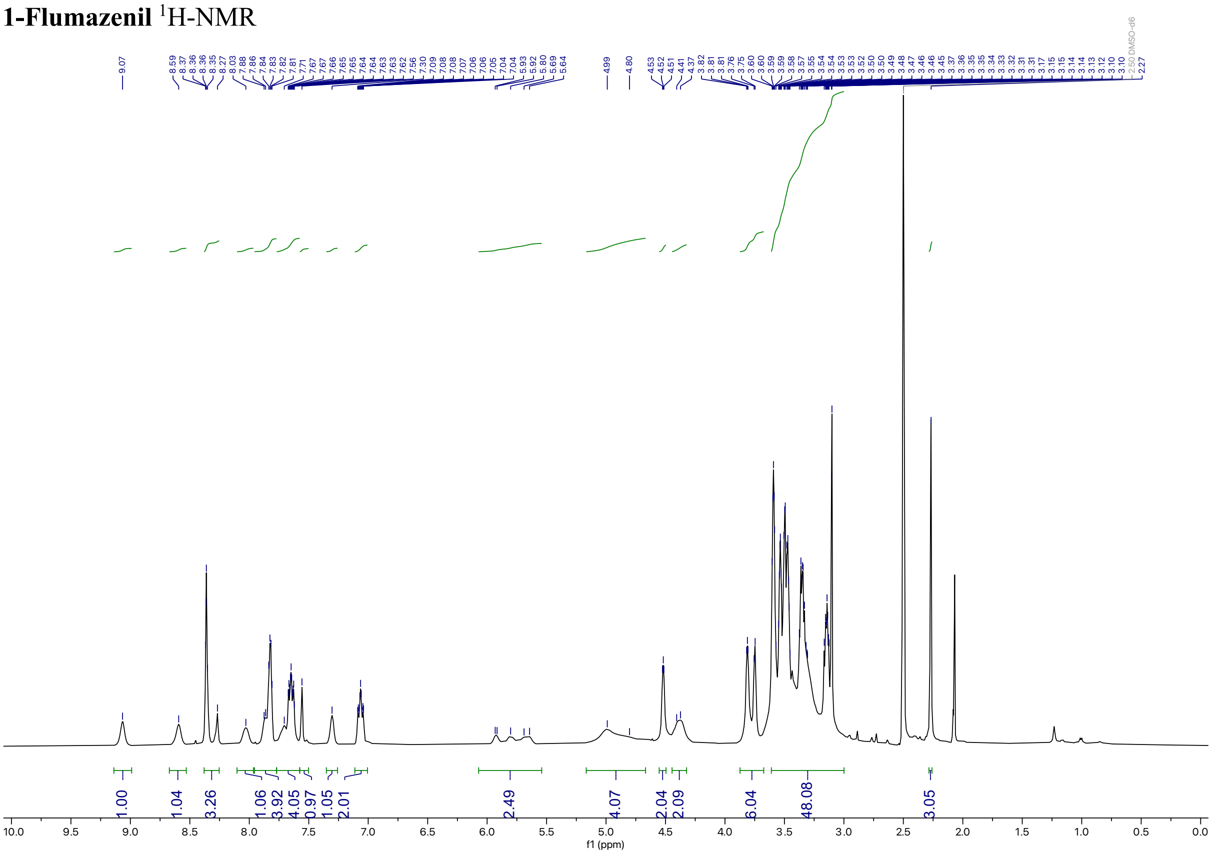

**1-Flumazenil** $^{19}$F-NMR

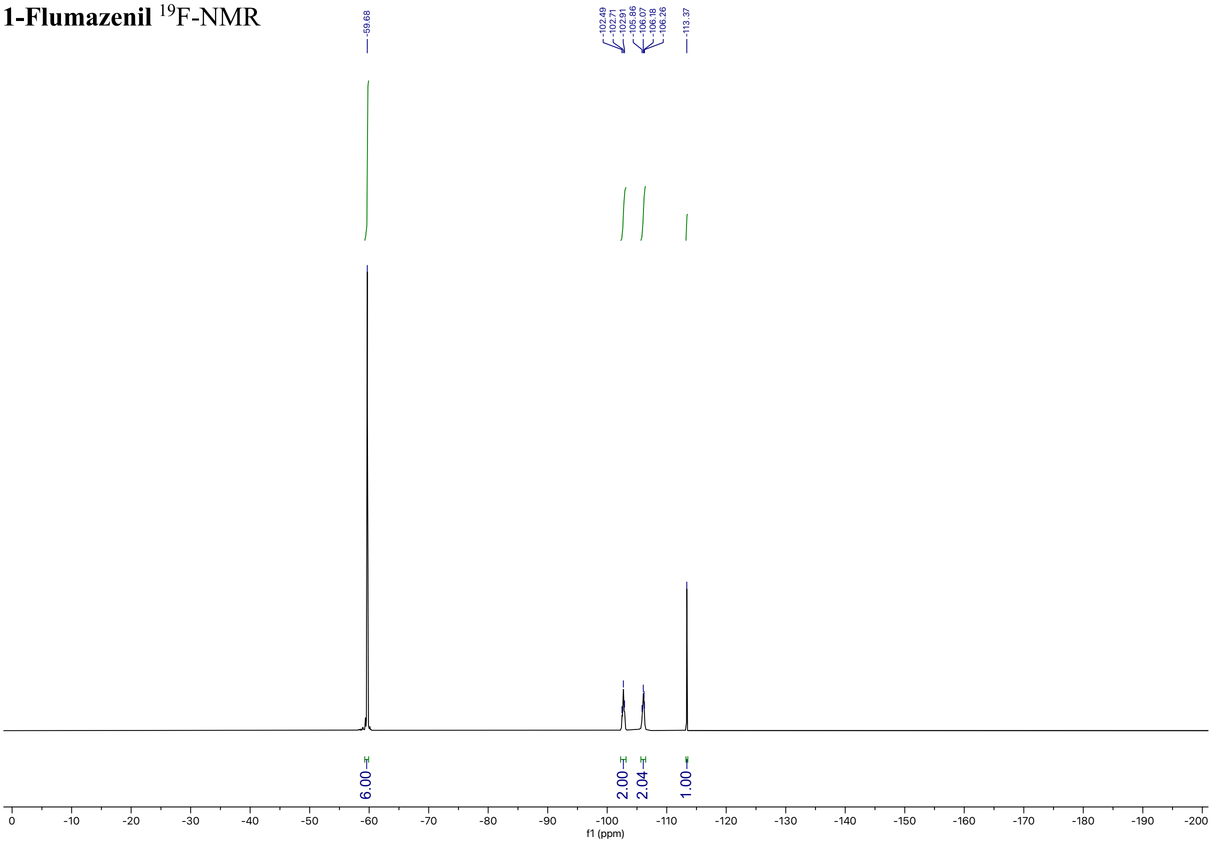

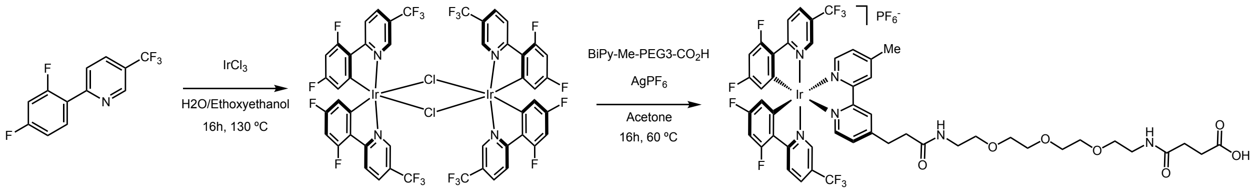

**2-$CO_2$H**

A flame-dried schlenk flask equipped with a PTFE-coated stir bar was charged with 2-(2,4-difluorophenyl)-5-(trifluoromethyl)pyridine (2 g, 7.72 mmol, 2.1 equiv.) and iridium(III) chloride hydrate (1.095 g, 3.677 mmol, 1.0 equiv.). The flask was placed under an argon atmosphere and 2-ethoxyethanol:$H_2O$ (2:1, 74 mL) was added. The mixture was heated to 130 °C and stirred for 16 hours. The reaction was cooled to room temperature, and quenched with cold water (400 mL) to afford a precipitate. The precipitate was filtered and washed with excess cold water, then dried to afford the title compound as a bright orange powder which was carried forward without further purification. (2.590 g, 47%).

A scintillation vial equipped with a PTFE-coated stir bar was charged with the dimeric intermediate (0.055 g, 0.037 mmol, 1.0 equiv.), **bipy-Me-PEG$_3$-$CO_2$H** (0.038 g, 0.074 mmol, 2 equiv.), and silver hexafluorophosphate (0.028 g, 0.111 mmol, 3.0 equiv.). Acetone (0.74 mL) was added, and the mixture was heated to 60 °C and stirred for 16 hours. The reaction was cooled to room temperature, filtered to remove insoluble salts, and the solvent was removed. The crude mixture was dissolved in DMSO (1.2 mL) and purified by preparative HPLC using a 35%-95% solvent B gradient over 15 minutes with 75 µL MeCN sample sandwiching during injection. Solvent was removed to afford the title compound as a yellow solid (25 mg, 28%).

**$^1$H NMR** (598 MHz, $CDCl_3$) δ 8.65 (d, *J* = 20.0 Hz, 2H), 8.48 (ddd, *J* = 18.5, 8.7, 3.0 Hz, 2H), 8.07 – 8.01 (m, 2H), 7.77 (dd, *J* = 25.2, 5.6 Hz, 2H), 7.60 (s, 1H), 7.51 (d, *J* = 6.2 Hz, 2H), 7.35 (d, *J* = 5.6 Hz, 1H), 7.07 (br, s, *J* = 10.5 Hz, 1H), 6.93 (br, s, *J* = 7.3 Hz, 1H), 6.64 (ddt, *J* = 12.2, 9.3, 3.1 Hz, 2H), 5.63 (dd, *J* = 6.8, 4.0 Hz, 2H), 3.68 – 3.59 (m, 9H), 3.54 (dd, *J* = 9.3, 4.9 Hz, 4H), 3.40 (dt, *J* = 20.2, 5.7 Hz, 4H), 3.22 – 3.14 (m, 2H), 2.75 (td, *J* = 7.5, 4.5 Hz, 2H), 2.67 (s, 3H), 2.61 – 2.50 (m, 4H).

**$^{13}$C NMR** (150 MHz, $CDCl_3$) δ 174.34, 173.90, 168.22, 168.06, 165.90, 164.24, 164.16, 162.44, 161.93, 155.48, 155.32, 154.97, 154.75, 154.64, 154.59, 150.00, 149.33, 145.07, 136.67, 129.94, 129.76, 127.22, 126.48, 125.94, 124.02, 123.88, 123.83, 123.69, 122.66, 122.59, 120.85, 120.78, 114.33, 114.28, 114.21, 114.16, 100.22, 100.11, 77.37, 77.16, 76.95, 70.66, 70.51, 70.31, 69.37, 39.84, 39.60, 35.54, 31.44, 30.89, 30.82, 28.30, 21.56, 21.41, 0.12.

**$^{19}$F NMR** (563 MHz, $CDCl_3$) δ -62.77 (s, 3F), -62.84 (s, 3F), -71.85 (d, *J* = 713.4 Hz, 5F), -101.40 (dd, *J* = 68.5, 12.8 Hz, 2F), -105.84 (dd, *J* = 78.8, 12.5 Hz, 2F).

**LC-MS** $R_t$ = 3.63 min (70%-95% B over 10 minutes); m/z calcd. For $C_{50}H_{46}F_{10}IrN_6O_7$ ($[M+H]^+$) = 1226.3, found 1226.8

**2-CO₂H** $^{1}$H-NMR

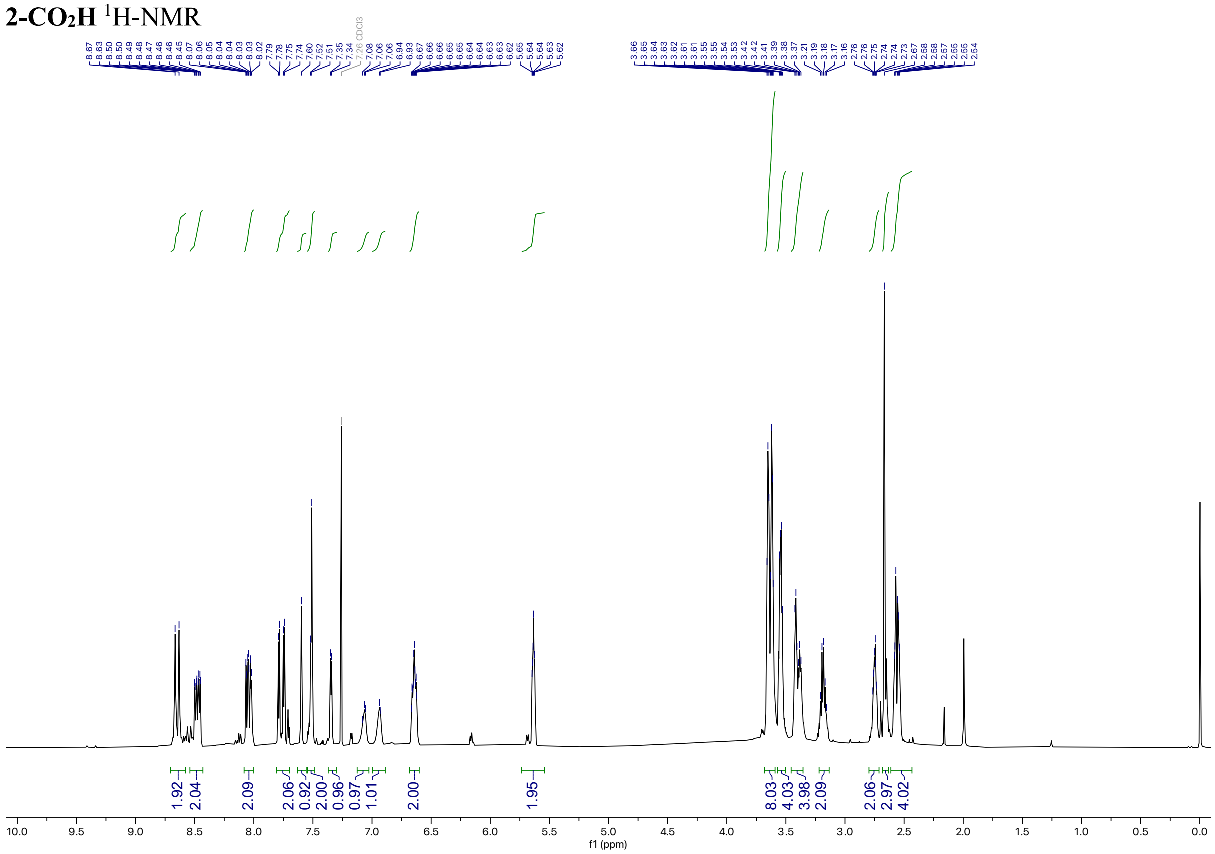

**2-CO₂H** $^{13}$C-NMR

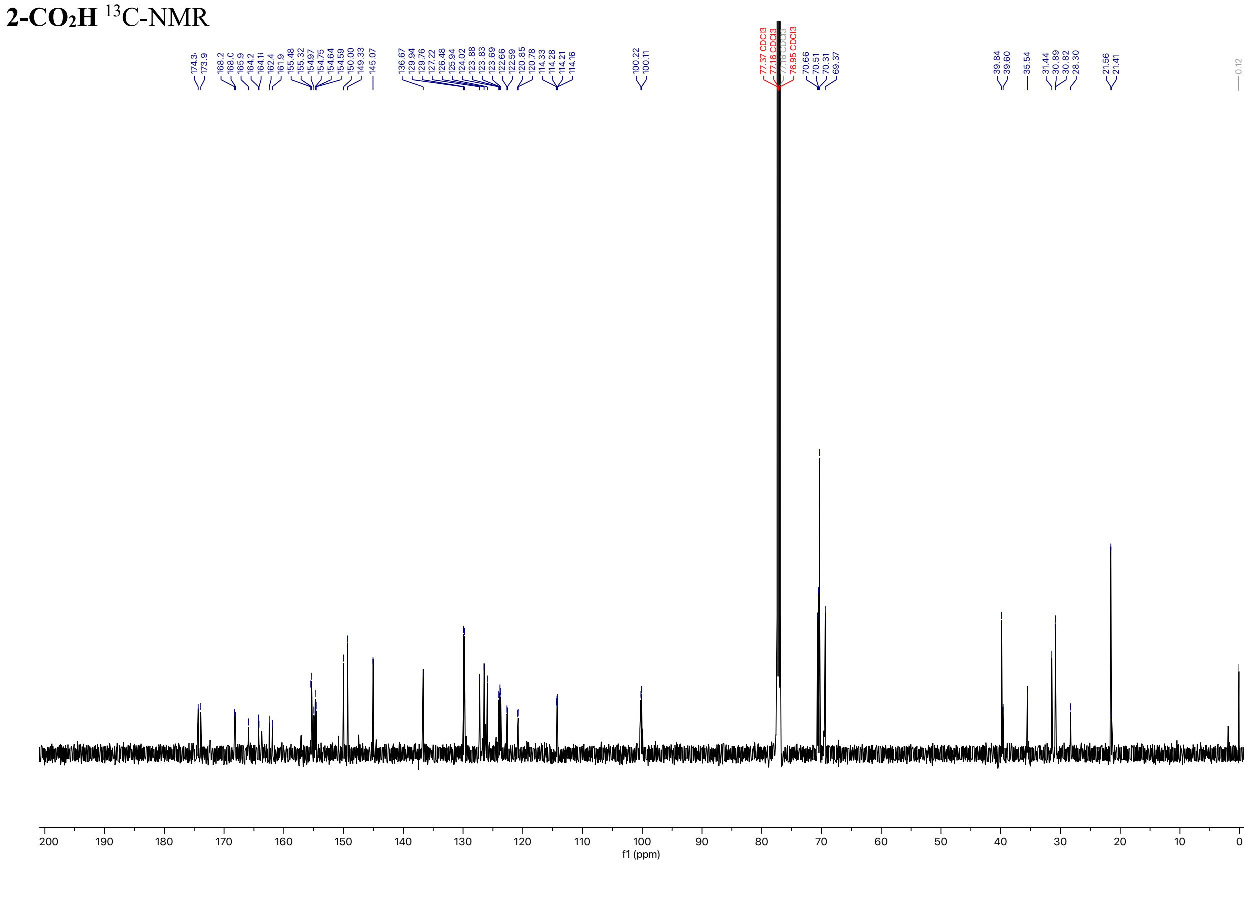

**2-CO₂H** $^{19}$F-NMR

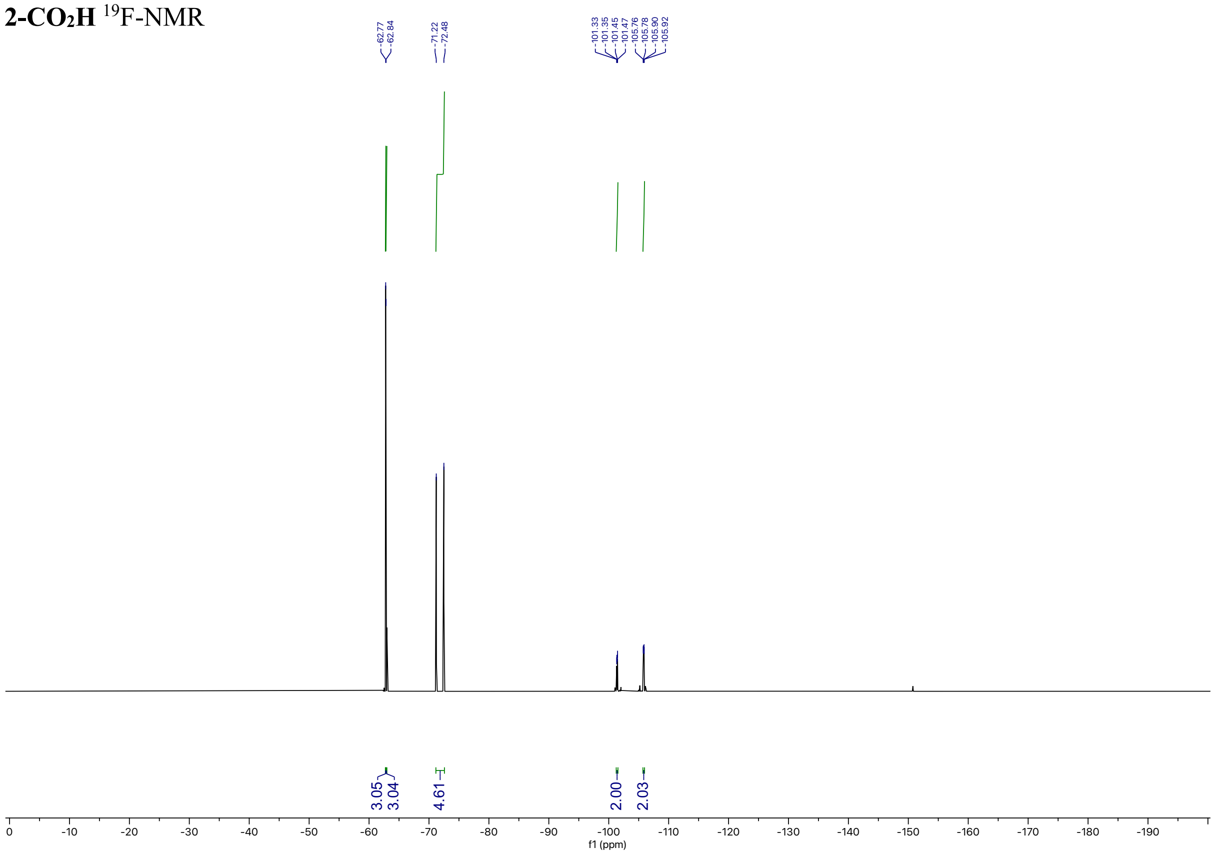

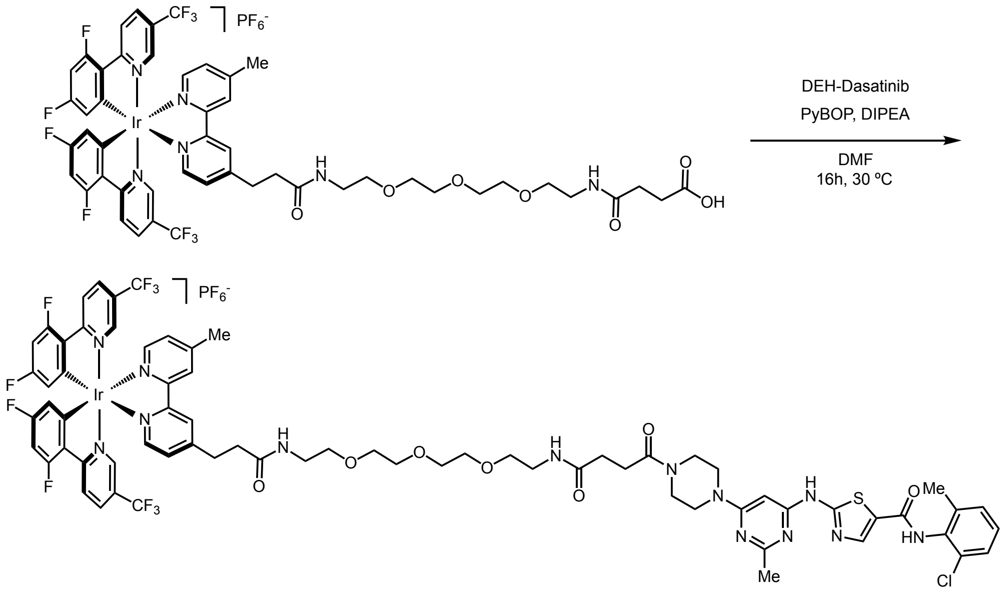

**2-Dasatinib**

A scintillation vial equipped with a PTFE-coated stir bar was charged with **2-CO₂H** (12.6 mg, 0.010 mmol, 1.0 equiv.), des-hydroxyethyl-dasatinib 1.131 mL, 0.01 M solution in DMF, 5.01 mg, 0.011 mmol, 1.1 equiv.), PyBOP (16.1 mg, 0.031 mmol, 3.0 equiv.), and DMF (0.2 mL). The solution was stirred at room temperature for 5 minutes, then DIPEA (14.33 uL, 0.082 mmol, 8 equiv.) was added and the reaction stirred for 16 hours. The reaction mixture was diluted with MeCN/$H_2O$ (1:1, 0.5 mL) and purified by semi-preparative HPLC using a 55% solvent B 10 minute isocratic method. The product eluted at 3.2 minutes, and solvent was removed to afford the title compound as a yellow solid (4.6 mg, 27%).

**$^1$H NMR** (598 MHz, DMSO) δ 11.53 (br, s, 1H), 9.89 (s, 1H), 8.81 (s, 2H), 8.49 – 8.39 (m, 4H), 8.22 (s, 1H), 7.99 (t, *J* = 4.5 Hz, 1H), 7.88 (q, *J* = 5.8 Hz, 1H), 7.82 (dd, *J* = 10.1, 5.7 Hz, 2H), 7.65 (s, 1H), 7.58 (ddd, *J* = 10.4, 5.8, 1.6 Hz, 2H), 7.52 (s, 1H), 7.40 (d, *J* = 7.7 Hz, 1H), 7.27 (dt, *J* = 15.4, 7.6 Hz, 2H), 7.06 (ddd, *J* = 12.2, 9.4, 2.4 Hz, 2H), 6.78 (br, s, 2H), 6.07 (s, 1H), 5.77 (ddd, *J* = 8.6, 6.6, 2.3 Hz, 2H), 3.63 – 3.40 (m, 15H), 3.16 (dd, *J* = 7.6, 4.0 Hz, 3H), 3.06 (t, *J* = 7.6 Hz, 2H), 2.57 (s, 4H), 2.56 – 2.53 (m, 1H), 2.41 (s, 3H), 2.34 (t, *J* = 7.0 Hz, 2H), 2.24 (s, 3H).

**$^{19}$F NMR** (563 MHz, DMSO) δ -61.57 (d, *J* = 56.2 Hz, 6F), -70.15 (d, *J* = 711.2 Hz, 3F), -103.31 (dd, *J* = 22.9, 11.8 Hz, 2F), -106.78 (d, *J* = 12.1 Hz, 2F).

**LC-MS** $R_t$ = 6.15 min; m/z calcd. For $C_{70}H_{66}ClF_{10}IrN_{13}O_7S$ ([M+2H]$^{2+}$) = 826.2, found 826.4

**2-dasatinib** $^{1}$H-NMR

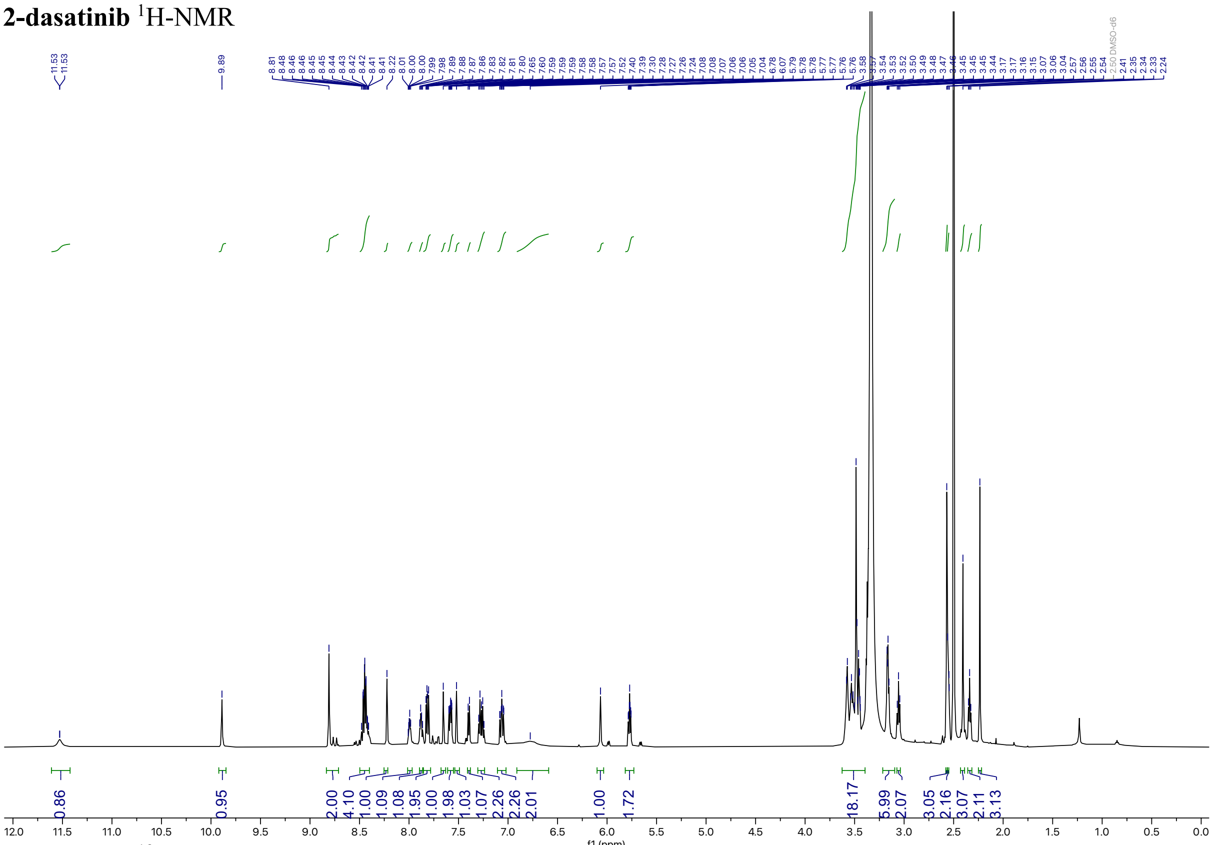

**2-dasatinib** $^{19}$F-NMR

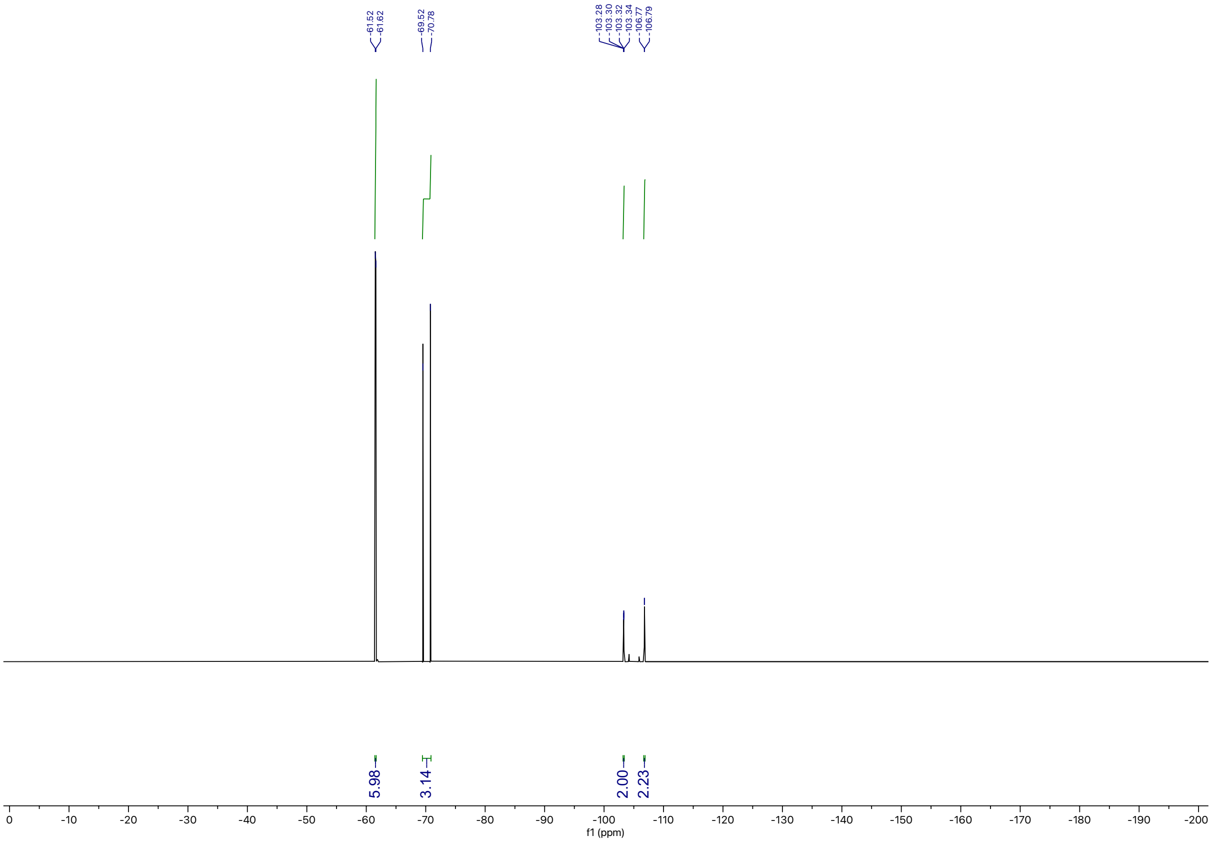

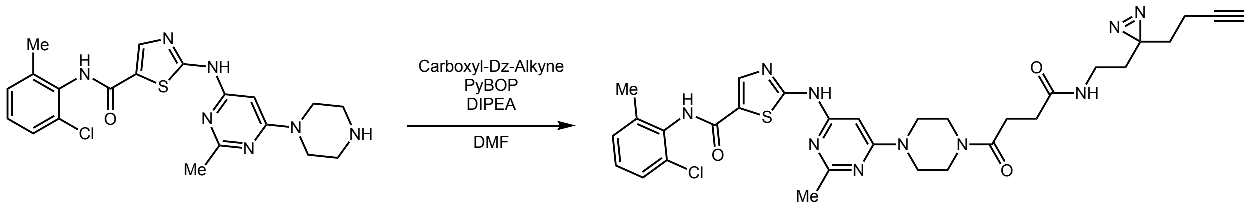

**Dasastinib-dz-alkyne**

A scintillation vial equipped with a PTFE-coated stir bar was charged with carboxyl-diazirine-alkyne (0.012 g, 0.051 mmol, 1.0 equiv.), PyBOP (0.080 g, 0.154 mmol, 3.0 equiv.), and DMF (0.264 mL). The solution was stirred at room temperature for 5 minutes then DIPEA (0.072 mL, 0.411 mmol, 8 equiv.) was added. After 25 minutes deshydroxyethyl-dasatinib (0.025 g, 0.057 mmol, 1.1 equiv.) dissolved in DMF (0.25 mL) was added dropwise. The reaction was stirred at room temperature for 16 hours in the dark. The reaction mixture was purified by semi-preparative HPLC using a 30%-95% solvent B over 10 minutes method. The product eluted at 4.5 minutes and solvent was removed to afford the title compound (19 mg, 56%).

**$^1$H NMR** (500 MHz, DMSO) δ 11.51 (s, 1H), 9.88 (s, 1H), 8.22 (s, 1H), 7.85 (t, $J$ = 5.6 Hz, 1H), 7.40 (dd, $J$ = 7.7, 1.8 Hz, 1H), 7.30 – 7.22 (m, 2H), 6.06 (s, 1H), 3.62 – 3.52 (m, 8H), 2.91 (q, $J$ = 7.0 Hz, 2H), 2.83 (t, $J$ = 2.6 Hz, 1H), 2.58 (t, $J$ = 7.1 Hz, 2H), 2.42 (s, 3H), 2.33 (t, $J$ = 7.1 Hz, 2H), 2.24 (s, 3H), 1.99 (td, $J$ = 7.4, 2.6 Hz, 2H).
**$^{13}$C NMR** (126 MHz, DMSO) δ 171.34, 170.11, 165.20, 162.52, 162.23, 159.90, 156.96, 140.82, 138.81, 133.51, 132.42, 129.03, 128.18, 127.00, 125.73, 83.17, 82.75, 71.77, 44.04, 43.43, 43.19, 40.58, 40.11, 40.02, 39.94, 39.85, 39.78, 39.69, 39.61, 39.52, 39.44, 39.35, 39.19, 39.02, 33.58, 32.07, 31.35, 30.37, 27.82, 27.22, 25.58, 18.30, 12.68.
**LC-MS** $R_t$ = 5.75 min; m/z calcd. For $C_{31}H_{35}ClN_{10}O_3S$ ($[M+H]^+$) = 663.2, found 663.4

**Dasatnib-Dz-Alkyne** $^{1}$H-NMR

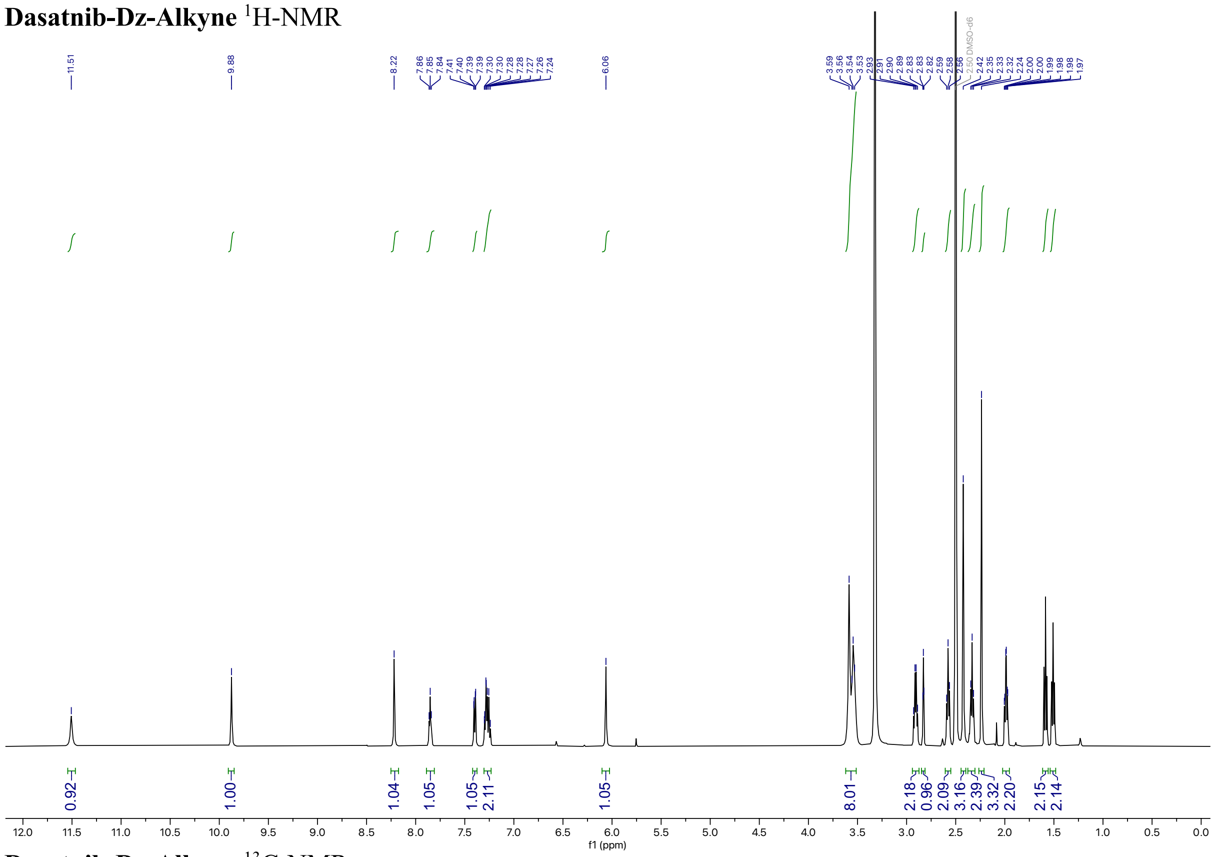

**Dasatnib-Dz-Alkyne** $^{13}$C-NMR

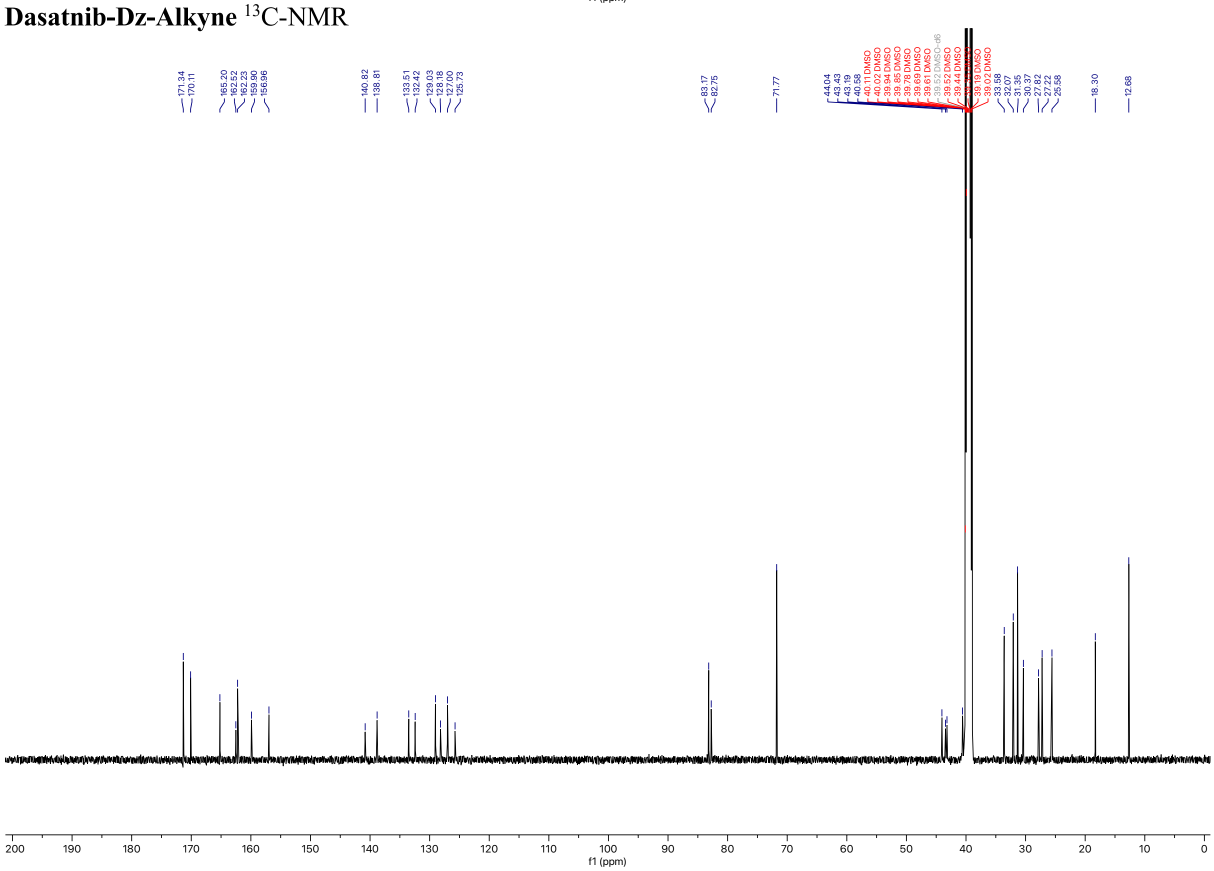

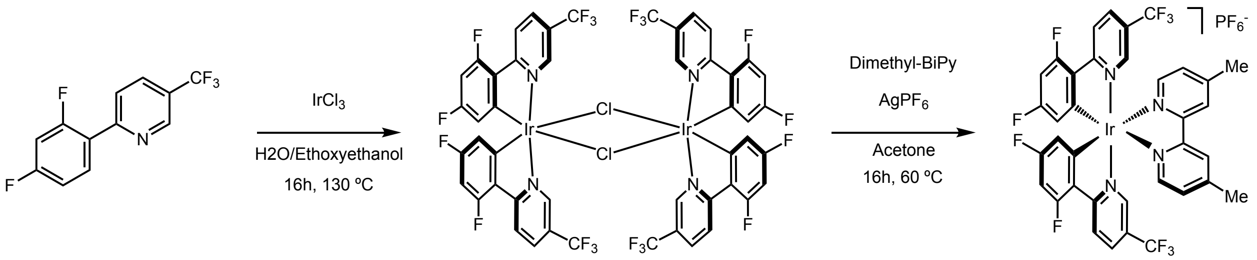

**Ir[dF(CF3)ppy]2(dMebipy)PF6 (16)**

A flame-dried schlenk flask equipped with a PTFE-coated stir bar was charged with 2-(2,4-difluorophenyl)-5-(trifluoromethyl)pyridine (2 g, 7.72 mmol, 2.1 equiv.) and iridium(III) chloride hydrate (1.095 g, 3.677 mmol, 1.0 equiv.). The flask was placed under an argon atmosphere and 2-ethoxyethanol:$H_2O$ (2:1, 74 mL) was added. The mixture was heated to 130 °C and stirred for 16 hours. The reaction was cooled to room temperature, quenched with cold water (300 mL), affording a precipitate. The precipitate was filtered and washed with excess cold water, then dried to afford the title compound as a bright orange powder which was carried forward without further purification. (2.590 g, 47%).

A scintillation vial equipped with a PTFE-coated stir bar was charged with the dimeric intermediate (0.060 g, 0.040 mmol, 1.0 equiv.), 4,4'-dimethyl-2,2'-bipyridine (0.016 g, 0.009 mmol, 2.2 equiv.), and silver hexafluorophosphate (0.012 g, 0.031 mmol, 3.0 equiv.). Acetone (0.8 mL) was added, the mixture was heated to 60 °C, and subsequently stirred for 16 hours. The reaction was cooled to room temperature, filtered to remove salts, and solvent was removed. The crude mixture was dissolved in DMSO (0.7 mL) and purified by preparative HPLC using a 50-95% solvent B gradient over 15 minutes with 150 µL MeCN sample sandwiching during injection. The product eluted at 5.65 minutes and was concentrated to afford the title compound as a yellow solid (70 mg, 97%). Spectral data were consistent with existing literature [31].

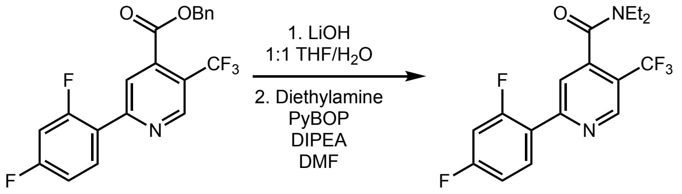

**2-(2,4-difluorophenyl)-*N,N*-diethyl-5-(trifluoromethyl)isonicotinamide (17)**
A scintillation vial equipped with a PTFE-coated stir bar was charged with **13** (0.1 g, 0.264 mmol, 1.0 equiv.). The solid was suspended in THF (1.31 mL) and aqueous LiOH (2 M, 1.31 mL, 10.0 equiv.) was added. The mixture was stirred vigorously at room temperature. After 16 hours solvent was removed and the reaction mixture partitioned between $H_2O$ (30mL) and EtOAc (30 mL). The aqueous layer was acidified to pH 2 with HCl, then extracted 3x with EtOAc (30 mL). The EtOAc extracts were washed with brine, dried with $Na_2SO_4$, and concentrated to remove solvent. The product was used without further purification.

A scintillation vial equipped with a PTFE-coated stir bar was charged with the hydrolysis product (0.075 g, 0.248 mmol, 1.0 equiv.) and PyBOP (0.644 g, 1.238 mmol, 5 equiv.). DMF (2.475 mL) and DIPEA (0.431 mL, 2.475 mmol, 10 equiv.) were added, and the solution was stirred at room temperature for 10 minutes. Diethylamine (0.102 mL, 0.990 mmol, 4 equiv.) was added dropwise and the reaction was heated to 70 °C and stirred for 16 hours. The reaction was cooled to room temperature and the solvent was removed. The crude residue was partitioned between $H_2O$ (30mL) and DCM (30 mL), and the aqueous layer was extracted 3x with DCM (30 mL) and the DCM extracts were washed 3x $H_2O$ (100 mL) to remove excess DMF. The organic phase was then washed with brine, dried with $Na_2SO_4$, and concentrated to remove solvent. The crude mixture was purified by flash column chromatography on silica (Hex:EtOAc) = 0% to 10% over 5 CVs followed by 10% to 30% over 10 CVs to afford the title compound as white solid (0.075 g, 85%).

**$^1$H NMR** (598 MHz, $CDCl_3$) δ 8.98 (s, 1H), 8.11 (s, 1H), 7.74 (s, 1H), 7.04 (s, 1H), 6.93 (s, 1H), 3.88 (s, 1H), 3.27 (s, 1H), 1.25 (d, $J$ = 14.3 Hz, 1H), 1.11 (s, 1H).
**$^{13}$C NMR** (150 MHz, $CDCl_3$) δ 165.70, 165.13, 165.05, 163.44, 163.36, 162.00, 161.92, 160.31, 160.23, 156.22, 147.98, 147.95, 147.91, 147.88, 144.07, 132.65, 132.62, 132.58, 132.56, 126.10, 124.28, 122.46, 122.01, 121.99, 121.94, 121.91, 121.24, 121.17, 120.84, 120.62, 112.64, 112.61, 112.50, 112.47, 104.97, 104.80, 104.62, 42.96, 39.00, 13.55, 12.17.
**$^{19}$F NMR** (563 MHz, $CDCl_3$) δ -59.76 (s, 3F), -106.31 (s, 1F), -111.75 (s, 1F).
**LC-MS** $R_t$ = 7.14 min; m/z calcd. For $C_{17}H_{15}F_5N_2O$ ($[M+H]^+$) = 359.1, found 359.3

**17** $^{1}$H-NMR

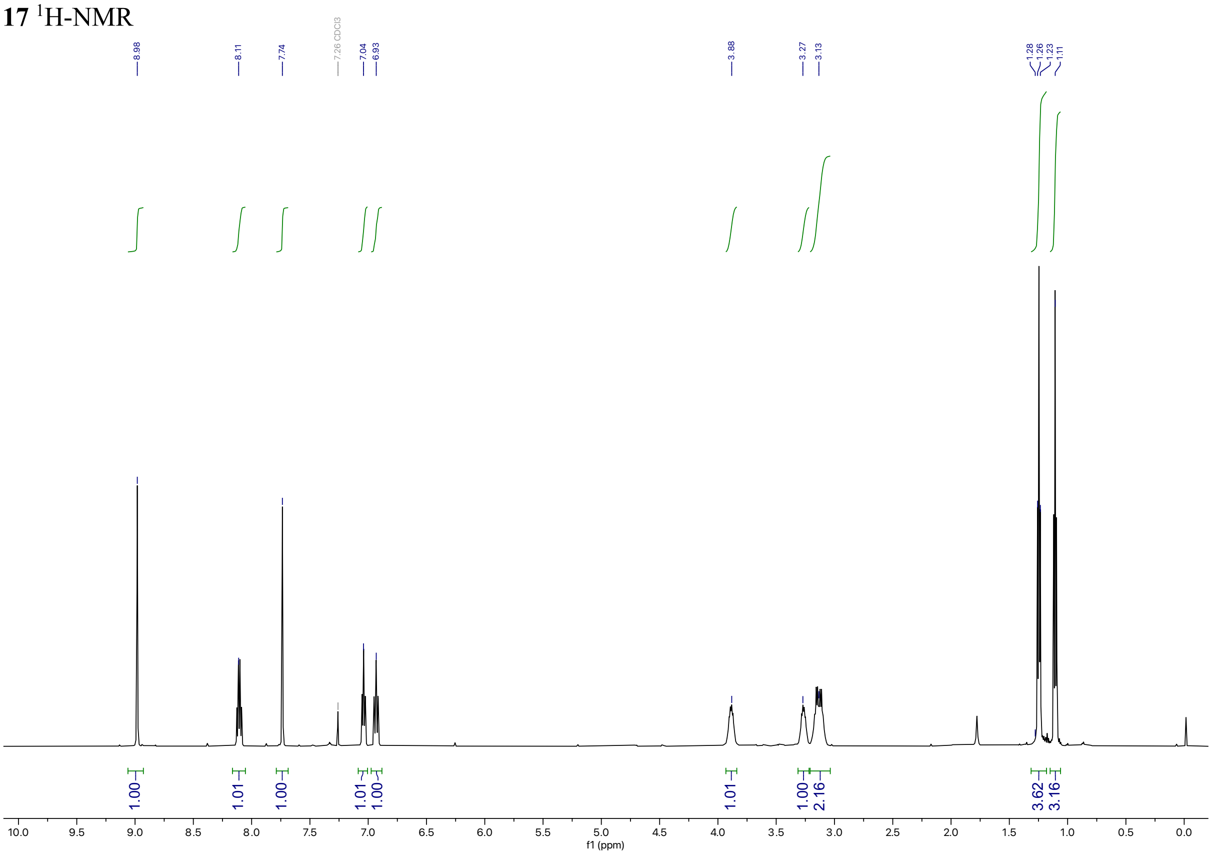

**17** $^{13}$C-NMR

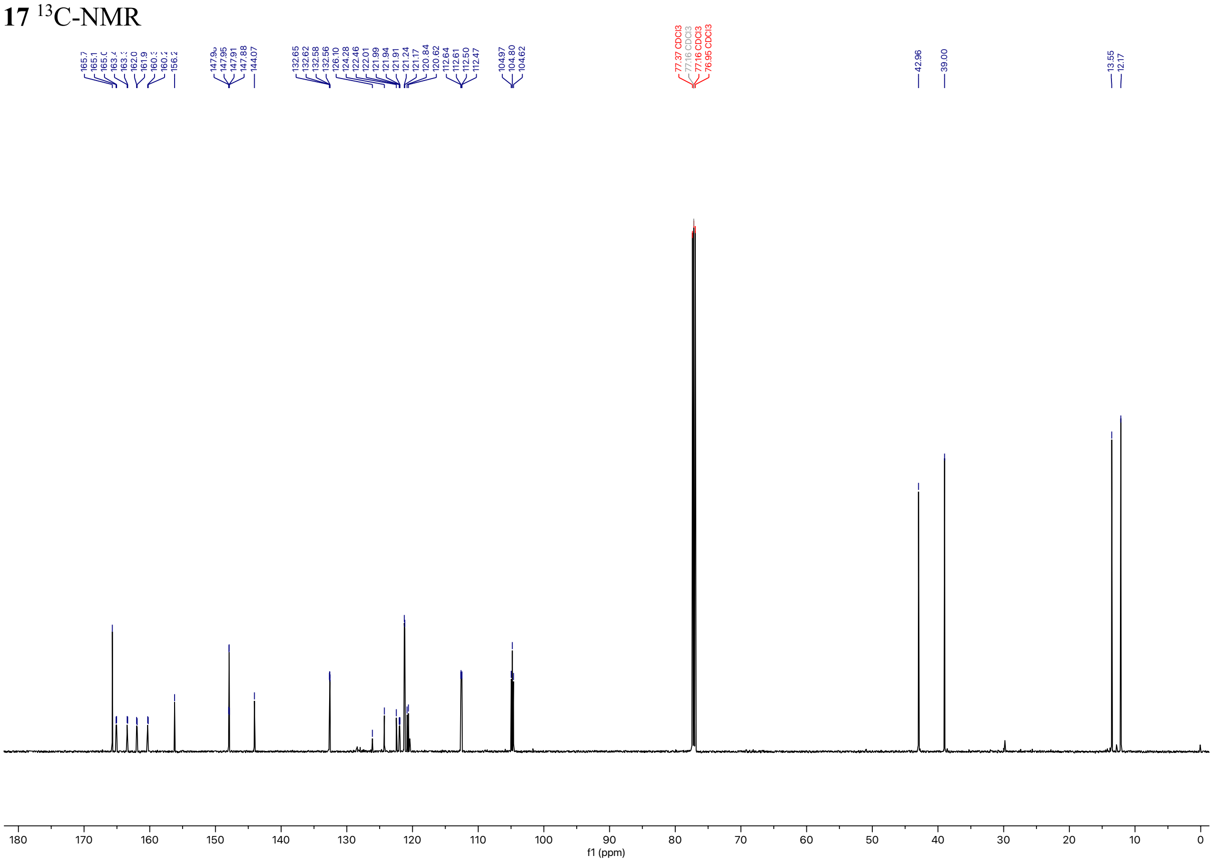

**17** $^{19}$F-NMR

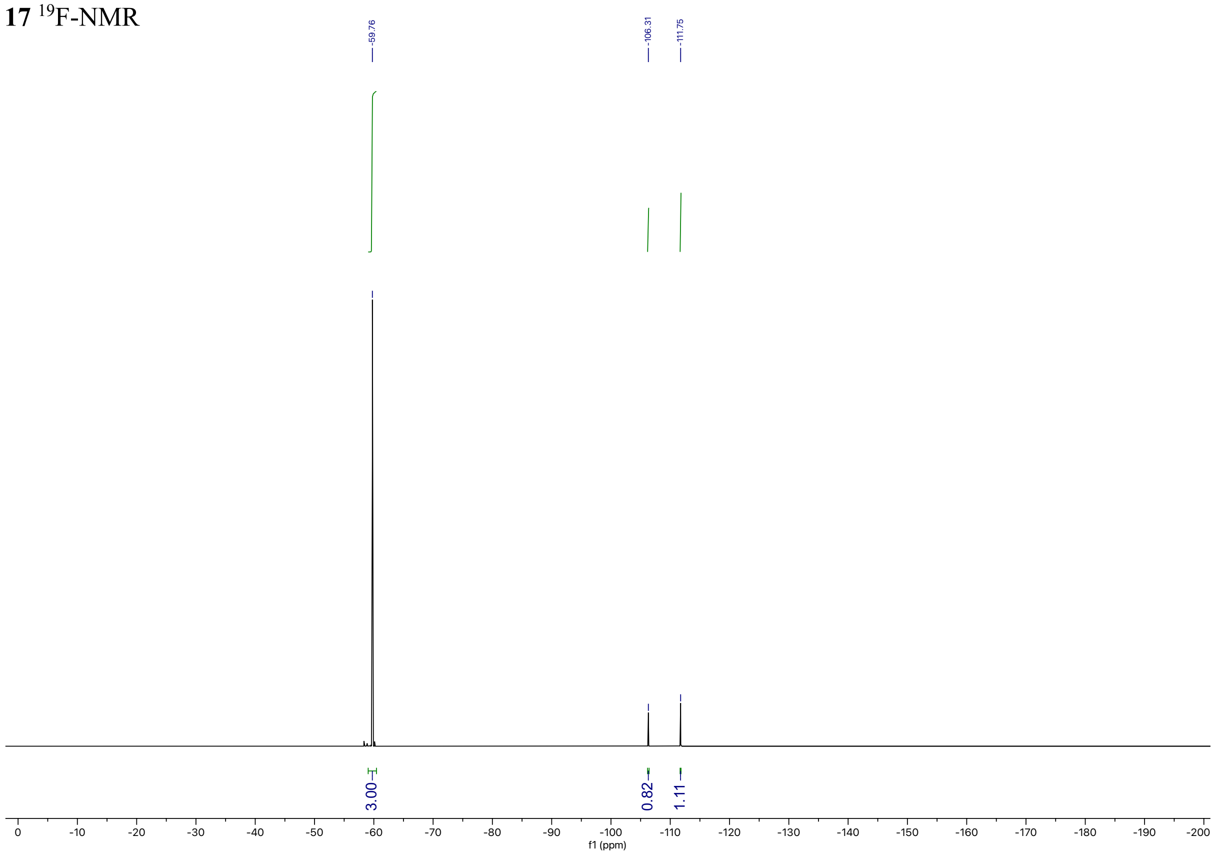
-59.76
-106.31
-111.75
3.00
0.82
1.11
0
-10
-20
-30
-40
-50
-60
-70
-80
-90
-100
-110
-120
-130
-140
-150
-160
-170
-180
-190
-200
f1 (ppm)

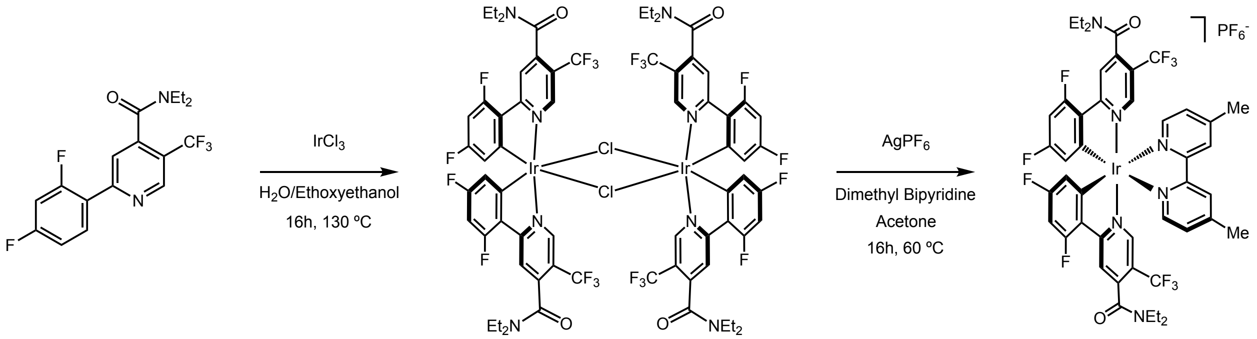

**Ir[diethylamideppy]$_2$(dMebipy)PF$_6$ (18)**
A flame-dried schlenk flask equipped with a PTFE-coated stir bar was charged with **17** (0.063 g, 0.035 mmol, 2.1 equiv.) and iridium(III) chloride hydrate (0.025 g, 0.083 mmol, 1.0 equiv.). The flask was placed under an argon atmosphere and 2-ethoxyethanol:$H_2O$ (2:1, 1.67 mL) was added. The mixture was heated to 130 °C and stirred for 16 hours. The reaction was cooled to room temperature, quenched with cold water (200 mL), generating a precipitate. The precipitate was filtered and washed with excess cold water then dried to afford the title compound as a bright orange powder which was used without further purification. (0.066 g, 85%).

A scintillation vial equipped with a PTFE-coated stir bar was charged with dimeric intermediate (0.033 g, 0.018 mmol, 1.0 equiv.), 4,4'-dimethyl-2,2'-bipyridine (0.007 g, 0.053 mmol, 2.2 equiv.), and silver hexafluorophosphate (0.013 g, 0.053 mmol, 3.0 equiv.). Acetone (0.7 mL) was added, and the mixture heated to 60 °C for 16 hours with stirring. The reaction was cooled to room temperature, filtered to remove salts, and the solvent was removed. The crude mixture was dissolved in MeCN/$H_2O$/DMSO (1:1:2, 1.5 mL) and purified by preparative HPLC using a 35-95% solvent B over 10 minute method with a 150 µL MeCN sample sandwich method during injection. The product eluted at 6.0 minutes and solvent was removed to afford the title compound as a yellow solid (25.1 mg, 65%).

**$^1$H NMR** (598 MHz, DMSO) δ 8.78 (dd, *J* = 29.4, 13.0 Hz, 2H), 8.32 (d, *J* = 4.8 Hz, 1H), 8.27 (d, *J* = 8.7 Hz, 1H), 7.80 – 7.56 (m, 6H), 6.95 (q, *J* = 11.8 Hz, 2H), 6.04 (d, *J* = 7.9 Hz, 1H), 5.52 (dd, *J* = 35.9, 8.2 Hz, 1H), 3.48 (dtdd, *J* = 38.2, 31.5, 13.9, 6.5 Hz, 5H), 2.98 (s, 3H), 2.62 – 2.57 (m, 6H), 1.28 – 1.14 (m, 6H), 1.02 (ddt, *J* = 46.3, 14.9, 7.1 Hz, 6H).
**$^{19}$F NMR** (563 MHz, DMSO) δ -61.44 (dd, *J* = 44.7, 30.5 Hz, 6F), -70.36 (d, *J* = 711.4 Hz, 2F), -95.47 (dt, *J* = 32.9, 12.9 Hz, 1F), -96.49 (d, *J* = 8.4 Hz, 1F), -102.95 (dd, *J* = 44.9, 11.0 Hz, 1F), -103.21 (dd, *J* = 140.9, 10.4 Hz, 1F).
**LC-MS** $R_t$ = 5.80 min; m/z calcd. For $C_{46}H_{40}F_{10}IrN_6O_2$ ($[M+H]^+$) = 1092.3, found 1092.6

**18** $^{1}$H-NMR

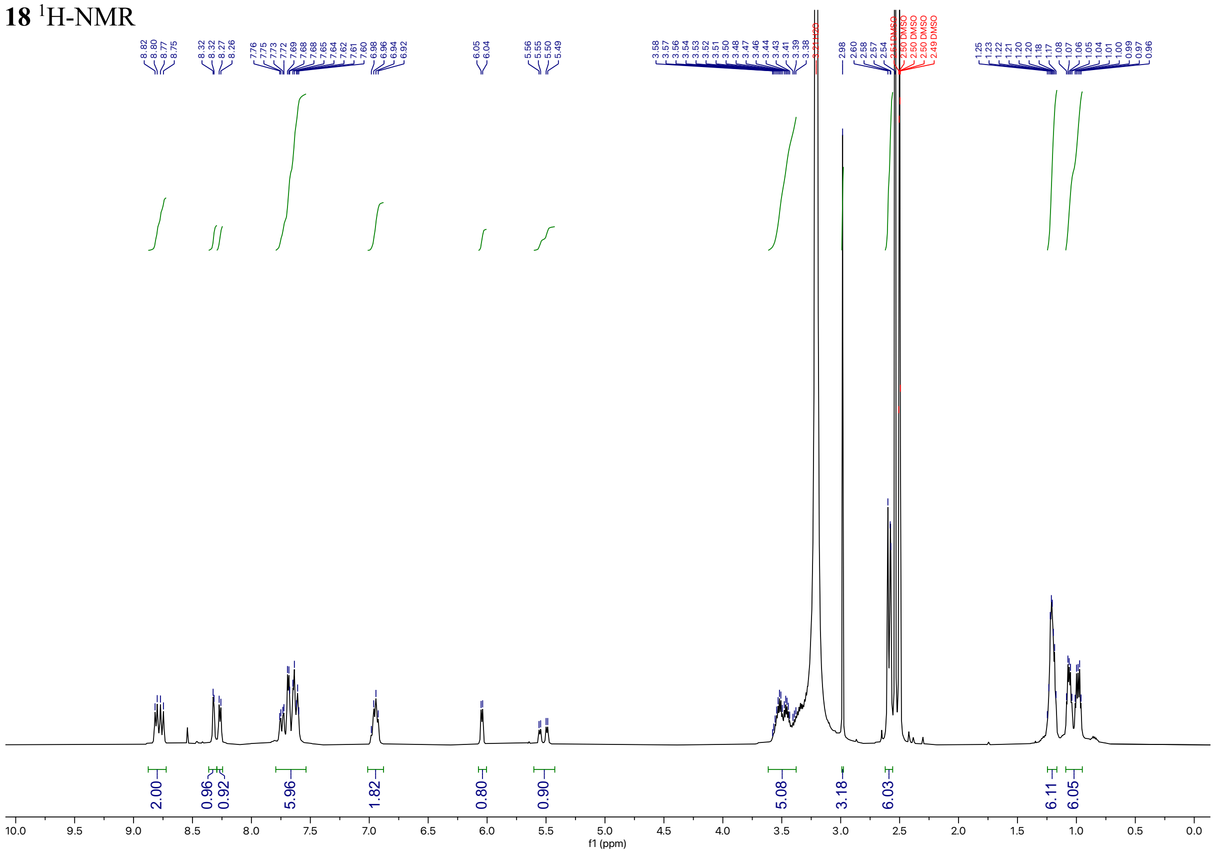

**18** $^{19}$F-NMR

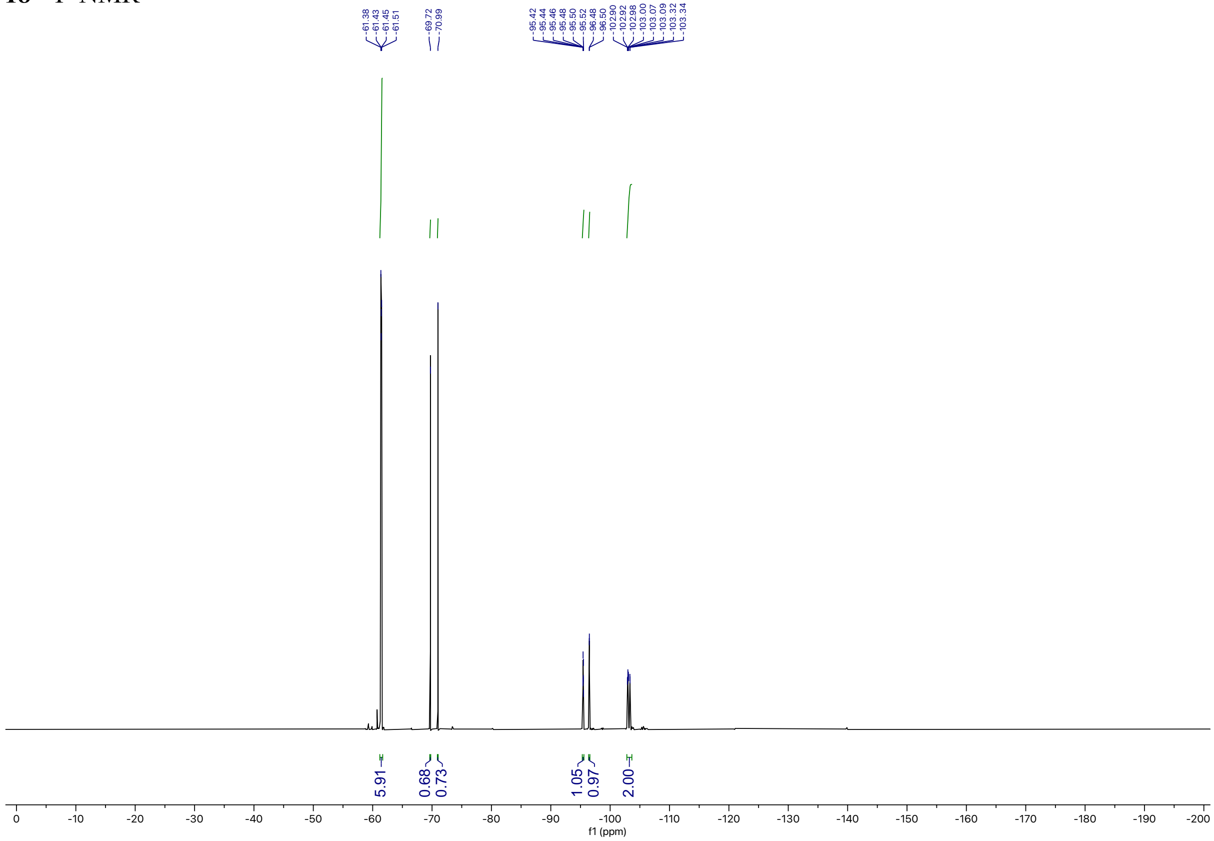

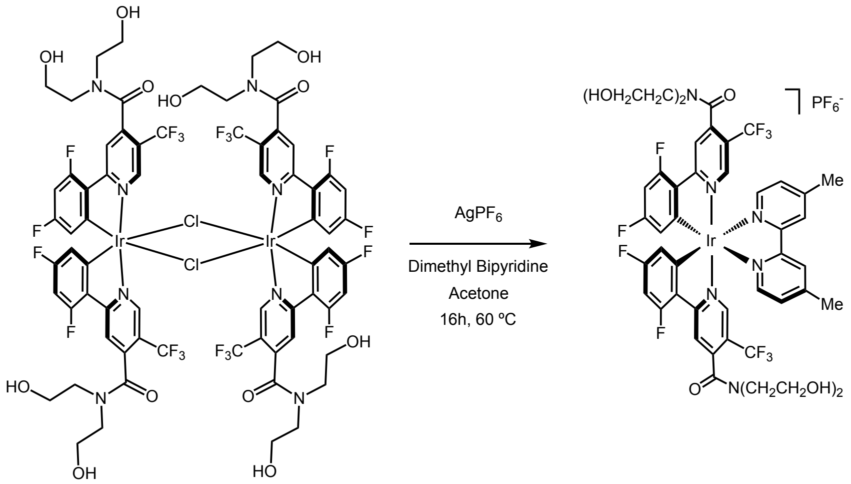

**Ir[dF(CF$_3$)(OBN)ppy]$_2$(dMebipy)PF$_6$ complex (19)**

A scintillation vial equipped with a PTFE-coated stir bar was charged with dimeric intermediate (**15**) (0.041 g, 0.020 mmol, 1.0 equiv.), 4,4'-dimethyl-2,2'-bipyridine (0.009 g, 0.050 mmol, 2.2 equiv.), and silver hexafluorophosphate (0.015 g, 0.061 mmol, 3.0 equiv.). Acetone (1 mL) was added, the mixture was heated to 60 °C, and stirred at this temperature for 16 hours. The reaction was cooled to room temperature, filtered to remove salts, and solvent was removed. The crude mixture was dissolved in DMSO (0.8 mL) and purified by preparative HPLC using a 20-95% solvent B over 10 minutes method with a 150 µL MeCN sample sandwiching method during injection. The product eluted at 4.4 minutes and was concentrated to afford the title compound as a yellow solid (37 mg, 79%).

**$^{1}$H NMR** (598 MHz, DMSO) δ 8.74 (d, $J$ = 48.8 Hz, 2H), 8.51 – 8.27 (m, 2H), 7.87 (d, $J$ = 35.0 Hz, 2H), 7.75 – 7.35 (m, 4H), 7.04 (t, $J$ = 10.6 Hz, 2H), 5.84 (br, s, 2H), 4.88 (br, s, 2H), 4.72 (s, 2H), 3.50 (m, 16H), 2.59 (s, 6H).
**$^{19}$F NMR** (563 MHz, DMSO) δ -59.67 (s, 6F), -102.54 (s, 2F), -106.00 (s, 2F).
**LC-MS** $R_t$ = 3.07 min (45%-95% B over 10 minutes); m/z calcd. For $C_{46}H_{40}F_{10}IrN_6O_6$ ($[M+H]^+$) = 1156.2, found 1156.7

**19** $^{1}$H-NMR

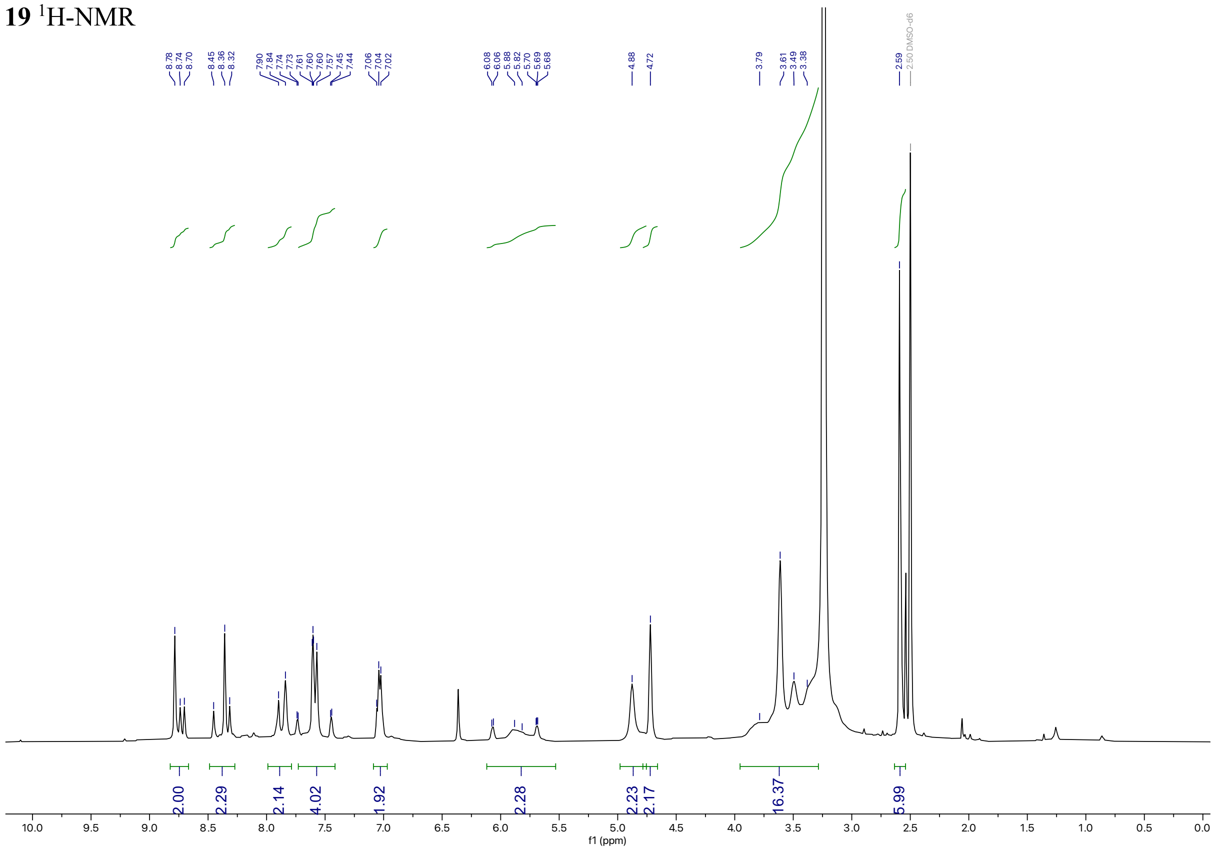

**19** $^{19}$F-NMR

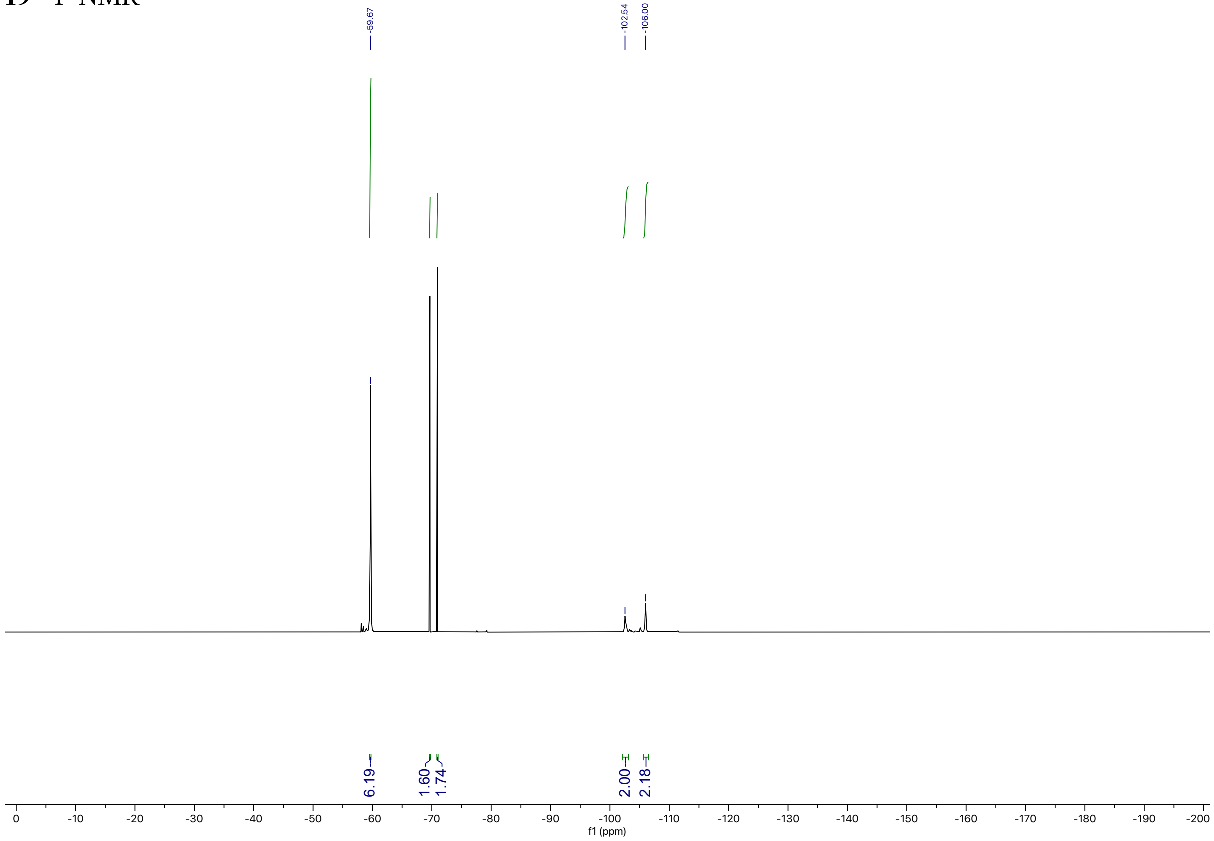

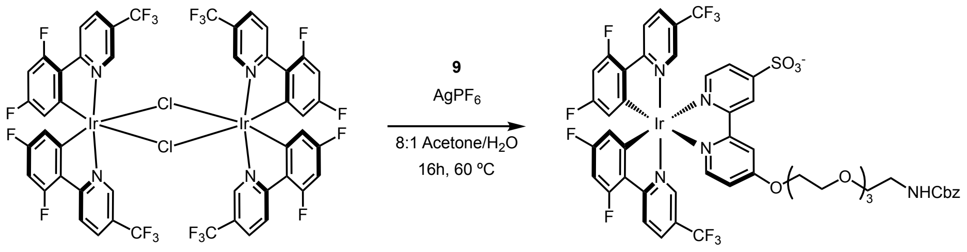

**Ir[dF(CF$_3$)ppy]$_2$(sulfo-ether-bipy) (20)**
A scintillation vial equipped with a PTFE-coated stir bar was charged with dimer (**16**) (0.008 g, 0.005 mmol, 1.0 equiv.), **9** (0.006 g, 0.011 mmol, 2.2 equiv.), and silver hexafluorophosphate (0.004 g, 0.016 mmol, 3.0 equiv.). Acetone (0.5 mL) was added, the mixture was heated to 60 °C, and stirred for 16 hours. The reaction was cooled to room temperature, filtered to remove salts, and the solvent was removed. The crude mixture was dissolved in DMSO (0.7 mL) and purified by semipreparative HPLC using a 5-95% solvent B over 10 minutes method with a 75 µL MeCN sample sandwich method during injection to afford the title compound as a yellow solid (6.3 mg, 46%).

**$^{1}$H NMR** (598 MHz, DMSO) δ 9.00 (s, 1H), 8.52 (s, 1H), 8.47 (d, $J$ = 9.4 Hz, 2H), 8.42 (d, $J$ = 9.5 Hz, 1H), 7.99 (d, $J$ = 6.2 Hz, 1H), 7.84 (d, $J$ = 5.7 Hz, 1H), 7.69 (s, 3H), 7.37 – 7.25 (m, 6H), 7.06 – 6.96 (m, H), 5.75 (dd, $J$ = 24.1, 8.1 Hz, 2H), 5.00 (s, 2H), 4.51 (d, $J$ = 4.9 Hz, 2H), 3.81 (d, $J$ = 4.9 Hz, 2H), 3.60 (d, $J$ = 4.7 Hz, 2H), 3.53 (d, $J$ = 4.5 Hz, 2H), 3.49 (s, 4H), 3.43 – 3.39 (m, 2H), 3.13 (d, $J$ = 6.9 Hz, 2H).
**$^{19}$F NMR** (563 MHz, DMSO) δ -61.15 – -61.93 (m, 6F), -103.43 (t, $J$ = 9.5 Hz, 2F), -106.79 (t, $J$ = 12.4 Hz, 2F).
**LC-MS** $R_t$ = 7.16 min; m/z calcd. For $C_{50}H_{40}F_{10}IrN_5O_9S$ ($[M+H]^+$) = 1270.2, found 1270.2

**20** $^{1}$H-NMR

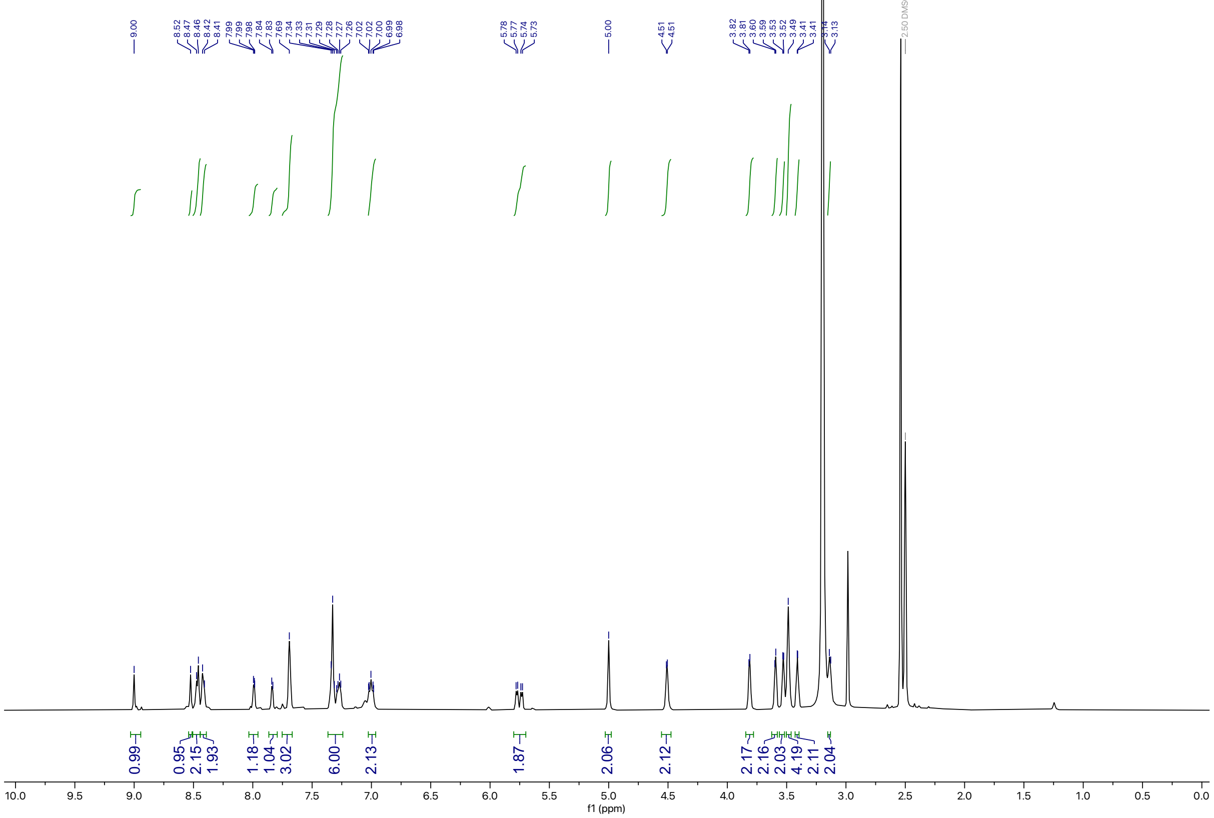

**20** $^{19}$F-NMR

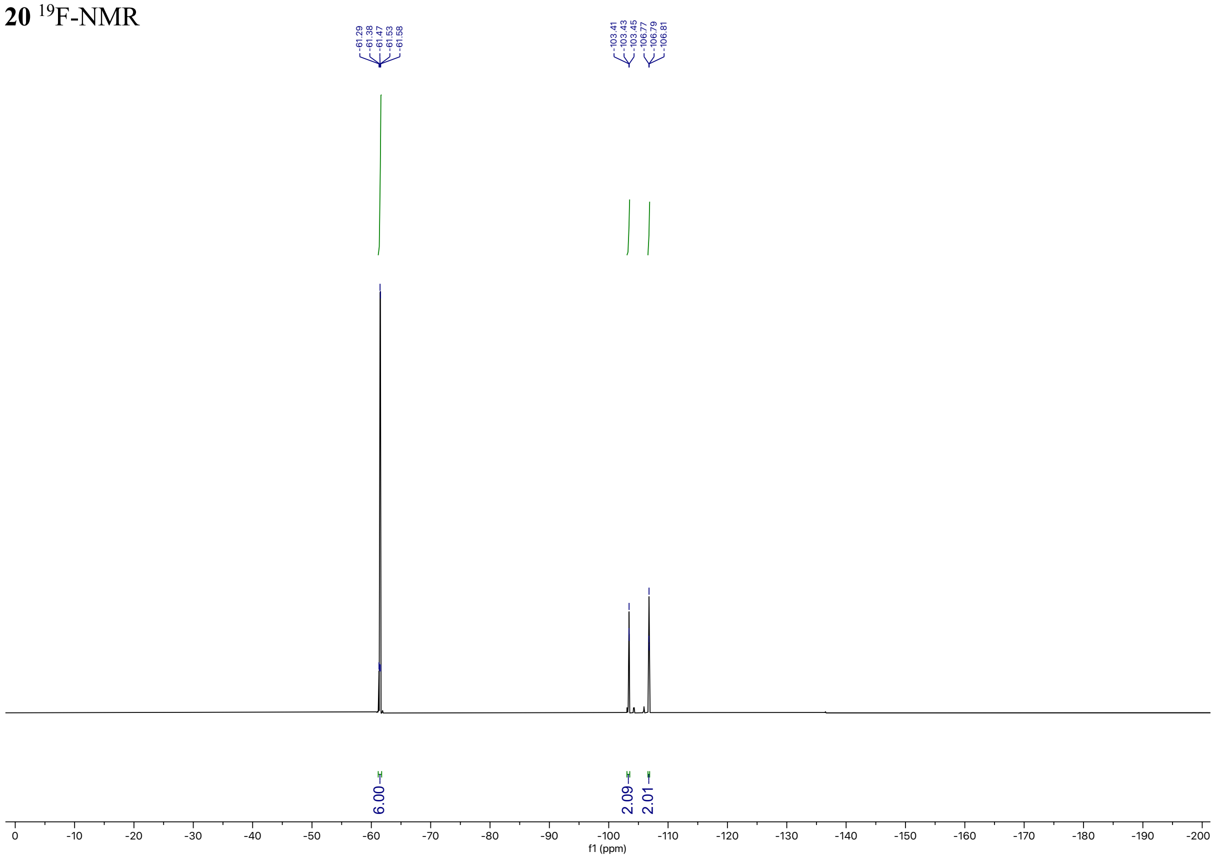

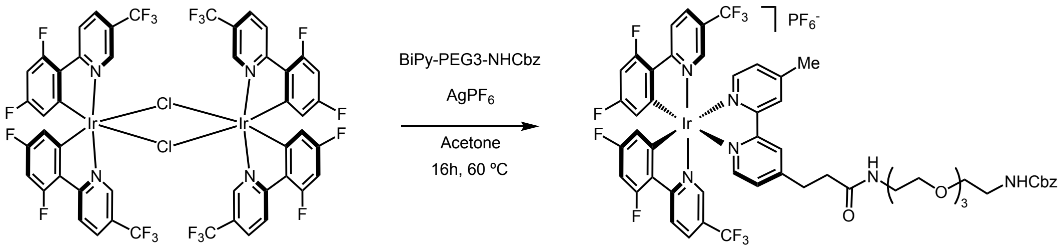

**2**

A scintillation vial equipped with a PTFE-coated stir bar was charged with dimer (**16**) (0.062 g, 0.042 mmol, 1.0 equiv.), **bipy-Me-PEG3-NHCbz** (0.050 g, 0.092 mmol, 2.2 equiv.), and silver hexafluorophosphate (0.031 g, 0.125 mmol, 3.0 equiv.). Acetone (0.83 mL) was added, the mixture was heated to 60 °C, and stirred for 16 hours. The reaction was cooled to room temperature, filtered to remove salts, and the solvent was removed. The crude mixture was dissolved in DMSO (1.2 mL) and purified by preparative HPLC using a 35-95% solvent B over 15 minutes method with a 75 µL MeCN sample sandwich method during injection to afford the title compound as a yellow solid (32.5 mg, 28%).

**$^{1}$H NMR** (598 MHz, $CDCl_3$) δ 8.71 – 8.40 (m, 4H), 8.03 (q, *J* = 10.4 Hz, 2H), 7.80 – 7.66 (m, 2H), 7.59 (s, 1H), 7.54 – 7.42 (m, 2H), 7.32 (d, *J* = 8.0 Hz, 5H), 6.93 (s, 1H), 6.70 – 6.58 (m, 2H), 5.63 (d, *J* = 7.8 Hz, 1H), 5.54 (s, 1H), 5.07 (s, 3H), 3.66 – 3.53 (m, 10H), 3.49 (t, *J* = 5.7 Hz, 2H), 3.36 (dt, *J* = 10.1, 5.3 Hz, 2H), 3.17 (dq, *J* = 14.0, 7.5 Hz, 2H), 2.76 – 2.70 (m, 4H), 1.99 (s, 3H).
**$^{19}$F NMR** (563 MHz, $CDCl_3$) δ -62.80 (d, *J* = 39.3 Hz, 6F), -71.98 (d, *J* = 712.8 Hz, 6F), -101.41 (dd, *J* = 53.7, 12.5 Hz, 2F), -105.86 (dd, *J* = 41.7, 12.6 Hz, 2F).
**LC-MS** $R_t$ = 4.68 min; m/z calcd. For $C_{54}H_{48}F_{10}IrN_6O_6$ ($[M+H]^+$) = 1260.3, found 1260.8

**2** $^{1}$H-NMR

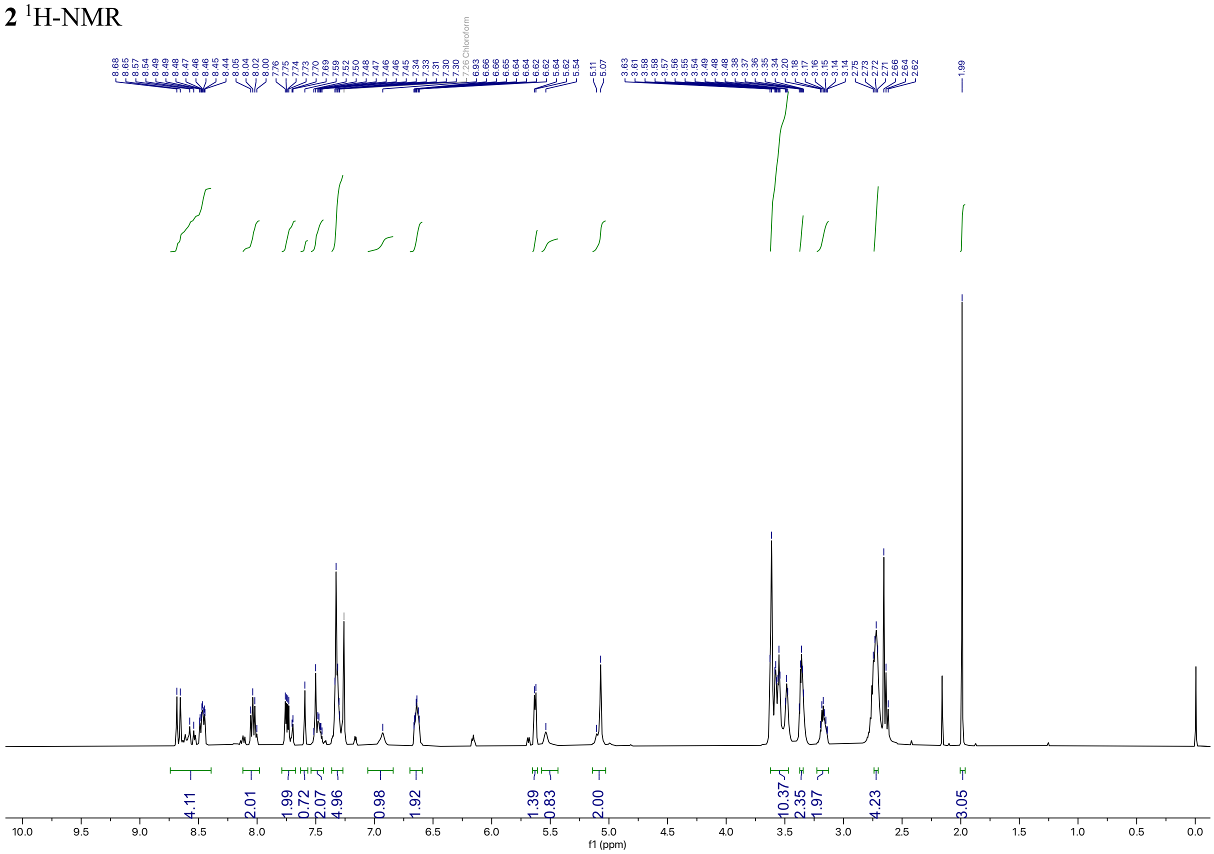

**2** $^{19}$F-NMR

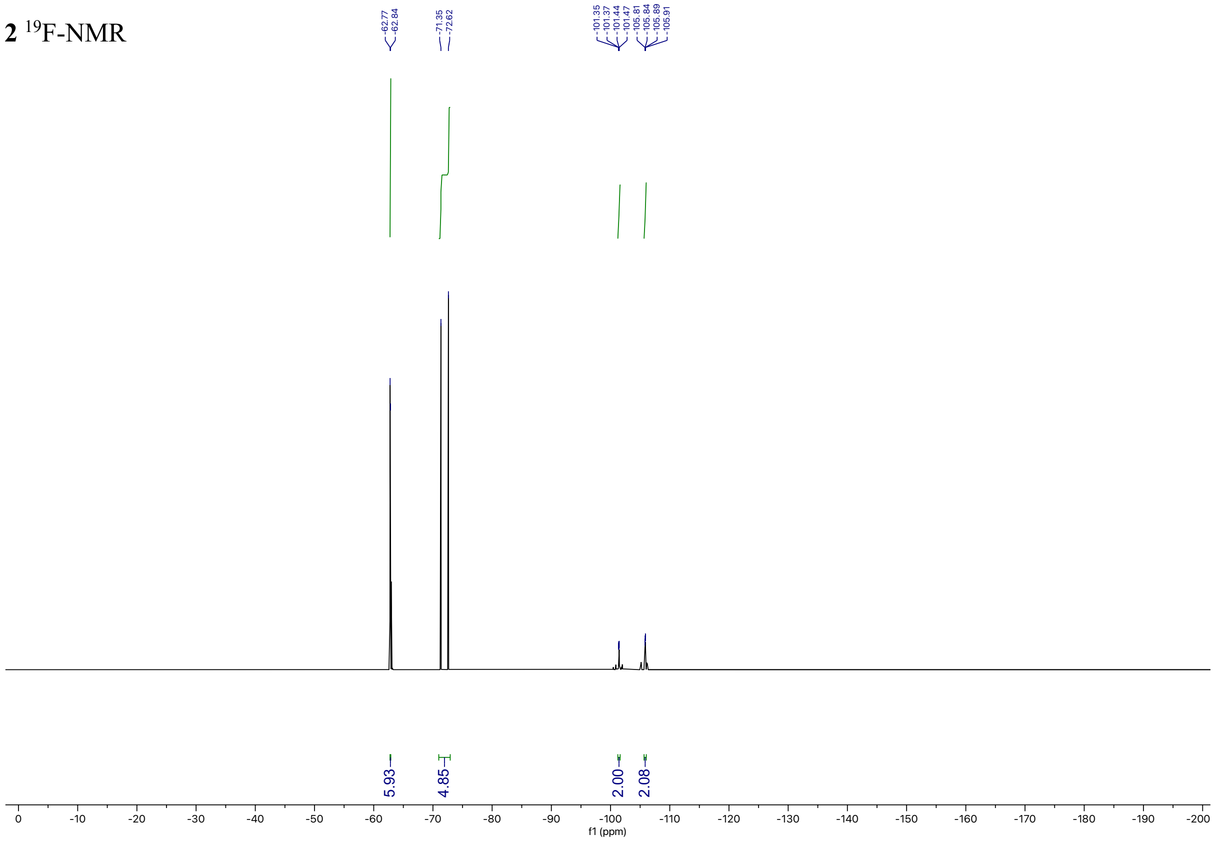

**Peptide photocatalyst conjugates**

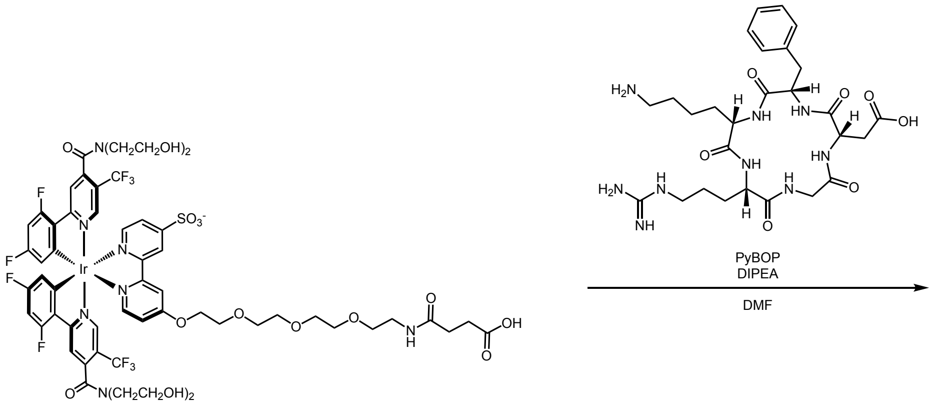

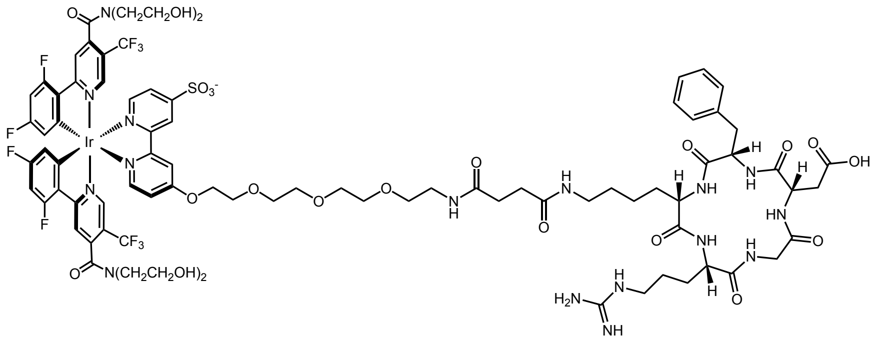

**1-cRGDFK**

A scintillation vial equipped with a PTFE-coated stir bar was charged with **1-CO$_2$H** (0.007 g, 0.005 mmol, 1.0 equiv.), PyBOP (0.002 g, 0.005 mmol, 1.0 equiv.), and in DMF (0.25 mL). The solution was stirred at room temperature for 5 minutes, then DIPEA (0.007 mL, 0.038 mmol, 8 equiv.) was added. After 35 minutes, cRGDFK (0.003 g, 0.005 mmol, 1.1 equiv.) dissolved in DMF (0.12 mL) was added dropwise and the reaction was stirred at room temperature for 16 hours. The reaction mixture was then diluted with $H_2O$ (0.2 mL) and purified by semi-preparative HPLC using a 22% solvent B 25 minute isocratic method. The product eluted at 18.4 minutes and solvent was removed to afford the title compound as a yellow solid (0.5 mg, 5%).

**LC-MS** $R_t$ = 4.65 min; m/z calcd. For $C_{83}H_{95}F_{10}IrN_{16}O_{22}S$ ($[M+2H]^{2+}$) = 1043.2, found 1043.2

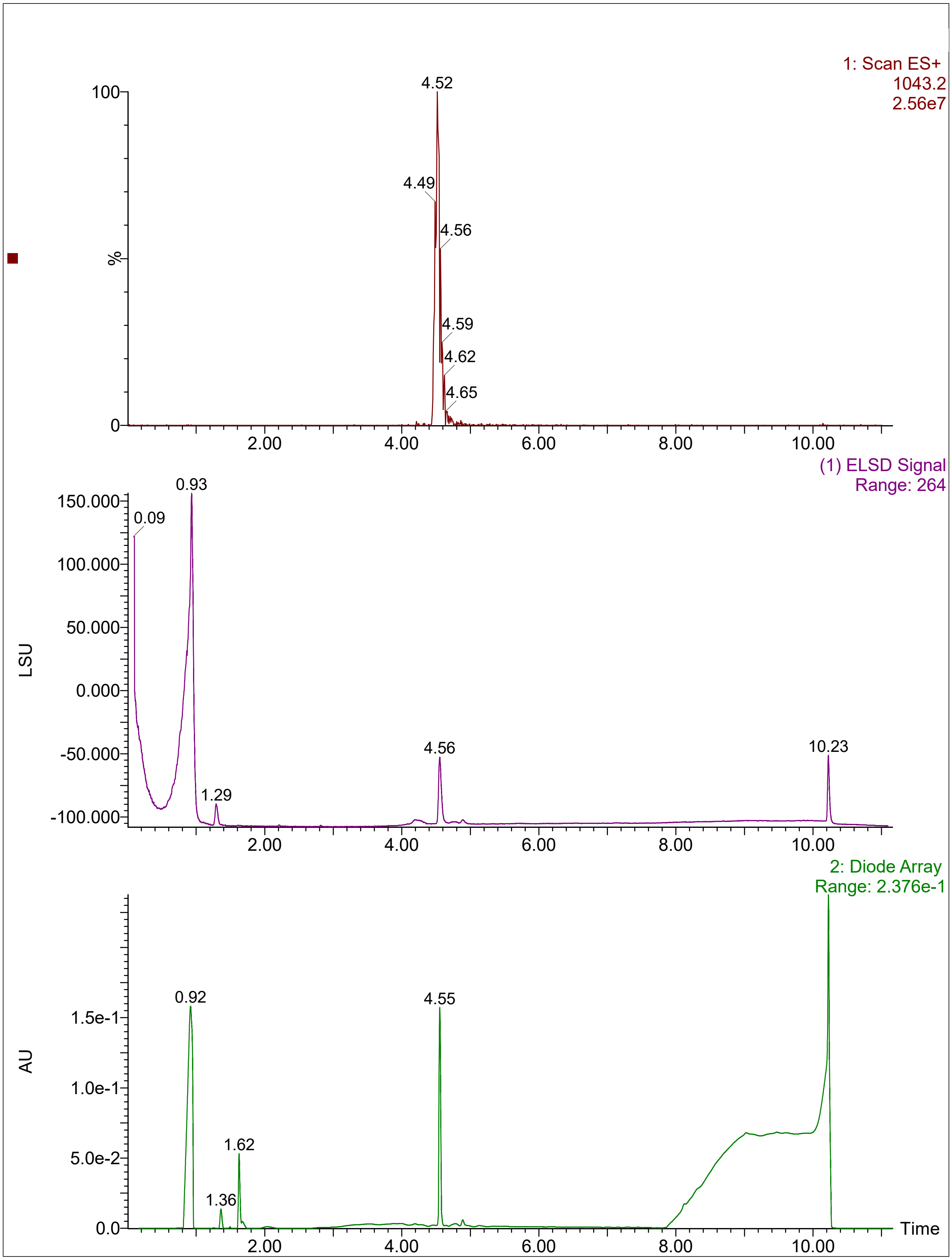
1: Scan ES+
1043.2
2.56e7
4.52
4.49
4.56
4.59
4.62
4.65
%
100
0
2.00
4.00
6.00
8.00
10.00
(1) ELSD Signal
Range: 264
LSU
150.000
100.000
50.000
0.000
-50.000
-100.000
0.93
0.09
1.29
4.56
10.23
2: Diode Array
Range: 2.376e-1
AU
1.5e-1
1.0e-1
5.0e-2
0.0
0.92
1.62
1.36
4.55
Time

Purification of peptide ligands lacking N or C-terminal cysteines:

Peptides were purchased as crude mixtures from GenScript at 9 mg scales. The lyophilized powders were dissolved in 100 µL DMSO. 50 µL of this solution was purified using semipreparative HPLC using a 20-60% B over 10 minutes method, and solvent was removed to afford pure peptides for competitive labeling experiments. Purity was validated by HPLC-MS-ELSD using a 5%-95% solvent B over 6 minutes method.

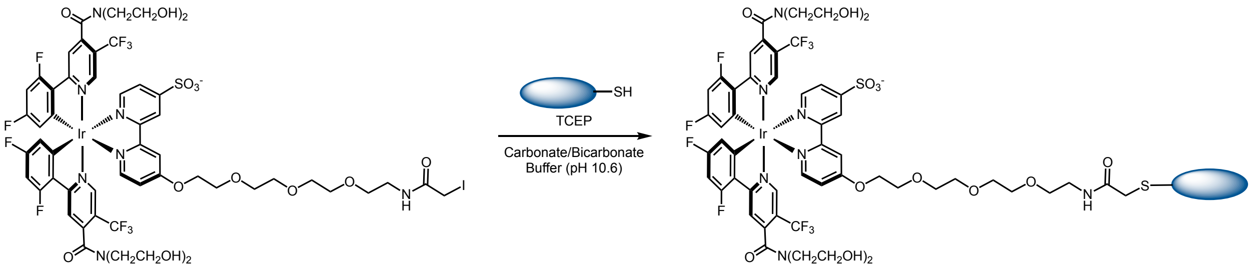

Peptide-photocatalyst conjugation:

Peptides containing terminal cysteines were purchased as crude mixtures from GenScript at 9 mg scales. The lyophilized powders were dissolved in 100 µL DMSO. 25 µL of this solution was transferred to a 1.5 mL centrifuge tube. To this tube was added DMSO (25 µL), $H_2O$ (45 µL) and TCEP (5 µL, 100 mM in $H_2O$, final concentration 5 mM). The mixture was sonicated for 10 minutes at room temperature to dissolve solids and reduce disulfide bonds. **1-IA** (1 mg, 50 µL of 12.5 mM solution in DMSO) was added followed by carbonate/bicarbonate buffer (100 mM, pH = 10.6, 50 µL). The reactions were then inverted in the dark at 37 °C for 16 hours. Conjugated peptides were isolated using an appropriate semipreparative HPLC-MS method. Products were concentrated in 1.5 mL centrifuge tubes under vacuum then redissolved in 100 µL DMSO for characterization and purity analysis via HPLC-MS-ELSD using a 5%-95% solvent B over 6 minutes method. Concentrations of the resulting stock solutions were determined by absorbance at 400 nm.

| Peptide Sequence | Name | Catalyst Conjugate or Unconjugated | HPLC Gradient | Retention Time (min.) | Molecular Weight | Mass(es) Observed |
|---|---|---|---|---|---|---|
| AVPIAQKSE | SI-Pep-1/ Smac peptide | Unconjugated | 20%-50% solvent B over 10 min. | 1.96 | 942.07 | $[M+H]^{+}$ = 942.9 $[M+2H]^{2+}$ = 472.2 |
| AVPIAQKSEC | SI-Pep-2/ **1**-Smac peptide | Conjugate | 20%-50% solvent B over 10 min. | 3.20 | 2484.21 | $[M+2H]^{2+}$ = 1242.4 $[M+3H]^{3+}$ = 828.6 |
| MRVKEKYQHLWRWGWRWGTMLLG | SI-Pep-3/ HIV-1 gp160 SPF | Unconjugated | 35%-95% solvent B over 10 min. | 5.42 | 3018.5 | $[M+4H]^{4+}$ = 755.8 $[M+5H]^{5+}$ = 604.9 |
| MRVKEKYQHLWRWGWRWGTMLLGC | SI-Pep-4/ **1**-HIV-1 gp160 SPF | Conjugate | 20%-50% solvent B over 10 min. | 3.87 | 4559 | $[M+6H]^{6+}$ = 728.8 |
| MPVWWRRRRLRARSWALRGARKPLR | SI-Pep-5/ SNHG6 lncRNA ORF | Unconjugated | 5%-95% solvent B over 10 min. | 2.71 | 3231.89 | $[M+6H]^{6+}$ = 539.7 $[M+7H]^{7+}$ = 462.7 |
| CMPVWWRRRRLRARSWALRGARKPLR | SI-Pep-6/ **1**- SNHG6 lncRNA ORF | Conjugate | 20%-60% solvent B over 10 min. | 2.41 | 4774.03 | $[M+7H]^{7+}$ = 682.9 |
| EVPVPPPVPPRRRP | SI-Pep-7/ SOS1 | Unconjugated | 20%-50% solvent B over 10 min. | 2.19 | 1592.89 | $[M+2H]^{2+}$ = 797.4 $[M+3H]^{3+}$ = 532.0 |
| EVPVPPPVPPRRRPC | SI-Pep-8 / **1**-SOS1 | Conjugate | 5%-95% solvent B over 10 min. | 3.14 | 3135.03 | $[M+3H]^{3+}$ = 1046.0 $[M+4H]^{4+}$ = 784.64 $[M+5H]^{5+}$ = 628.0 |
| MPSSRAV | SI-Pep-9 / STAT1 uORF | Unconjugated | 20%-60% solvent B over 10 min. | 2.01 | 746.88 | $[M+H]^{+}$ = 747.7 $[M+2H]^{2+}$ = 374.8 |
| MPSSRAVC | SI-Pep-10 / **1**-STAT1 uORF | Conjugate | 20%-60% solvent B over 10 min. | 3.34 | 2289.02 | $[M+2H]^{2+}$ = 1145.5 |

**Characterization Data: HPLC-MS Spectra:**
SI-Pep-1

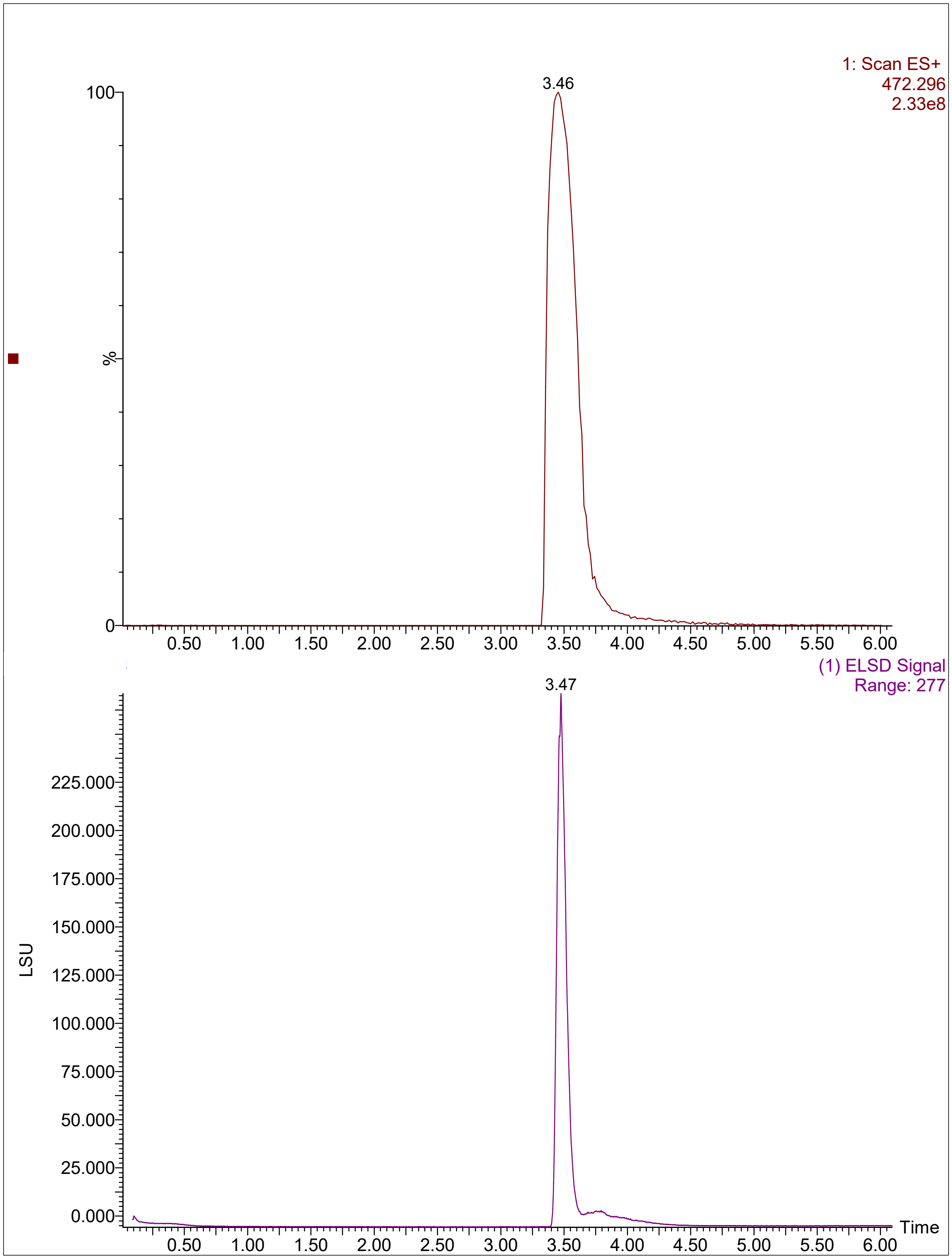

SI-Pep-2

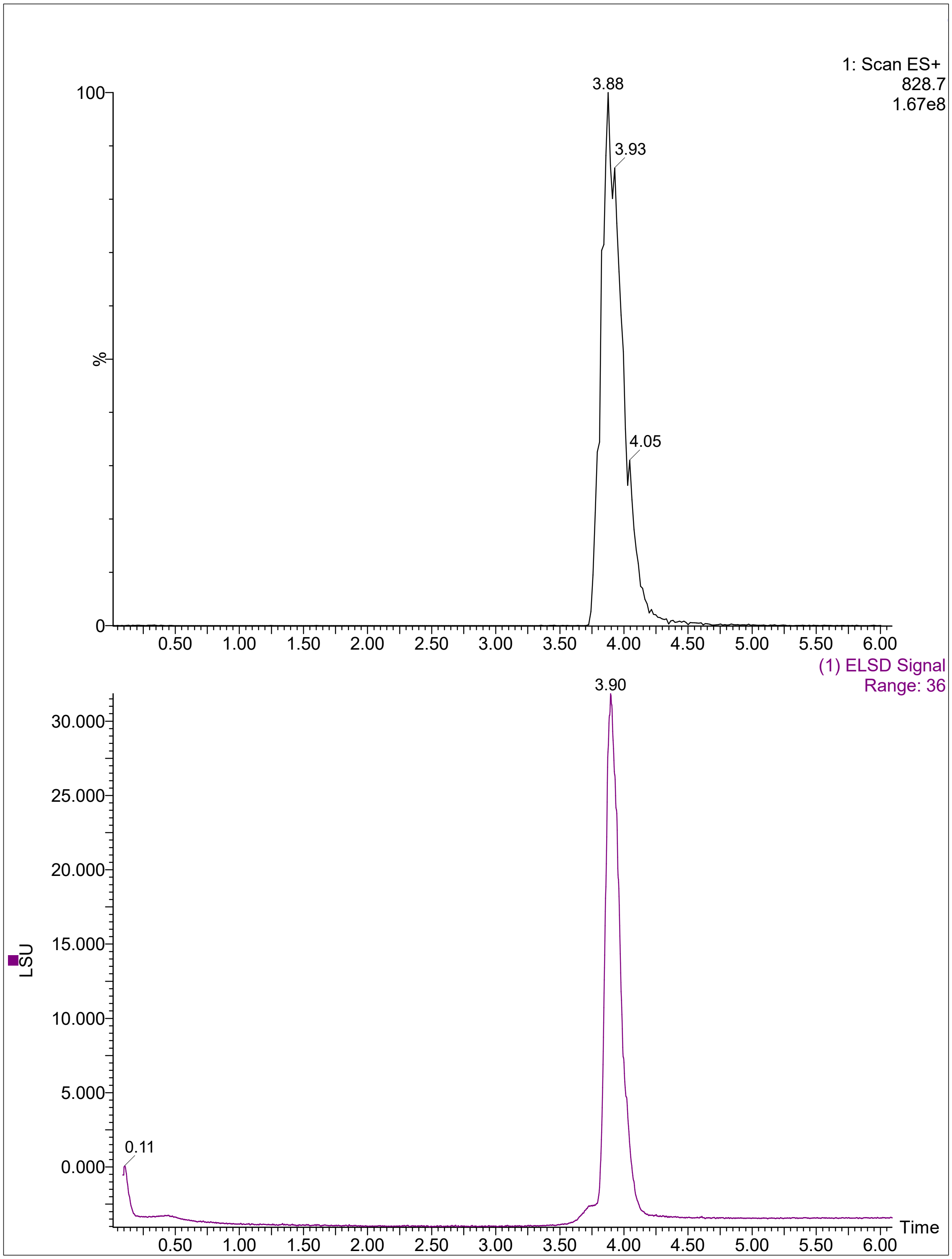

1: Scan ES+
828.7
1.67e8
3.88
3.93
4.05
100
%
0
0.50
1.00
1.50
2.00
2.50
3.00
3.50
4.00
4.50
5.00
5.50
6.00
(1) ELSD Signal
Range: 36
3.90
30.000
25.000
20.000
15.000
10.000
5.000
0.000
0.11
LSU
Time

SI-Pep-3

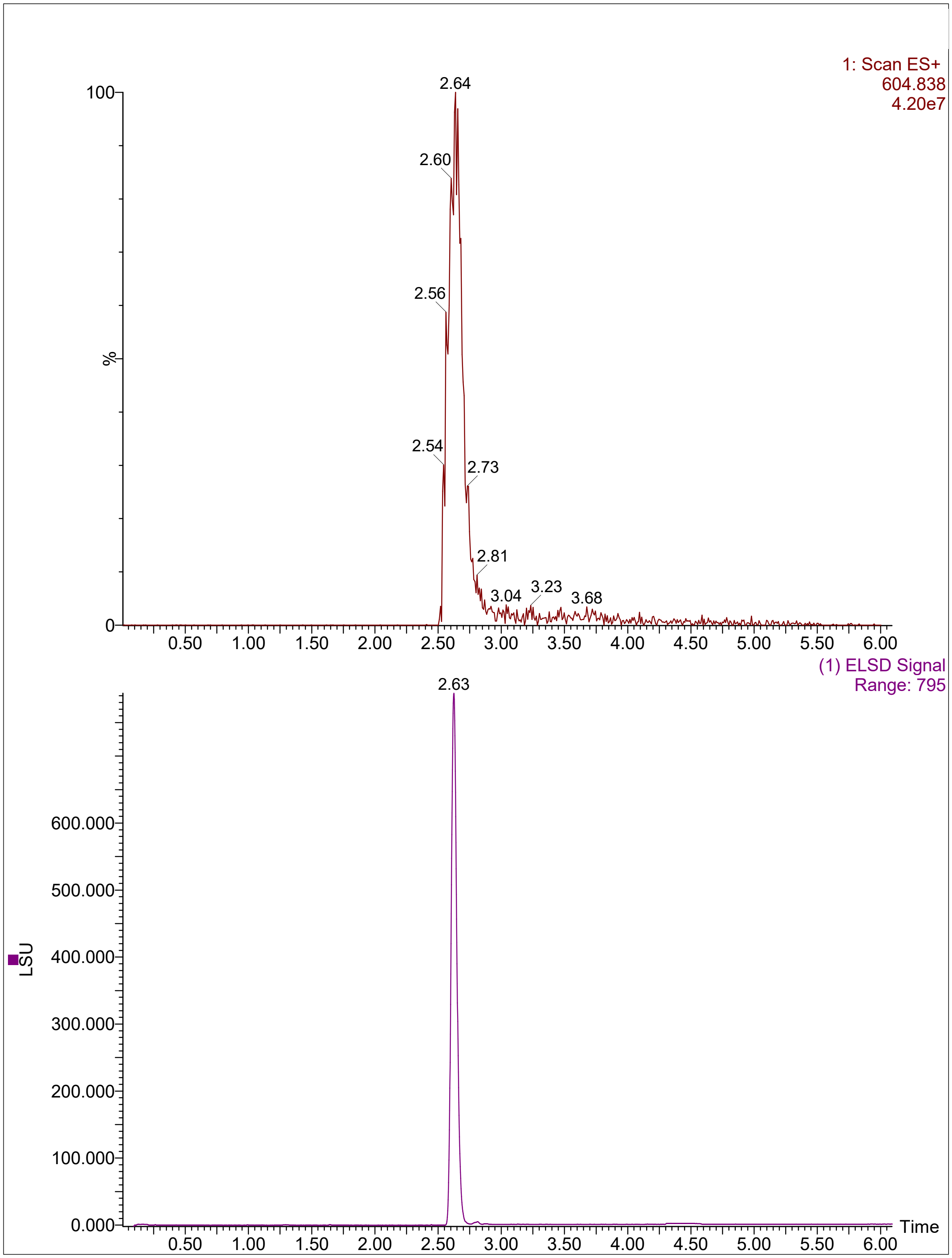
1: Scan ES+
604.838
4.20e7
100
%
0
2.64
2.60
2.56
2.54
2.73
2.81
3.04
3.23
3.68
0.50 1.00 1.50 2.00 2.50 3.00 3.50 4.00 4.50 5.00 5.50 6.00
(1) ELSD Signal
Range: 795
2.63
600.000
500.000
400.000
300.000
200.000
100.000
0.000
LSU
Time
0.50 1.00 1.50 2.00 2.50 3.00 3.50 4.00 4.50 5.00 5.50 6.00

SI-Pep-4

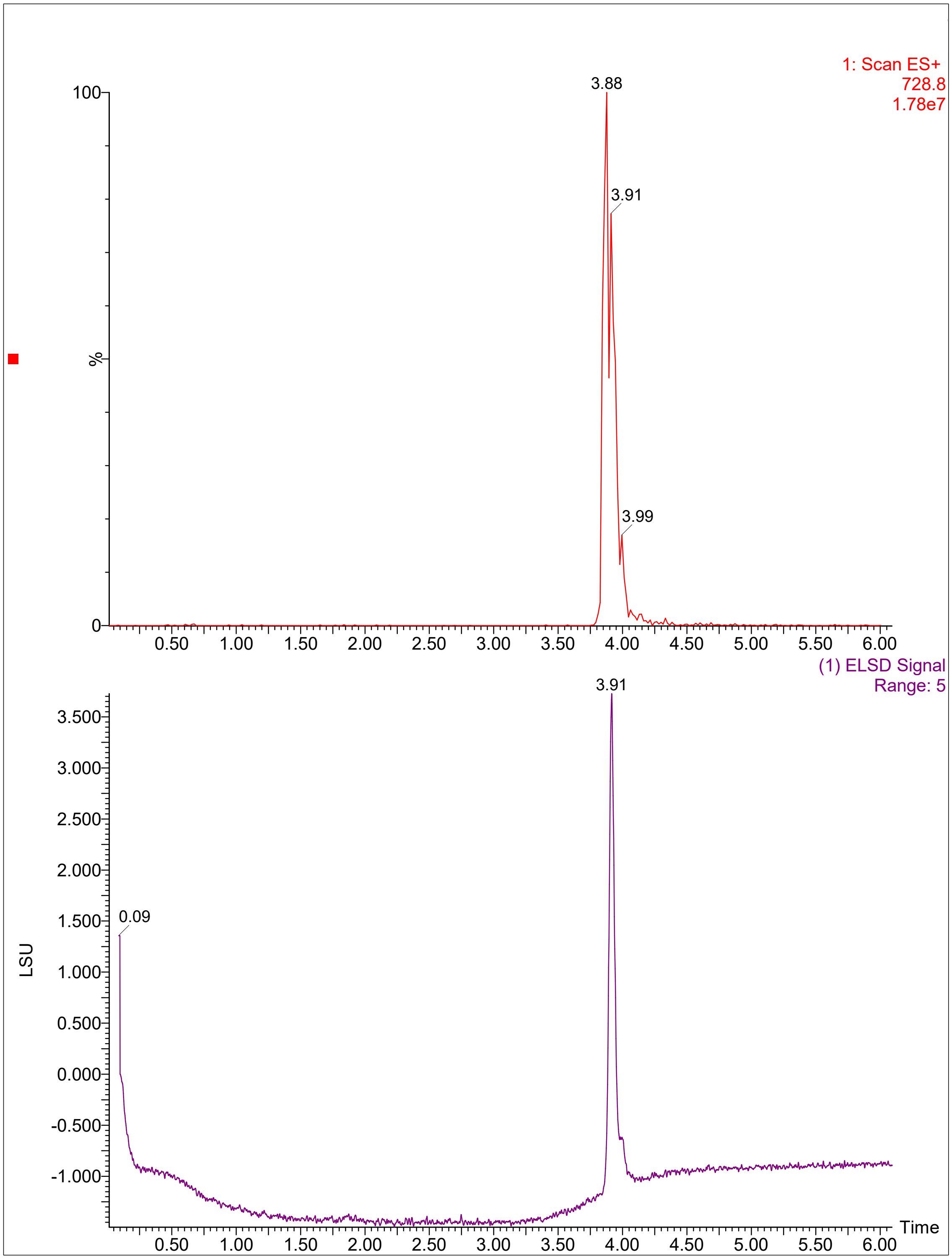
1: Scan ES+
728.8
1.78e7
3.88
3.91
3.99
%
100
0
0.50 1.00 1.50 2.00 2.50 3.00 3.50 4.00 4.50 5.00 5.50 6.00
(1) ELSD Signal
Range: 5
3.91
0.09
LSU
3.500
3.000
2.500
2.000
1.500
1.000
0.500
0.000
-0.500
-1.000
Time

SI-Pep-5

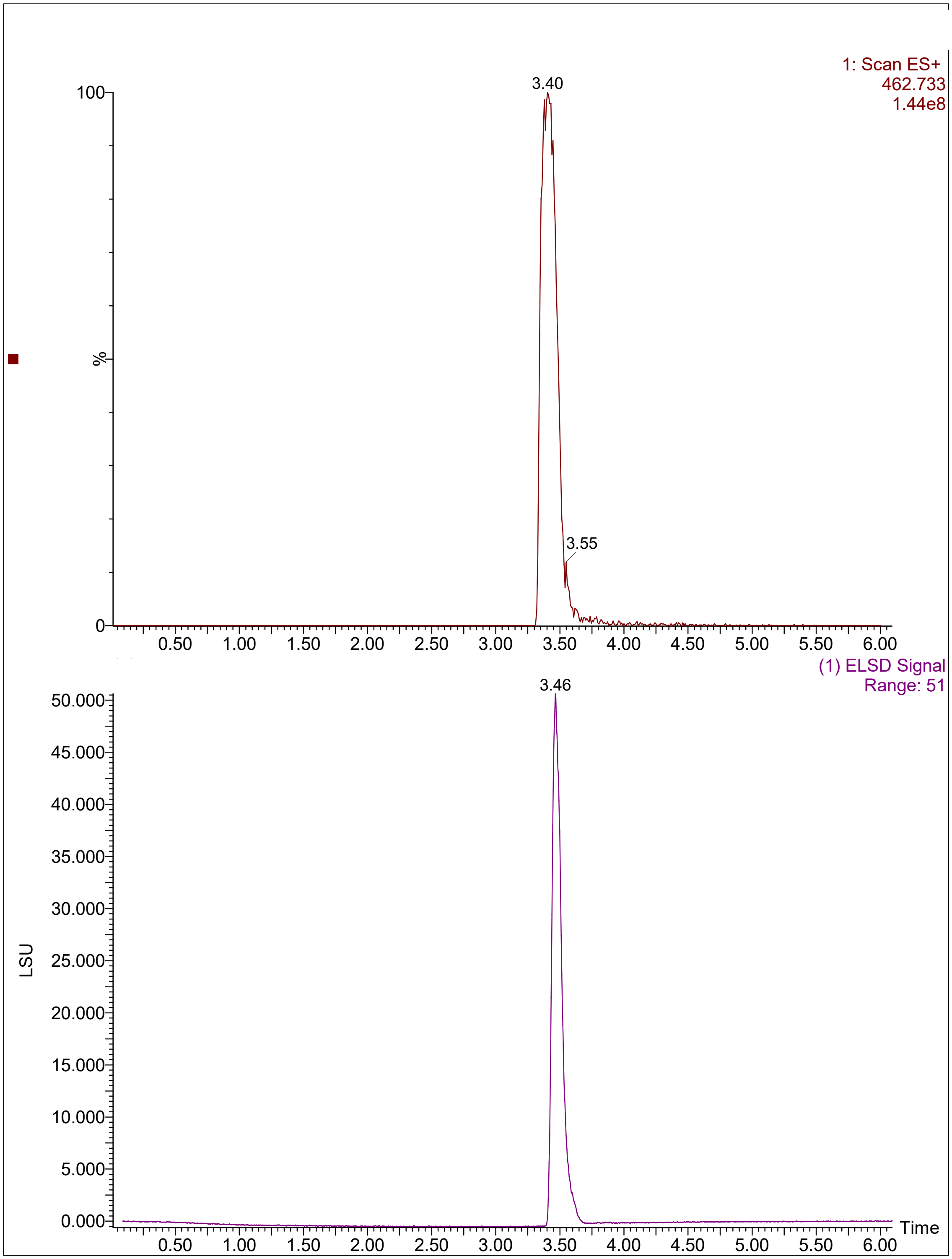
1: Scan ES+
462.733
1.44e8
3.40
3.55
100
%
0
0.50
1.00
1.50
2.00
2.50
3.00
3.50
4.00
4.50
5.00
5.50
6.00
(1) ELSD Signal
Range: 51
3.46
50.000
45.000
40.000
35.000
30.000
25.000
20.000
15.000
10.000
5.000
0.000
LSU
Time

SI-Pep-6

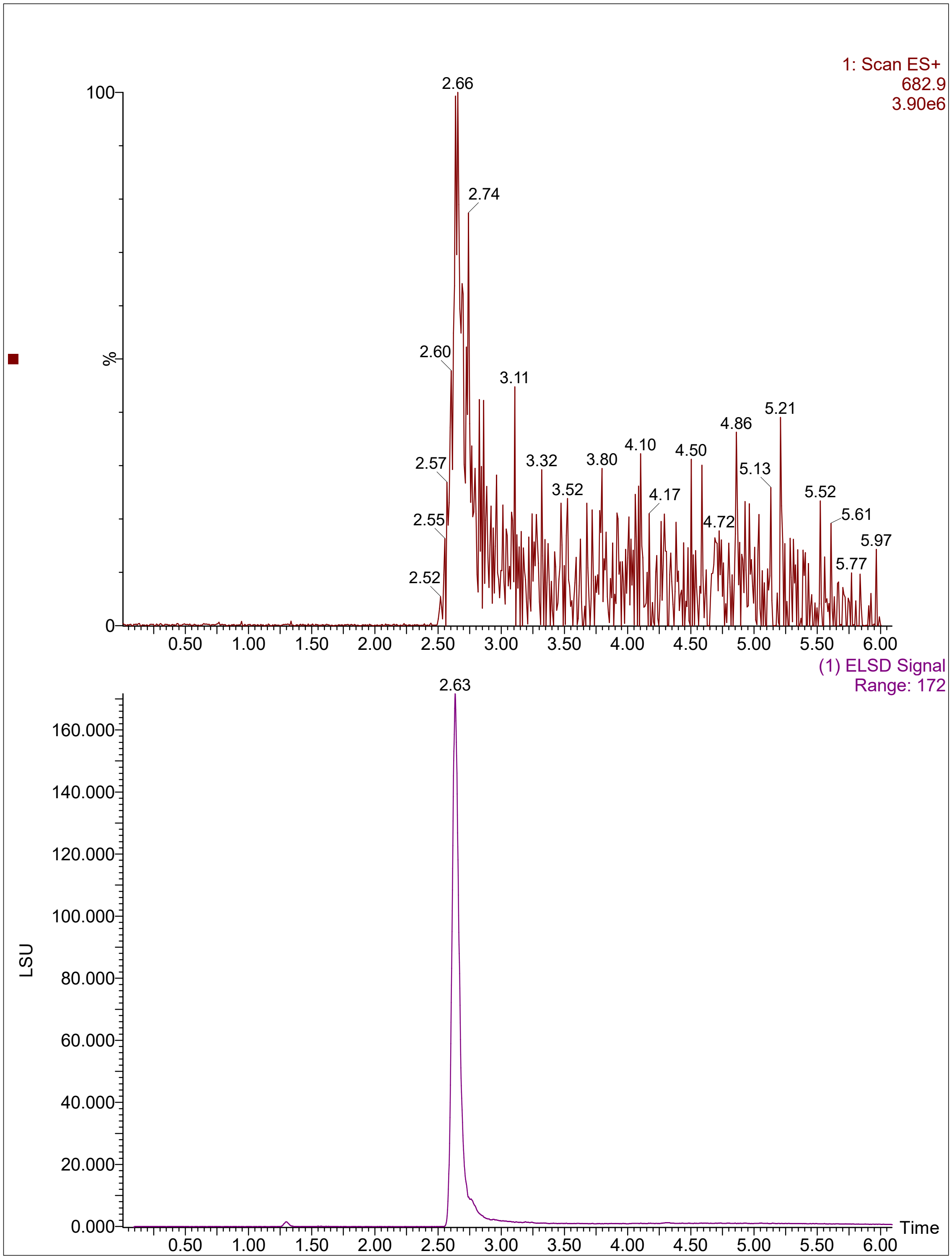
1: Scan ES+
682.9
3.90e6
2.66
2.74
2.60
3.11
2.57
3.32
3.52
3.80
4.10
4.17
4.50
4.72
4.86
5.13
5.21
5.52
5.61
5.77
5.97
2.55
2.52
100
%
0
0.50
1.00
1.50
2.00
2.50
3.00
3.50
4.00
4.50
5.00
5.50
6.00
(1) ELSD Signal
Range: 172
2.63
LSU
160.000
140.000
120.000
100.000
80.000
60.000
40.000
20.000
0.000
Time

SI-Pep-7

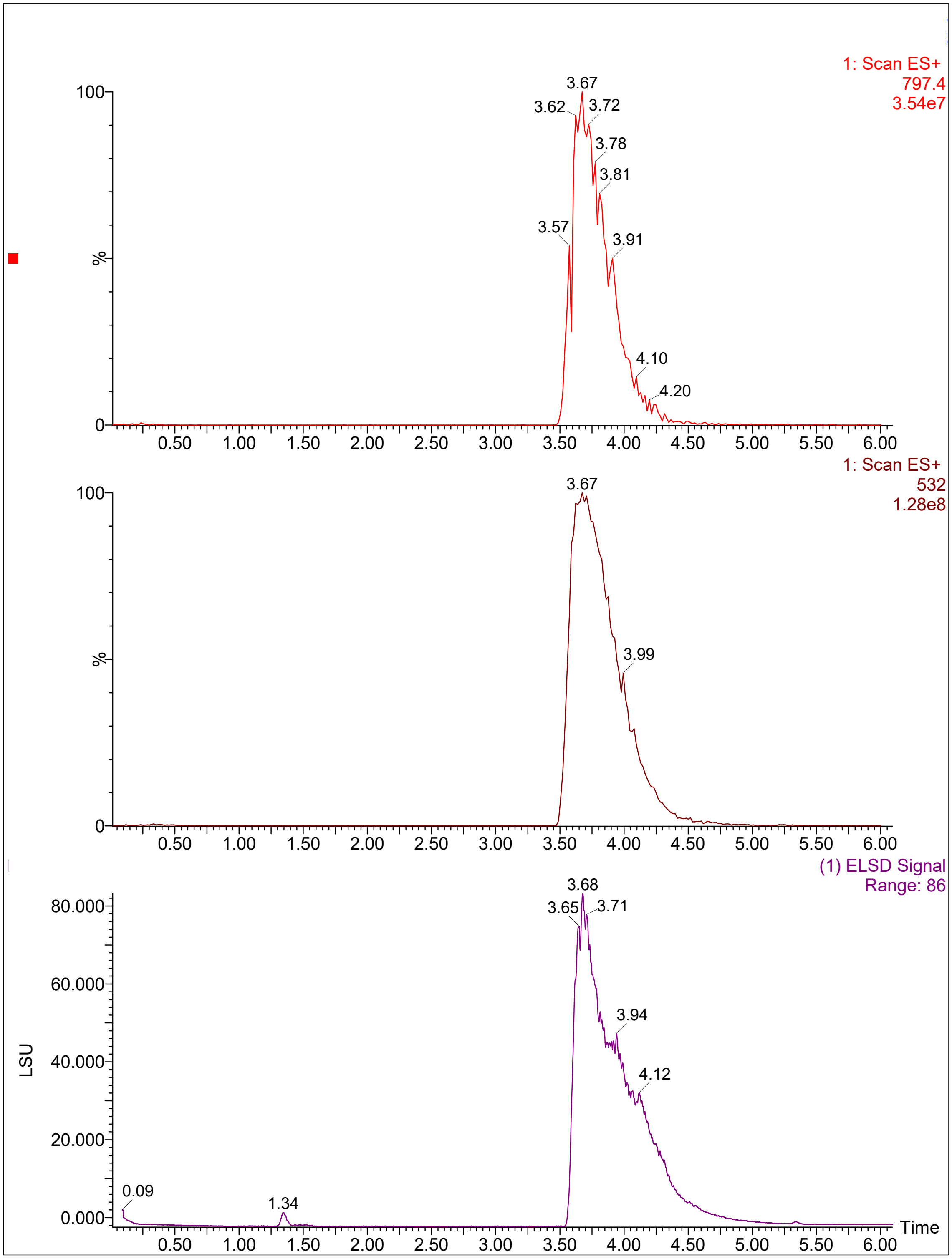

1: Scan ES+
797.4
3.54e7
3.67
3.62
3.72
3.78
3.81
3.57
3.91
4.10
4.20
100
%
0
0.50
1.00
1.50
2.00
2.50
3.00
3.50
4.00
4.50
5.00
5.50
6.00
1: Scan ES+
532
1.28e8
3.67
3.99
(1) ELSD Signal
Range: 86
3.68
3.65
3.71
3.94
4.12
0.09
1.34
80.000
60.000
40.000
20.000
0.000
LSU
Time

SI-Pep-8

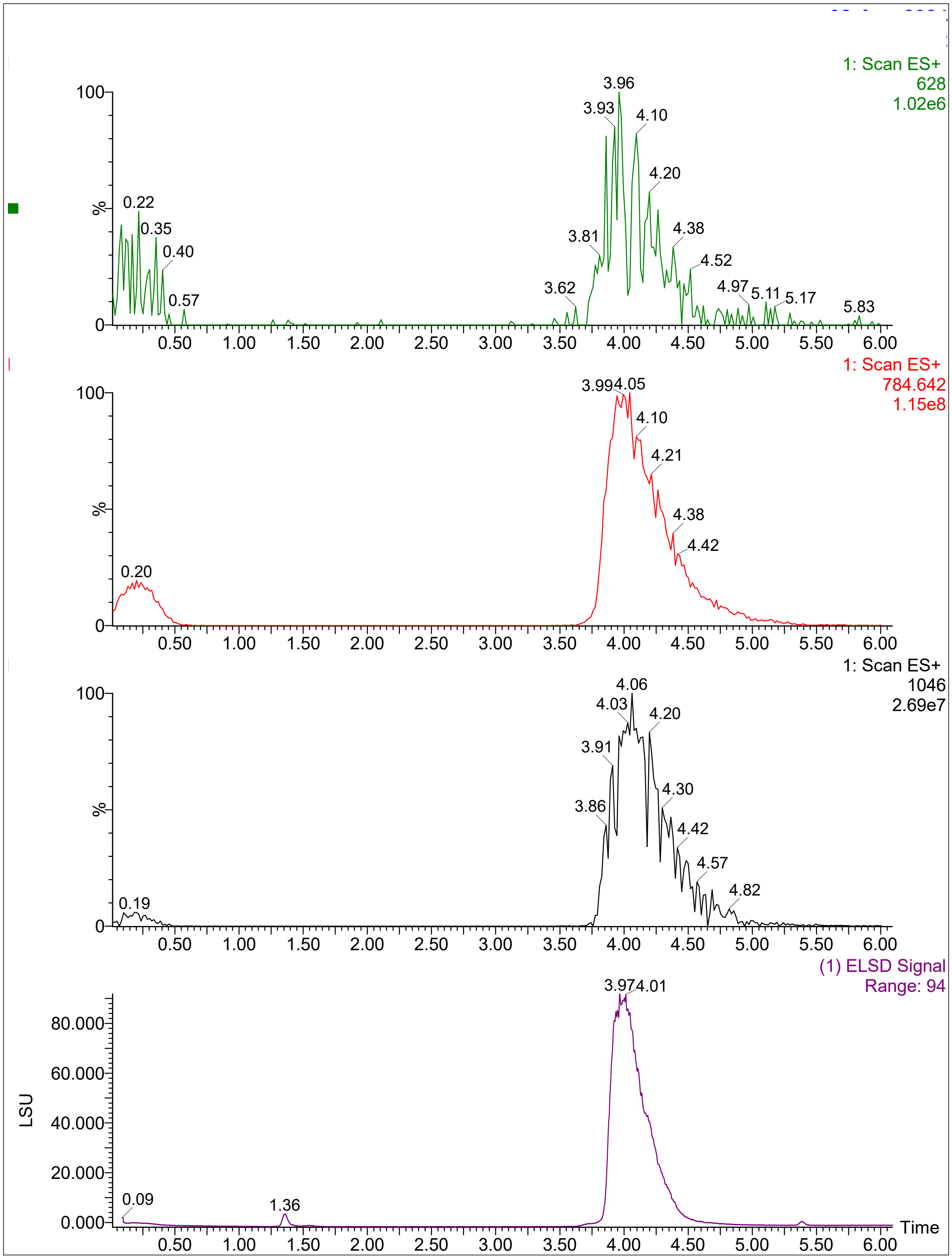
1: Scan ES+
628
1.02e6
3.96
3.93
4.10
4.20
0.22
0.35
0.40
3.81
4.38
4.52
3.62
4.97
5.11
5.17
0.57
5.83
1: Scan ES+
784.642
1.15e8
3.994.05
4.10
4.21
4.38
4.42
0.20
1: Scan ES+
1046
2.69e7
4.06
4.03
4.20
3.91
4.30
3.86
4.42
4.57
4.82
0.19
(1) ELSD Signal
Range: 94
3.974.01
LSU
0.09
1.36
Time
100
%
0
0.000
20.000
40.000
60.000
80.000
0.50 1.00 1.50 2.00 2.50 3.00 3.50 4.00 4.50 5.00 5.50 6.00

SI-Pep-9

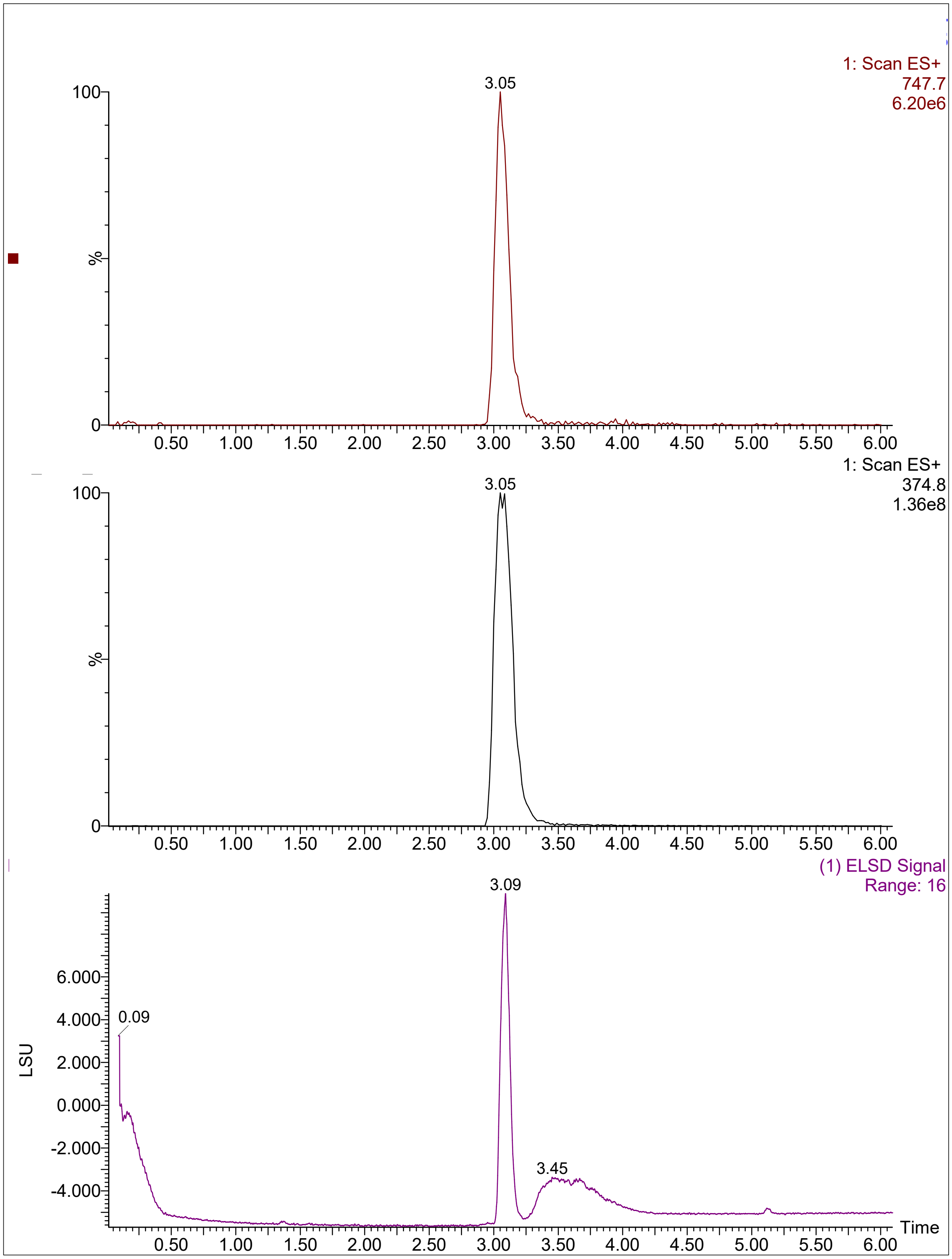

1: Scan ES+
747.7
6.20e6
3.05
100
%
0
0.50
1.00
1.50
2.00
2.50
3.00
3.50
4.00
4.50
5.00
5.50
6.00
1: Scan ES+
374.8
1.36e8
3.05
100
%
0
(1) ELSD Signal
Range: 16
3.09
6.000
4.000
2.000
0.000
-2.000
-4.000
LSU
0.09
3.45
Time

SI-Pep-10

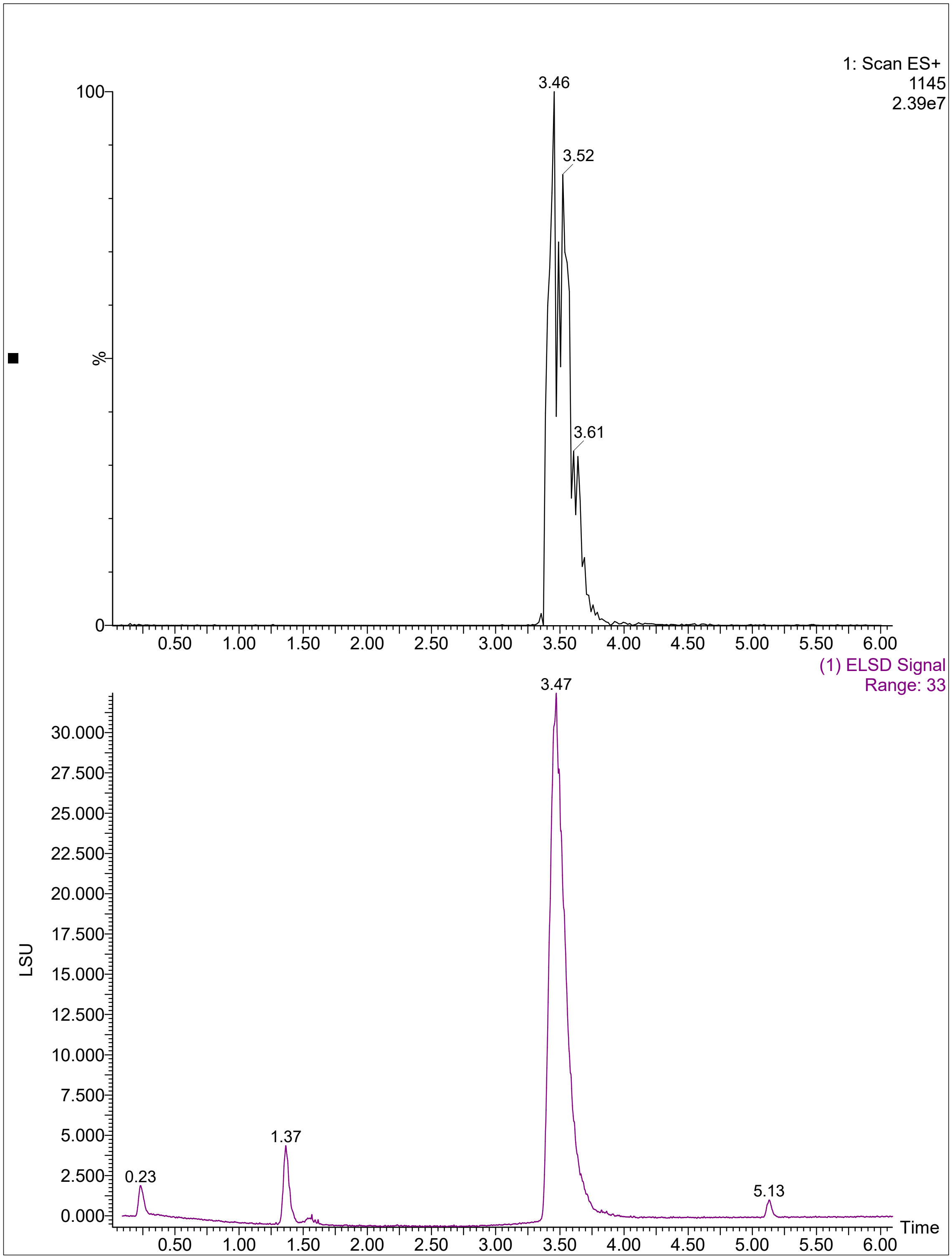
1: Scan ES+
1145
2.39e7
100
%
0
3.46
3.52
3.61
0.50
1.00
1.50
2.00
2.50
3.00
3.50
4.00
4.50
5.00
5.50
6.00
(1) ELSD Signal
Range: 33
3.47
LSU
30.000
27.500
25.000
22.500
20.000
17.500
15.000
12.500
10.000
7.500
5.000
2.500
0.000
0.23
1.37
5.13
Time
